# HILIC-Enabled Mass Spectrometric Discovery of Novel Endogenous and Glycosylated Neuropeptides in the American Lobster Nervous System

**DOI:** 10.1101/2025.06.26.661634

**Authors:** Vu Ngoc Huong Tran, Gaoyuan Lu, Thao Duong, Angel E. Ibarra, Feixuan Wu, Margot Beaver, Lingjun Li

## Abstract

Neuropeptides are a highly conserved and diverse class of intercellular signaling molecules that regulate a broad range of neural and hormonal processes across animal phyla. The American lobster, *Homarus americanus*, has long served as a powerful invertebrate model for the discovery and functional investigation of neuropeptides. Among common post-translational modifications (PTMs) found in neuropeptides, glycosylation remains underexplored due to the inherently low *in vivo* abundance and intrinsically complex structural heterogeneity. In this study, we employed hydrophilic interaction liquid chromatography (HILIC) enrichment coupled with oxonium-ion triggered EThcD fragmentation strategy to simultaneously profile novel endogenous and glycosylated neuropeptides across eight distinct neural tissues and neuroendocrine organs of *Homarus americanus*. This integrative mass spectrometry-based approach led to the identification of 154 endogenous neuropeptides derived from 25 families, approximately one-third of which are newly reported, and uncovered 24 glycosylated neuropeptides in this species for the first time. These peptides exhibit strong tissue-specific expression, distinct proteolytic cleavage patterns, and confidently localized glycosylation sites. Our results highlight the utility of integrated sampling enrichment and hybrid fragmentation strategies for deep neuropeptidomic profiling and provide a valuable resource for future studies on the functional roles of newly identified neuropeptides and glycosylation in crustacean neuromodulation and peptidergic signaling.

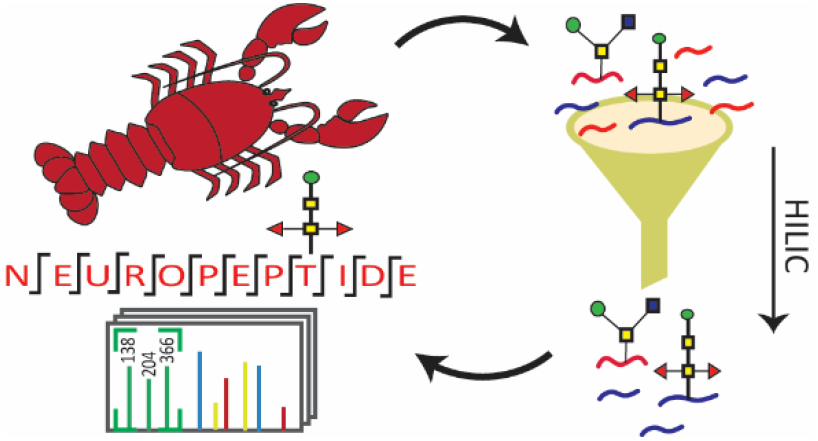

## Introduction

Neuropeptides represent the most diverse class of secreted signaling molecules that play essential roles in numerous physiological and neurological processes including appetite control, stress response, energy homeostasis, and psychostimulant addiction^1-4^. However, elucidating their precise functions in mammals remains challenging due to the intricacies of the mammalian nervous system. As an alternative, crustaceans, such as the American lobster, *Homarus americanus*, have long served as valuable model organisms in neuroscience research, owing to their relatively simple, accessible, and well-defined neural circuits. These circuits express a wide array of neuropeptides that are structurally and functionally homologous to those found in vertebrates^5^. The crustacean nervous system consists of multiple anatomically distinct yet functionally interconnected regions, including both neural tissues and neuroendocrine organs. These regions engage in reciprocal signaling, where neuropeptides act both locally within specific circuits and systemically as hormonal messengers to coordinate neuromodulation, neuronal plasticity, and physiological responses^6^. Given the dynamic interplay among these tissues, a comprehensive profiling of neuropeptides across the entire nervous system is essential to fully understand the molecular underpinnings of neurochemical signaling in crustaceans.

Neuropeptides are initially encoded in the genome as part of larger precursor proteins known as preprohormones. Following multiple proteolytic cleavages and extensive post-translational modifications (PTMs), mature bioactive neuropeptides are released, exhibiting high sequence diversity, structural similarity, and numerous PTMs. A novel PTMs found in neuropeptides is glycosylation, where a glycan (a group of monosaccharides) is attached to the side chain of the peptide backbone, capable of altering bioactivity, receptor binding, and peptide stability^7^. Glycans are commonly classified into two categories: N-linked glycosylation, where the sugar is attached to the nitrogen (N) atom in the side chain of asparagine (Asn) within the consensus motif Asn-X-Ser/Thr (with X being any amino acid except proline), and O-linked glycosylation, where the sugar is attached to the oxygen (O) atom of serine (Ser) or threonine (Thr) residues on the peptide backbone, with no strict consensus motif. Depending on their composition and branching, N-glycans can be further classified into high-mannose, hybrid, and complex types^8^. Meanwhile, the first monosaccharide of O-glycans linked to the peptide backbone is used to further classify them into sub-categories such as O-GalNAcylation (mucin-type O-glycosylation with predominantly core 1, core 2, and Tn antigen structures), O-fucosylation, O-mannosylation, or O-glucosylation^9^.

Given this complexity, mass spectrometry (MS) has emerged as a powerful platform for global neuropeptide profiling with high throughput, specificity, and sensitivity^10, 11^. Following the release of the high-quality American lobster draft genome, our lab previously employed MS-based analysis to identify 101 unique mature neuropeptides across five lobster neural tissues, laying the ground foundation for expanded neurochemical discovery^12, 13^. However, several important limitations remained. Firstly, many additional putative neuropeptides predicted from the genome-derived precursor protein database were not detected in the previous study, likely due to incomplete recovery during sample preparation and limitations of MS sensitivity. Thus, a more optimized workflow, including strategic enrichment and alternative MS acquisition method, is required to capture a broader neuropeptide repertoire from complex tissue matrices. Secondly, although glycosylation is a well-recognized and functionally significant PTM in neuropeptides, it is often overlooked in our previous publications. This is largely attributed to the inherently lower *in vivo* abundance of glycosylated species compared to their non-modified counterparts, as well as their remarkably diverse structural heterogeneity and poor ionization efficiency, which poses substantial analytical challenges.

To overcome the low *in vivo* abundance of glycosylated neuropeptides, glycopeptide enrichment strategies are essential. Affinity-based assays, such as lectin affinity chromatography, titanium dioxide, hydrazide and boronic acid chemistries are commonly employed; however, these methods often rely on specific chemical interactions between the enrichment material and particular glycan motifs, leading to enrichment biases toward certain glycan classes.^14-17^ On the other hand, hydrophilic interaction liquid chromatography (HILIC) offers an unbiased method for enriching both glycopeptides and a broad range of hydrophilic peptides based solely on their hydrophilic properties^18, 19^. Given the significant variation in hydrophobicity and glycosylation of endogenous neuropeptides, HILIC enrichment has been showed to be effective in neuropeptidomic applications^20^. Furthermore, a hallmark feature of glycopeptide spectra is the presence of oxonium ions, which correspond to low-mass glycan-derived fragments (*e*.*g*., *m/z* 204.08 and 366.14) and serve as diagnostic indicators of glycopeptide precursors. While the conventional higher-energy collision dissociation (HCD) fragmentation is effective for peptide sequence identification, the glycan moieties on glycopeptides are labile and prone to cleavage during HCD, hindering accurate determination of glycosylation sites. On the other hand, electron-transfer dissociation (ETD) offers gentler fragmentation that preserves glycan structures, though it provides limited peptide backbone fragmentation. Therefore, we employed a widely adapted oxonium ion-triggered electron-transfer/higher-energy collision dissociation (EThcD) strategy, where precursors are first subjected to HCD fragmentation, and those producing oxonium ions trigger additional ETD fragmentation^21-25^. This hybrid acquisition method combines the complementary benefits of both ETD and HCD, providing maximum fragment information in a single run. It enables the concurrent collection of HCD and EThcD spectra, with HCD scans facilitating the discovery of endogenous neuropeptides, and EThcD scans providing confident characterization of glyconeuropeptide sequences and glycosylation sites.

Leveraging advanced sample preparation and hybrid fragmentation strategies, this study aimed to simultaneously discover novel endogenous neuropeptides and, for the first time, characterize the glycosylated neuropeptidome of the American lobster nervous system. Our findings offer a deeper coverage of the American lobster neurochemical landscape and significantly expand the repertoire of bioactive neuropeptide candidates. This work lays the foundation for future functional studies investigating the roles of newly identified neuropeptides and glycosylation in crustacean neuromodulation and peptidergic signaling.

## Materials and Methods

### Chemicals

Unless specified otherwise, all solvents and reagents were obtained from Thermo Fisher Scientific (Pittsburgh, PA) or MilliporeSigma (St. Louis, MO). Poly HYDROXYETHYL A bulk material was acquired from PolyLC (Columbia, MD).

### Animals and Tissue Collection

American lobsters, *Homarus americanus*, were purchased from Global Market and Food Hall (Madison, WI) and allowed to acclimate to artificial seawater tanks (made with Instant Ocean Sea Salt, 10– 12°C, alternating 12-hr light/dark cycle) for at least 2 weeks before use. The lobsters were housed, treated, and sacrificed in accordance with the animal care and study protocol approved by the UW-Madison Animal Care and Use Committee. Lobsters were cold anesthetized on ice for 30 min prior to the dissection conducted in chilled physiological saline solution (479 mM NaCl; 12.74 mM KCl; 20 mM MgSO_4_, 3.91 mM Na_2_SO_4_, 13.67 mM CaCl_2_, 5 mM HEPES, pH 7.45). Eight neural tissues, including a pair of sinus glands (SG), the brain, the oesophageal ganglion (OG), a pair of commissural ganglia (CoG), the stomatogastric ganglion (STG), the thoracic ganglion (TG), a pair of pericardial organs (PO), and the cardiac ganglion (CG), were collected and immediately placed in 200 µL ice-cold acidified methanol (90% methanol, 9% water, and 1% glacial acetic acid), snap-frozen on dry ice, and stored at -80°C until needed. Due to the limited amount of the endogenous glycosylated neuropeptide per tissue, samples were pooled from 10 to 20 lobsters to obtain 100 – 300 *µ*g of starting material per analysis.

### Neuropeptide Extraction and HILIC Enrichment

Tissues were homogenized in 200 *µ*L acidified methanol using a Fisher Scientific sonic dismembrator (Pittsburgh, PA) set to 50% amplitude in an ice-water bath. Sonication was performed in 8-second on / 15-second off pulses for 7 to 12 cycles, adjusted based on tissue size and consistency. The homogenized samples were centrifuged at 20,000 rcf at 4°C for 40 min. The supernatant was collected and pooled, and an aliquot (approximately 10%) was reserved as the non-enriched sample. All samples were dried using a SpeedVac concentrator and stored at -80°C for subsequent HILIC enrichment.

The spin-tip cotton HILIC enrichment method was conducted as previously described^20^. Briefly, dried samples were reconstituted in 100 *µ*L of 95% ACN containing 1% TFA, and peptide concentration was determined using a Thermo Scientific NanoDrop One microvolume spectrophotometer. HILIC beads were weighed at a bead-to-peptide mass ratio of 90:1 and activated in 60 mg/mL 1% TFA for 15 min prior to use. HILIC beads were loaded on top of 3 mg of cotton wool in a pipette tip, flushed, and conditioned with 95% ACN/1% TFA solution. Peptide samples were then loaded and passed through multiple times to ensure binding. After washing several times with 95% ACN/1% TFA solution, bound peptides were eluted in a stepwise manner using 50% and 10% ACN containing 5% FA solutions. Enriched samples were further dried, reconstituted in 30 µL of 0.1% FA, and desalted by C18 ZipTip pipette tips following the manufacturer’s instructions. The desalted samples were then resuspended in 12 µL of 0.1% formic acid in water, transferred into MS sample vials, and peptide concentrations were measured (**Figure S1**). A 2 µL aliquot was injected into the mass spectrometer for analysis.

### LC-MS/MS Analysis

Samples were analyzed on the Thermo Orbitrap Fusion Lumos Tribrid mass spectrometer coupled with a Dionex UltiMate 3000 UPLC system. A self-packed Waters BEH C18 particles 19-cm microcapillary column was used for a 136-minute chromatographic separation of mobile phase A (0.1% FA in water), and mobile phase B (0.1% FA in 80% acetonitrile) as follows: 0−18 min 3% B; 18−80 min 3−35% B; 80−100 min 35−80% B; 100−105 min 80% B; 105−110 min 80-100% B; 110−120 min 100% B; 120−120.5 min 100–3% B; 120.5–136 min 3% B. The flow rate was maintained at 0.3 *µ*L/min. For pd-EThcD analysis, the MS scan range was 400 to 2000 *m/z* at a resolution of 60 000, RF Lens of 30%, a standard AGC target, and a maximum injection time of 50 ms. Precursors were dynamically excluded for 20 s with a 10-ppm tolerance. A cursory HCD scan was performed to fragment the most abundant precursor ions between 2+ to 7+ charge state with a normalized collision energy of 35%, a resolution of 30 000, a dynamic exclusion of 20 s, a normalized AGC target of 60%, and a maximum injection time of 60 ms. When oxonium ions at *m/z* 138.0545, *m/z* 204.0867, or *m/z* 366.1396 (± *m/z* 0.01) were detected within the top 30 most abundant peaks, pd-EThcD MS/MS fragmentation would be triggered and further applied to the same precursor in the Orbitrap at a resolution of 60 000, a subsequent supplemental activation energy of 33%, a normalized AGC target of 500%, and a maximum injection time of 250 ms. For pd-EThcD scans, a decision tree was used to subject precursors of different charges to different ETD reaction times (z = 2 – 4, 30 ms; z = 5 – 8, 15 ms).

### Database Search and Data Analysis

The MS raw data were searched by PEAKS Xpro Studio and PEAKS GlycanFinder v.2.5. (Bioinformatics Solutions Inc.) against a manual curated neuropeptide precursor database filtered from the Uniprot genome-derived *Homarus americanus* proteome database (Feb 12^th^, 2023). The built-in 1867 basic N-glycan and 265 basic O-glycans databases in PEAKS GlycanFinder were employed to search for glycosylated neuropeptides. Database search parameter settings are listed as follows: 10 ppm parent mass error tolerance, 0.02 Da fragment mass error tolerance, 20 ppm glycan fragment mass tolerance, and an unspecific digest mode. Each peptide can have a maximum of 3 variable PTMs including oxidation on the methionine, pyro glutamylation on the N-term glutamic acid and glutamine, and amidation on the C-termini. The identified peptides were further filtered by these criteria: Proteins -10lgP ≥ 0, ≥1 unique peptide, and with significant peptides; Peptides -10lgP ≥ 15. Peptides were considered positively identified by manually reviewing the MS/MS fragmentation, the presence of oxonium ions in MS/MS spectra, protein and peptide scores, sequence coverage, and mass accuracy. Data were analyzed and visualized using Microsoft Excel and in-house-built Python scripts.

## Results

### Optimized Workflow for Enhanced Coverage of the *Homarus americanus* Neuropeptidome

To comprehensively characterize the dynamic neurochemical landscape of the American lobster peptidergic signaling system, we collected and analyzed eight distinct neural tissues (**Figure 1A**). These included the central nervous system (CNS) (*i*.*e*., brain), the stomatogastric nervous system (STNS) (*i*.*e*., STG, OG, and CoG), the CG, the TG, and two key neuroendocrine organs: the SG in the eyestalk and the PO adjacent to the heart. The American lobster features several well-defined central pattern generator (CPG) circuits, whose activity are modulated by neuropeptides and peptide hormones in response to physiological conditions and environmental stimuli^26^. For example, the CNS integrates sensory input and initiates command signals; the STNS controls rhythmic chewing and pyloric movements through its gastric mill and pyloric CPGs; the CG regulates rhythmic heart contractions; and the TG governs claw movement and associated motor output^26, 27^. Moreover, the SG and PO serve as neuroendocrine sites that synthesize and secrete neuropeptides into the circulating fluid (*i*.*e*., hemolymph) to reach distant organs, influencing both local and downstream physiological processes. Altogether, profiling neuropeptides across these interconnected tissues provides critical insights into their sites of origin and modes of action, whether they act locally within neural circuits or systemically as hormones, thereby enhancing our understanding of neuromodulation in crustaceans.

**Figure 1.**
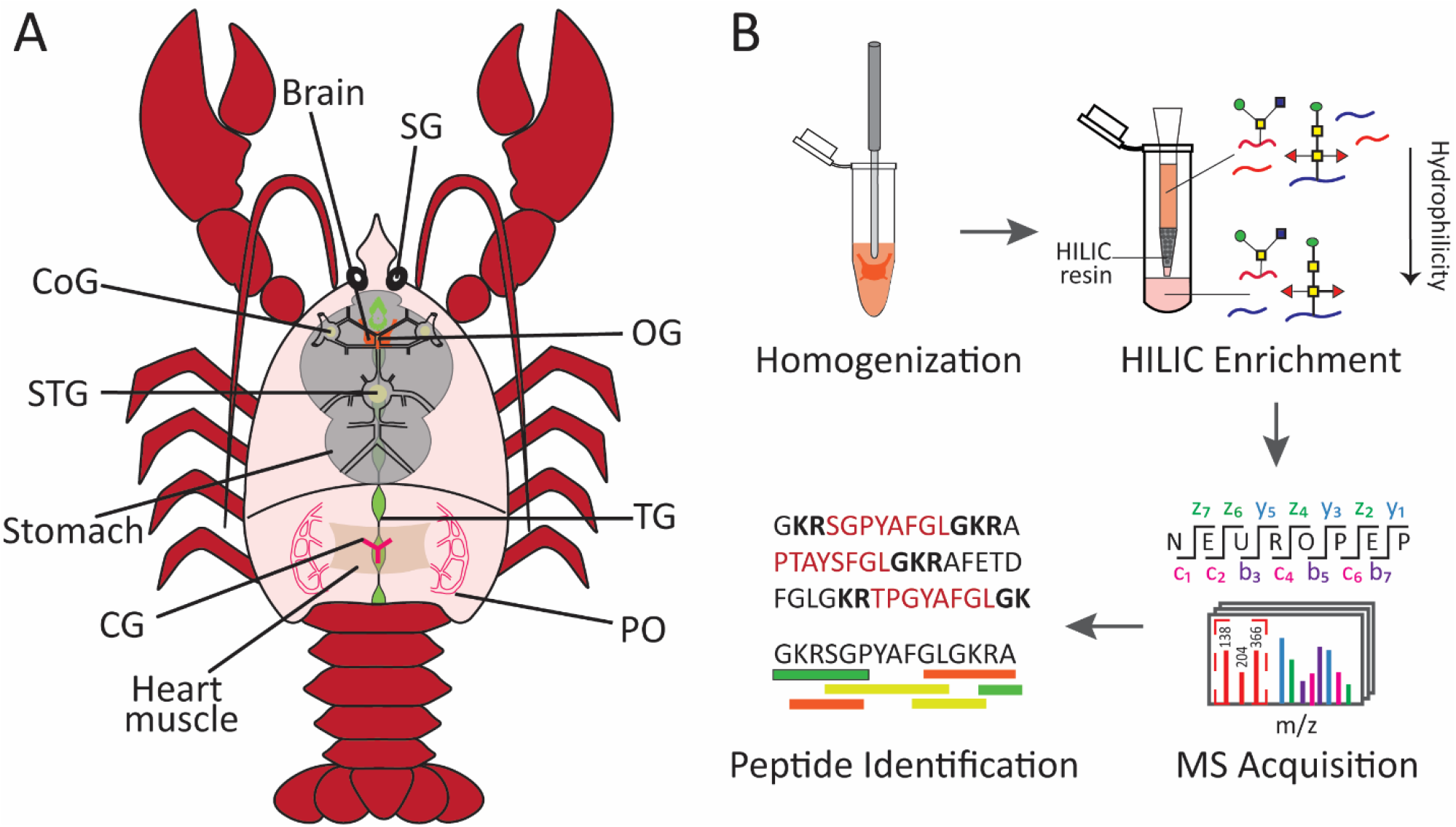
(A) General anatomy of an American lobster *Homarus americanus*, with key neural tissues and neuroendocrine organs highlighted, including the brain, sinus glands (SG), oesophageal ganglion (OG), commissural ganglia (CoG), stomatogastric ganglion (STG), cardiac ganglion (CG), thoracic ganglion (TG), and pericardial organs (PO). (B) Schematic overview of the experimental workflow for extraction, HILIC enrichment, and MS-based analysis of endogenous (glyco)neuropeptides from lobster neural tissues.

However, recovering low-abundance neuropeptides from complex tissue extracts is particularly challenging, necessitating an enrichment step in the traditional crustacean neuropeptidomic workflow to improve neuropeptidome coverage (**Figure 1B**)^10^. Briefly described, lobster tissues were dissected, immediately snap-frozen in acidified methanol to inactivate proteases that could degrade or modify neuropeptides, and thoroughly homogenized. A HILIC enrichment step was incorporated to clean-up and selectively enrich glycosylated and endogenous neuropeptides. Because HILIC involved multiple wash steps that could result in significant sample loss, a relatively large amount of starting material was required. Lobster tissues were pooled according to their relative size and neuropeptide content, with 10 lobsters pooled per sample for brain, TG, CG, paired CoGs, SGs, POs, and 20 lobsters for smaller STG and OG tissues, yielding 100–300 µg material per tissue. Unlike tryptic peptides, neuropeptides vary in length ranging from 4 to more than 80 amino acids and contain a higher proportion of basic residues, thus requiring conventional HILIC protocols for tryptic glycopeptides to be optimized. Increasing the organic composition (*e*.*g*., acetonitrile) in reconstitution and wash buffers from 80% to 95% significantly improved neuropeptide retention on HILIC materials^20^. Utilizing this optimized workflow, HILIC enrichment substantially improved neuropeptide recovery, yielding 20% to 50% more neuropeptides than non-enriched samples across all eight neural tissues (**Figure 2A**). Notably, the number of glycosylated neuropeptides identified from HILIC-enriched samples enhanced sevenfold compared to non-enriched samples (**Figure S2**).

**Figure 2.**
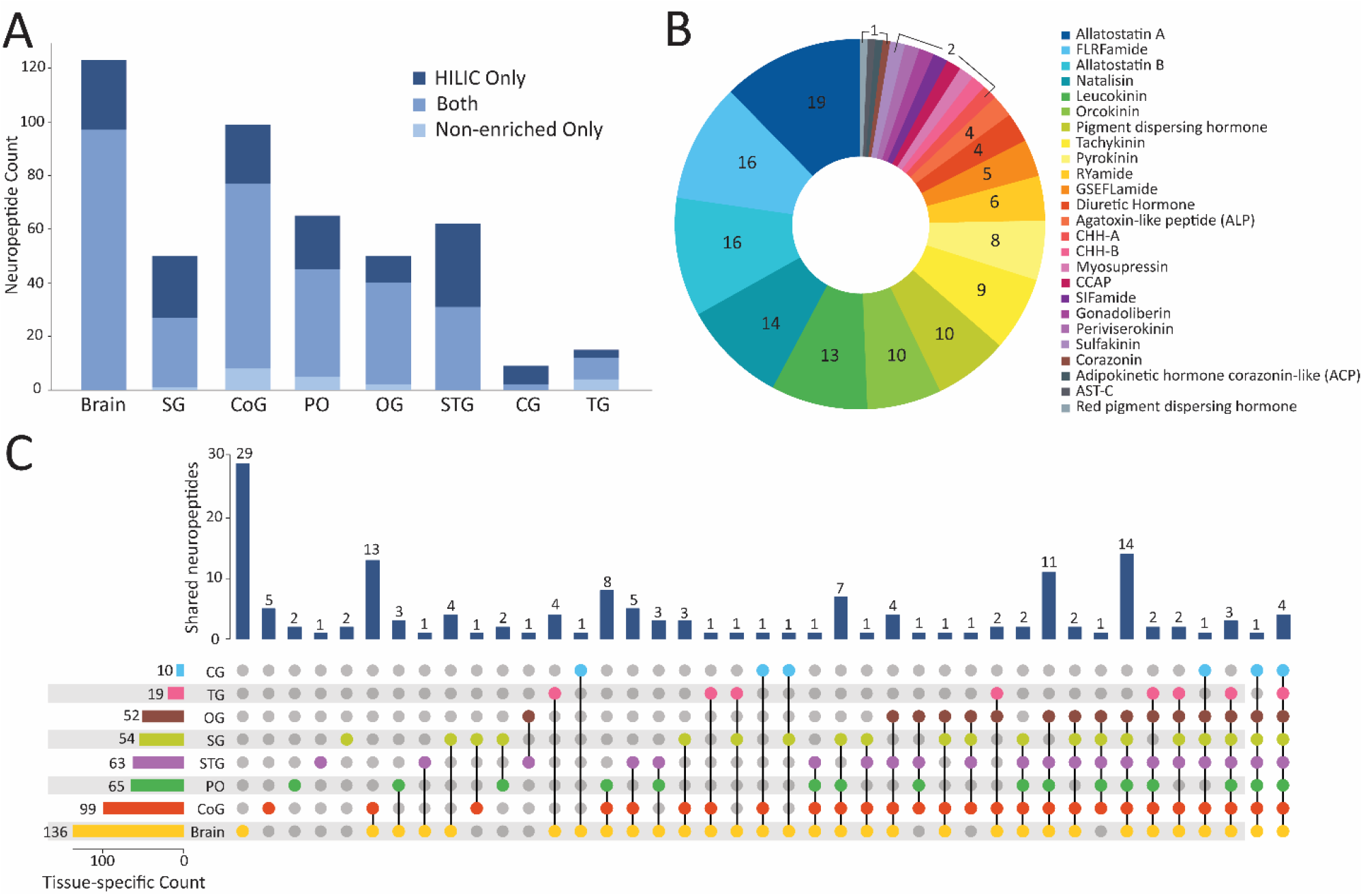
(A) Stacked bar chart showing the number of neuropeptides detected in HILIC *versus* Non-enriched samples across eight neural tissues. (B) Pie chart depicting the distribution of neuropeptide families identified in the American lobster nervous system, grouped by their corresponding prohormone origin. (C) Upset plot illustrating the tissue-specific distribution of neuropeptides among the eight neural tissues of the American lobster. The left bar chart indicates the total number of neuropeptides detected per tissue, while the top bar chart represents the number of shared neuropeptides across tissues connected by their respective dots.

To strengthen the software’s search capability of identifying endogenous peptides, we utilized a manually curated database of 87 known or predicted neuropeptide precursor sequences rather than the large Uniprot lobster proteome database comprising over 24,000 protein entries. This refined database search strategy stemmed from the inherent limitations associated with large-scale proteome database searching in peptidomics workflows, which substantially increase search space and statistical correction required during false discovery rate (FDR) estimation^28, 29^. As a result, low-abundance, short, post-translationally modified neuropeptides may be overlooked or scored below significance thresholds. By employing a curated neuropeptide-specific database, we effectively reduced search complexity, improved spectral matching sensitivity, and increased neuropeptidome coverage. We applied a peptide confidence threshold of -10lgP ≥ 15, consistent with software recommendations, and further validated each assigned spectrum through manual inspection of fragmentation patterns. A key cleavage pattern at dibasic residues (lysine and arginine) or monobasic (lysine) residue, with occasional cleavages at glutamate, leucine, and serine residues, typically occurs at both terminals of the mature bioactive neuropeptide^30, 31^. Common PTMs, such as N-terminal pyroglutamation and C-terminal amidation followed by glycine residue, can also serve as reliable indicators of bioactive peptides^32^. Given these cleavage and modification patterns, we meticulously reviewed and documented the mature neuropeptides for inclusion in **Table S1** and their annotated MS/MS spectra were shown in **Figure S5**. Additionally, a careful inspection of oxonium ions in the lower *m/z* range, a strong indicator of glycopeptide spectra, enabled the identification of glyconeuropeptides with their corresponding MS/MS spectra (**Figure S6**).

In our study, peptides cleaved from the same protein precursor were broadly classified as members of the corresponding neuropeptide family. Overall, we identified 154 neuropeptides derived from 25 distinct neuropeptide families, with their distribution shown in **Figure 2B**, highlighting the diversity of crustacean neuropeptides involved in various physiological processes. This list includes highly conserved neuropeptide families, such as allatostatin, tachykinin, and FLRFamide, which are commonly found across animal phyla^5^. The properties of endogenous neuropeptides in the American lobster nervous system were further analyzed (**Figure S3**). Peptide lengths predominantly ranged from 8 to 15 amino acids, with some comprising up to 40 residues. This distribution aligns with the MS-based peptidomics capability, which is most effective at detecting peptides within the 5-25 amino acid range. Very short peptides (3-5 amino acids) may fail to generate sufficient fragmentation, while longer peptides may ionize poorly or undergo incomplete fragmentation. Interestingly, longer peptides were mostly detected from hormone families (*e*.*g*, CHH, DH, and PDH), likely reflecting their hormonal roles where their size may provide greater stability and facilitate their transport through the circulatory system. Furthermore, the grand average of hydropathy (GRAVY) scores, primarily clustered around -0.5, indicate that most neuropeptides are relatively hydrophilic, which highlights the effectiveness of HILIC in recovering a broad range of neuropeptides, particularly those with hydrophilic properties. Additionally, the isoelectric point (pI) distribution revealed a bimodal pattern, with an enrichment of acidic and basic peptides and a noticeable depletion of neutral ones. Peptides near neutral pI are less soluble, poorly retained in HILIC, and less likely to ionize efficiently in positive mode electrospray ionization (ESI) due to their lower net charge, further reducing their detectability in MS analysis.

We conducted a tissue-specific distribution analysis of the identified neuropeptides across all eight tissues, as illustrated in **Figure 2C**. Overall, the brain exhibited the highest number of unique neuropeptides (136), followed by CoG and PO with 98 and 65 neuropeptides, respectively. Fewer neuropeptides were detected in the CG and TG, likely due to their higher lipid content and lower neuropeptide abundance. In fact, the CG only contains 9 neurons but plays a crucial role in modulating the cardiac circuit’s robustness^33^. Four neuropeptides, including myosuppressin (pQDLDHVFLRFamide) and three orcokinin neuropeptides (FDAFTTGFGHN, NFDEIDRSGFGFH, and VYGPRDIANLY), were ubiquitously expressed across all tissues. Myosuppressin pQDLDHVFLRFamide, a well-characterized cardioactive neuropeptide, has been shown to decrease heartbeat frequency in the American lobster^34-36^. While the orcokinin NFDEIDRSGFGFN proved to enhance mid- and hindgut motility in the crayfish *Procambarus clarkii*, it does not influence gut motility in the American lobster^37^. The widespread expression of several orcokinin family members raises intriguing questions regarding their regulatory roles and physiological functions in the American lobster. Notably, 29 unique neuropeptides were found exclusively in the brain, suggesting their localized release and specific functions within this tissue. However, most neuropeptides were shared across multiple tissues, suggesting that they play systemic roles and coordinate neuromodulatory actions throughout the nervous system. Additionally, Val^1^-SIFamide (VYRKPPFNGSIFamide), a specific isoform of the SIFamide unique to the genus *Homarus*, was previously shown to be widely distributed in the STNS, including the STG, CoG, and OG, occasionally observed in the PO, but not in the SG via immunohistochemistry^38^. However, our study detected Val^1^-SIFamide in all examined tissues except the CG, underscoring the high throughput, sensitivity, and accuracy of our MS-based analysis. In summary, our study presents the tissue-specific distribution of the American lobster neuropeptidome, establishing a foundation for future research on the physiological roles and regulatory mechanisms of neuropeptides in the lobster nervous system.

### Novel Neuropeptide Families

#### Leucokinin (LK)

Since LK was first discovered in a cockroach *L. maderae* in 1986, there have been numerous reports of LKs in many invertebrates, including arthropods, annelids, and mollusks^39^. Members of this family are characterized by an FX_1_X_2_WX_3_amide C-teminus, where X_3_ is S, A, or G. In crustaceans, C-terminally amidated LKs were detected in the white shrimp *P. vannamei* brain, in the water flea *D. pulex* via immunohistochemistry, and *in silico* predictions suggest their presence in the spiny lobster *P. argus* and American lobster *H. americanus*^40-43^. Previous MS-based study also detected LK precursor-related peptides in the American lobster^13^ (**Figure 3A**). Interestingly, our current analysis revealed six amidated LKs in the brain, CoG, OG, and TG of the American lobster, all possessing the conserved C-terminal pentapeptide motif FXXWXamide, consistent with findings in other invertebrates (**Figure 3B**). A representative MS/MS spectrum of one amidated LK was also shown in **Figure 3C**, with the fragment ions matching well to the peptide sequence QAFHPWGamide. Currently, the biological function of LKs in crustaceans remains underexplored. A shrimp LKs (PevK-2) was applied to an isolated *C. borealis* STG, showing an increase in both the frequency of the ongoing pyloric rhythm and the activity of multiple pyloric neurons^44^. In insects, LKs function as circulating hormone by acting as diuretic factors, regulating gut contractions, and exerting myotropic activity^39^. Subsequent physiological studies are needed to uncover the role of newly identified LK neuropeptides in the lobster nervous system.

**Figure 3.**
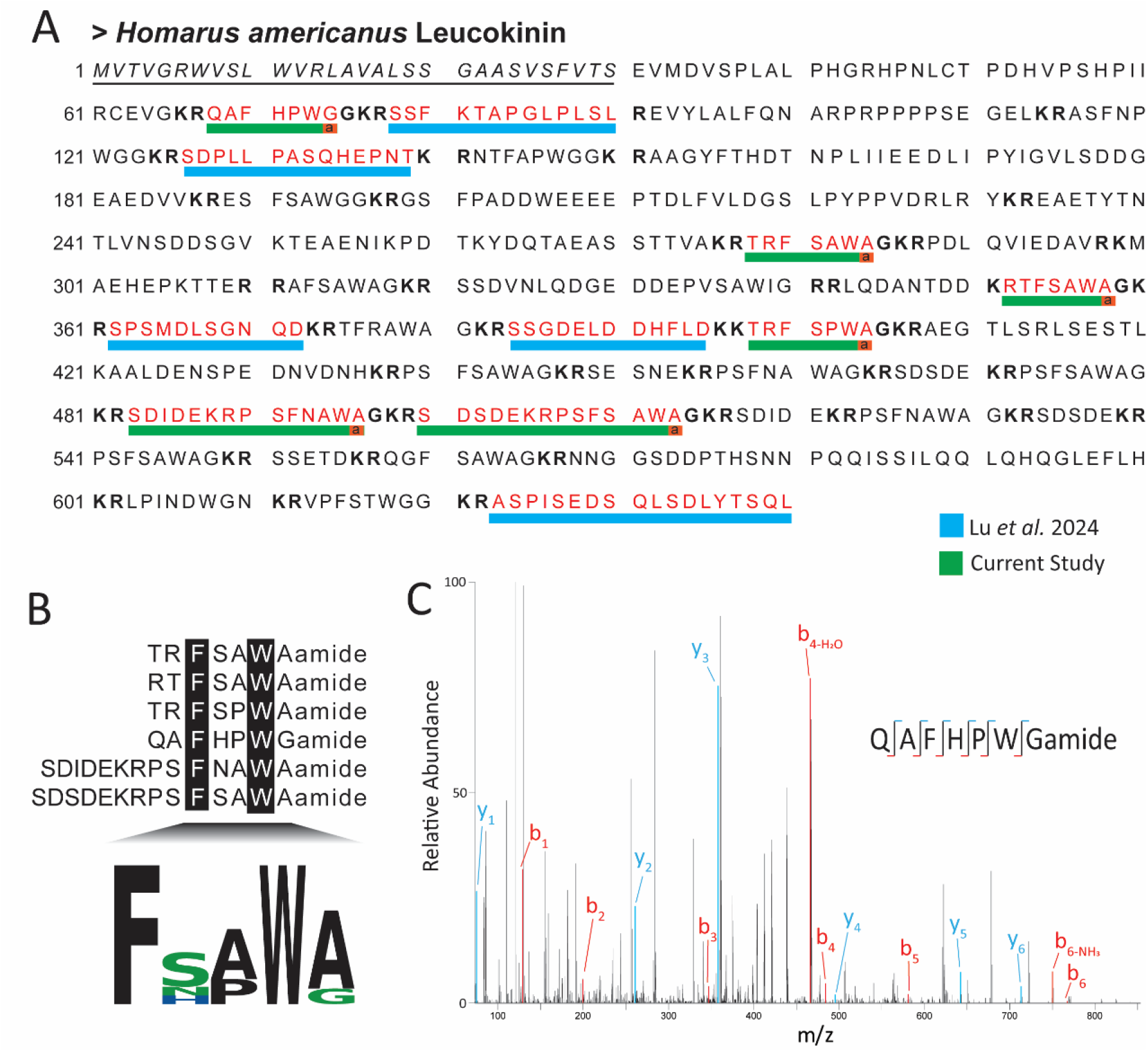
Schematic representation of the predicted H. americanus leucokinin precursor, with identified peptides highlighted using red single-letter amino acid codes. Peptides previously reported by Lu et al. 2024 are indicated by a blue bar, while peptides identified in this study are shown with a green bar. Dibasic cleavage sites are shown in bold, and the predicted signaling peptide at the N-terminus is italicized and underlined. (B) Sequence alignment of six amidated leucokinin peptides, illustrating a conserved C-terminal pentapeptide motif. Identical residues across sequences are highlighted against a black background. (C) Representative MS/MS spectrum of the doubly charged leucokinin peptide ion corresponding to QAFHPWGamide.

#### Periviscerokinin (PVK)

The initial isolation of PVK peptides from the perisympathetic organs of the American cockroach *P. americana* was achieved based on their myotropic activity^45^. In insects, PVK effects are species-specific, demonstrating myotropic activity on various muscle types and contributing to diuresis regulation^46^. The hallmark motif of this novel family is the C-terminal LX_1_X_2_PRX_3_amide. PVK members present in many insects, an arachnid, molluscs and annelid worms^47^. In crustaceans, PVK precursors were found in the transcriptomes of several species such as the American lobster *H. americanus*, the spiny lobster *P. argus*, the copepod *C. finmarchicus* and validated by MS in the water flea *D. pulex* ^12, 42, 48, 49^. Fascinatingly, our study detected QDLIPFPRVamide and pQDLIPFPRVamide in the brain and CG samples, both sharing the conserved C-terminal motif LX_1_X_2_PRX_3_amide. To the best of our knowledge, this is the first recorded identification of PVK neuropeptides in decapods, opening the door for further research into this specific peptide family. The presence of PVK peptides in the lobster CG suggests their involvement in modulating the neurogenic cardiac neuromuscular system, making them prime candidates for further functional studies.

### Gonadotropin-Releasing Hormone (GnRH) Superfamily – Corazonin (CRZ), Adipokinetic hormone CRZ-related Peptides (ACP) and Red Pigment Concentrating Hormone (RPCH)

The GnRH superfamily comprises five members, classified based on structural similarities: vertebrate GnRH, CRZ, ACP, RPCH, and invertebrate GnRH^50, 51^. In insects, ACP is a decapeptide with a WXXamide C-terminus, whereas in crustaceans, it is an undecapeptide with a WVPQamide C-terminus. In our study, we identified the neuropeptide CRZ pQTFQYSRGWTNamide in the brain, ACP pQITFSRSWVPQamide in the brain and CoG, whereas RPCH pQLNFSPGWamide in the brain, CoG, SG, and PO indicating its hormonal role. While CRZs are involved in cardio-acceleratory effects, metabolism, and nutritional stress in insects, their functions in crustaceans remain inconclusive^50, 52, 53^. Similarly, ACPs show no distinct function in most examined insects and crustaceans^50^. Though neural transcriptomic analysis of the female prawn *M. rosenbergii* suggests that ACPs may regulate lipid metabolism and inhibit oocyte proliferation, physiological experiments are needed to confirm this hypothesis^54^. RPCHs in crustaceans primarily regulate pigment concentration in chromatophores and may also influence reproductive processes^55-57^.

### Discovery and Structural Elucidation of Novel Glycosylated Neuropeptides

In our workflow, we employed HILIC enrichment and oxonium ion-triggered EThcD fragmentation to selectively characterize the glycosylated neuropeptide population in the American lobster nervous system. This approach led to the identification of 24 glycosylated neuropeptides from four different families across three neural tissues, displaying a high level of micro- and macro-heterogeneity, as summarized in **Figure 4A**. There is diversity in the types of glycan structures attached to the same peptide backbone or at different glycosylation sites (micro-heterogeneity), as well as variability in the glycoforms of the same neuropeptide family (macro-heterogeneity). Remarkably, the CHH-B family exhibited the highest degree of glycosylation, comprising over 80% of the identified glyconeuropeptides, and was predominantly found in the PO samples. CHH-B neuropeptides are known to regulate glucose homeostasis, molting, osmoregulation, and reproductive processes^58, 59^. Interestingly, O-linked glycosylation was also observed on insulin peptides extracted from mouse and human pancreatic islets^25^. Given the functional parallels in glucose regulation between CHH-B and insulin, O-linked glycosylation may play an important role in modulating their bioactivity, stability, or receptor interactions, suggesting a conserved regulatory mechanisms in peptide hormone signaling. In blue crab *C. sapidus*, the heavily glycosylated neuropeptide families are AST-B and RYamide^20^, contrasting with our findings in the American lobster, which suggests that glycosylation patterns may be species-specific.

**Figure 4.**
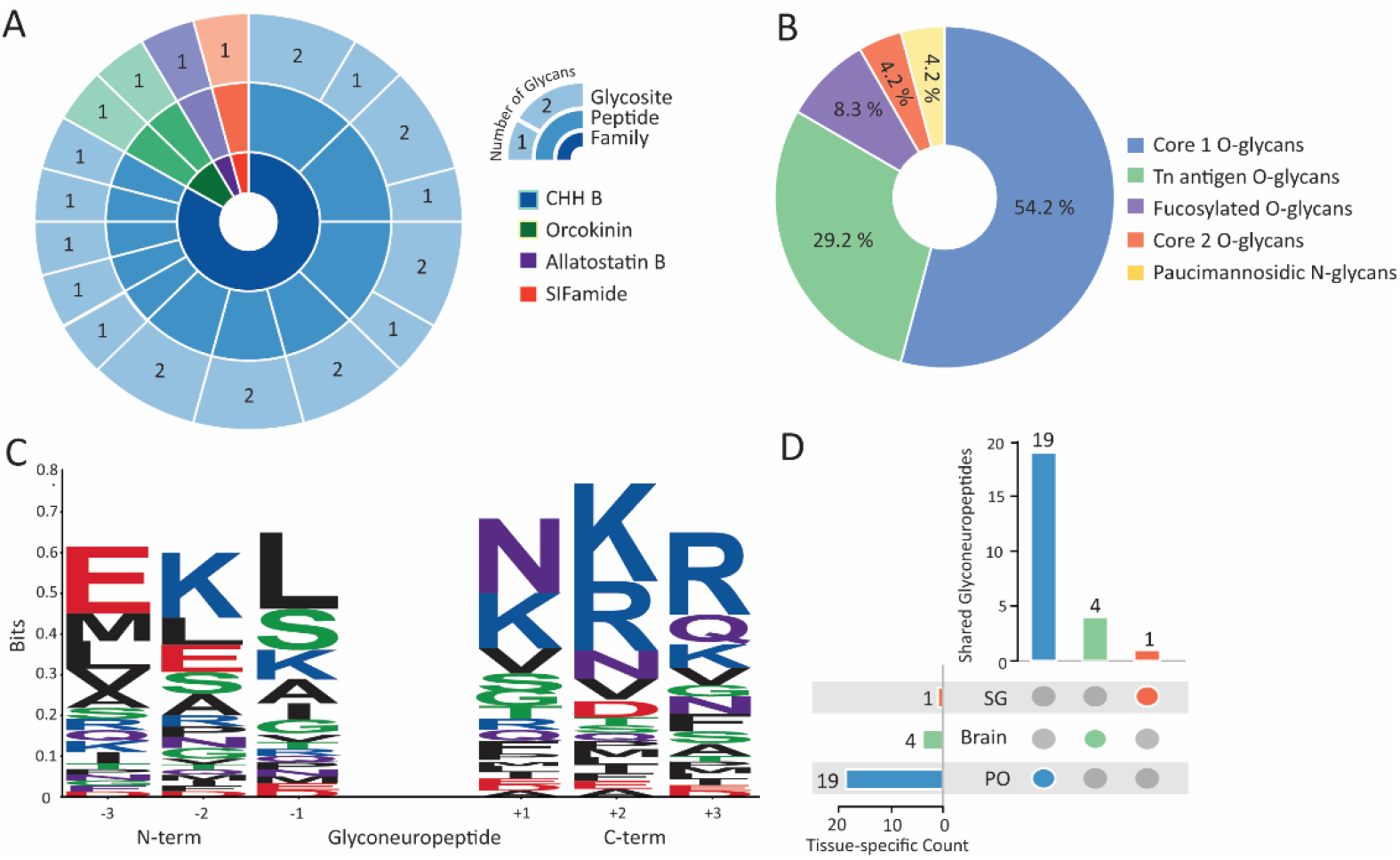
(A) Sunburst chart visualizing the diversity of detected glycosylated neuropeptides. The inner ring represents neuropeptide families, the middle ring displays individual neuropeptides (unique peptide backbones) within each family, and the outer ring maps the glycosylation sites, annotated with the number of distinct glycans observed at each site. (B) Distribution profile of glycan types identified among glyconeuropeptides in the American lobster nervous system. (C) Sequence logo analysis revealing conserved sequence motifs at both the N- and C-terminal regions of the glyconeuropeptides. Amino acids are color-coded by chemical properties: red for acidic, blue for basic, black for hydrophobic, green for polar, and purple for neutral residues. (D) Upset plot summarizing the tissue-specific occurrence of glyconeuropeptides across three neural tissues.

Intriguingly, our study revealed that all identified peptides were O-linked glycosylated, except a novel N-linked Val^1^-SIFamide glycopeptide (**Table 1**). This observation aligns well with previous study of the blue crab *C. sapidus* glyconeuropeptidome where the majority of identified glyconeuropeptides were O-linked, although several N-linked counterparts were detected^20, 21^. This prevalence could be driven by the inherent structural flexibility, the enzyme specificity in the biosynthesis of O-glycans, and the biological advantages provided by this modification. Unlike N-linked glycosylation, which requires a specific consensus motif (N-X-S/T, where N is asparagine, X can be any amino acid except proline, and S/T is serine or threonine), O-glycosylation possibly occurs at nearly any serine or threonine residue along the peptide backbone. Since many neuropeptides are short and contain multiple serine and threonine residues, the N-X-S/T motif could be less common whereas O-glycosylation is likely to be more widespread in these signaling molecules. Additionally, to resolve glycan isomer ambiguity, we employed the GlcNAc/GalNAc ratio as a diagnostic tool to differentiate between GlcNAc and GalNAc residues originating from the HexNAc fragment ion at *m/z* 204.09, typically attached to Ser/Thr residues^60^. This ratio was calculated by dividing the summed intensities of the diagnostic ions at *m/z* 138.06 ([C_7_H_8_NO_2_]^+^) and *m/z* 168.07 ([C_8_H_10_NO_3_]^+^) by those at *m/z* 126.06 ([C_6_H_7_NO_2_]^+^) and *m/z* 144.07 ([C_6_H_10_NO_3_]^+^), all of which arise from HexNAc-derived neutral losses. A ratio below 1 indicates the presence of GalNAc, while a ratio above 1 suggests GlcNAc. As summarized in **Table 1**, all O-glycans displayed GalNAc-type glycosylation with ratios below 1. Although PEAKSGlycanFinder initially assigned some structures as GlcNAc-linked, we revised these annotations based on the calculated GlcNac/GalNac ratio (**Figure S6**).

**Table 1.** Glycosylated neuropeptides detected in the peptidergic signaling system of the American lobster *Homarus americanus*. The bold character represents a glycan position confirmed by EThcD spectra, while the underlined character indicates a putative glycan site suggested by PEAKSGlycanFinder search results. SG, sinus gland; PO, pericardial organ; AST, allatostatin; CHH, crustacean hyperglycemic hormone. (-.98) indicates amidated C-terminal.

| Family | Peptide Sequence | Glycan | Linkage | GlcNAc/<br>GalNAc<br>ratio | Brain | SG | PO |
| --- | --- | --- | --- | --- | --- | --- | --- |
| AST-B | VSS.<u>S</u>(+365.13)SSSPQQDDPASSPSHIEE.KRV | (HexNAc)1(Hex)1 | O | 0.80 | X |  |  |
| CHH-B | EKL.L<u>S</u>(+203.08)SISPSSTPLGFLSQDHSVN.KRQ | (HexNAc)1 | O | 0.42 |  | X |  |
|  | EKL.LSSISPS(+365.13)STPLGFLSQDHSV.NKR | (HexNAc)1(Hex)1 | O | 0.55 |  |  | X |
|  | MEK.LLSSISPS(+365.13)STPLGFLSQDHSV.NKR | (HexNAc)1(Hex)1 | O | 0.61 |  |  | X |
|  | MEK.LLSSISPS(+203.08)STPLGFLSQDHSV.NKR | (HexNAc)1 | O | 0.59 |  |  | X |
|  | EKL.LSSISPS(+365.13)STPLGFLSQDHS.VNK | (HexNAc)1(Hex)1 | O | 0.68 |  |  | X |
|  | LLS.SSISPS(+365.13)STPLGFLSQDHSV.NKR | (HexNAc)1(Hex)1 | O | 0.55 |  |  | X |
|  | LLS.SSISPS(+365.13)STPLGFLSQDHSVN.KRQ | (HexNAc)1(Hex)1 | O | 0.62 |  |  | X |
|  | LLS.SSISPS(+365.13)STPLGFLSQDHS.VNK | (HexNAc)1(Hex)1 | O | 0.57 |  |  | X |
|  | EKL.LSSISPS(+365.13)STPLGFLSQDHSVN.KRQ | (HexNAc)1(Hex)1 | O | 0.69 |  |  | X |
|  | EKL.LSSISPS(+203.08)STPLGFLSQDHSVN.KRQ | (HexNAc)1 | O | 0.54 |  |  | X |
|  | EKL.LSSISPS(+203.08)STPLGFLSQDHS.VNK | (HexNAc)1 | O | 0.43 |  |  | X |
|  | EKL.LSSISPS(+203.08)STPLGFLSQDHSV.NKR | (HexNAc)1 | O | 0.34 |  |  | X |
|  | LLS.SSISPS(+203.08)STPLGFLSQDHSV.NKR | (HexNAc)1 | O | 0.53 |  |  | X |
|  | EKL.LSSISPS(+203.08)STPLGFLSQDHSV(-.98).NKR | (HexNAc)1 | O | 0.56 |  |  | X |
|  | EKL.LSSISPS(+552.22)STPLGFLSQDH(-.98).SVN | (HexNAc)2(Fuc)1 | O | 0.66 |  |  | X |
|  | EKL.LSSISPS(+365.13)STPLGFLSQDHSV(-.98).NKR | (HexNAc)1(Hex)1 | O | 0.65 |  |  | X |
|  | MEK.LLSSISPS(+552.22)STPLGFLSQDH(-.98).SVN | (HexNAc)2(Fuc)1 | O | 0.65 |  |  | X |
|  | MEK.LL<u>S</u>(+365.13)SISPSSTPLGFLSQDHSV.NKR | (HexNAc)1(Hex)1 | O | 0.63 |  |  | X |
|  | VGG.RSVEGVSRMEKLLSSISPSSTPLGFL<u>S</u>(+365.13)QDHSV(-.98).NKR | (HexNAc)1(Hex)1 | O | 0.97 |  |  | X |
|  | EKL.L<u>S</u>(+365.13)SISPSSTPLGFLSQDHSV.NKR | (HexNAc)1(Hex)1 | O | 0.43 |  |  | X |
| Orcokinin | AAA.GPIKAAPARSSPQQDAAAGY**T**(+365.13)DGAPV.KRF | (HexNAc)1(Hex)1 | O | 0.53 | X |  |  |
|  | IPI.AAPAR<u>S</u>(+568.21)SPQQDAAAGY(-.98).TDG | (HexNAc)2(Hex)1 | O | 0.74 | X |  |  |
| SIFamide | VSA.VYRKPPFN(+1022.38)GSIF(-.98).GKR | (HexNAc)2(Hex)2(Fuc)2 | N | 6.52 | X |  |  |

Upon analyzing the glycan composition of these glyconeuropeptides, we putatively assigned them to different core classes based on their composition and the most likely structural configurations (**Figure 4B**). Notably, more than half of the observed O-glycans exhibited a Core 1 structure, defined by a hexose linked to GalNac (N-acetylgalactosamine). This structure is one of the hallmark of mucin-type O-glycans, which play a crucial role in the development of the CNS and neuromuscular junctions in *D. melangogaster*, controlling synaptic molecular assemblies, neurotransmission strength, and synaptic plasticity^61^. Moreover, Tn antigen, the simplest form of O-GalNAcylation consisting of a single GalNAc linked to the peptide backbone, emerged as the second most prevalent glycan observed on neuropeptides in our study. This glycan is often associated with various cellular processes including cell adhesion, migration, and signal transduction, as well as implicated in pathological conditions such as tumor progression, suggesting its potential importance in neuromodulation and neuropeptide function^62, 63^. Previous studies have reported the expression of the Tn antigen in *Drosophila melanogaster* embryos and nervous system, reinforcing its relevance in neurodevelopment and neural signaling^61, 64^. In addition, several fucosylated O-glycans were also detected in the lobster glyconeuropeptidome, highlighting the diversity of glycosylation modifications in this species. Intriguingly, many O-fucosylated peptides, where a single deoxyhexose (fucose) is added to the hydroxyl group of Ser or Thr, were reported in the PEAKSGlycanFinder search results. These assignments were supported by the presence of a diagnostic oxonium ion at *m/z* 147.08 (data not shown). However, O-linked fucosylation is typically associated with specific structural domains, such as Epidermal Growth Factor (EGF)-like repeats or thrombospondin type 1 repeats (TSRs), within larger glycoproteins^65^. Its occurrence on short, secreted neuropeptides is highly unusual and may represent a non-canonical or artifactual modification. Therefore, we excluded these putative O-fucosylated peptides from the final characterization of the lobster glyconeuropeptidome. Overall, the glycan composition in the American lobster nervous system mirrors patterns observed in other invertebrates, suggesting shared functional roles.

We performed a linear sequence analysis to investigate the cleavage patterns of glycosylated neuropeptides (**Figure 4C)**. Interestingly, while basic residues were enriched at the -2 N-terminal and +1, +2 C-terminal positions, leucine (L), serine (S), and asparagine (N) residues were also prominent at the first position of both the C- and N-termini. This pattern contrasts with the typical cleavage sites of many endogenous neuropeptides, which generally exhibit a preference for dibasic or monobasic residues (*e*.*g*., lysine and arginine, as illustrated in **Figure S4**). We hypothesize that the glycan modification on the neuropeptide backbone might alter the steric structure of the neuropeptide precursor, potentially making certain bonds more accessible to proteases or affecting enzyme specificity. These observations imply that glycosylation not only affects peptide stability and receptor interactions but may also modify proteolytic cleavage patterns, resulting in the generation of functional neuropeptide fragments with altered bioactivities. The tissue-specific distribution of glycosylated neuropeptides was also plotted in **Figure 4D**. Among the eight target neural tissues, the PO, one of the primary neuroendocrine sites in the American lobster, unsurprisingly exhibited the richest and most unique repertoire of glyconeuropeptides. Several glycopeptides were also found in the SG and brain samples. However, no glycosylated neuropeptides were detected in any other tissues, which can likely be attributed to their very low *in vivo* abundance in these tissues, making them particularly challenging for MS detection, rather than the absence of glycosylation itself. Furthermore, all of the identified glyconeuropeptides were not widely shared across tissues, indicating a strong degree of tissue-specific glycosylation or localized peptide function, which is consistent with similar findings in the blue crab *C. sapidus*^20^.

EThcD provides gentle fragmentation that preserves intact glycans on the peptide backbone, yielding rich spectra containing glycan fragments with both *b/y*- and *c/z*-type ions for confident glycopeptide identification. A representative EThcD fragmentation spectrum of the glycosylated form of the well-characterized neuropeptide Val^1^-SIFamide was shown in **Figure 5**. Val^1^-SIFamide has the sequence VYRKPPFNGSIFamide, with potential N-glycosylation at Asn8 within the conserved N-G-S motif, or potential O-glycosylation at Ser10. Initially, the PEAKSGlycanFinder software assigned Ser10 as the glycosylation site due to the presence of glycan-containing *y*3 ions (**Figure S6.27**). However, the difucosylated paucimannosidic glycan structure is commonly observed in many insect glycopeptides and has previously been reported on an N-linked glycosylated *Drosophila melanogaster* sex-peptide pheromone^66, 67^. Upon manual inspection and annotation, the presence of a glycan-retaining *z*5 and *c*8 ions in the MS/MS spectrum supports that glycosylation should occur at Asn8. Moreover, the GlcNAc/GalNAc ratio was calculated to be 6.52, strongly suggesting that the attached HexNAc is GlcNAc, consistent with N-linked glycosylation. Taken together, these findings support that glycosylation occurs at Asn8, and we therefore classified this as an N-linked Val^1^-SIFamide glycoform. In our study, Val^1^-SIFamide was broadly expressed across all tissues except CG, but its glycosylated counterpart was detected only in the brain, suggesting a brain-specific functional role. Val^1^-SIFamide was shown to activate the pyloric motor pattern by increasing burst amplitude and duration in the pyloric dilator neurons in the American lobster^38^. Future electrophysiological experiments could reveal whether glycosylation could alter its modulatory effects on the STG compared to the non-modified form.

**Figure 5.**
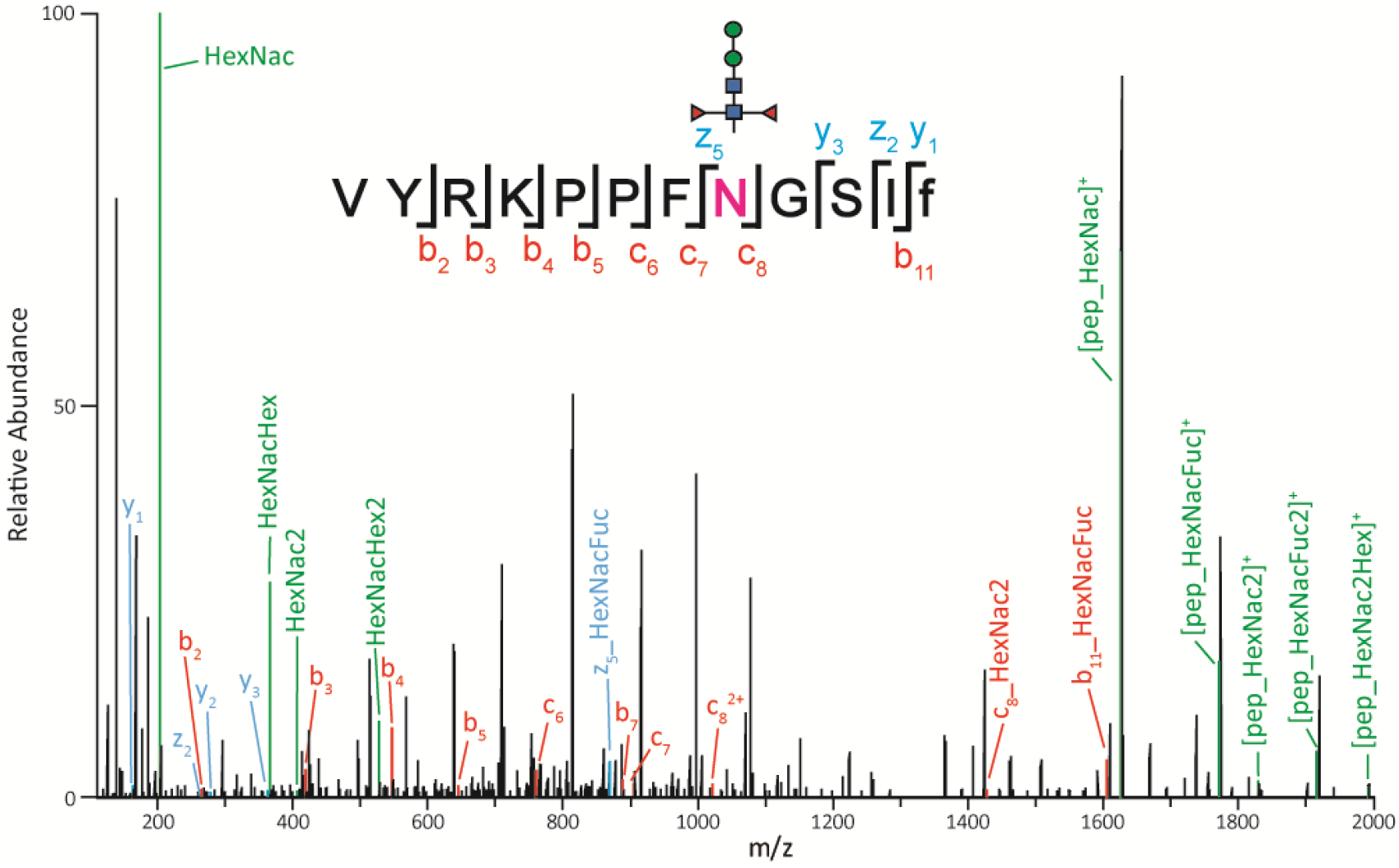
Representative MS/MS spectrum of the doubly charged N-linked glycosylated Val^1^-SIFamide neuropeptide ion, highlighting the diagnostic oxonium ions along with *b/y* and *c/z* fragment ions used for peptide sequencing and glycosylation site localization.

## Discussion

Although prior studies have made significant efforts to profile the neuropeptidome of the American lobster’s peptidergic signaling via high-throughput nucleotide sequencing^43^, transcriptomic mining^68^, MS-based *de novo* sequencing^69^, or ion mobility MS-based analysis^13^, the published high-quality genome draft of the American lobster suggests that many novel neuropeptides remain to be discovered in this important neuroscience model organism. By incorporating HILIC enrichment into our sample preparation workflow, we have significantly expanded its known neuropeptidome landscape, identifying 154 unique neuropeptides from 25 families, with approximately one-third of them being reported for the first time. Using oxonium ion-triggered EThcD fragmentation approach, we also characterized 24 glycosylated neuropeptides for the first time, greatly advancing our understanding of the American lobster nervous system. Our analysis elucidates key cleavage patterns, peptide properties, tissue-specific, and species-specific distribution that further suggests their distinct roles in neuromodulation and neuroendocrine regulation.

Nonetheless, we were still unable to detect many of the novel neuropeptides predicted by the high-quality genome draft of the American lobster. This gap likely stems from the inherent limitations of the current sample preparation and enrichment strategies, which may not capture all peptide variants due to significant sample loss and a bias toward hydrophilic species. Moreover, neuropeptides carrying liable PTMs (*e*.*g*., sulfation), or those with reduced ionization efficiency due to intramolecular disulfide bonds (*e*.*g*., neuroparsin, bursicon, trissin, CCRFamide) pose substantial challenges for MS detectability. To address these issues, alternative sample preparation techniques, such as high pH fractionation, reduction and alkylation of disulfide bonds or the implementation of different glycopeptide enrichment strategies, should be explored. Furthermore, numerous studies demonstrated that some neuropeptides are expressed only under specific external stimuli or are present at very low *in vivo* abundance under control conditions^70-72^. Thus, incorporating external stimuli could offer a more comprehensive understanding of the full spectrum of neuropeptides expressed in the lobster nervous system. Most importantly, while MS enables confident identification of the peptide sequence, glycan composition, and precise glycosylation site, it does not provide information about glycosidic linkages or the specific isomeric forms of the glycans. Therefore, the newly discovered glyconeuropeptides should be further validated through spectral comparison with glycopeptide standards before proceeding with functional studies. Exploring alternative MS data acquisition techniques, including different fragmentation approaches (*e*.*g*., Electron-Activated Dissociation^73^) or advanced separation methods (*e*.*g*., ion mobility spectrometry^74^), could also improve sensitivity and detection of more elusive glyconeuropeptides.

It is also crucial to conduct future research that investigates how the American lobster neuropeptidome responds to external environmental stimuli (*e*.*g*., temperature variations, pH fluctuations) or behavioral conditions (*e*.*g*., feeding). These factors could trigger significant shifts in neuropeptide expression, shedding light on the adaptive and regulatory mechanisms of the nervous system under stress conditions. Furthermore, exploring the neuropeptidome of the lobster hemolymph presents an intriguing opportunity to better understand the hormonal roles of the newly identified neuropeptides. Given that hemolymph is a complex matrix containing salts, lipids, and large proteins, developing a tailored experimental protocol will be essential for effectively extracting and recovering neuropeptides from this medium. Altogether, this study significantly expands the understanding of the neurochemical composition of the American lobster neural circuits, paving the way for future research into the functional roles of these neuropeptides in neuroendocrine regulation and adaptive responses.

## Data Availability Statement

The mass spectrometry proteomics data have been deposited to the ProteomeXchange Consortium via the MassIVE partner repository with dataset identifier MSV000097908

## Author Contributions

Conceptualization: V.N.H.T., G.L., and L.L.; Experimentation: V.N.H.T., T.D., and G.L.; Data acquisition and analysis: V.N.H.T., G.L., A.E.I., and F.W.; Visualization: V.N.H.T and M.B.; Initial manuscript drafting: V.N.H.T.; Funding acquisition: L.L. All authors have given approval to the final version of the manuscript.

## Notes

The authors declare no competing financial interest.

## Acknowledgments

The authors would like to thank Prof. Iain Wilson (BOKU University) for his valuable discussions and insightful advice on the annotation of glycan structures. The study was supported in part by a National Science Foundation grant CHE-2108223 and National Institutes of Health grants R01DK071801, R01AG078794, and R01AG052324. Some of the mass spectrometers were acquired using NIH shared instrument grants S10 OD028473, S10 RR029531, and S10 OD025084. A.E.I. was supported by the National Science Foundation Graduate Research Fellowship Program under Grant No. DGE-2137424. L.L. wishes to acknowledge additional funding support from the Vilas Distinguished Achievement Professorship, the Charles Melbourne Johnson Professorship, the Wisconsin Alumni Research Foundation, and the University of Wisconsin-Madison School of Pharmacy.

