## Supplemental Information for "HILIC-Enabled Mass Spectrometric Discovery of Novel Endogenous and Glycosylated Neuropeptides in the American Lobster Nervous System"

### Table of Contents

**Table S1.** Neuropeptides detected in the American lobster *Homarus americanus* nervous system

**Figure S1.** The peptide concentration in the HILIC-enriched LC vials of eight neural tissues.

**Figure S2.** Ven diagram showing the number of glycosylated neuropeptides detected in HILIC *versus* Non-enriched samples across six neural tissues.

**Figure S3.** Sequence length, hydrophobicity, and net-charge behavior of the endogenous neuropeptides in the American lobster nervous system.

**Figure S4.** Sequence logo analysis revealing the cleavage patterns at both the N- and C-termini of the endogenous neuropeptides in the American lobster nervous system.

**Figure S5.1.-144.** MS/MS spectrum of neuropeptide sequences detected in the peptidergic signaling system of the American lobster *Homarus american*us generated by PEAKS XPro software.

**Figure S6.1.-24.** MS/MS spectrum of O-linked glycosylated neuropeptide sequences detected in the peptidergic signaling system of the American lobster *Homarus american*us generated by PEAKSGlycanFinder v.2.5. software.

**Table S1. Neuropeptides detected in the American lobster *Homarus americanus* nervous system.** TG, thoracic ganglion; SG, sinus gland; CoG, commissural ganglion; PO, pericardial organ; OG, oesophageal ganglion; STG, stomatogastric ganglion; CG, cardiac ganglion; AST, allatostatin; CHH, crustacean hyperglycemic hormone; CCAP, crustacean cardioactive peptide; DH, diuretic hormone; PDH, pigment dispersing hormone; ACP, adipokinetic hormone corazonin-related peptides; RPCH, red pigment concentrating hormone; ALP, agatoxin-like peptide. (-.98) indicates amidated C-terminal; (-17.03) and (-18.01) indicate the N-terminal pyroglutamate formation; (+15.99) indicates the oxidation on methionine. Peptides previously reported in *H. americanus*[1] are shown in regular font; Peptides reported in prior studies[1] but not detected in the current analysis are shown in *italics*, with their tissue distribution based on those earlier reports; peptides identified for the first time in this study are shown in ***bold italics***.

| Family | Peptide Sequence | Mass | TG | Brain | SG | CoG | PO | OG | STG | CG |
|---|---|---|---|---|---|---|---|---|---|---|
| AST-A | GKR.HSNYGFGL(-.98).GKR | 892.42 | | X | X | X | X | X | X | |
| | GKR.SVGDLPEVSKVEDGASPRT.KRD | 1941.96 | | X | | X | X | X | X | |
| | NKR.SKLYGFGL(-.98).GKR | 882.5 | | X | X | X | X | X | X | |
| | EKR.PRNYAFGL(-.98).GKR | 935.5 | | X | X | X | X | X | X | |
| | DKR.PRDYAFGL(-.98).GKR | 936.48 | X | X | X | X | X | X | X | |
| | *GKR.ASSDEDDEERYYAYEQ(-.98).GKR* | 1967.77 | | X | X | | | | | X |
| | GKR.AGRYAFGL(-.98).GKR | 852.46 | | X | X | X | X | X | X | |
| | GKK.AGHYAFGL(-.98).GKR | 833.42 | | X | X | X | X | X | X | |
| | ***GKR.TPGYAFGL(-.98).GKR*** | 823.43 | | X | X | X | X | | | |
| | ***GKK.AGQYSFGL(-.98).GKR*** | 840.42 | | X | X | X | X | | X | |
| | ***GKR.SDAPDSGFGRRSYDFGL(-.98).GKR*** | 1844.85 | X | X | | X | X | X | X | |
| | ***GKR.TGPYAFGL(-.98).GKR*** | 823.43 | | X | X | X | X | | | |
| | ***DKR.SDLYSFGL(-.98).GKK*** | 899.44 | | X | X | X | X | | | |
| | ***GKK.SGSYNFGL(-.98).GKR*** | 842.40 | | X | X | X | X | | | |
| | ***DKR.SQMYSFGL(-.98).GKR*** | 930.43 | | X | | | X | | | |
| | ***DKR.SQM(+15.99)YSFGL(-.98).GKR*** | 946.42 | | X | X | X | X | | X | |
| | ***GKR.PTAYSFGL(-.98).GKR*** | 853.44 | | X | X | X | X | | | |
| | ***GKR.ADPYAFGL(-.98).GKK*** | 851.42 | | | | | X | | | |
| | GKR.SDSDSDQYTL(-.98).GRR | 1128.46 | | | | X | | | | |
| AST-B | VSS.SSSSPQQDDPASSPSHIEE.KRV | 1983.83 | | X | | | X | | X | |
| | EKR.VGWSSMHGTW(-.98).GKR | 1145.51 | | X | | X | X | | | |
| | EKR.VGWSSM(+15.99)HGTW(-.98).GKR | 1161.5 | | X | | | X | | | |
| | GKR.PHLEDAQLDAAEV.KRT | 1406.67 | | X | X | X | X | X | X | |
| | ***SLR.GTWGKRSADWNKL.RGA*** | 1517.77 | | X | | X | | | X | |

| | | | | | | | | | |
|---|---|---|---|---|---|---|---|---|---|
| | ***GKR.GEELQAAED.KRT*** | 960.40 | | X | | X | X | X | X |
| | VKR.TNWNKFQGSW(-.98).GKR | 1265.59 | | X | | X | X | | |
| | GKR.NNWRSLQGSW(-.98).GKR | 1245.6 | | X | | | X | | X |
| | GKR.AWNKLQGAW(-.98).GKR | 1071.56 | | X | | X | X | | X |
| | SPR.STNWSSLRGTW(-.98).GKR | 1292.63 | | X | | | X | | X |
| | GKR.SADWNKLRGAW(-.98).GKR | 1301.66 | | X | | X | X | X | X |
| | GKR.ASDWGQFRGSW(-.98).GKR | 1294.58 | | X | | X | X | X | X |
| | GKR.APDMMSVAAPNQA | 1301.57 | | X | | | X | | |
| | ***GKR.APDM(+15.99)MSVAAPNQA*** | 1317.56 | | X | | | | | |
| | ***GKR.APDMM(+15.99)SVAAPNQA*** | 1317.56 | | X | | | | | |
| | GKR.APDM(+15.99)M(+15.99)SVAAPNQA | 1317.57 | | X | | X | X | | |
| AST-C | RLR.NNADIKDLQ.RKR | 1030.51 | | X | | | | | |
| CHH-A | VGG.RSVEGASRMEKLLSSSNSPSSTPLGFLSQDHSVN.KRQ | 3603.76 | | | X | | X | | |
| | VGG.RSVEGASRM(+15.99)EKLLSSSNSPSSTPLGFLSQDHSVN.KRQ | 3619.75 | | | X | | X | | |
| CHH-B | VGG.RSVEGVSRMEKLLSSISPSSTPLGFLSQDHSVN.KRQ | 3543.8 | | | X | | | | |
| | *VGG.RSVEGVSRM(+15.99)EKLLSSISPSSTPLGFLSQDHSVN.KRQ* | 3559.79 | | | X | | | | |
| Corazonin | AAA.Q(-17.03)TFQYSRGWTN(-.98).GRK | 1368.62 | | X | | | | | |
| CCAP | AKR.DIGDLLEGKD.KRP | 1073.52 | X | X | | X | X | X | X |
| | *RKR.STPHTQPRQHLTSTPQQKVETEKQ* | 2785.41 | X | X | | | | | |
| DH | *EAR.AVVQIEDPDYVLELLTRLGHSIIRANELEKFVRSSGSA.KRG* | 4224.25 | | X | | | | | |
| | AKR.GLDLGLGRGFSGSQAAKHLMGLAAANFAGGP(-.98).GRR | 2939.52 | | X | | X | | X | X |
| | ***AKR.GLDLGLGRGFSGSQAAKHLM(+15.99)GLAAANFAGGP(-.98).GRR*** | 2955.51 | | X | | X | | X | X |
| | RRR.SSDDGLDLHHDDNLYAQDQAADLAESS.R | 2901.22 | | X | | | | | X |
| FLRFamide | EKR.LLKYFLPASQAWGGDAYPIGQEGT.KRG | 2581.29 | | X | | X | X | X | X |
| | TKR.GYSDRNYLRF(-.98).GRS | 1288.63 | | X | X | X | X | X | X |
| | FGR.SDTNDYEGEEMPESPE.KRN | 1827.66 | | X | | | | | |
| | ***VLR.Q(-17.03)INAHRI.KRA*** | 833.45 | | X | | X | X | X | X |
| | ***QNR.NFLRFGRS(-.98).GSP*** | 994.55 | | X | | X | X | X | X |
| | ***SDRNFLRF(-.98)*** | 1052.56 | | | | | X | | |
| | EKR.NRNFLRF(-.98).GRD | 964.54 | | X | | X | X | X | X |

| | | | | | | | | | | |
|---|---|---|---|---|---|---|---|---|---|---|
| | FGR.DQNRNFLRF(-.98).GRS | 1207.62 | | X | X | X | X | X | X | |
| | FGR.SGSPMEFATDLQEDVELPVEE.KRG | 2321.03 | | | | X | X | X | X | |
| | ***FGR.SGSPM(+15.99)EFATDLQEDVELPVEE.KRG*** | 2337.02 | | | | | | X | X | |
| | EKR.GAHKNYLRF(-.98).GRG | 1103.6 | | X | | X | X | X | X | |
| | FGR.GNRNFLRF(-.98).GRG | 1021.56 | | X | X | X | X | X | X | |
| | FGR.GDRNFLRF(-.98).GRS | 1022.54 | | X | X | X | X | X | X | |
| | AKR.FSHDRNFLRF(-.98).GKR | 1336.68 | | X | X | X | X | X | X | |
| | GKR.DGSDDYPSSSSSAESPPVEYPRYV.RAP | 3310.56 | | X | | X | X | X | X | |
| | YVR.APSKNFLRF(-.98).G | 1077.61 | | X | | X | X | X | X | |
| GSEFLamide | SAA.LPTHLPDELDDPVV.KRL | 1558.79 | X | X | | X | | | | |
| | ***GVR.RIGSEFL(-.98).GKR*** | 819.46 | | X | X | X | | X | | |
| | *VKR.LAGTPHESMIRYFLMAMSNPAGRYKSPQLLNRGV.RRI* | 3804.94 | | X | | | | | | |
| | ***GKR.QYEPEFAHTLDYDT.KRA*** | 1727.73 | | X | | X | | | | |
| | GKR.Q(-17.03)YEPEFAHTLDYDT.KRA | 1727.73 | | X | | | | | | |
| Myosuppressin | VKR.QDLDHVFLRF(-.98).GRS | 1287.67 | | X | X | X | | X | X | X |
| | VKR.Q(-17.03)DLDHVFLRF(-.98).GRS | 1270.65 | X | X | X | X | X | X | X | X |
| Orcokinin | *AAA.GPIKVRFLSAIFIPIAAPARSSPQQDAAAGYTDGAPV.KRF* | 3752 | | X | | | | | | |
| | AAA.GPIKAAPARSSPQQDAAAGYTDGAPV.KRF | 2495.24 | | X | X | X | X | | | |
| | VKR.FDAFTTGFGHN.KRS | 1212.52 | X | X | X | X | X | X | X | X |
| | NKR.SSEDMDRLGFGFN.KRN | 1473.62 | | X | X | X | X | X | X | |
| | NKR.SSEDM(+15.99)DRLGFGFN.KRN | 1489.61 | | X | X | X | X | X | X | |
| | NKR.NFDEIDRSGFGFH.KRN | 1539.67 | X | X | X | X | X | X | X | X |
| | HKR.NFDEIDRSGFGFN.KRN | 1516.66 | | X | X | X | X | X | X | X |
| | HKR.GDYDVYPE.KRN | 956.38 | | X | X | X | X | X | X | |
| | EKR.NFDEIDRSGFGFV.KRV | 1501.68 | X | X | X | X | X | X | X | |
| | VKR.VYGPRDIANLY.KRN | 1279.66 | X | X | X | X | X | X | X | X |
| PDH | *IQA.Q(-17.03)ELKYPEREVVADMAAQILRVALGPWGSVAAVPR.KRN* | 3701.97 | | X | | | | | | |
| | ***IQA.Q(-17.03)ELKYPEREVVADM(+15.99)AAQILRVALGPWGSVAAVPR.KRN*** | 3717.96 | | X | | | | | | |
| | ***IQA.Q(-17.03)ELKYPEREVVADMAAQIL.RVA*** | 2185.10 | | X | | | | | | |
| | ***IQA.Q(-17.03)ELKYPEREVVADM(+15.99)AAQIL.RVA*** | 2201.09 | | X | | | | | | |
| | ***TQA.Q(-17.03)ELKYPEREVVAELAAQIL.RVI*** | 2181.16 | | X | | | | | | |

| | | | | | | | | | |
|---|---|---|---|---|---|---|---|---|---|
| | RKR.NSELINSLLGIPKVMNDA(-.98).GRR | 1926.02 | | X | X | | | | |
| | RKR.NSELINSLLGIPKVM(+15.99)NDA(-.98).GRR | 1942.02 | | X | X | | | | |
| | HKR.NSELINSILGLPKVMNDA(-.98).GRD | 1926.02 | | X | X | X | | | |
| | HKR.NSELINSILGLPKVM(+15.99)NDA(-.98).GRD | 1942.02 | | X | X | X | | | |
| | HKR.NSEILNTLLGSQDLSNMRSA(-.98).GRR | 2161.08 | | | | X | | | |
| Pyrokinin | GKR.GDGFAFSPRL(-.98).GKR | 1064.54 | | X | | X | X | | |
| | GKR.GADFAFSPRL(-.98).GRR | 1078.56 | | X | | X | | | |
| | GRR.SEFVFSSRP(-.98).GKK | 1053.52 | | X | | X | X | | |
| | ***FVA.VRRSLFSPRL(-.98).GKR*** | 1228.76 | | X | | | | | |
| | ***PGR.AYFSPRL(-.98).G*** | 851.47 | | X | | X | X | | |
| | ***PKR.LYYSQRP(-.98).GKR*** | 924.49 | | X | | X | X | | |
| | ***VRR.SLFSPRL(-.98).GKR*** | 817.48 | | X | | X | X | | |
| | GKK.SDFAFSPRL(-.98).GKK | 1037.53 | | X | | X | | | |
| SIFamide | VSA.VYRKPPFNGSIF(-.98).GKR | 1422.78 | X | X | X | X | X | X | X |
| | AVY.RKPPFNGSIF(-.98).GKR | 1160.65 | X | X | | | | | |
| Tachykinin | ERR.APSGFLGMR(-.98).GKK | 933.49 | X | X | X | X | | X | X |
| | ERR.APSGFLGM(+15.99)R(-.98).GKK | 949.48 | | X | X | X | X | X | X |
| | ***VSA.AGEGQDTPQDRE.RRA*** | 1301.55 | | X | X | X | | | X |
| | GKK.SDEEVFSDATADNDLEILL.KRA | 2094.95 | | | X | X | | X | X |
| | GKK.YYDDDSDMDAYIQALTAVVDGQQQQ.KRA | 2851.21 | | | | X | | | |
| | GKK.AYYSENPDEEISM(+15.99)TGVD.KRT | 1934.77 | | X | | X | | | |
| | *GKK.AYYSENPDEEISMTGVD.KRT* | 1918.78 | X | X | X | | | | |
| | DKR.TPSGFLGMR(-.98).G | 963.5 | X | X | X | X | | X | X |
| | DKR.TPSGFLGM(+15.99)R(-.98).G | 979.49 | | X | X | X | | X | X |
| Natalisin | GKR.PSSELLHQHHQ.KRS | 1311.63 | | X | | X | | | |
| | GKR.DGGGPFWIAR(-.98).GKR | 1073.54 | | X | | X | | | |
| | GKR.Q(-17.03)ETEGNGGPFWIAR(-.98).GKK | 1542.72 | | X | | | | | |
| | *GKR.QETEGNGGPFWIAR(-.98).GKK* | 1559.75 | | X | | | | | |
| | HKR.Q(-17.03)DGGPFWISR(-.98).GKK | 1143.55 | | X | | X | | | |
| | HKR.QDGGPFWISR(-.98).GKK | 1160.57 | | X | | X | | | |
| | *GRK.SEDERTFWVAR(-.98).GKK* | 1393.67 | | X | | | | | |
| | *GKK.ENKGNESELFWISR(-.98).GKR* | 1706.84 | | X | | | | | |

| | | | | | | | | | | |
|---|---|---|---|---|---|---|---|---|---|---|
| | GKR.EGEAPPFWVSR(-.98).GKK | 1272.63 | | X | | X | | | X | |
| | GKK.EGEETHPFWVSR(-.98).GKK | 1471.68 | | X | | X | | | X | |
| | GKK.E(-18.01)GEETHPFWVSR(-.98).GKK | 1453.67 | | X | | | | | | |
| | ***GKK.DAVDGRAPFWISR(-.98).GKK*** | 1487.77 | | X | | X | | | | |
| | ***GKK.DTPALLPVGHPSLWGNR(-.98).GRK*** | 1827.98 | | X | | | | | | |
| | GKK.DTTYGPIDDPFVKGFLALR(-.98).G | 2123.11 | | X | | X | | | X | |
| RYamide | TAA.Q(-17.03)GFYTQRY(-.98).GKR | 1043.48 | | X | X | X | | X | X | |
| | ***TAA.QGFYTQRY(-.98).GKR*** | 1060.51 | | X | | X | | | X | |
| | TVR.SGFYANRN(-.98).GRS | 926.44 | | X | | X | | X | X | |
| | NGR.SSPSQGLPEIKIRSSRFIGGSRY(-.98).GKR | 2520.36 | | | | | | | X | |
| | ***NGR.SSPSQGLPEI.KIR*** | 1013.50 | | X | | X | | | | |
| | KIR.SSRFIGGSRY(-.98).GKR | 1127.58 | | X | | X | | X | X | |
| Gonadoliberin | ATS.Q(-17.03)IHWNRGWGAGGSM(+15.99)(-.98).GKR | 1553.69 | | X | | X | | | | |
| | ATS.Q(-17.03)IHWNRGWGAGGSM(-.98).GKR | 1537.7 | | X | X | X | | | | |
| ACP | ***TLA.Q(-17.03)ITFSRSWVPQ(-.98).GKR*** | 1329.68 | | X | | X | | | | |
| RPCH | ***VSA.Q(-17.03)LNFSPGW(-.98).GKR*** | 929.44 | | X | X | X | X | | | |
| ALP | ***QKR.DDVAGSDPIK.RWR*** | 1015.48 | | X | X | X | | | | |
| | ***VMA.Q(-17.03)PLLEEGREEDGVQQAEPDYAADLLERLLART Q.KRD*** | 3734.83 | | | | X | | X | X | X |
| | ***CQR.M(+15.99)GIFQQW(-.98).GK*** | 923.43 | | X | X | | | | | |
| | ***CQR.MGIFQQW(-.98).GK*** | 907.44 | | X | | | | | | |
| Periviscerokinin | ***RKR.QDLIPFPRV(-.98).GKR*** | 1082.63 | | X | | X | | | | X |
| | ***RKR.Q(-17.03)DLIPFPRV(-.98).GKR*** | 1065.60 | | X | | | | | | X |
| Sulfakinin | ***GKR.EFDEYGHMRF(-.98).GKR*** | 1328.56 | | | X | X | | | | |
| | ***GKR.EFDEYGHM(+15.99)RF(-.98).GKR*** | 2655.11 | | | | X | | | | |
| Leucokinin | GKR.SDPLLPASQHEPNT.KRN | 1504.72 | | X | | X | | | | |
| | *GKR.SPSMDLSGNQD.KRT* | 1149.46 | | X | | | | | | |
| | ***GKR.SPSM(+15.99)DLSGNQD.KRT*** | 1165.45 | | | | X | | | | |
| | *GKR.SSGDELDDHFLD.KKT* | 1348.54 | X | X | | | | | | |
| | ***AKR.TRFSAWA(-.98).GKR*** | 836.43 | X | X | | X | | X | | |
| | ***DDK.RTFSAWA(-.98).GKR*** | 836.43 | | X | | | | | | |
| | ***DKK.TRFSPWA(-.98).GKR*** | 862.45 | X | X | | X | | X | | |

| | | | |
|---|---|---|---|
| ***GKR.QAFHPWG(-.98).GKR*** | 840.41 | | X |
| ***GKR.SSFKTAPGLPLSL.REV*** | 1316.73 | | X |
| ***GKR.SDIDEKRPSFNAWA(-.98).GKR*** | 1633.79 | | X |
| ***GKR.SDSDEKRPSFSAWA(-.98).GKR*** | 1580.73 | | X |
| *GKR.AEGTLSRLSESTLKAALDENSPEDNVDNH.KRP* | 3111.46 | | X |
| *GKR.ASPISEDSQLSDLYTSQL* | 1952.92 | X | X |

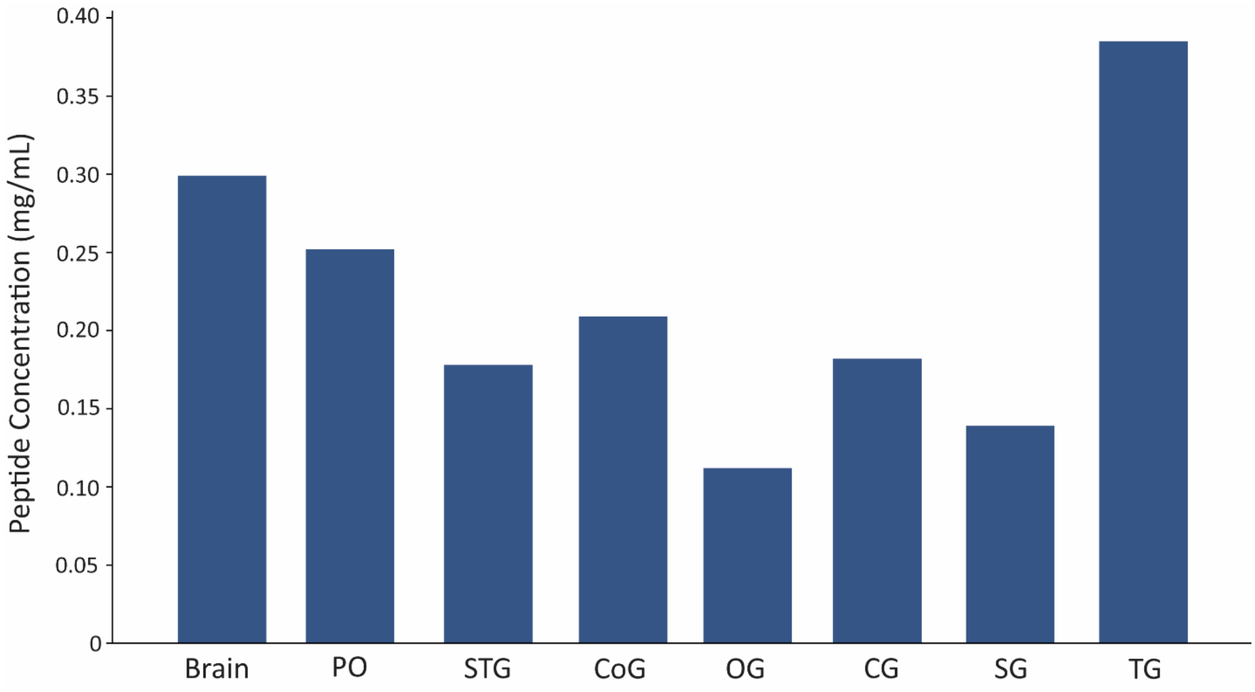


**Figure S1.** The peptide concentration in the HILIC-enriched LC vials of eight neural tissues.

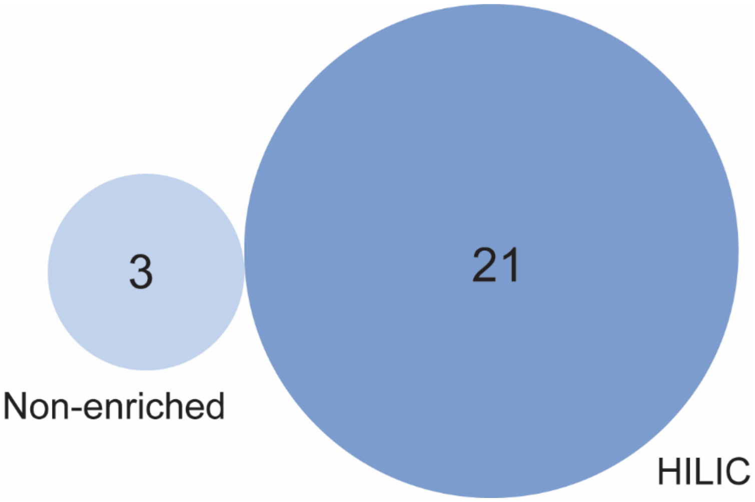


**Figure S2.** Ven diagram showing the number of glycosylated neuropeptides detected in HILIC *versus* Non-enriched samples across six neural tissues.

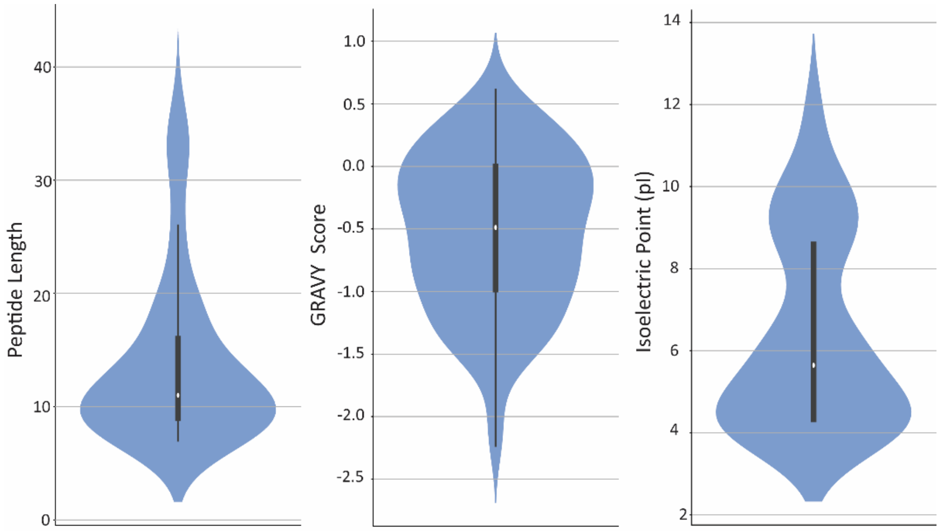


**Figure S3.** Sequence length, hydrophobicity, and net-charge behavior of the endogenous neuropeptides in the American lobster nervous system.

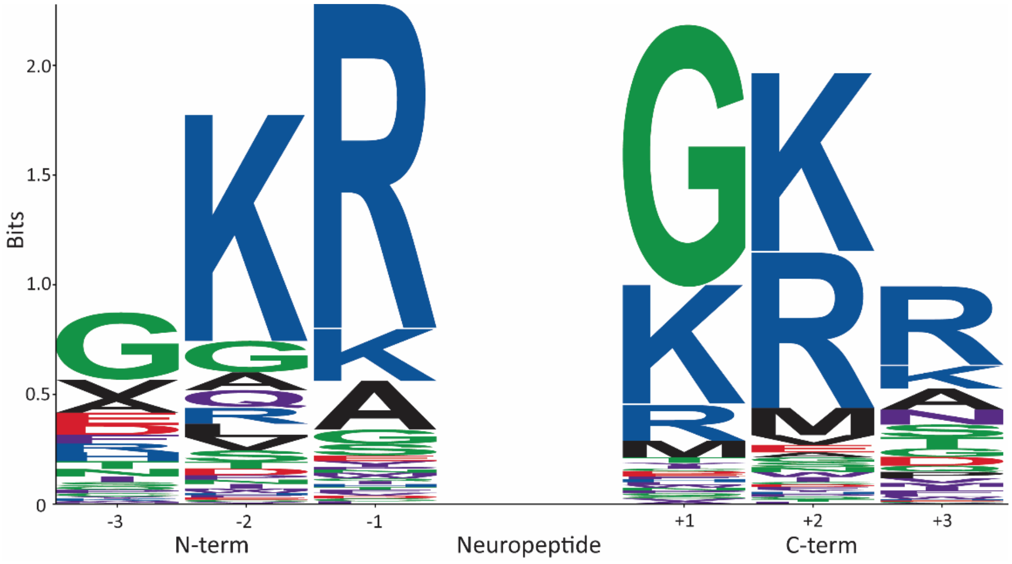


**Figure S4.** Sequence logo analysis revealing the cleavage patterns at both the N- and C-termini of the endogenous neuropeptides in the American lobster nervous system.

**Figure S5.1.-144.** MS/MS spectrum of neuropeptide sequences detected in the peptidergic signaling system of the American lobster *Homarus americanus* generated by PEAKS XPro software.

**Figure S5.1.** MS/MS spectrum of a AST-A neuropeptide with the sequence HSNYGFGL(-.98) detected in the peptidergic signaling system of the American lobster *Homarus americanus* generated by PEAKS XPro software.

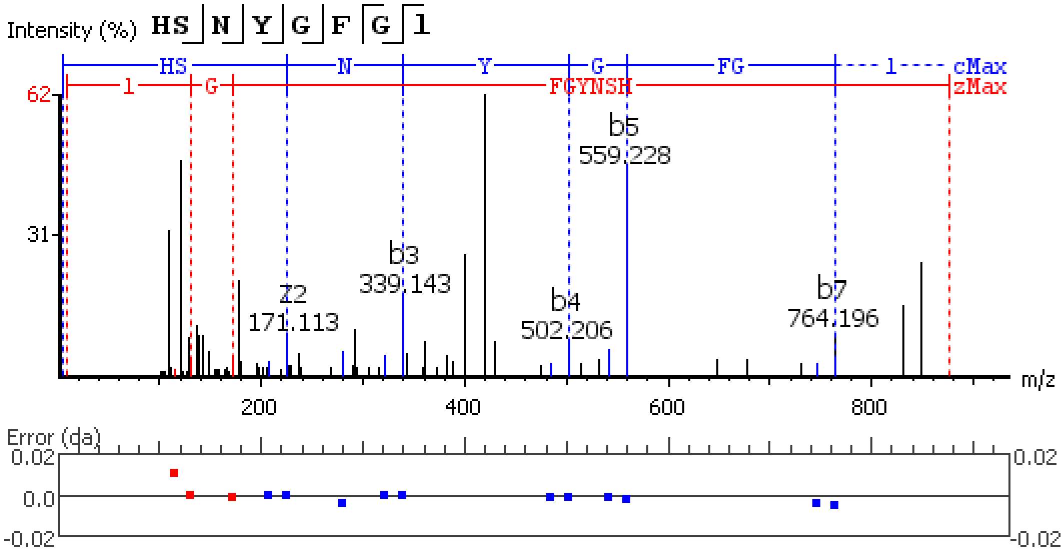


**Figure S5.2.** MS/MS spectrum of a AST-A neuropeptide with the sequence SVGDLPEVSKVEDGASPRT detected in the peptidergic signaling system of the American lobster *Homarus americanus* generated by PEAKS XPro software.

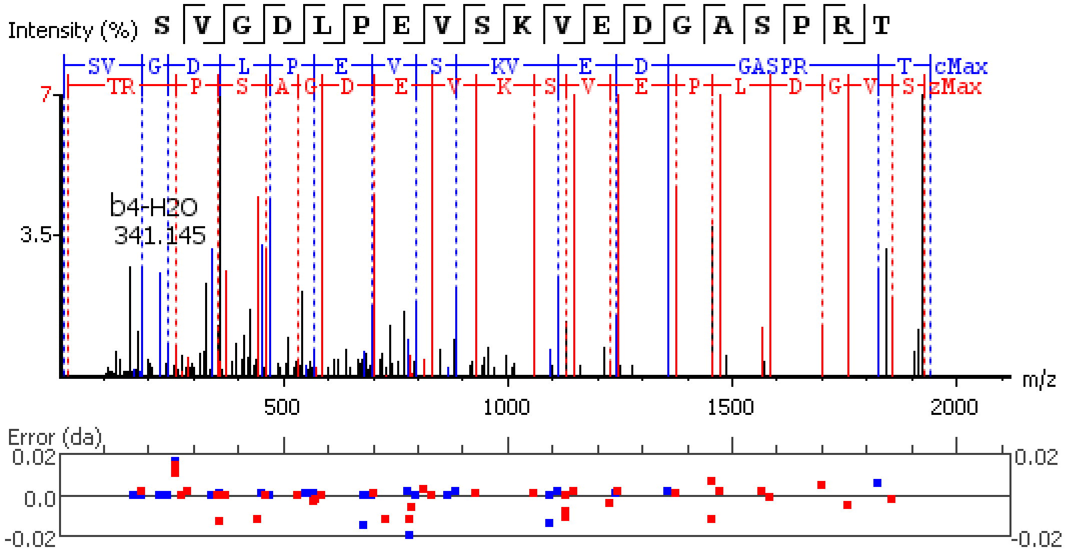


**Figure S5.3.** MS/MS spectrum of a AST-A neuropeptide with the sequence SKLYGFGL(-.98) detected in the peptidergic signaling system of the American lobster *Homarus americanus* generated by PEAKS XPro software.

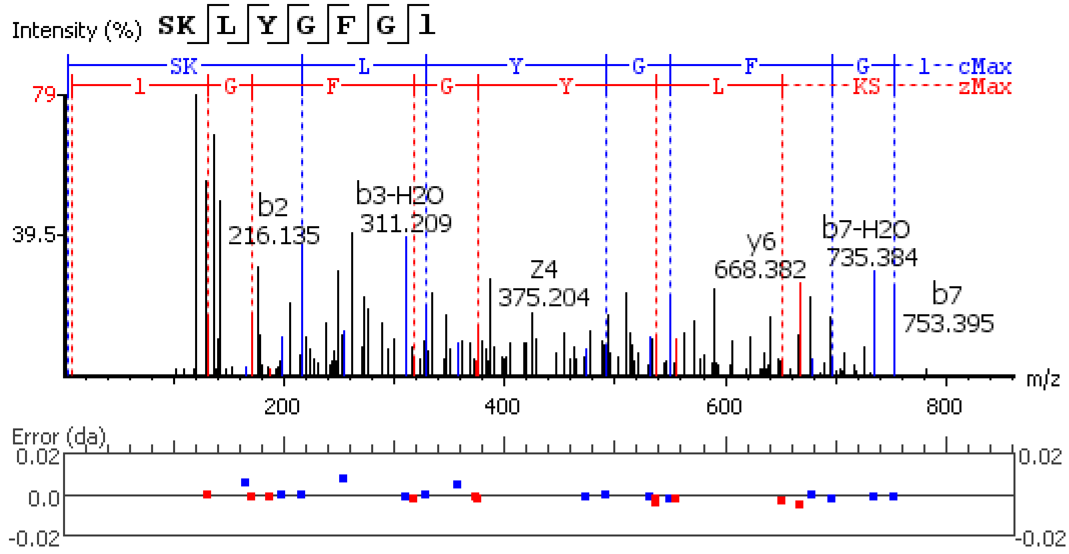


**Figure S5.4.** MS/MS spectrum of a AST-A neuropeptide with the sequence PRNYAFGL(-.98) detected in the peptidergic signaling system of the American lobster *Homarus american*us generated by PEAKS XPro software.

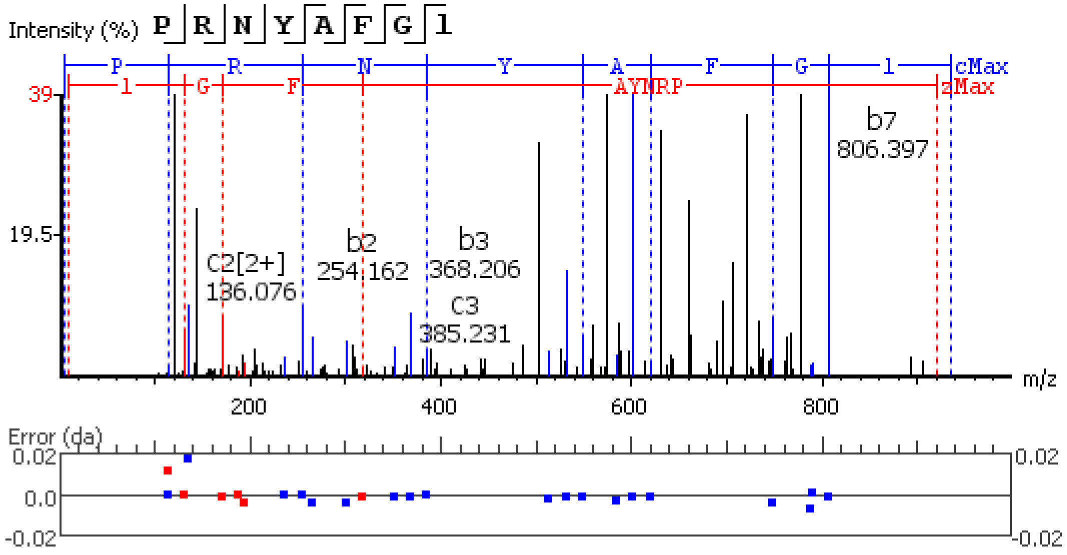


**Figure S5.5.** MS/MS spectrum of a AST-A neuropeptide with the sequence PRDYAFGL(-.98) detected in the peptidergic signaling system of the American lobster *Homarus american*us generated by PEAKS XPro software.

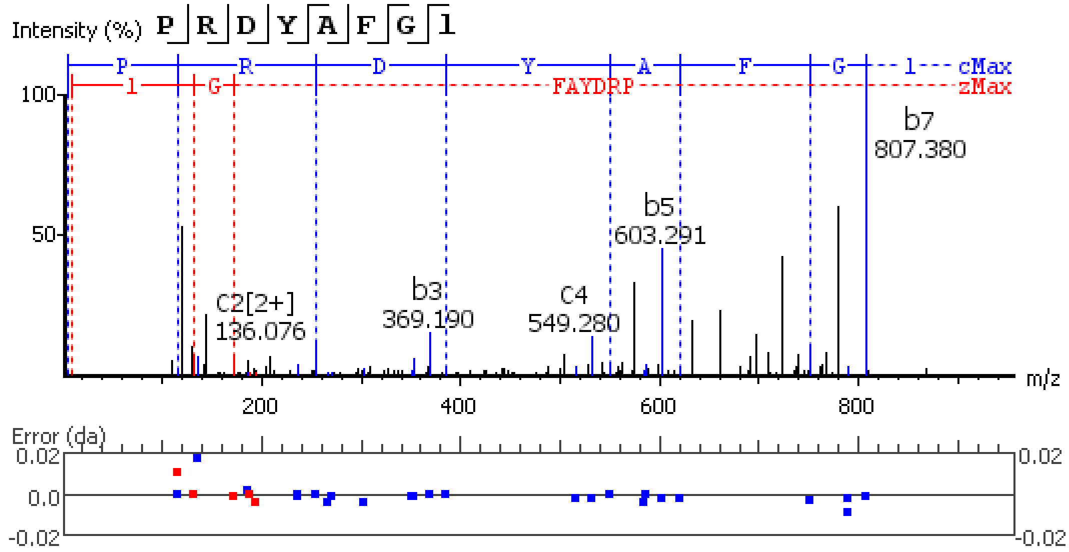


**Figure S5.6.** MS/MS spectrum of a AST-A neuropeptide with the sequence AGRYAFGL(-.98) detected in the peptidergic signaling system of the American lobster *Homarus americanus* generated by PEAKS XPro software.

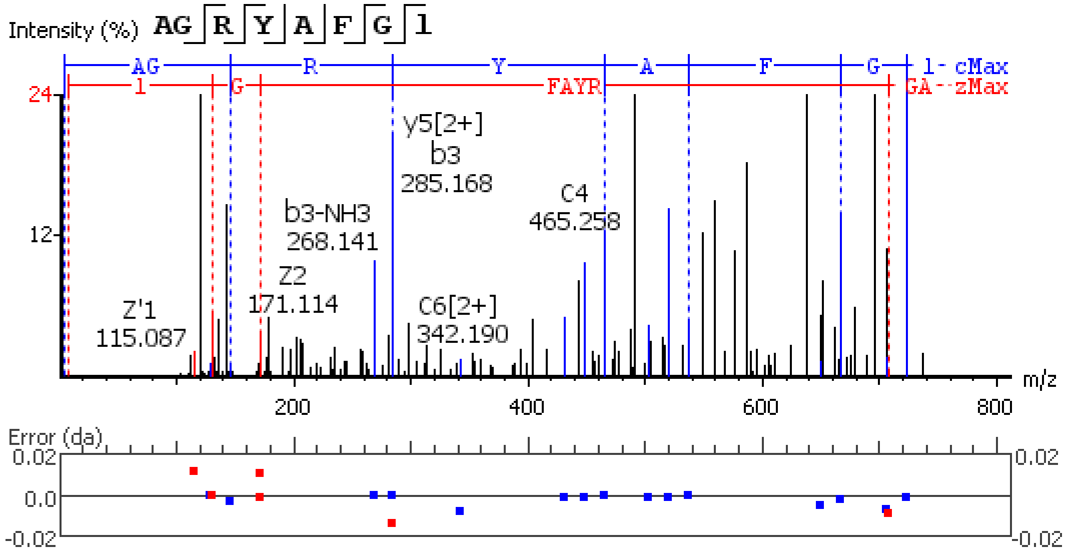


**Figure S5.7.** MS/MS spectrum of a AST-A neuropeptide with the sequence AGHYAFGL(-.98) detected in the peptidergic signaling system of the American lobster *Homarus americanus* generated by PEAKS XPro software.

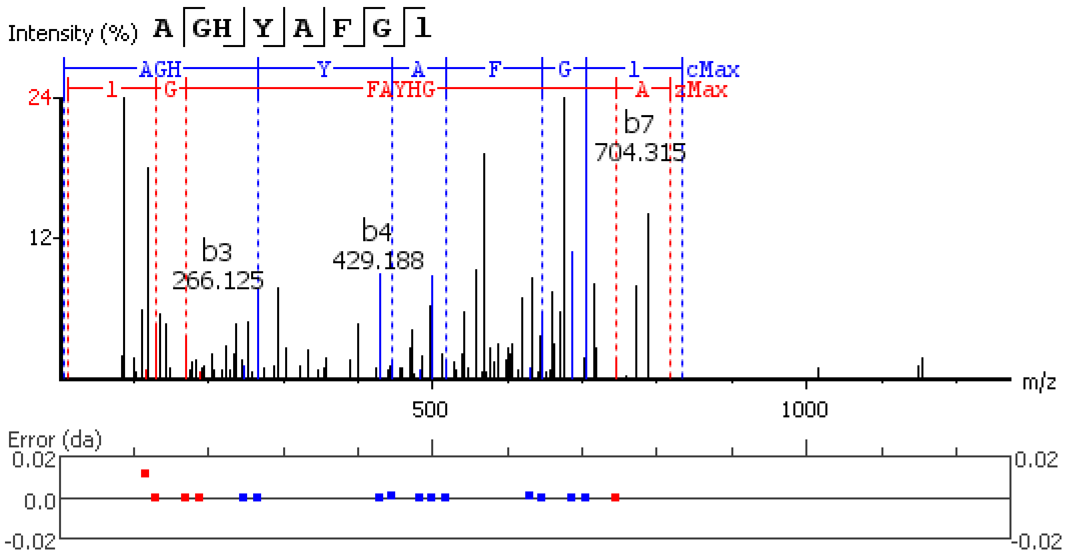


**Figure S5.8.** MS/MS spectrum of a AST-A neuropeptide with the sequence TPGYAFGL(-.98) detected in the peptidergic signaling system of the American lobster *Homarus americanus* generated by PEAKS XPro software.

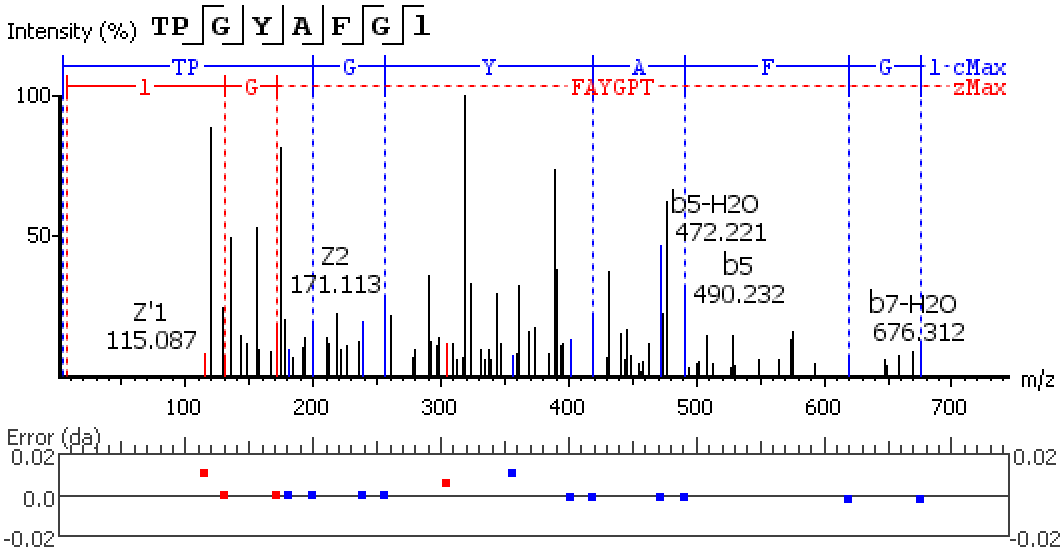


**Figure S5.9.** MS/MS spectrum of a AST-A neuropeptide with the sequence AGQYSFGL(-.98) detected in the peptidergic signaling system of the American lobster *Homarus americanus* generated by PEAKS XPro software.

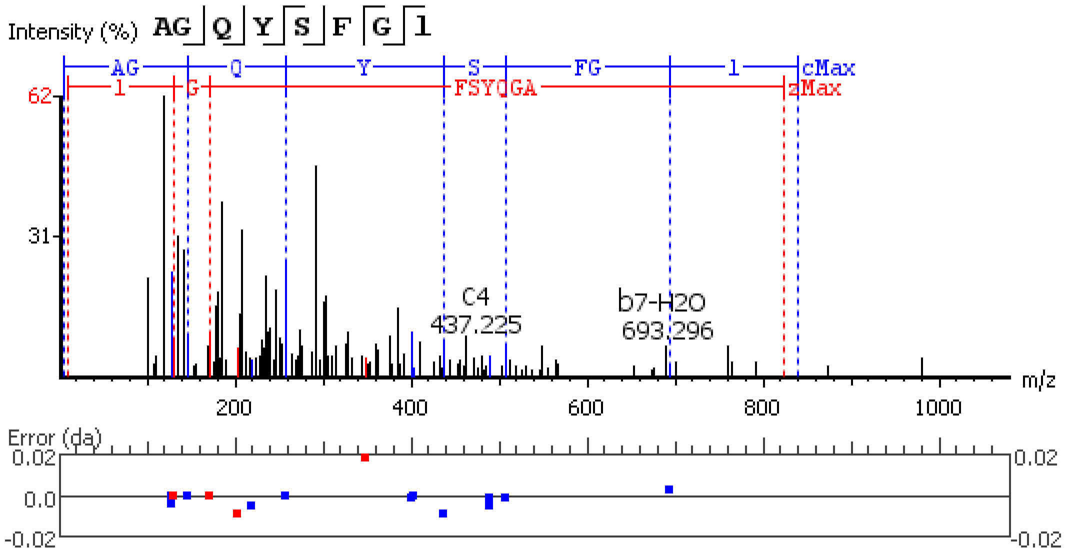


**Figure S5.10.** MS/MS spectrum of a AST-A neuropeptide with the sequence SDAPDSGFGRRSYDFGL(-.98) detected in the peptidergic signaling system of the American lobster *Homarus american*us generated by PEAKS XPro software.

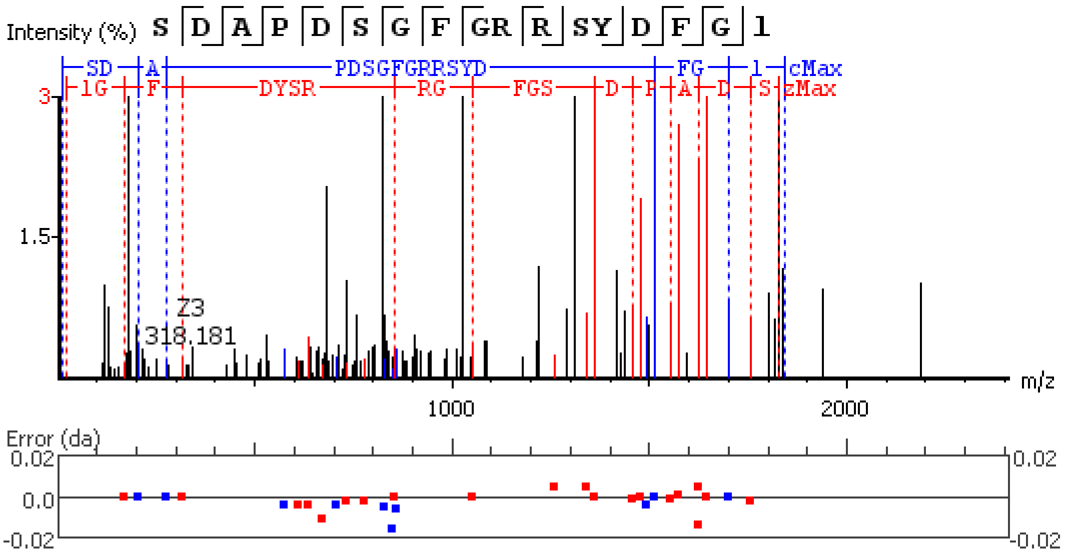


**Figure S5.11.** MS/MS spectrum of a AST-A neuropeptide with the sequence TGPYAFGL(-.98) detected in the peptidergic signaling system of the American lobster *Homarus american*us generated by PEAKS XPro software.

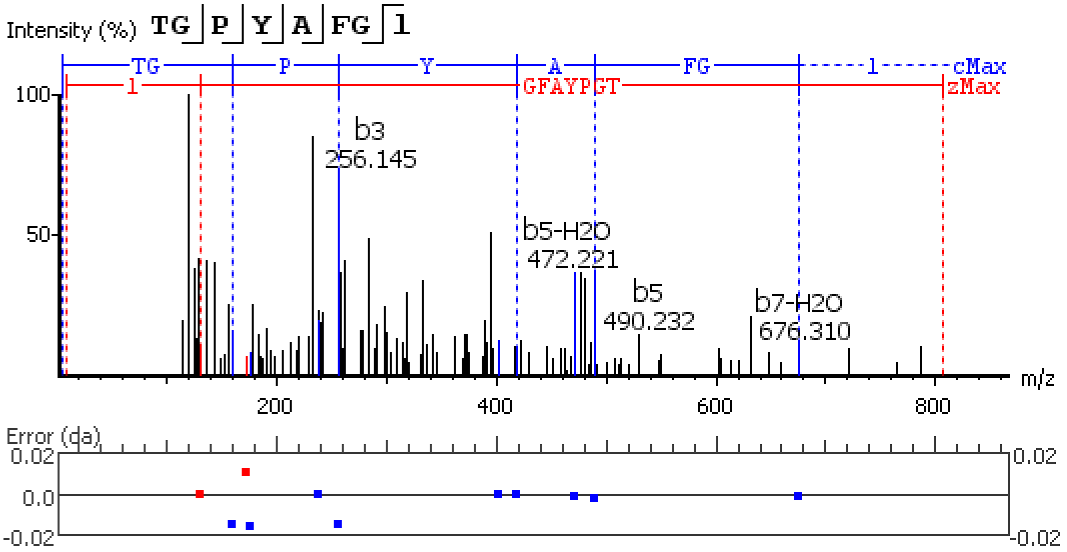


**Figure S5.12.** MS/MS spectrum of a AST-A neuropeptide with the sequence SDLYSFGL(-.98) detected in the peptidergic signaling system of the American lobster *Homarus american*us generated by PEAKS XPro software.

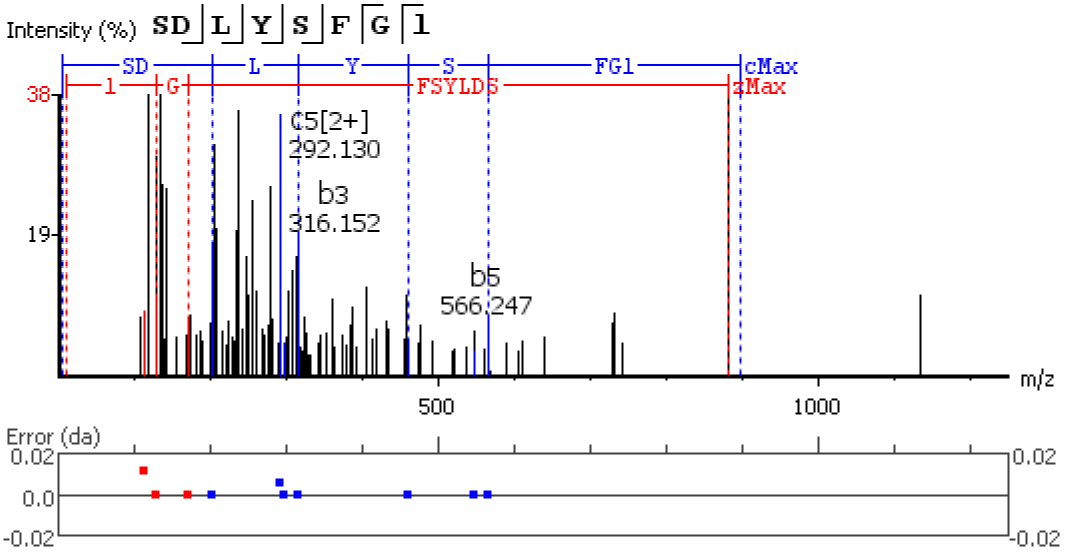


**Figure S5.13.** MS/MS spectrum of a AST-A neuropeptide with the sequence SGSYNFGL(-.98) detected in the peptidergic signaling system of the American lobster *Homarus american*us generated by PEAKS XPro software.

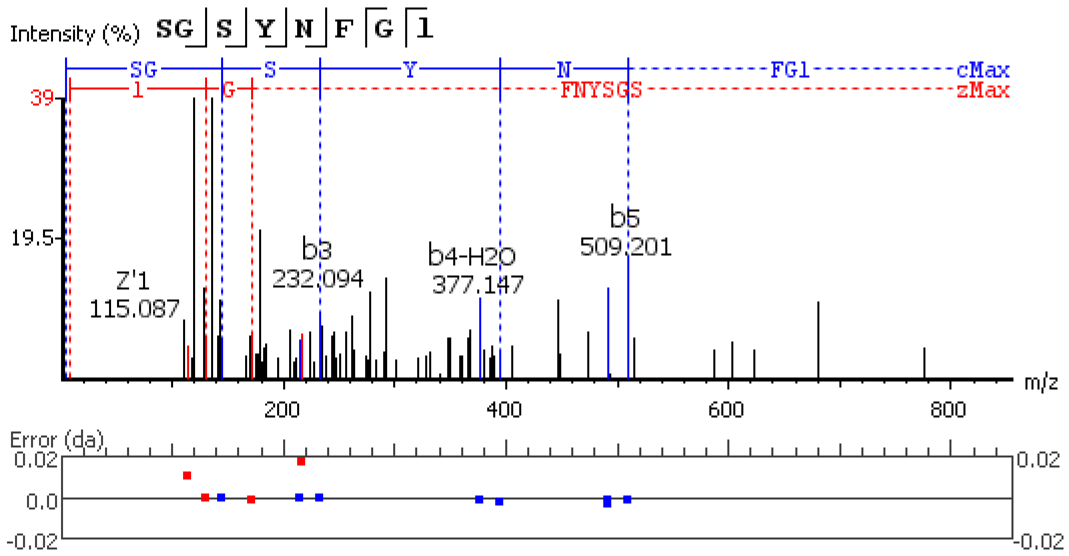


**Figure S5.14.** MS/MS spectrum of a AST-A neuropeptide with the sequence SQMYSFGL(-.98) detected in the peptidergic signaling system of the American lobster *Homarus american*us generated by PEAKS XPro software.

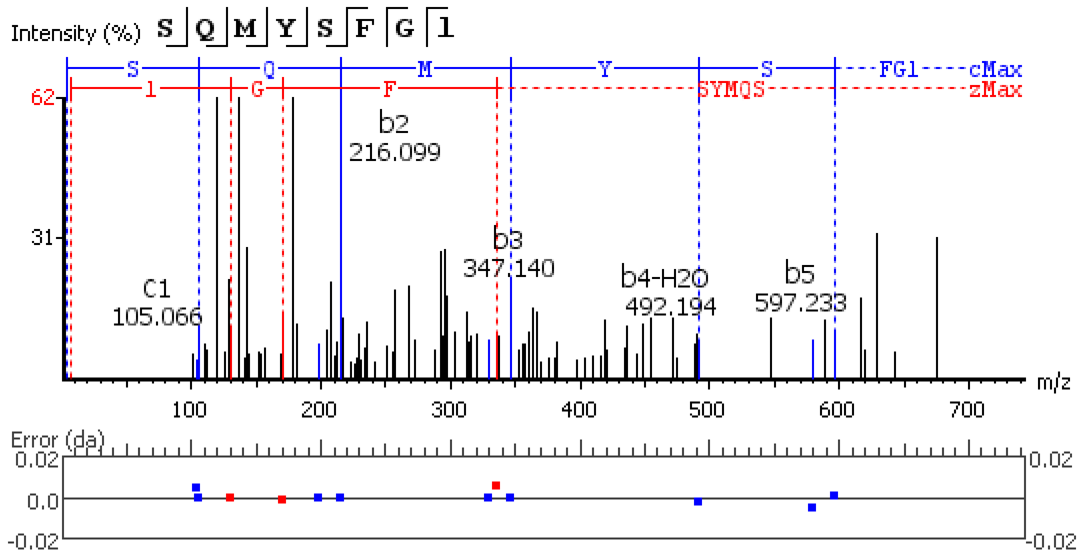


**Figure S5.15.** MS/MS spectrum of a AST-A neuropeptide with the sequence SQM(+15.99)YSFGL(-.98) detected in the peptidergic signaling system of the American lobster *Homarus american*us generated by PEAKS XPro software.

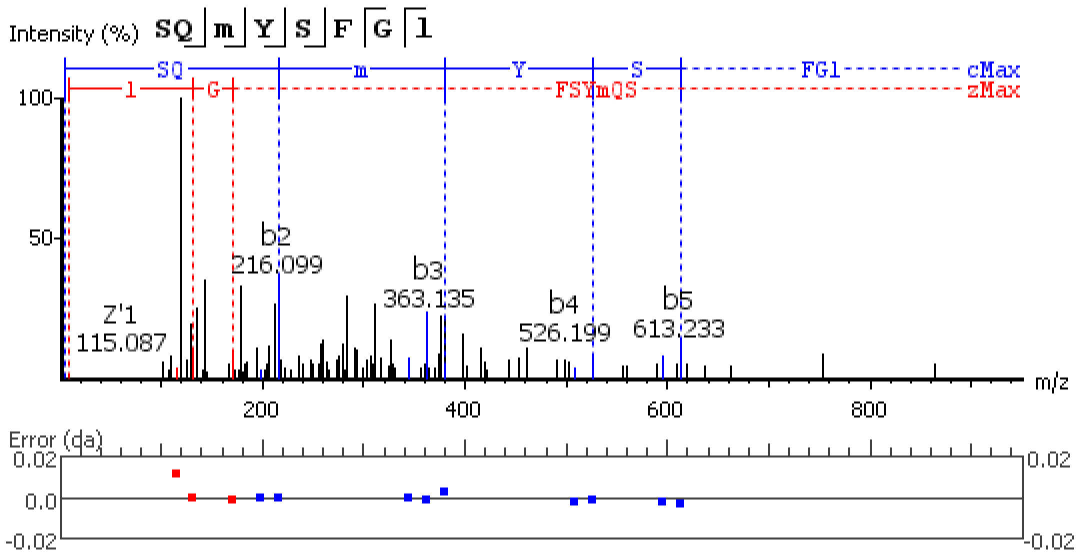


**Figure S5.16.** MS/MS spectrum of a AST-A neuropeptide with the sequence PTAYSFGL(-.98) detected in the peptidergic signaling system of the American lobster *Homarus american*us generated by PEAKS XPro software.

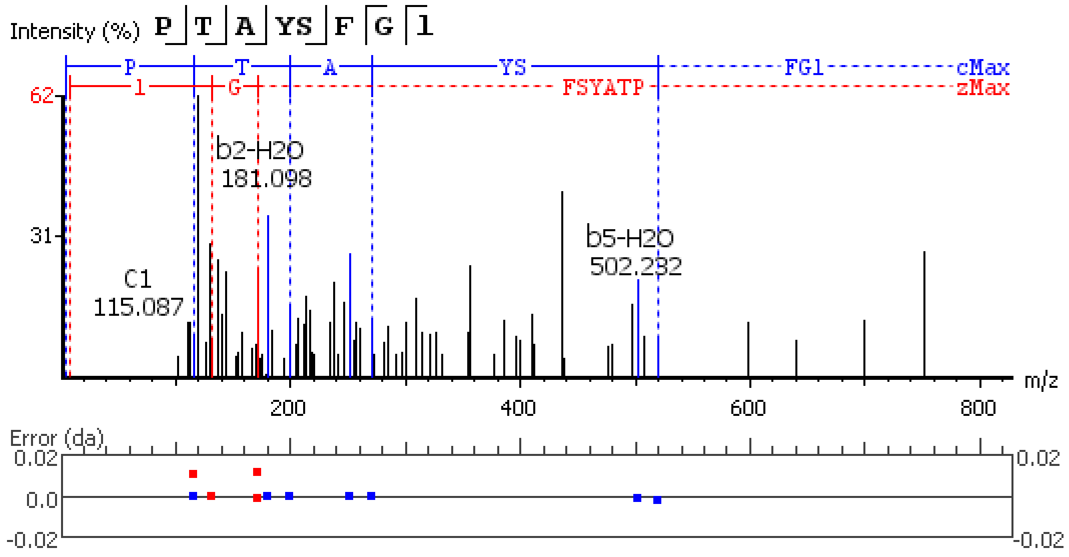


**Figure S5.17.** MS/MS spectrum of a AST-A neuropeptide with the sequence ADPYAFGL(-.98) detected in the peptidergic signaling system of the American lobster *Homarus american*us generated by PEAKS XPro software.

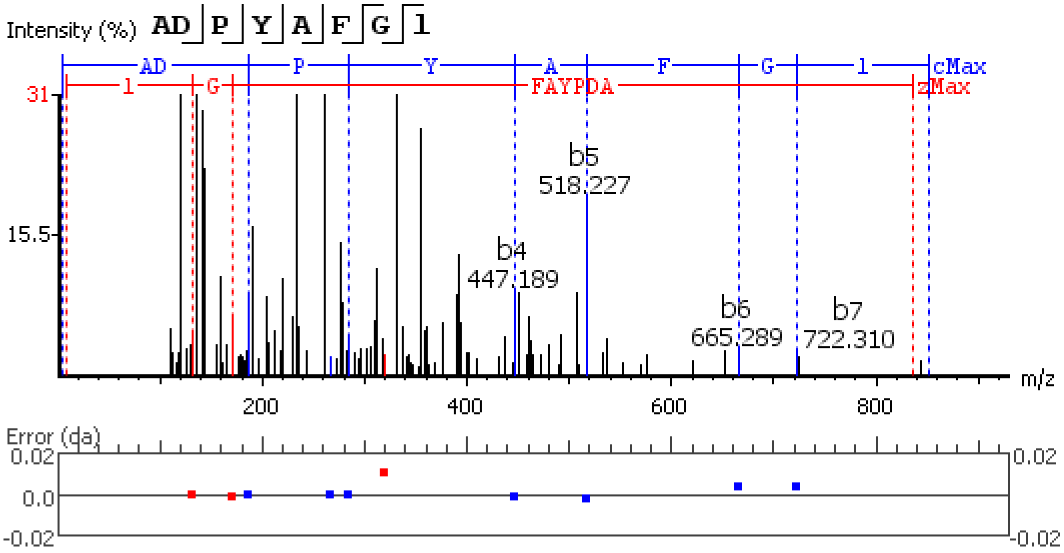


**Figure S5.18.** MS/MS spectrum of a AST-A neuropeptide with the sequence SDSDSDQYTL(-.98) detected in the peptidergic signaling system of the American lobster *Homarus american*us generated by PEAKS XPro software.

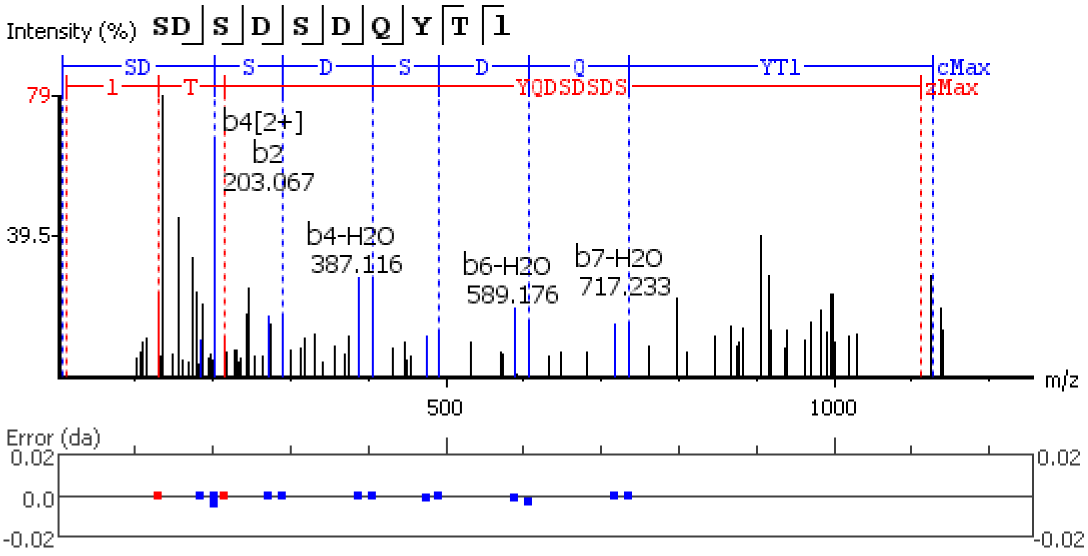


**Figure S5.19.** MS/MS spectrum of a AST-B neuropeptide with the sequence SSSSPQQDDPASSPSHIEE detected in the peptidergic signaling system of the American lobster *Homarus american*us generated by PEAKS XPro software.

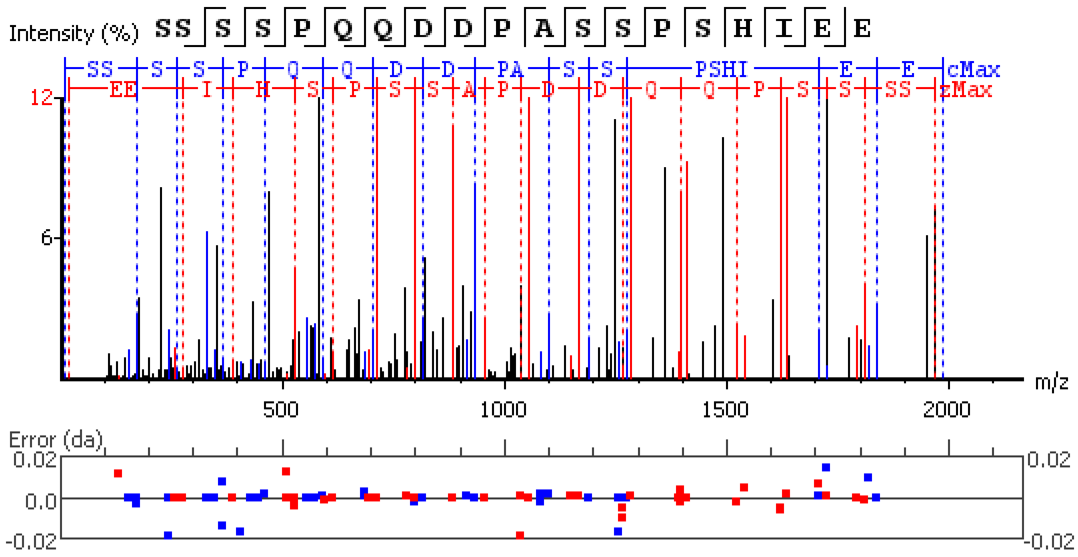


**Figure S5.20.** MS/MS spectrum of a AST-B neuropeptide with the sequence VGWSSMHGTW(-.98) detected in the peptidergic signaling system of the American lobster *Homarus american*us generated by PEAKS XPro software.

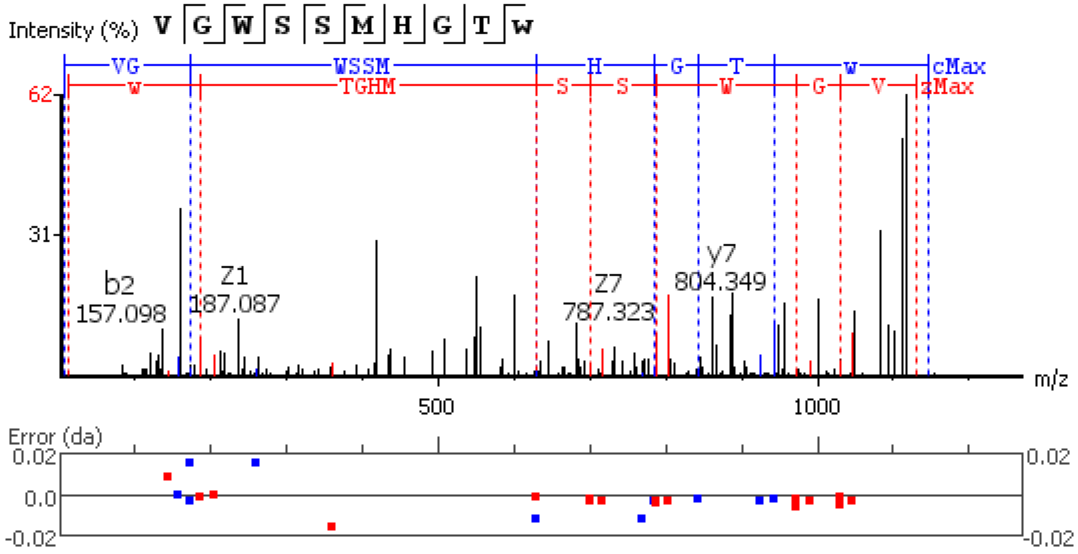


**Figure S5.21.** MS/MS spectrum of a AST-B neuropeptide with the sequence VGWSSM(+15.99)HGTW(-.98) detected in the peptidergic signaling system of the American lobster *Homarus american*us generated by PEAKS XPro software.

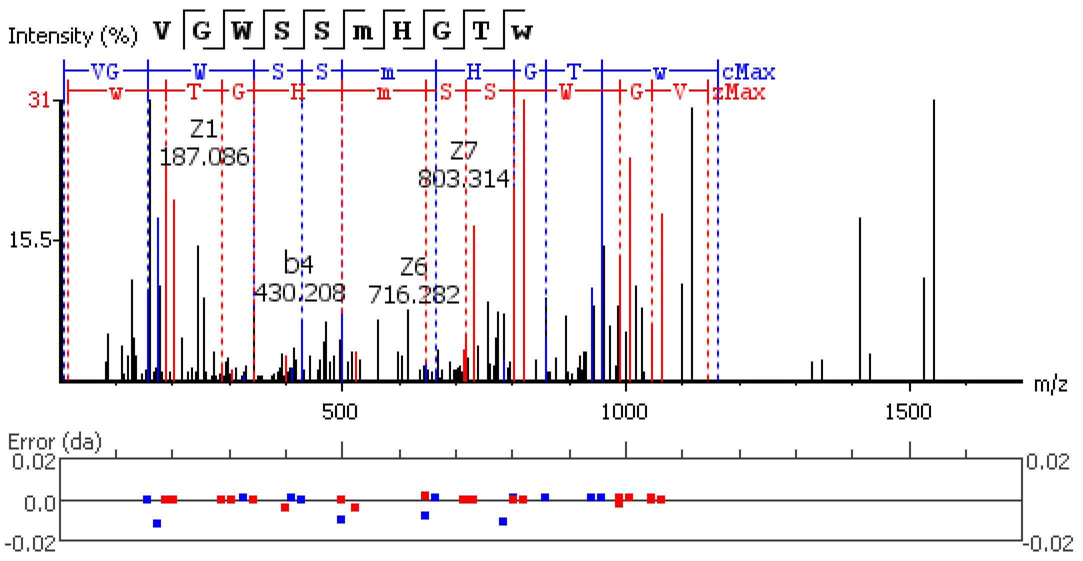


**Figure S5.22.** MS/MS spectrum of a AST-B neuropeptide with the sequence PHLEDAQLDAAEV detected in the peptidergic signaling system of the American lobster *Homarus americanus* generated by PEAKS XPro software.

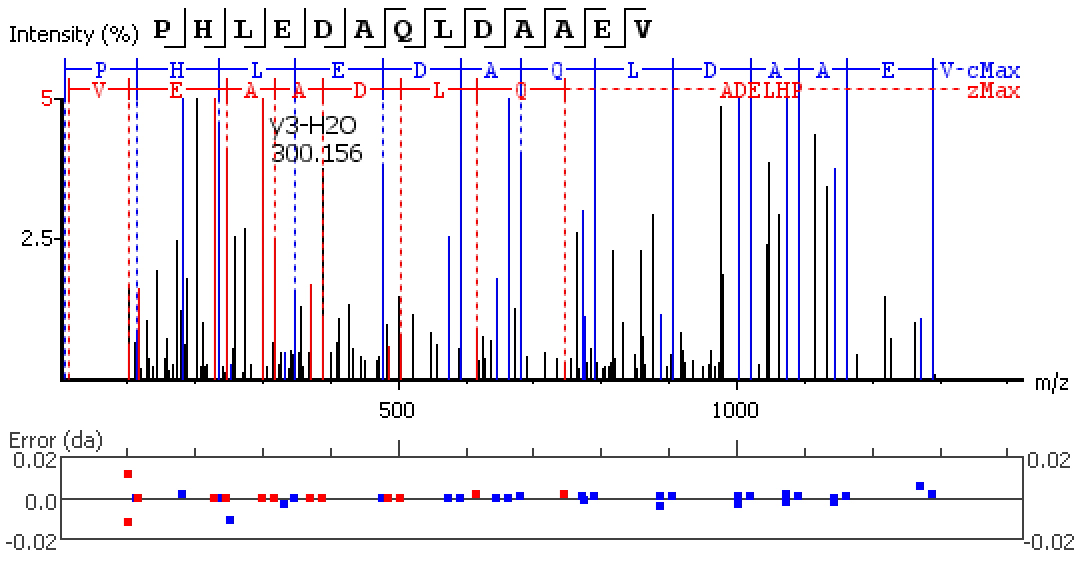


**Figure S5.23.** MS/MS spectrum of a AST-B neuropeptide with the sequence GTWGKRSADWNKL detected in the peptidergic signaling system of the American lobster *Homarus americanus* generated by PEAKS XPro software.

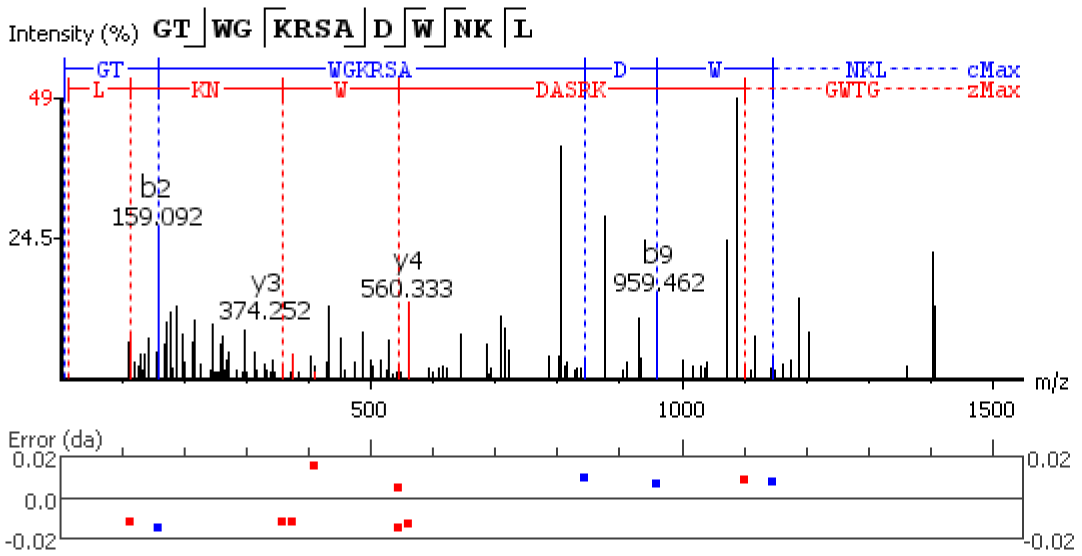


**Figure S5.24.** MS/MS spectrum of a AST-B neuropeptide with the sequence GEELQAAED detected in the peptidergic signaling system of the American lobster *Homarus americanu*s generated by PEAKS XPro software.

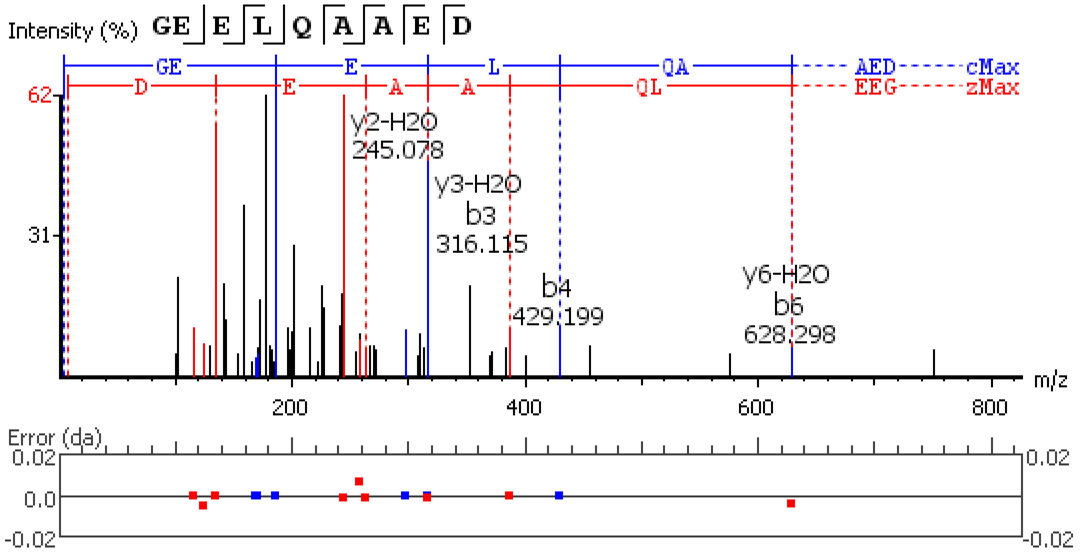


**Figure S5.25.** MS/MS spectrum of a AST-B neuropeptide with the sequence TNWNKFQGSW(-.98) detected in the peptidergic signaling system of the American lobster *Homarus americanu*s generated by PEAKS XPro software.

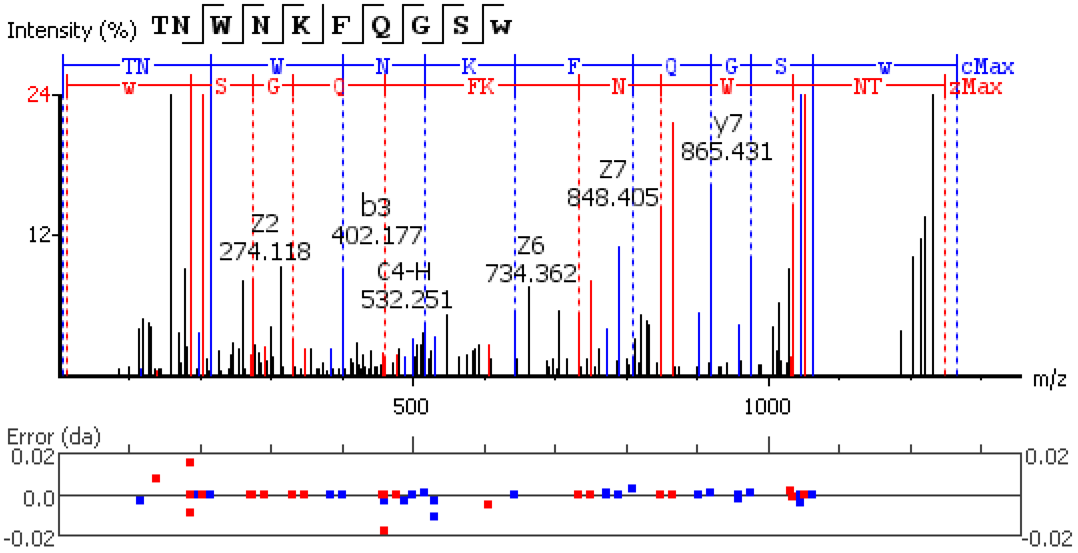


**Figure S5.26.** MS/MS spectrum of a AST-B neuropeptide with the sequence NNWRSLQGSW(-.98) detected in the peptidergic signaling system of the American lobster *Homarus americanus* generated by PEAKS XPro software.

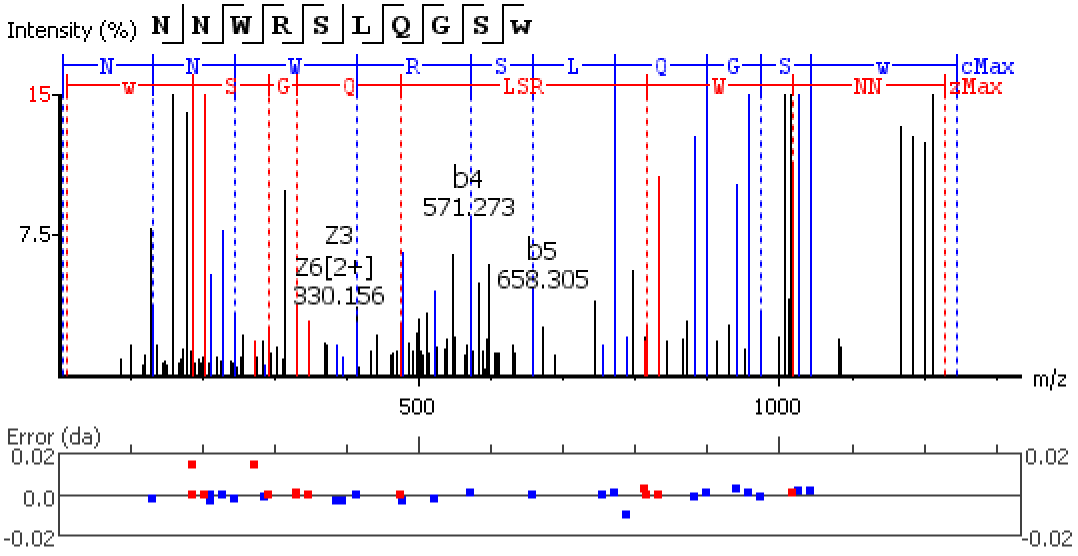


**Figure S5.27.** MS/MS spectrum of a AST-B neuropeptide with the sequence AWNKLQGAW(-.98) detected in the peptidergic signaling system of the American lobster *Homarus americanus* generated by PEAKS XPro software.

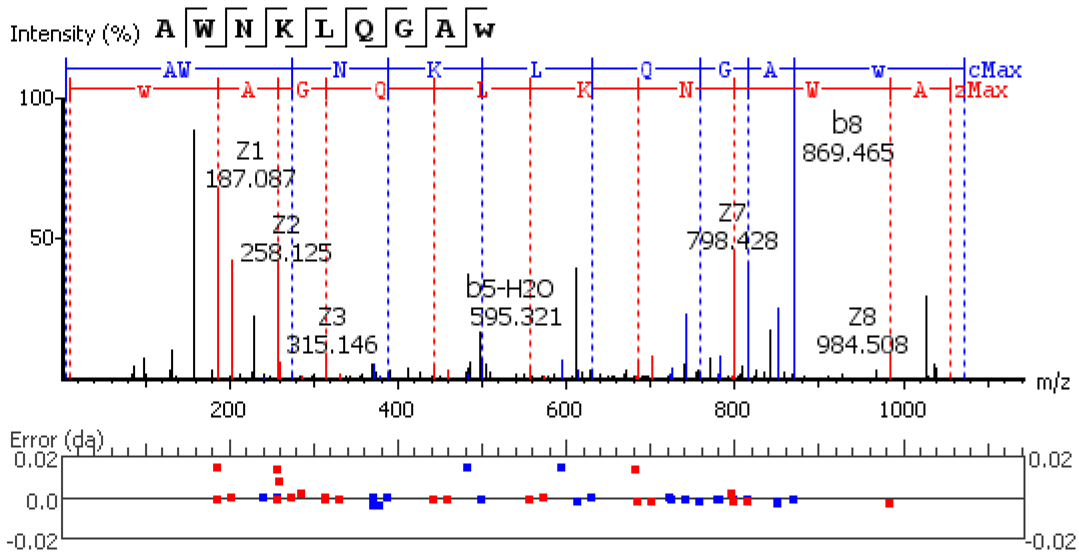


**Figure S5.28.** MS/MS spectrum of a AST-B neuropeptide with the sequence STNWSSLRGTW(-.98) detected in the peptidergic signaling system of the American lobster *Homarus american*us generated by PEAKS XPro software.

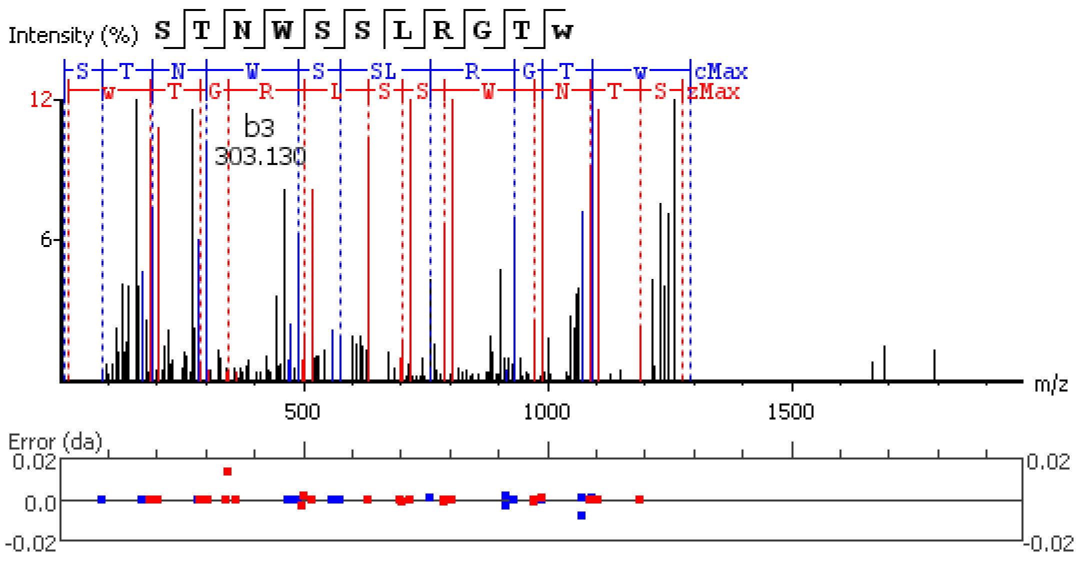


**Figure S5.29.** MS/MS spectrum of a AST-B neuropeptide with the sequence SADWNKLRGAW(-.98) detected in the peptidergic signaling system of the American lobster *Homarus american*us generated by PEAKS XPro software.

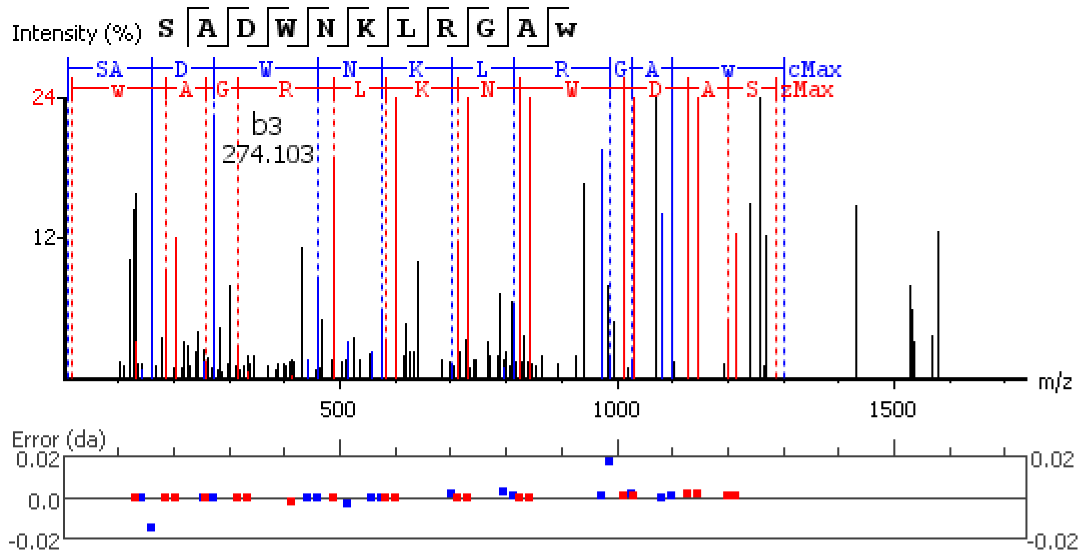


**Figure S5.30.** MS/MS spectrum of a AST-B neuropeptide with the sequence ASDWGQFRGSW(-.98) detected in the peptidergic signaling system of the American lobster *Homarus american*us generated by PEAKS XPro software.

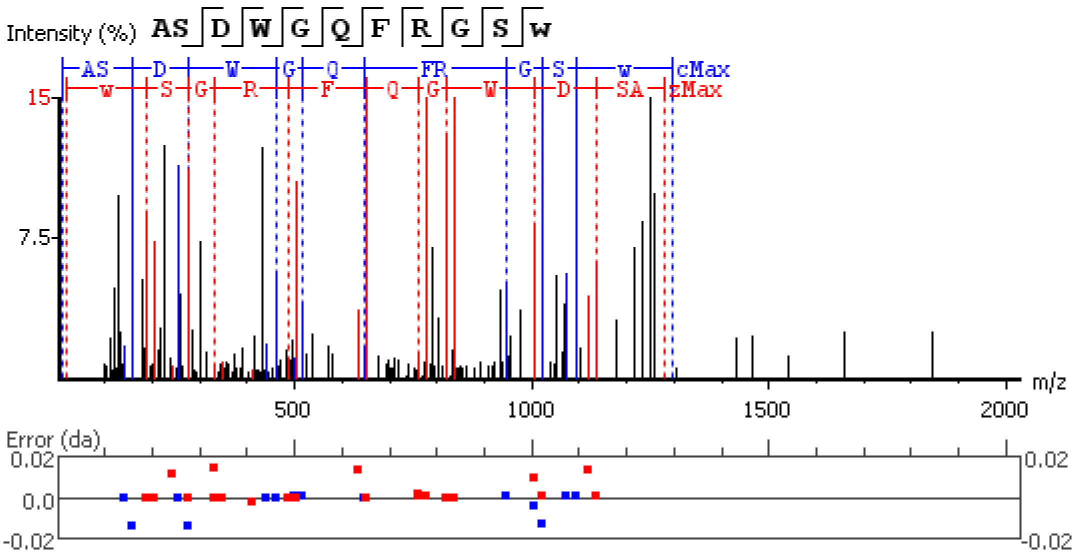


**Figure S5.31.** MS/MS spectrum of a AST-B neuropeptide with the sequence APDMMSVAAPNQA detected in the peptidergic signaling system of the American lobster *Homarus american*us generated by PEAKS XPro software.

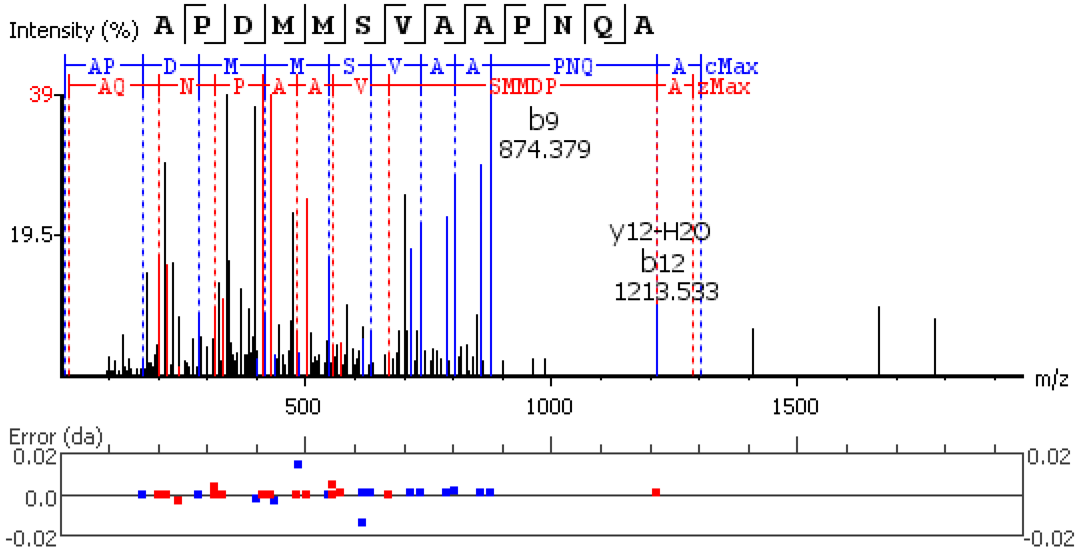


**Figure S5.32.** MS/MS spectrum of a AST-B neuropeptide with the sequence APDM(+15.99)MSVAAPNQA detected in the peptidergic signaling system of the American lobster *Homarus american*us generated by PEAKS XPro software.

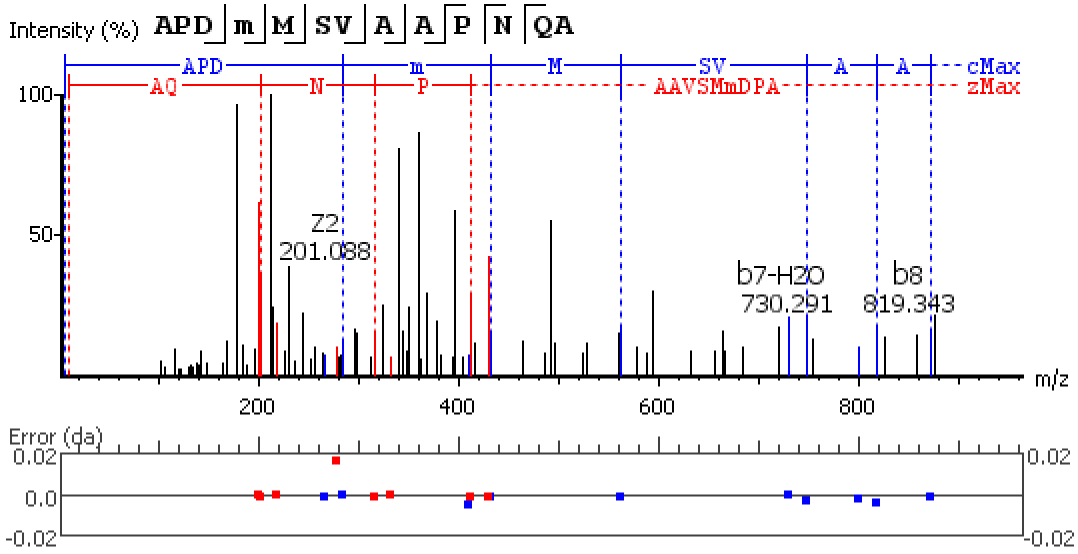


**Figure S5.33.** MS/MS spectrum of a AST-B neuropeptide with the sequence APDMM(+15.99)SVAAPNQA detected in the peptidergic signaling system of the American lobster *Homarus american*us generated by PEAKS XPro software.

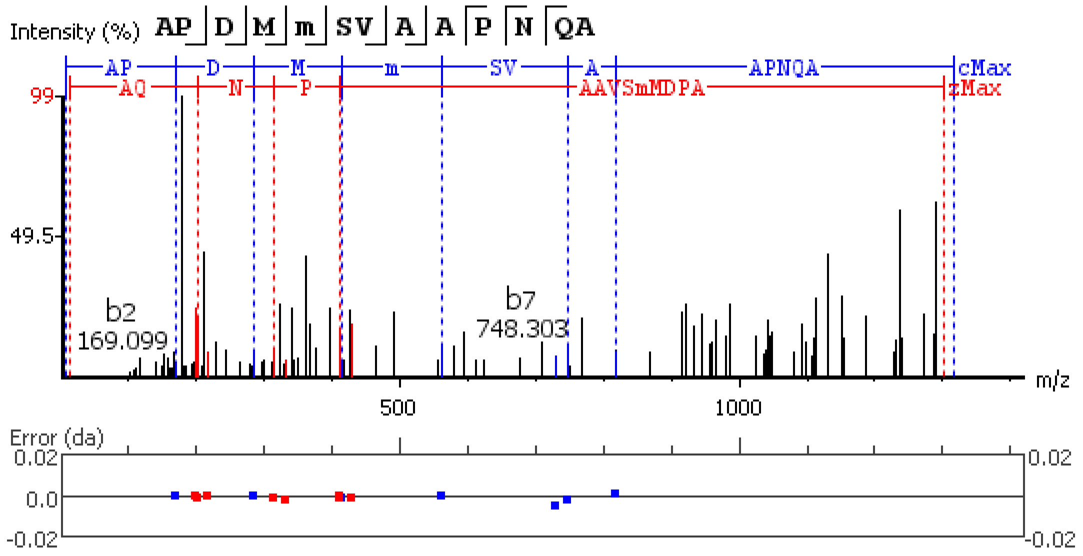


**Figure S5.34.** MS/MS spectrum of a AST-B neuropeptide with the sequence APDM(+15.99)M(+15.99)SVAAPNQA detected in the peptidergic signaling system of the American lobster *Homarus american*us generated by PEAKS XPro software.

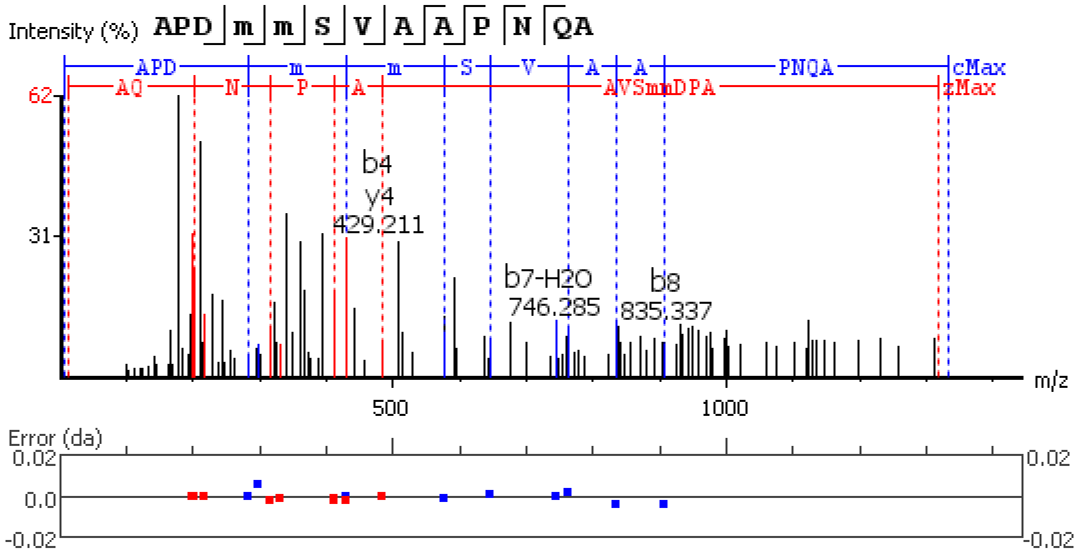


**Figure S5.35.** MS/MS spectrum of a AST-C neuropeptide with the sequence NNADIKDLQ detected in the peptidergic signaling system of the American lobster *Homarus american*us generated by PEAKS XPro software.

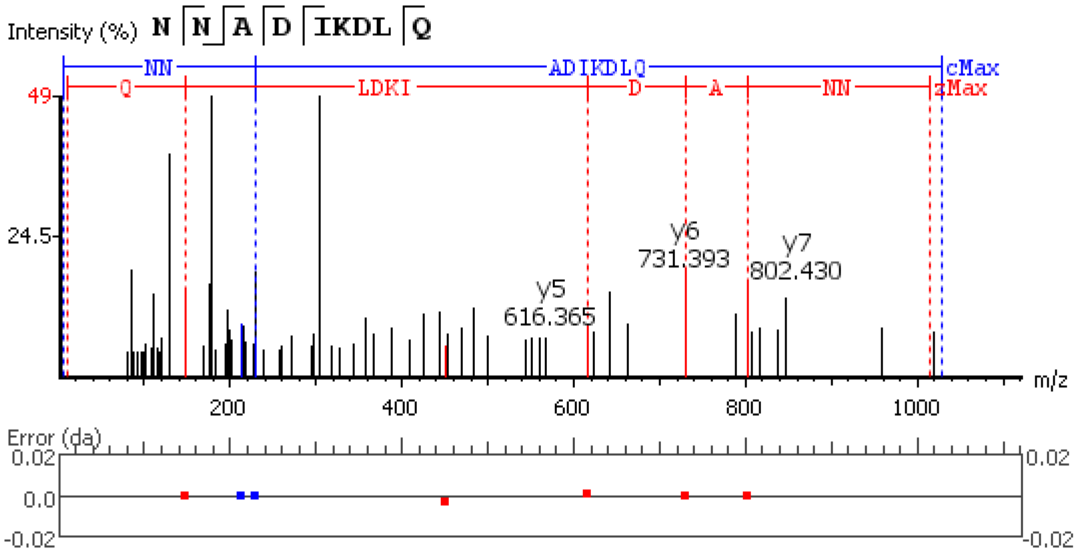


**Figure S5.36.** MS/MS spectrum of a CHH-A neuropeptide with the sequence RSVEGASRMEKLLSSSNSPSSTPLGFLSQDHSVN detected in the peptidergic signaling system of the American lobster *Homarus american*us generated by PEAKS XPro software.

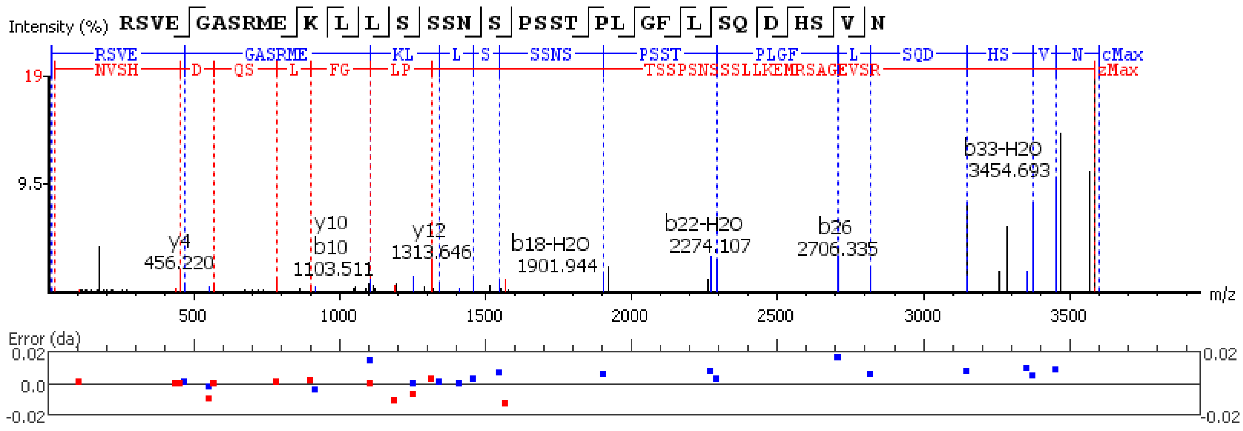


**Figure S5.37.** MS/MS spectrum of a CHH-A neuropeptide with the sequence RSVEGASRM(+15.99)EKLLSSSNSPSSTPLGFLSQDHSVN detected in the peptidergic signaling system of the American lobster *Homarus american*us generated by PEAKS XPro software.

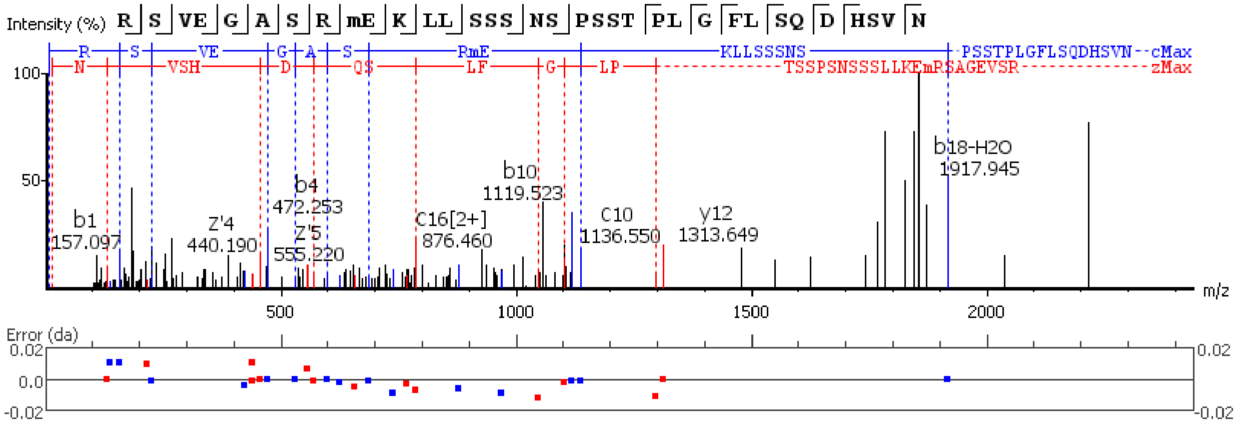

**Figure S5.38.** MS/MS spectrum of a CHH-B neuropeptide with the sequence RSVEGVSRMEKLLSSISPSSTPLGFLSQDHSVN detected in the peptidergic signaling system of the American lobster *Homarus americanus* generated by PEAKS XPro software.

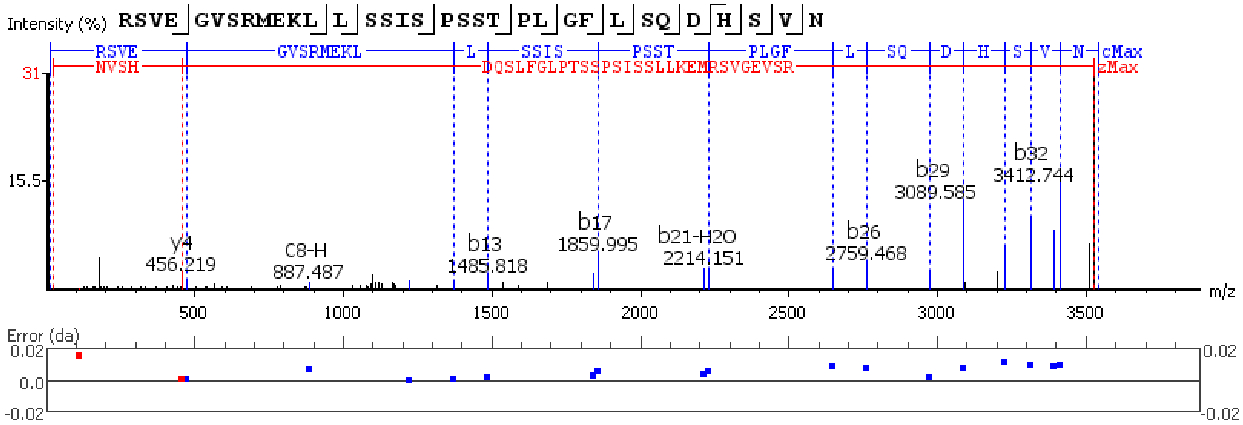


**Figure S5.39.** MS/MS spectrum of a Corazonin neuropeptide with the sequence Q(-17.03)TFQYSRGWTN(-.98) detected in the peptidergic signaling system of the American lobster *Homarus americanus* generated by PEAKS XPro software.

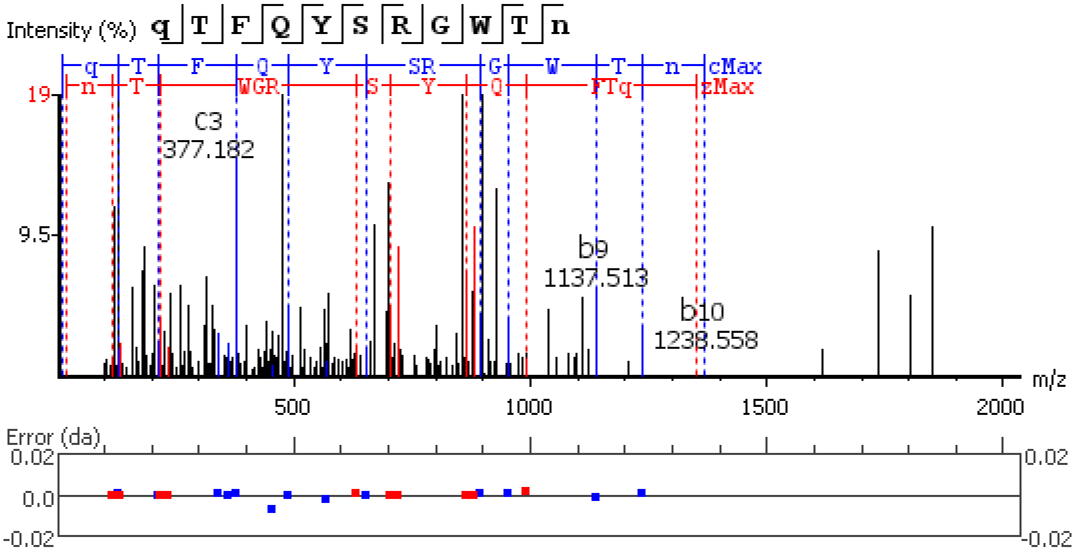


**Figure S5.40.** MS/MS spectrum of a crustacean cardioactive peptide (CCAP) neuropeptide with the sequence DIGDLLEGKD detected in the peptidergic signaling system of the American lobster *Homarus americanus* generated by PEAKS XPro software.

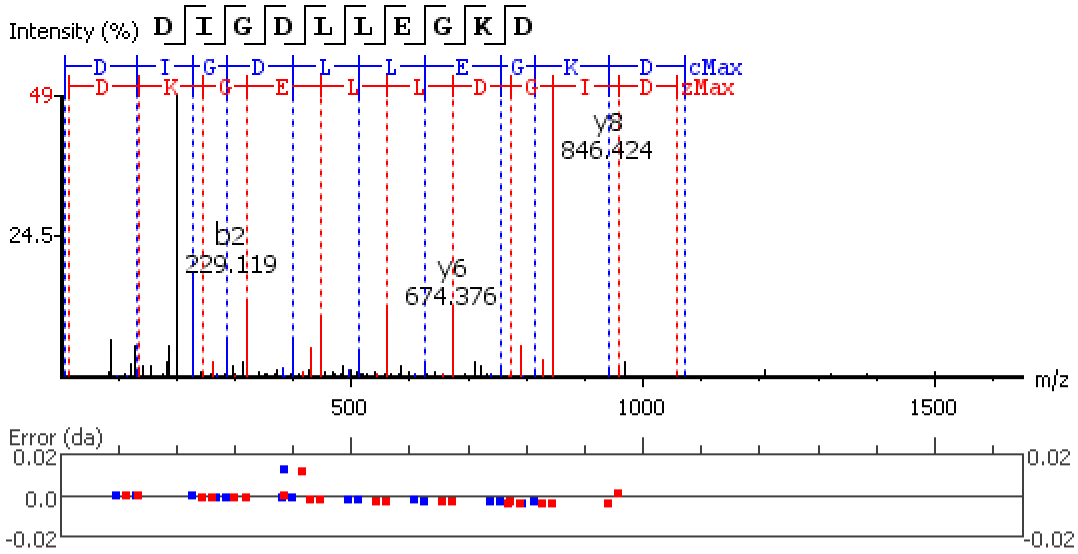


**Figure S5.41.** MS/MS spectrum of a crustacean cardioactive peptide (CCAP) neuropeptide with the sequence STPHTQPRQHLTSTPQQKVETEKQ detected in the peptidergic signaling system of the American lobster *Homarus american*us generated by PEAKS XPro software.

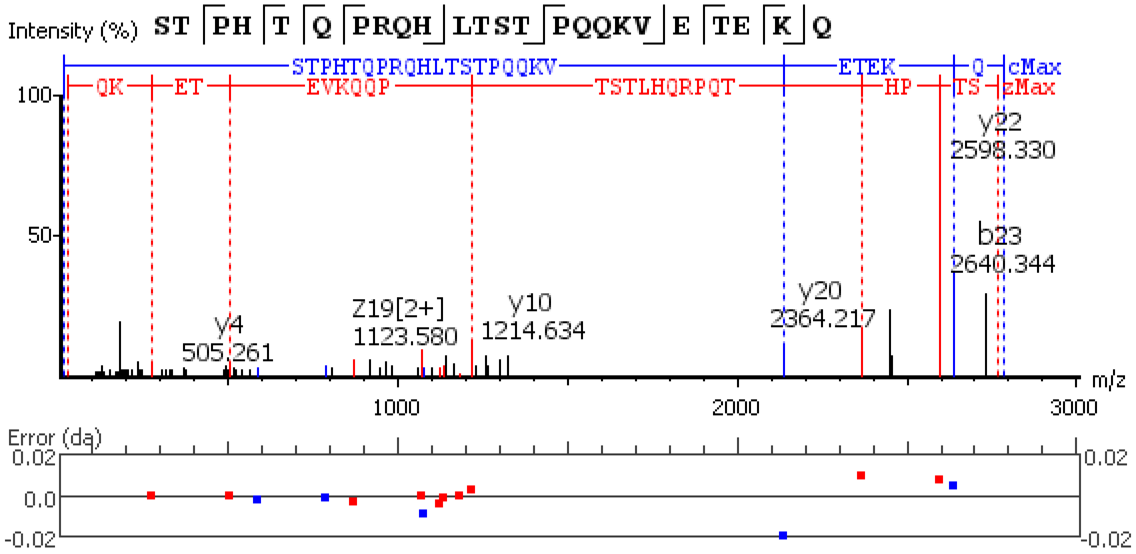


**Figure S5.42.** MS/MS spectrum of a Diuretic Hormone neuropeptide with the sequence GLDLGLGRGFSGSQAAKHLMGLAAANFAGGP(-.98) detected in the peptidergic signaling system of the American lobster *Homarus american*us generated by PEAKS XPro software.

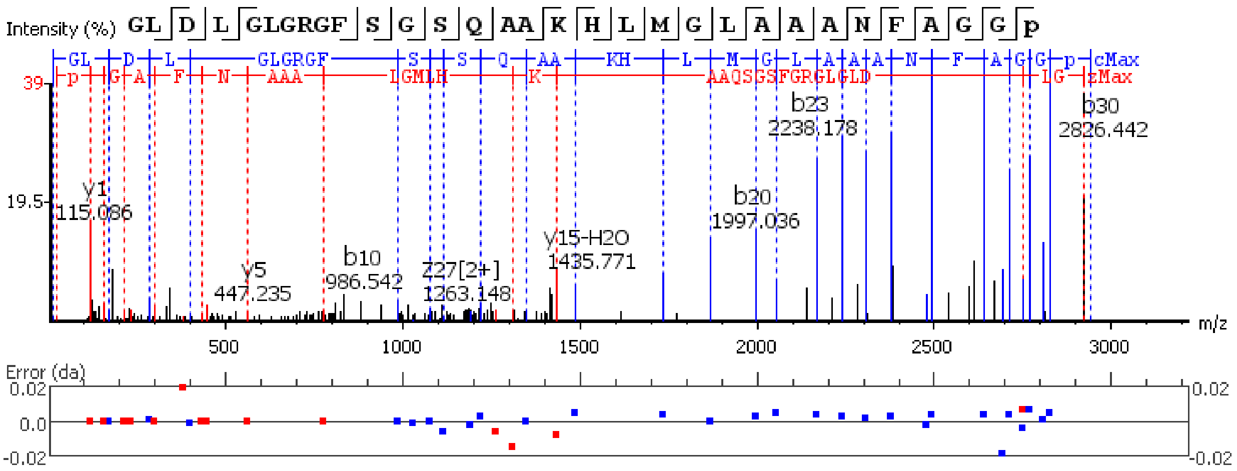


**Figure S5.43.** MS/MS spectrum of a Diuretic Hormone neuropeptide with the sequence GLDLGLGRGFSGSQAAKHLM(+15.99)GLAAANFAGGP(-.98) detected in the peptidergic signaling system of the American lobster *Homarus american*us generated by PEAKS XPro software.

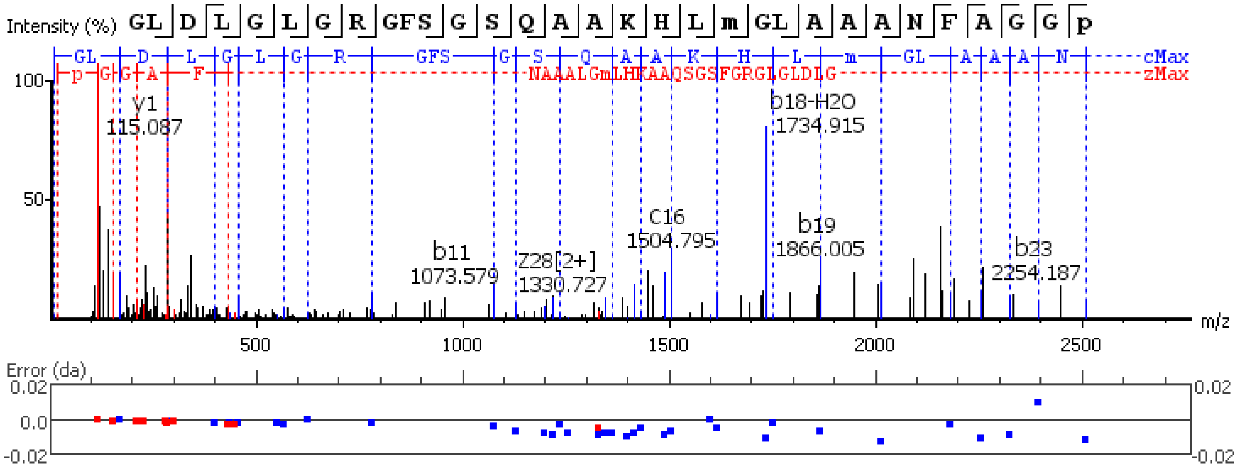


**Figure S5.44.** MS/MS spectrum of a Diuretic Hormone neuropeptide with the sequence SSDDGLDLHHDDNLYAQDQAADLAESS detected in the peptidergic signaling system of the American lobster *Homarus american*us generated by PEAKS XPro software.

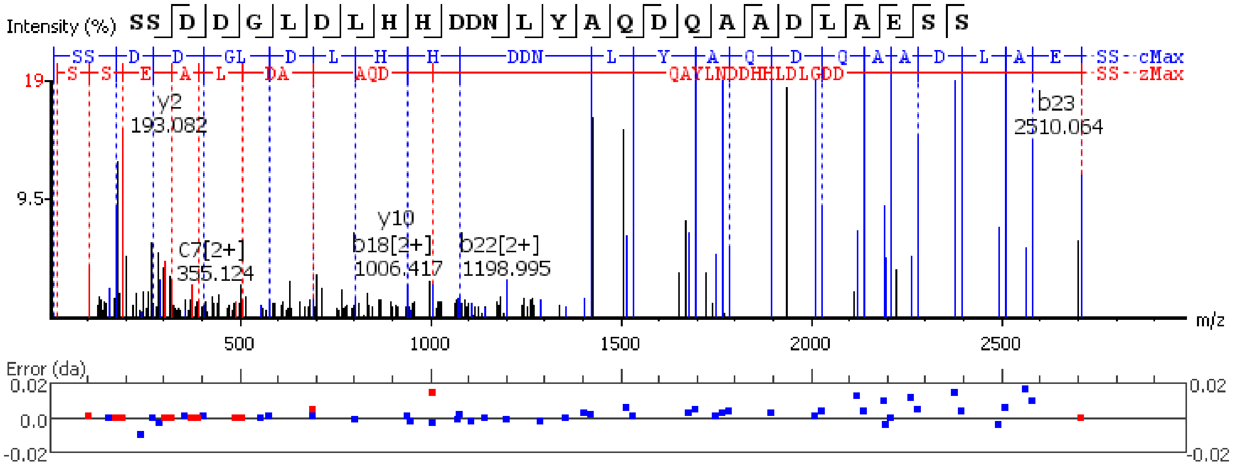

**Figure S5.45.** MS/MS spectrum of a FMRFamide neuropeptide with the sequence LLKYFLPASQAWGGDAYPIGQEGT detected in the peptidergic signaling system of the American lobster *Homarus american*us generated by PEAKS XPro software.

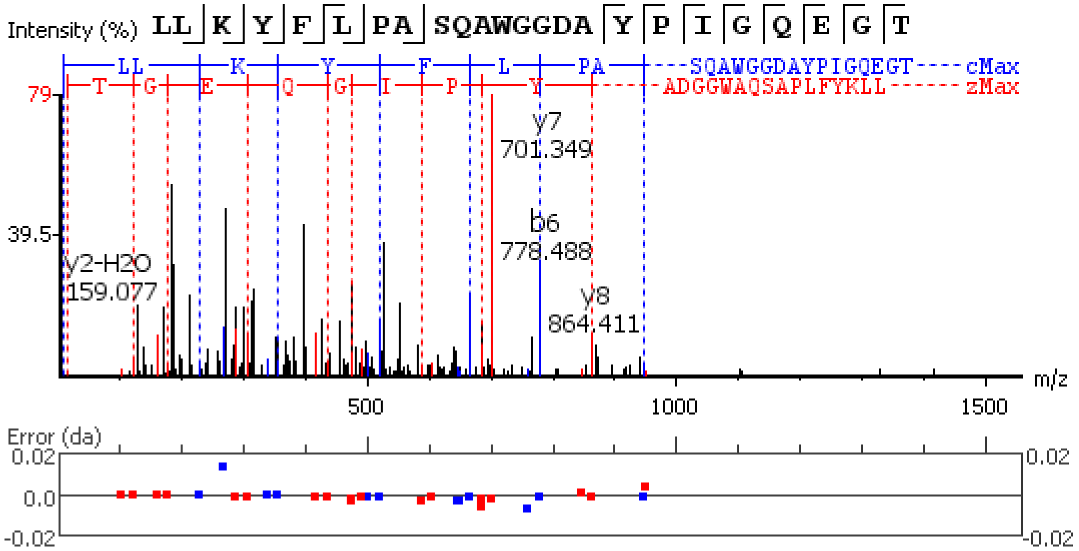


**Figure S5.46.** MS/MS spectrum of a FMRFamide neuropeptide with the sequence GYSDRNYLRF(-.98) detected in the peptidergic signaling system of the American lobster *Homarus american*us generated by PEAKS XPro software.

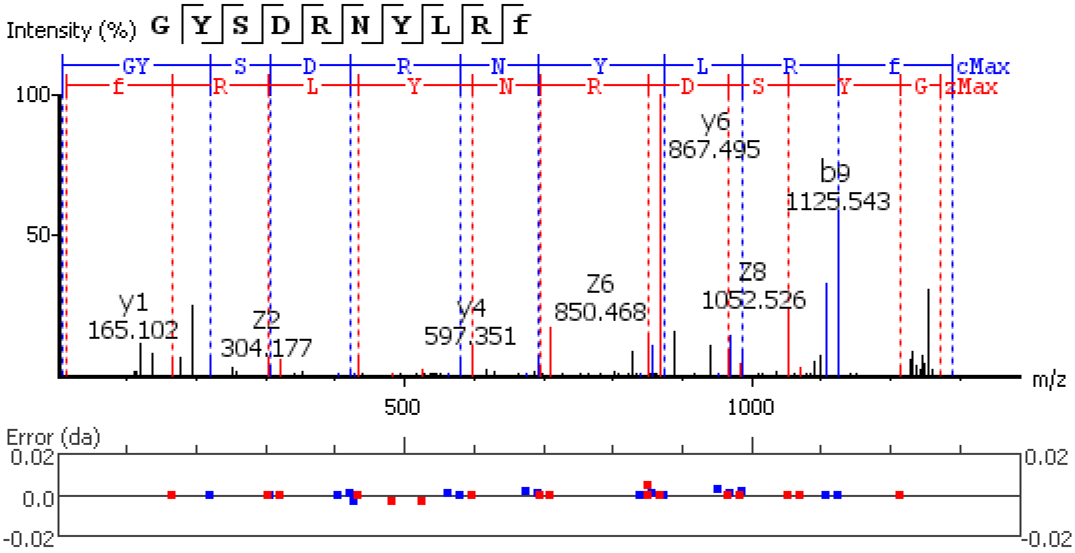


**Figure S5.47.** MS/MS spectrum of a FMRFamide neuropeptide with the sequence SDTNDYEGEEMPESPE detected in the peptidergic signaling system of the American lobster *Homarus american*us generated by PEAKS XPro software.

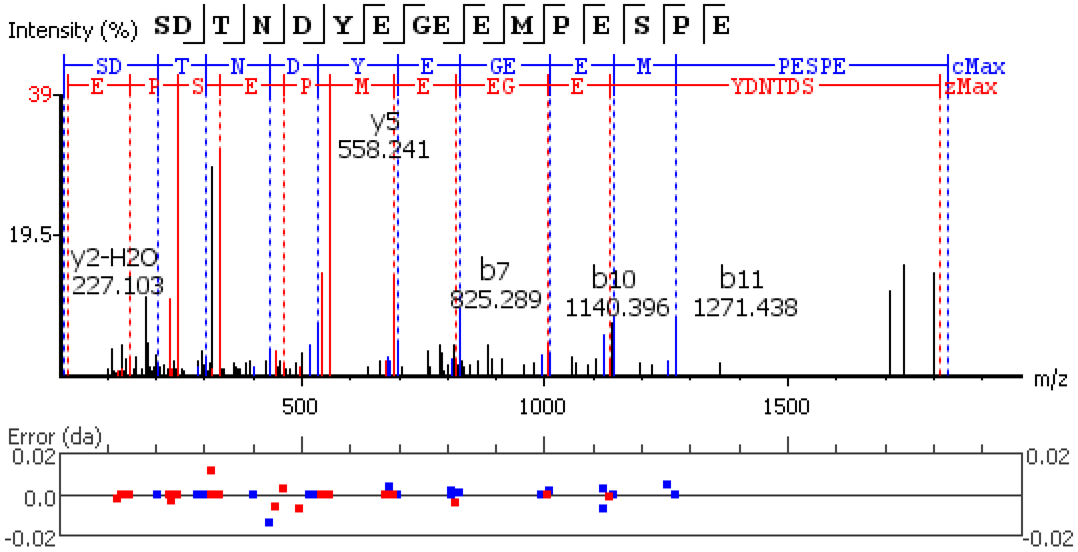


**Figure S5.48.** MS/MS spectrum of a FMRFamide neuropeptide with the sequence Q(-17.03)INAHRI detected in the peptidergic signaling system of the American lobster *Homarus americanus* generated by PEAKS XPro software.

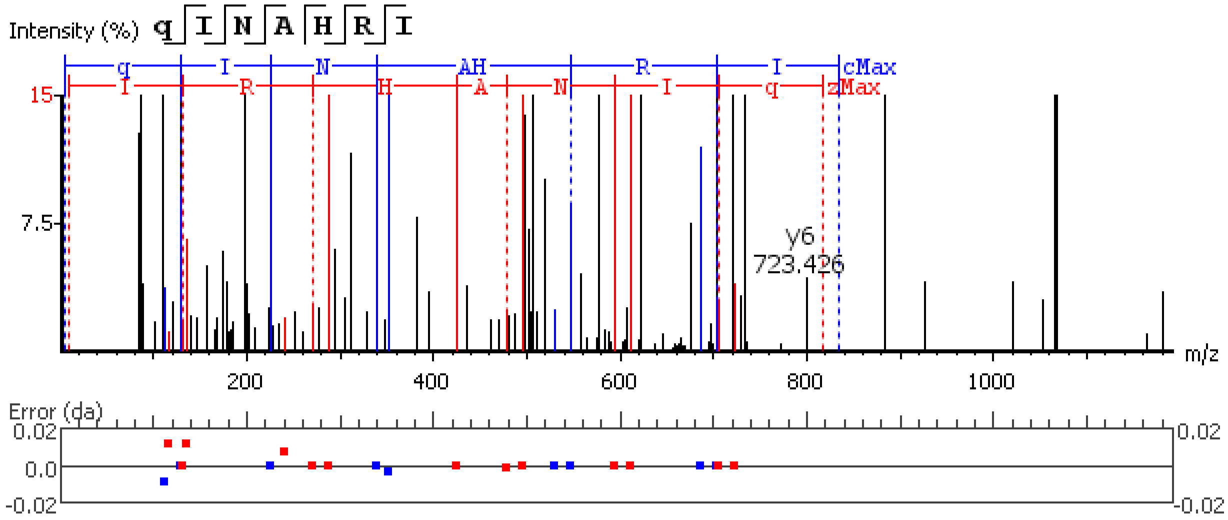


**Figure S5.49.** MS/MS spectrum of a FMRFamide neuropeptide with the sequence NFLRFGRS(-.98). detected in the peptidergic signaling system of the American lobster *Homarus americanus* generated by PEAKS XPro software.

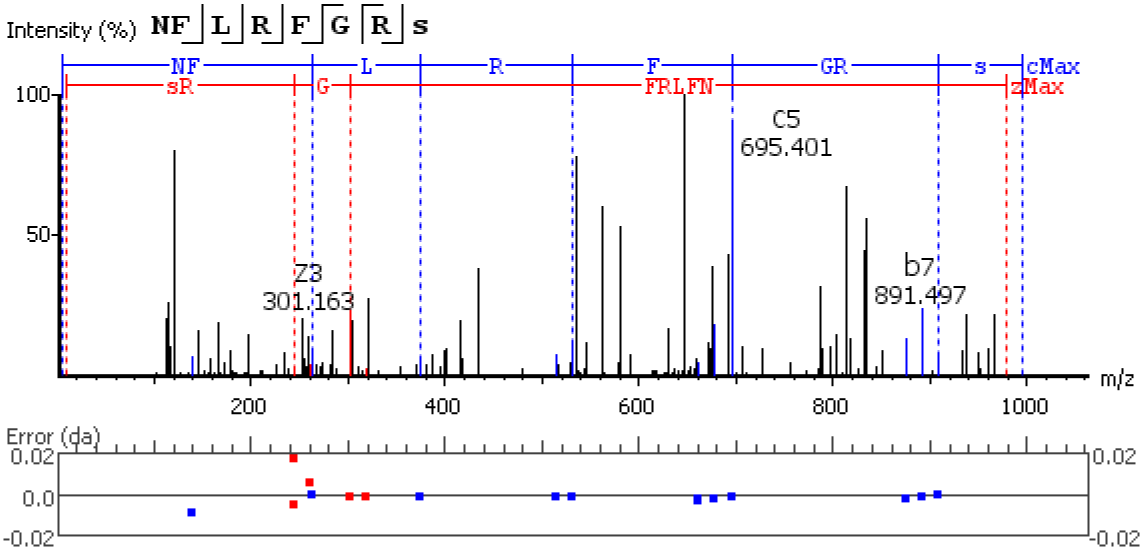


**Figure S5.50.** MS/MS spectrum of a FMRFamide neuropeptide with the sequence SDRNFLRF(-.98)

detected in the peptidergic signaling system of the American lobster *Homarus american*us generated by PEAKS XPro software.

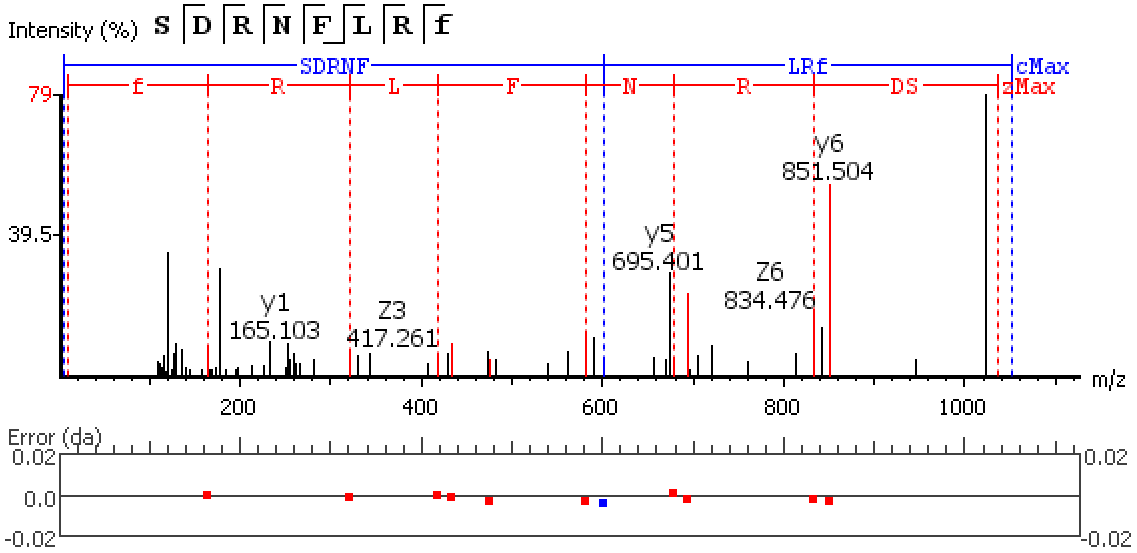


**Figure S5.51.** MS/MS spectrum of a FMRFamide neuropeptide with the sequence NRNFLRF(-.98)

detected in the peptidergic signaling system of the American lobster *Homarus american*us generated by PEAKS XPro software.

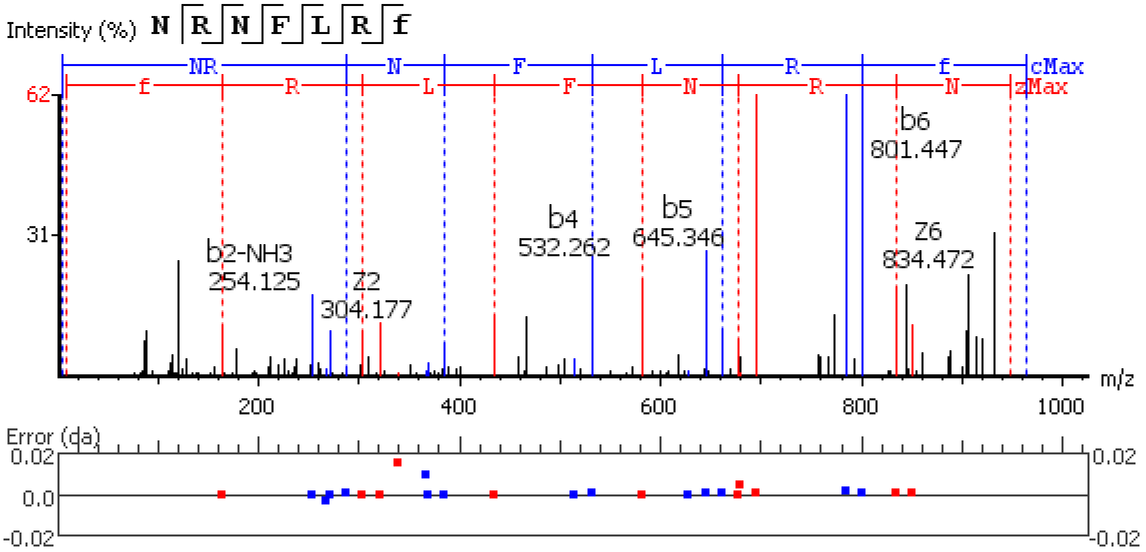


**Figure S5.52.** MS/MS spectrum of a FMRFamide neuropeptide with the sequence DQNRNFLRF(-.98)

detected in the peptidergic signaling system of the American lobster *Homarus american*us generated by PEAKS XPro software.

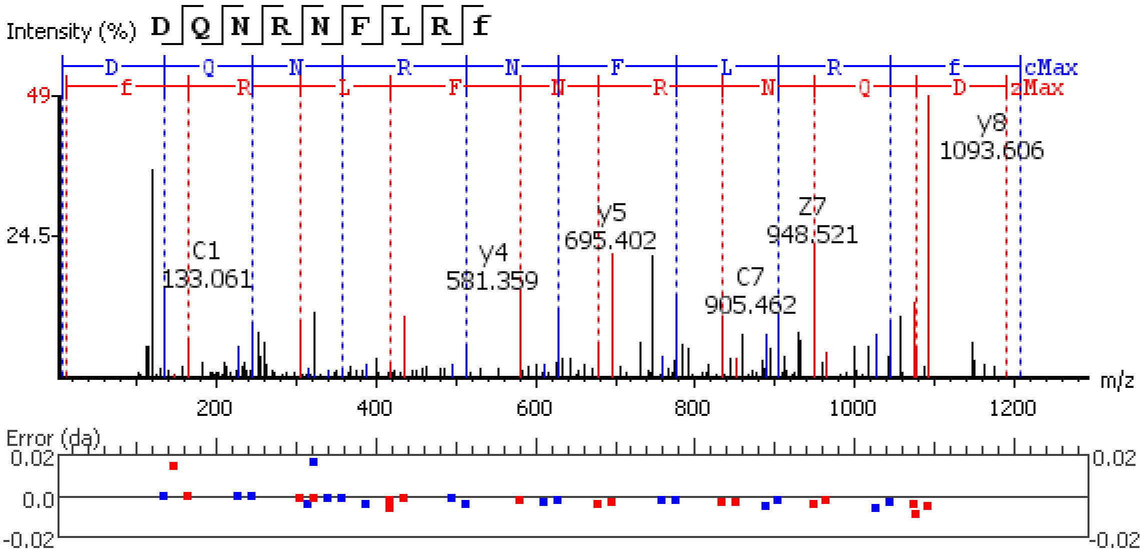


**Figure S5.53.** MS/MS spectrum of a FMRFamide neuropeptide with the sequence

SGSPMEFATDLQEDVELPVEE detected in the peptidergic signaling system of the American

lobster *Homarus american*us generated by PEAKS XPro software using data dependent acquisition (DDA)

method.

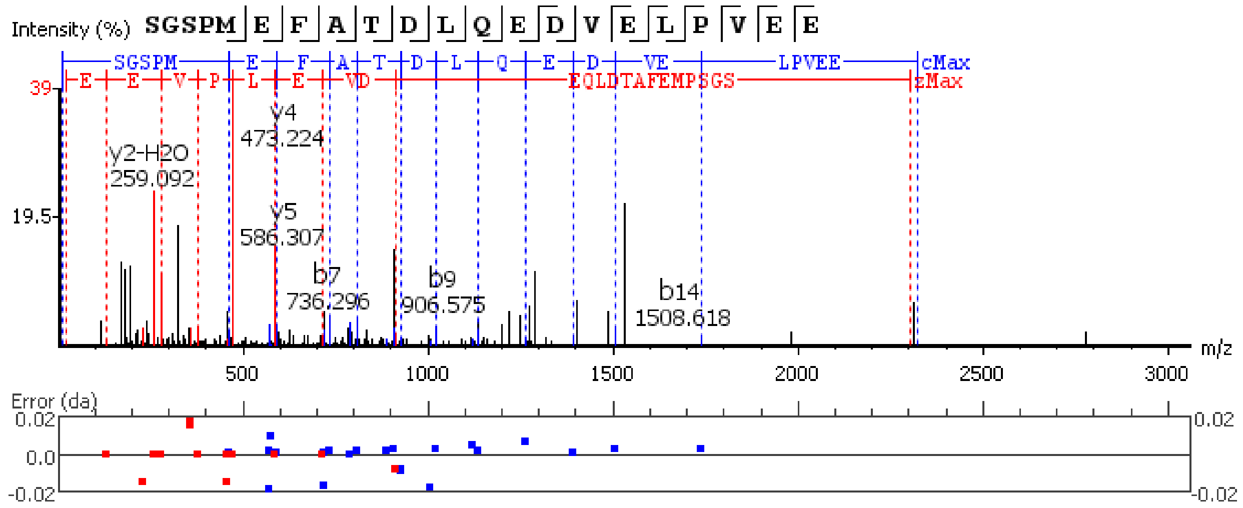


**Figure S5.54.** MS/MS spectrum of a FMRFamide neuropeptide with the sequence

SGSPM(+15.99)EFATDLQEDVELPVEE detected in the peptidergic signaling system of the

American lobster *Homarus american*us generated by PEAKS XPro software using data dependent acquisition

(DDA) method.

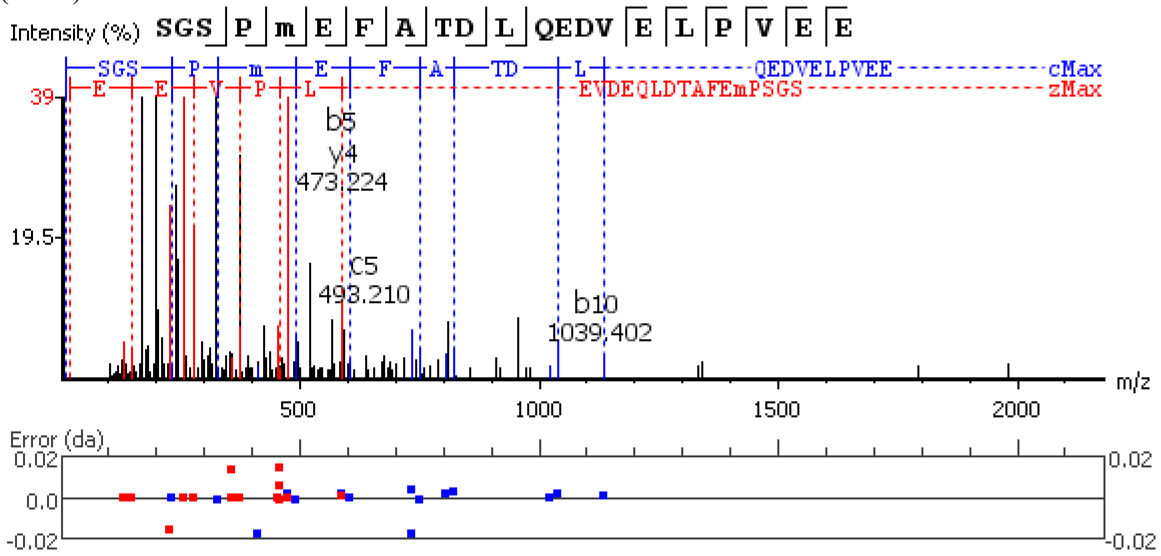


**Figure S5.55.** MS/MS spectrum of a FMRFamide neuropeptide with the sequence

GAHKNYLRF(-.98) detected in the peptidergic signaling system of the

American lobster *Homarus american*us generated by PEAKS XPro software using data dependent acquisition

(DDA) method.

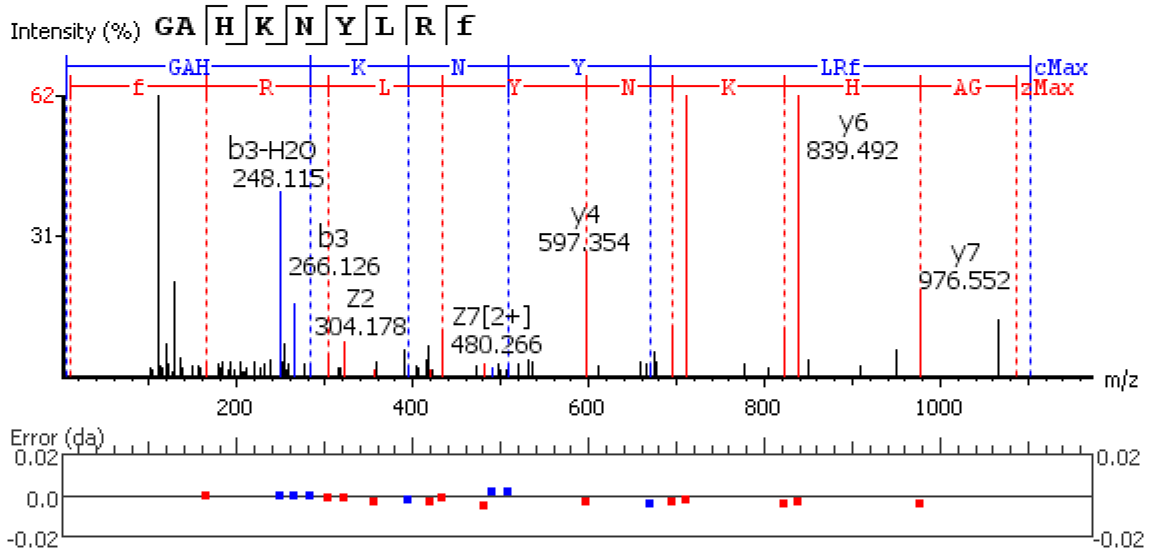


**Figure S5.56.** MS/MS spectrum of a FMRFamide neuropeptide with the sequence GNRNFLRF(-.98) detected in the peptidergic signaling system of the American lobster *Homarus american*us generated by PEAKS XPro software.

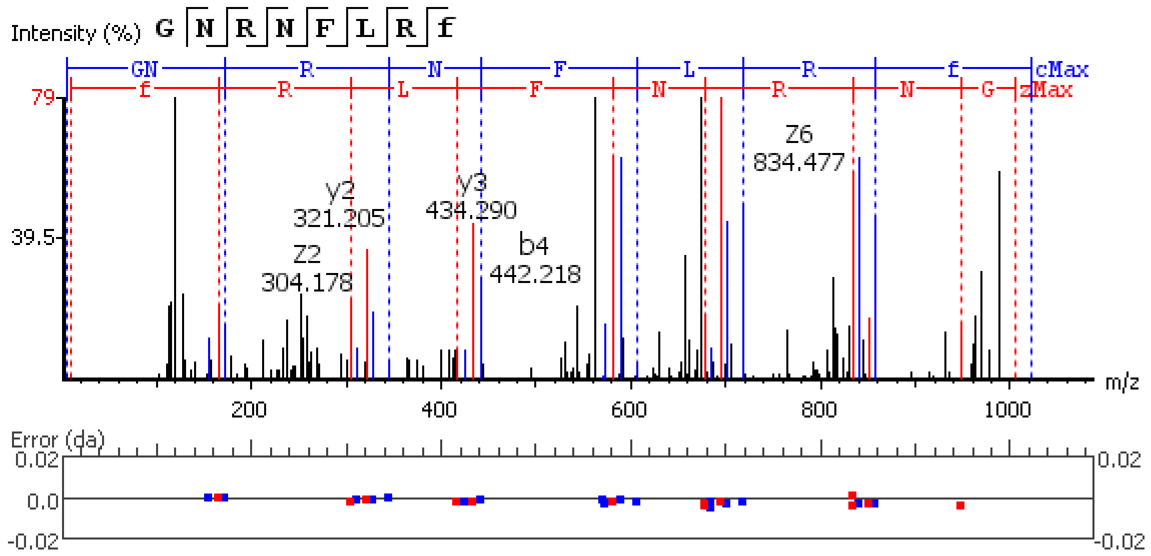


**Figure S5.57.** MS/MS spectrum of a FMRFamide neuropeptide with the sequence GDRNFLRF(-.98) detected in the peptidergic signaling system of the American lobster *Homarus american*us generated by PEAKS XPro software.

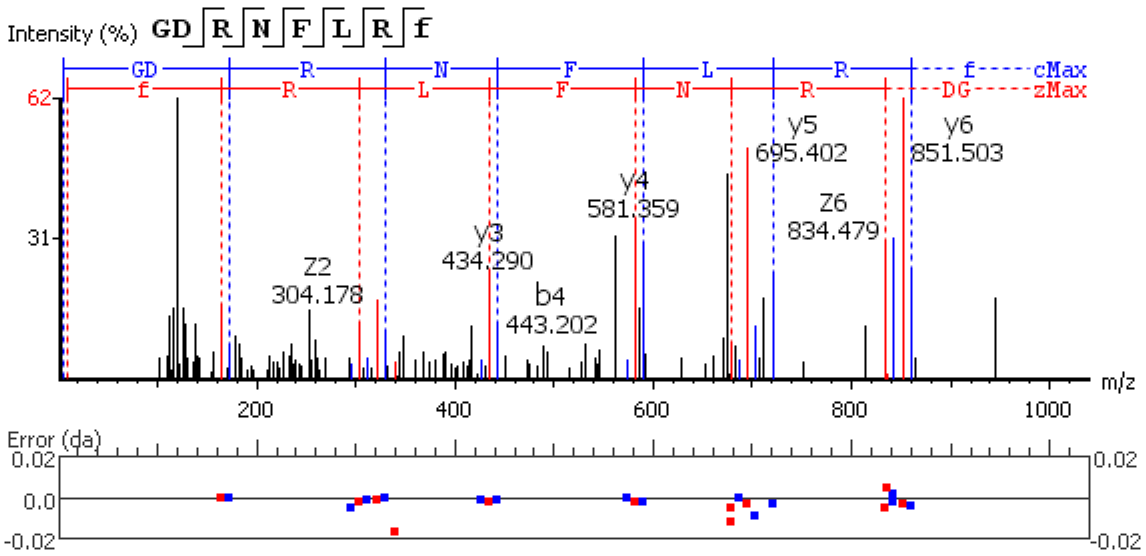


**Figure S5.58.** MS/MS spectrum of a FMRFamide neuropeptide with the sequence FSHDRNFLRF(-.98) detected in the peptidergic signaling system of the American lobster *Homarus american*us generated by PEAKS XPro software.

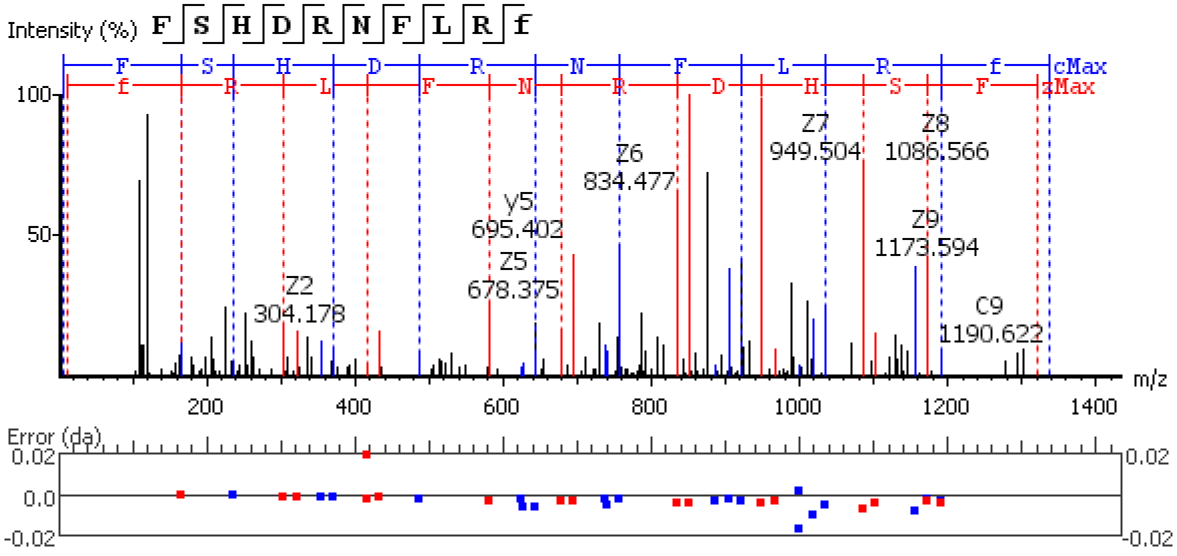


**Figure S5.59.** MS/MS spectrum of a FMRFamide neuropeptide with the sequence DGSDDYPSSSSSAESPAPVVVVRPVEYPRYV detected in the peptidergic signaling system of the American lobster *Homarus american*us generated by PEAKS XPro software.

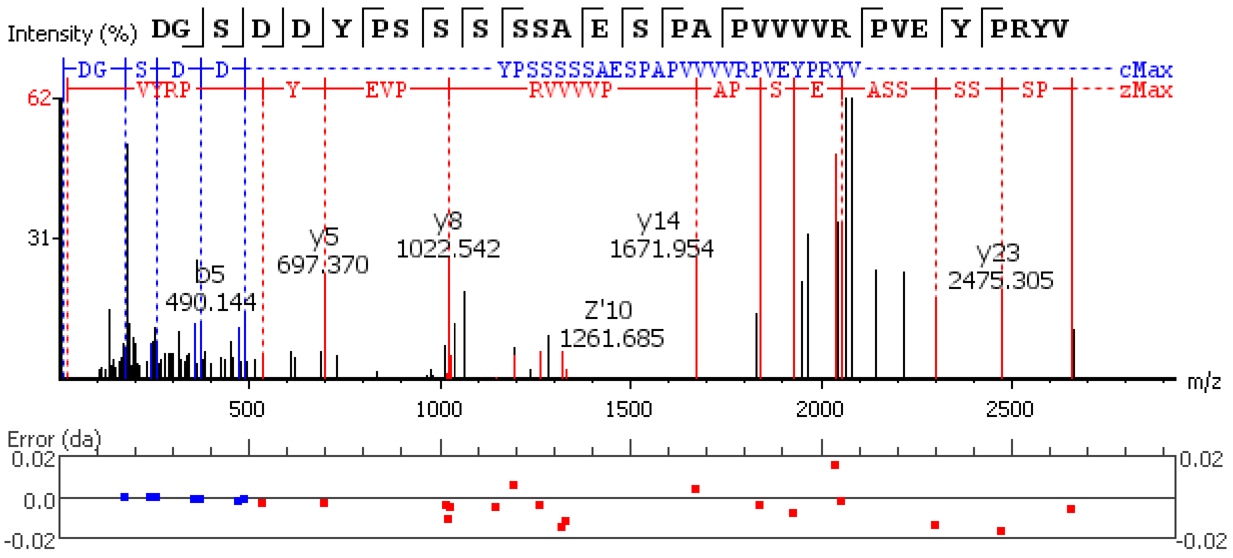


**Figure S5.60.** MS/MS spectrum of a FMRFamide neuropeptide with the sequence APSKNFLRF(-.98) detected in the peptidergic signaling system of the American lobster *Homarus americanus* generated by PEAKS XPro software.

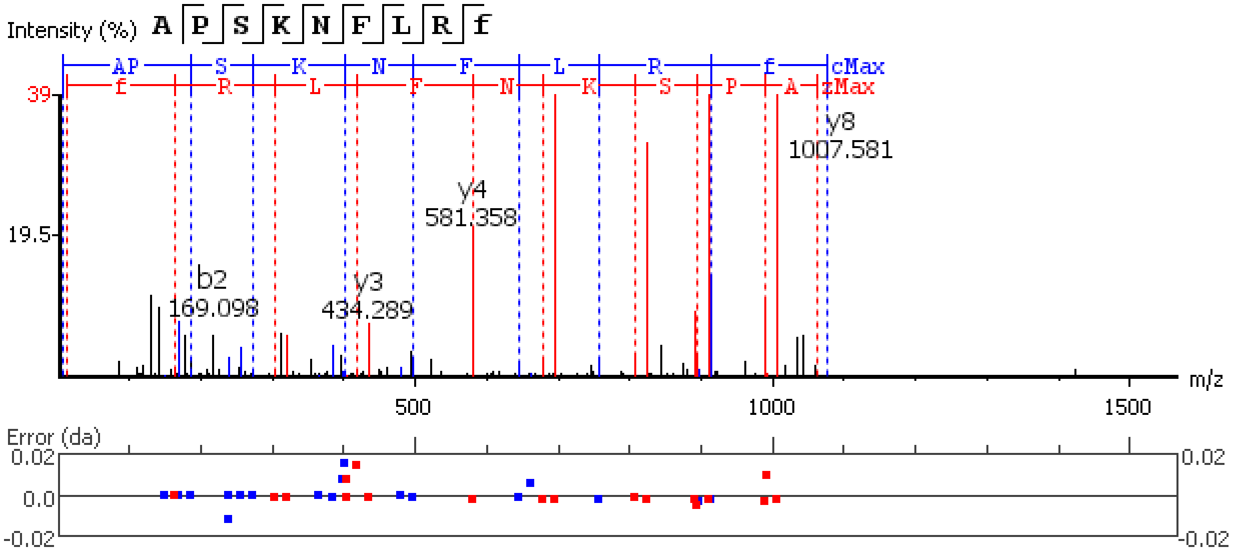


**Figure S5.61.** MS/MS spectrum of a GSEFLamide neuropeptide with the sequence LPTHLPDELDDPVV detected in the peptidergic signaling system of the American lobster *Homarus americanus* generated by PEAKS XPro software.

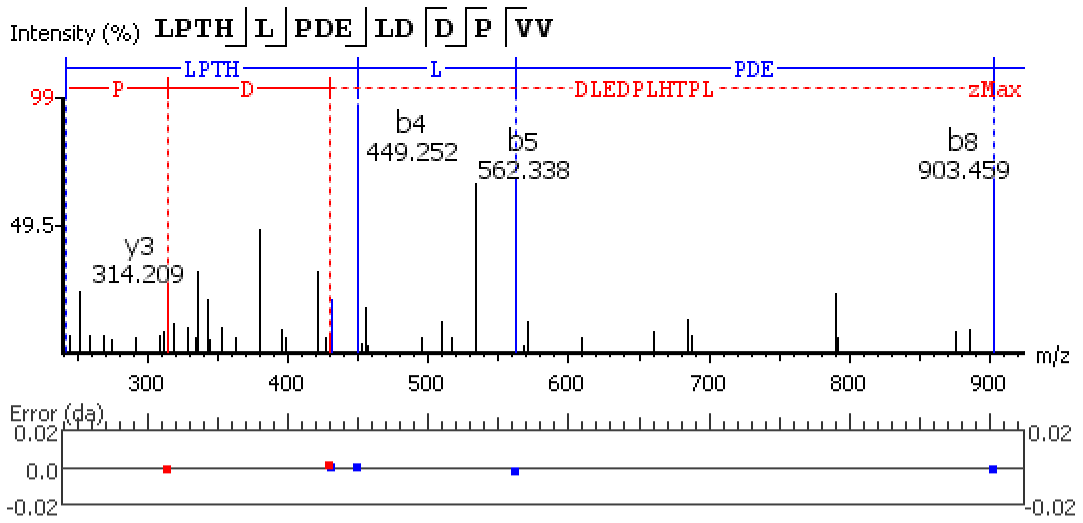


**Figure S5.62.** MS/MS spectrum of a GSEFLamide neuropeptide with the sequence RIGSEFL(-.98) detected in the peptidergic signaling system of the American lobster *Homarus american*us generated by PEAKS XPro software.

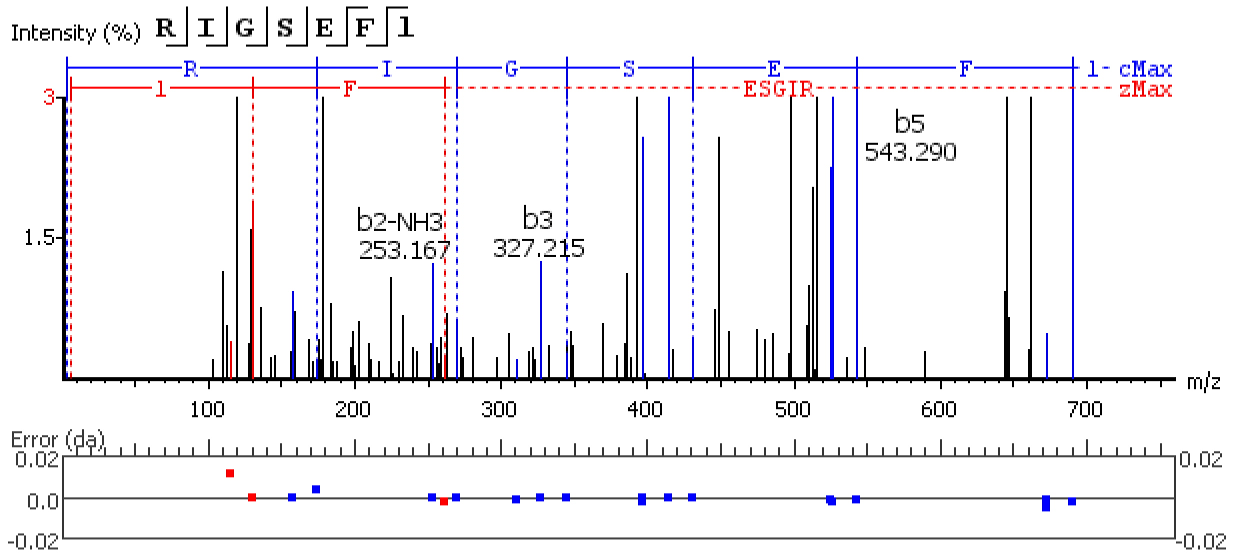


**Figure S5.63.** MS/MS spectrum of a GSEFLamide neuropeptide with the sequence QYEPEFAHTLDYDT detected in the peptidergic signaling system of the American lobster *Homarus american*us generated by PEAKS XPro software.

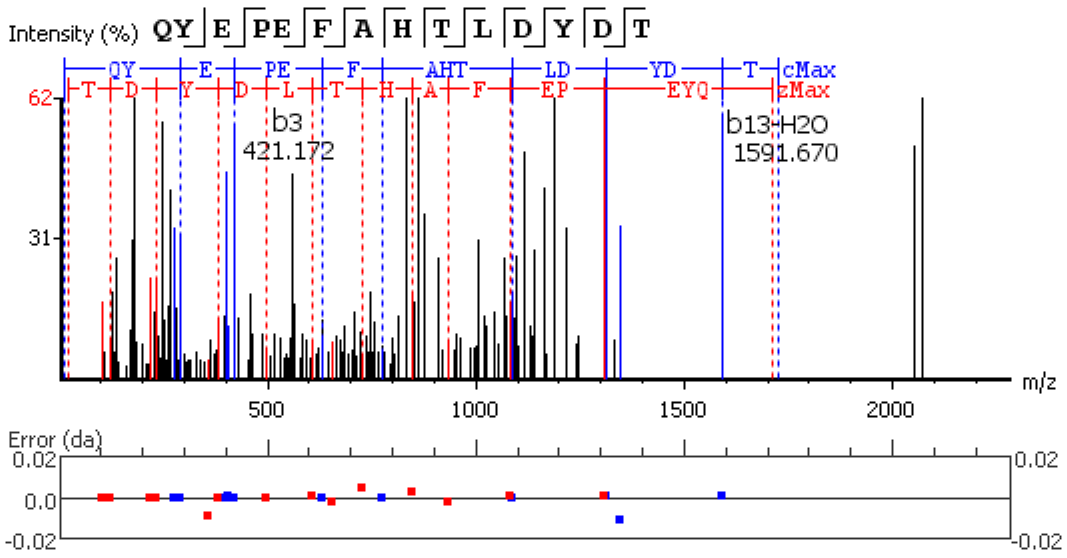


**Figure S5.64.** MS/MS spectrum of a GSEFLamide neuropeptide with the sequence

Q(17.03)YEPEFAHTLDYDT detected in the peptidergic signaling system of the American

lobster *Homarus american*us generated by PEAKS XPro software using data dependent acquisition (DDA)

method.

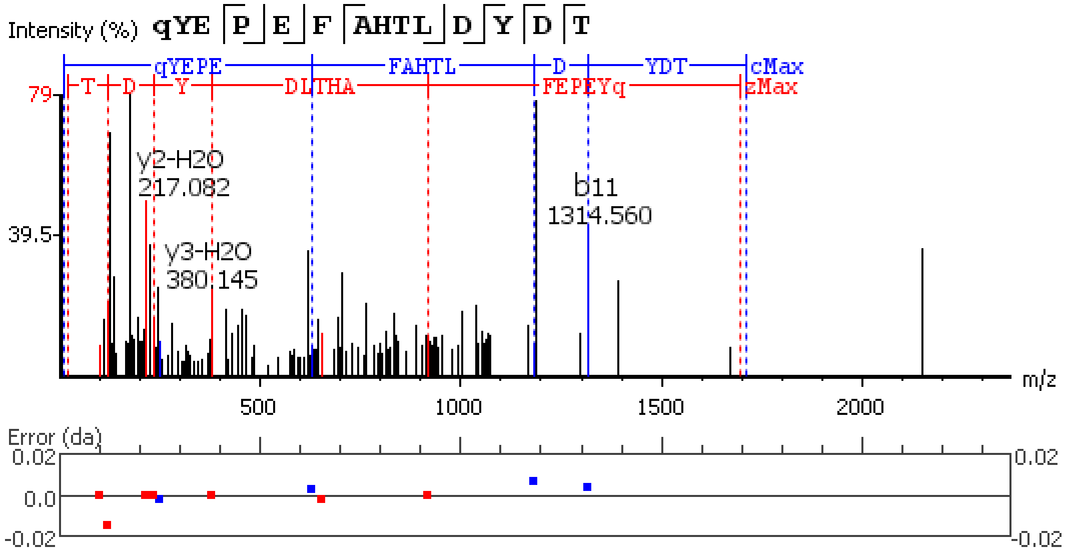


**Figure S5.65.** MS/MS spectrum of a Myosuppressin neuropeptide with the sequence QDLDHVFLRF(-.98) detected in the peptidergic signaling system of the American lobster *Homarus american*us generated by PEAKS XPro software.

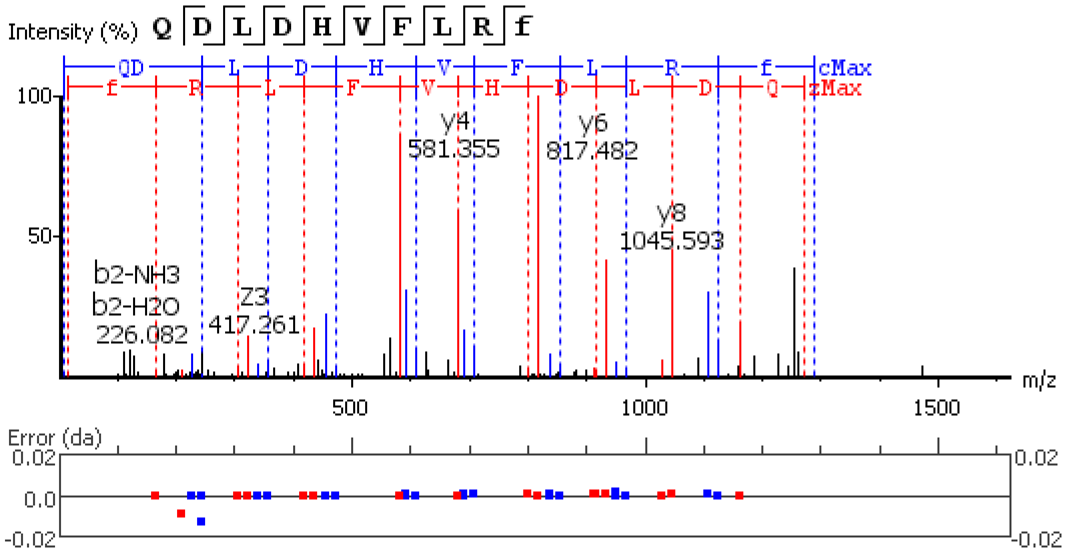


**Figure S5.66.** MS/MS spectrum of a Myosuppressin neuropeptide with the sequence Q(-17.03)DLDHVFLRF(-.98) detected in the peptidergic signaling system of the American lobster *Homarus american*us generated by PEAKS XPro software.

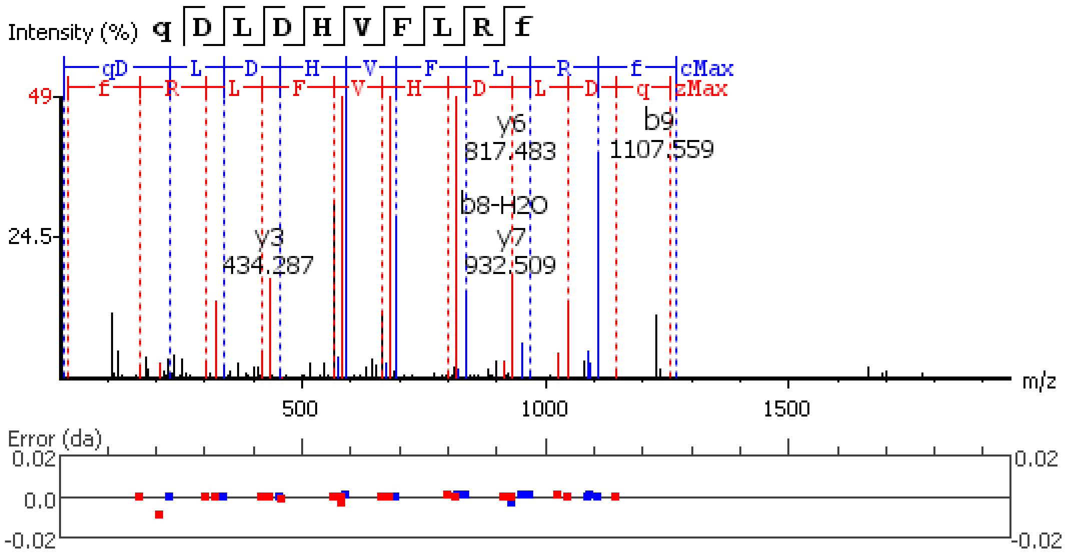


**Figure S5.67.** MS/MS spectrum of a Orcokinin neuropeptide with the sequence GPIKAAPARSSPQQDAAAGYTDGAPV detected in the peptidergic signaling system of the American lobster *Homarus american*us generated by PEAKS XPro software.

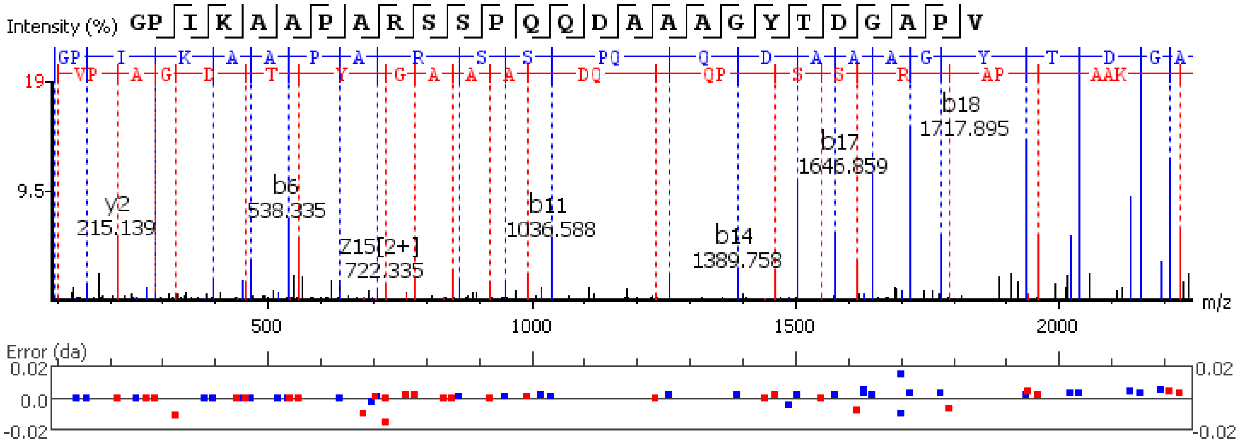


**Figure S5.68.** MS/MS spectrum of a Orcokinin neuropeptide with the sequence FDAFTTGFGHN detected in the peptidergic signaling system of the American lobster *Homarus americanus* generated by PEAKS XPro software.

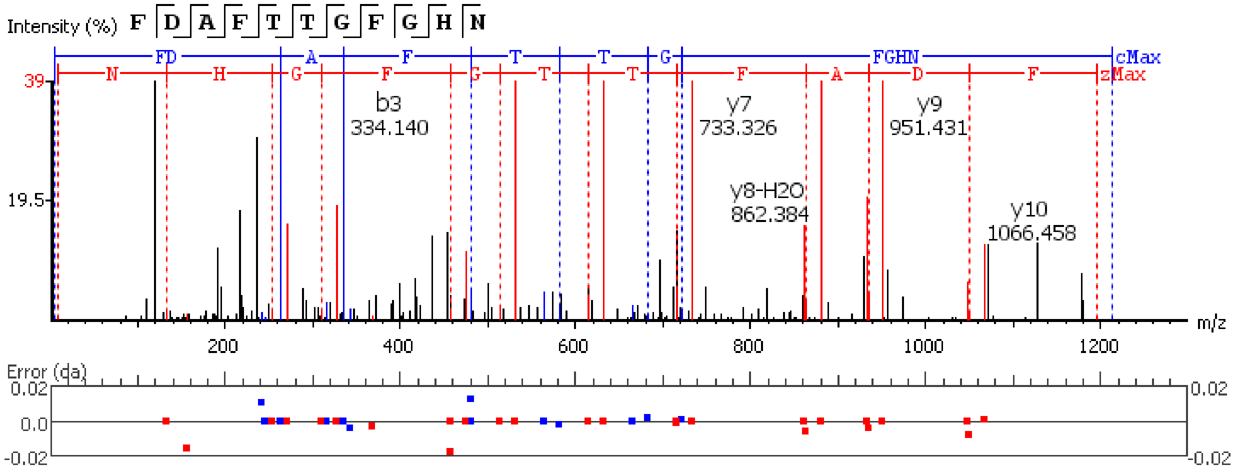


**Figure S5.69.** MS/MS spectrum of a Orcokinin neuropeptide with the sequence SSEDMDRLGFGFN detected in the peptidergic signaling system of the American lobster *Homarus americanus* generated by PEAKS XPro software.

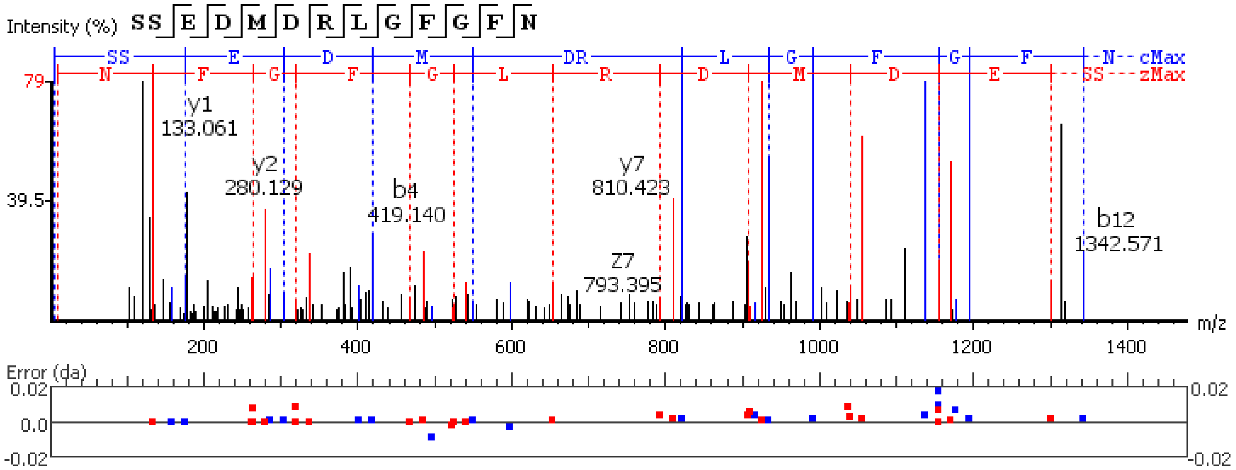

**Figure S5.70.** MS/MS spectrum of a Orcokinin neuropeptide with the sequence SSEDM(+15.99)DRLGFGFN detected in the peptidergic signaling system of the American lobster *Homarus americanus* generated by PEAKS XPro software.

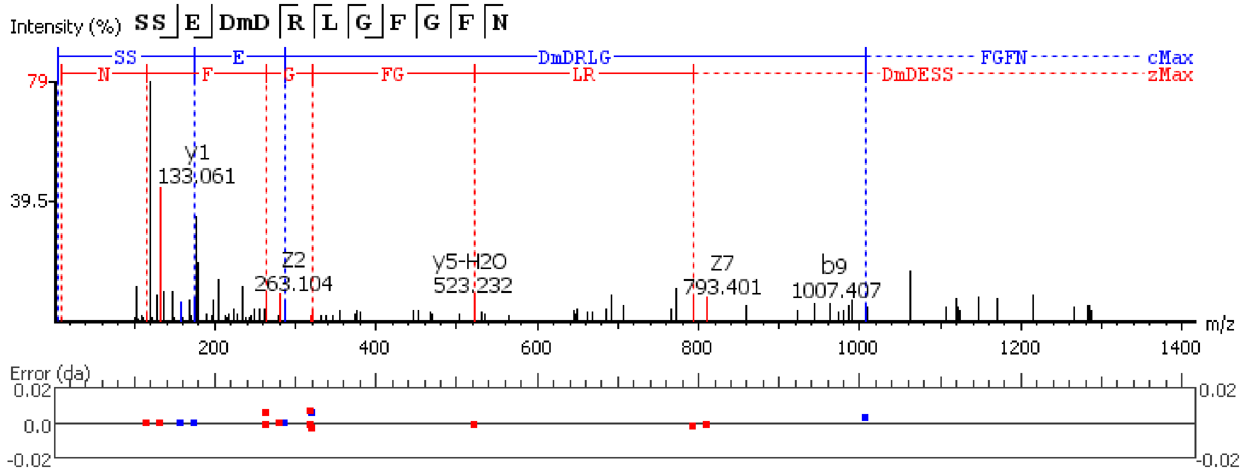


**Figure S5.71.** MS/MS spectrum of a Orcokinin neuropeptide with the sequence NFDEIDRSGFGFH detected in the peptidergic signaling system of the American lobster *Homarus americanus* generated by PEAKS XPro software.

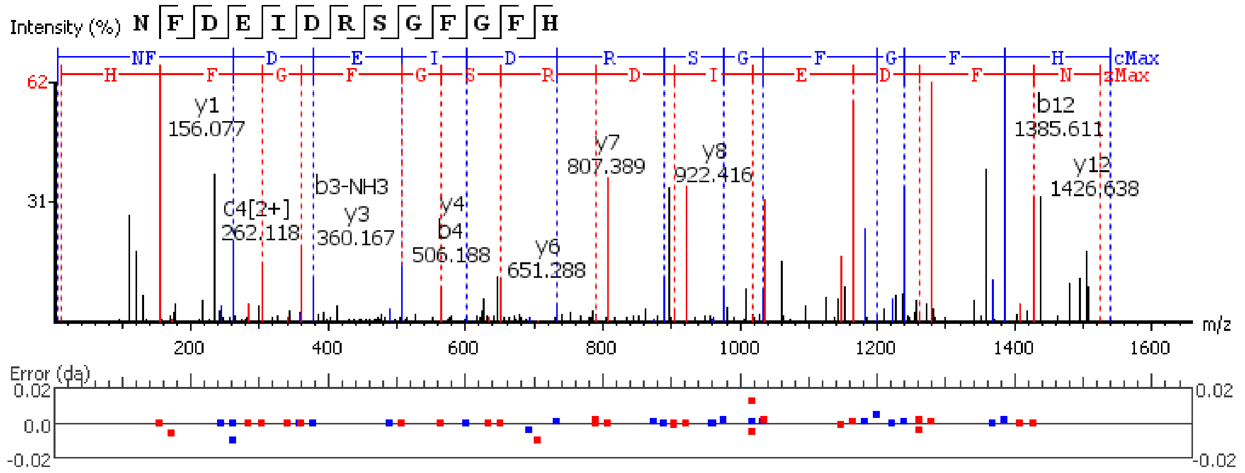


**Figure S5.72.** MS/MS spectrum of a Orcokinin neuropeptide with the sequence NFDEIDRSGFGFN detected in the peptidergic signaling system of the American lobster *Homarus americanus* generated by PEAKS XPro software.

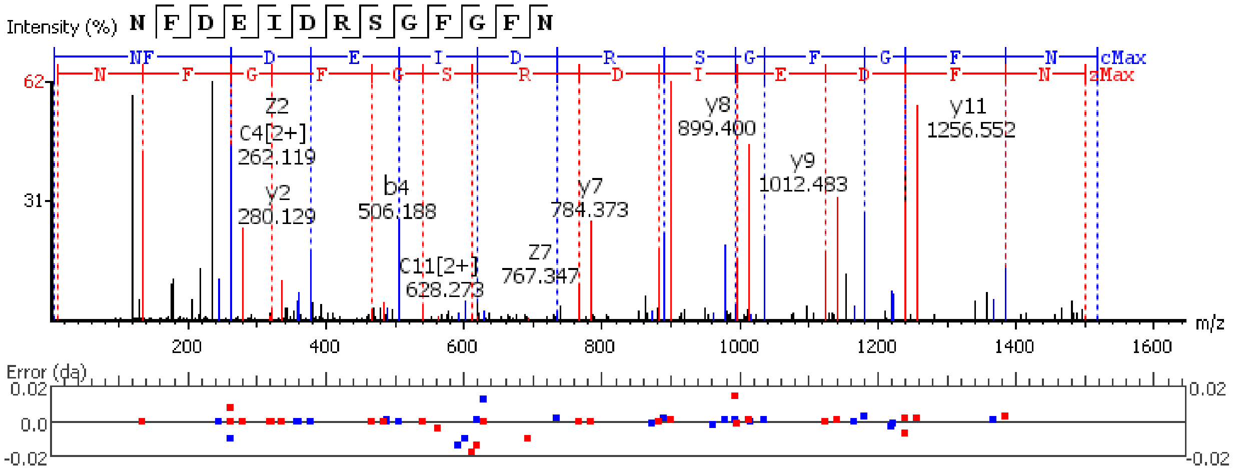


**Figure S5.73.** MS/MS spectrum of a Orcokinin neuropeptide with the sequence GDYDVYPE detected in the peptidergic signaling system of the American lobster *Homarus american*us generated by PEAKS XPro software.

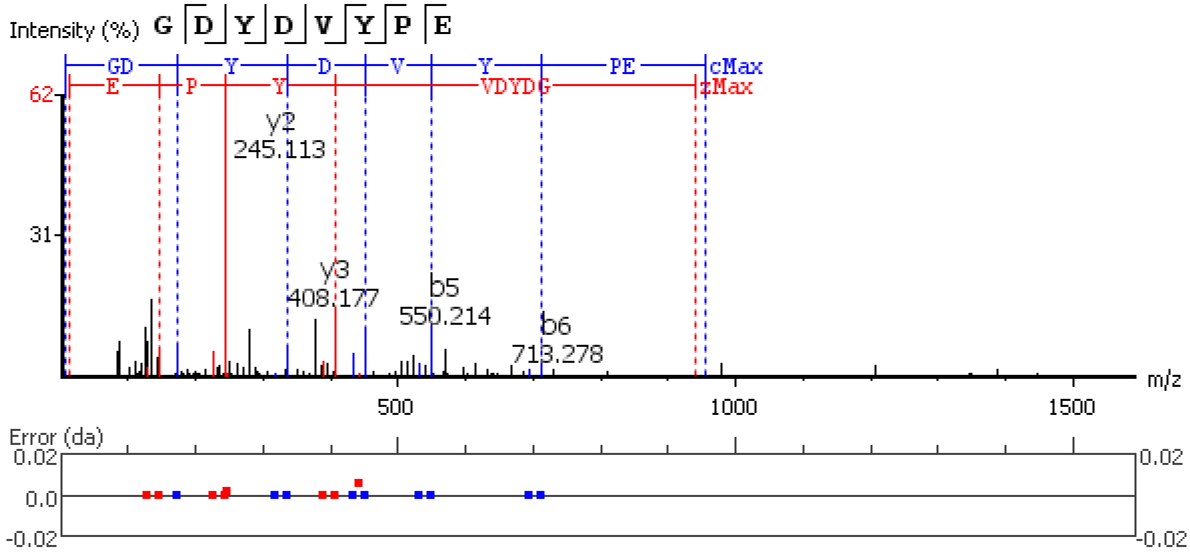


**Figure S5.74.** MS/MS spectrum of a Orcokinin neuropeptide with the sequence NFDEIDRSGFGFV detected in the peptidergic signaling system of the American lobster *Homarus american*us generated by PEAKS XPro software.

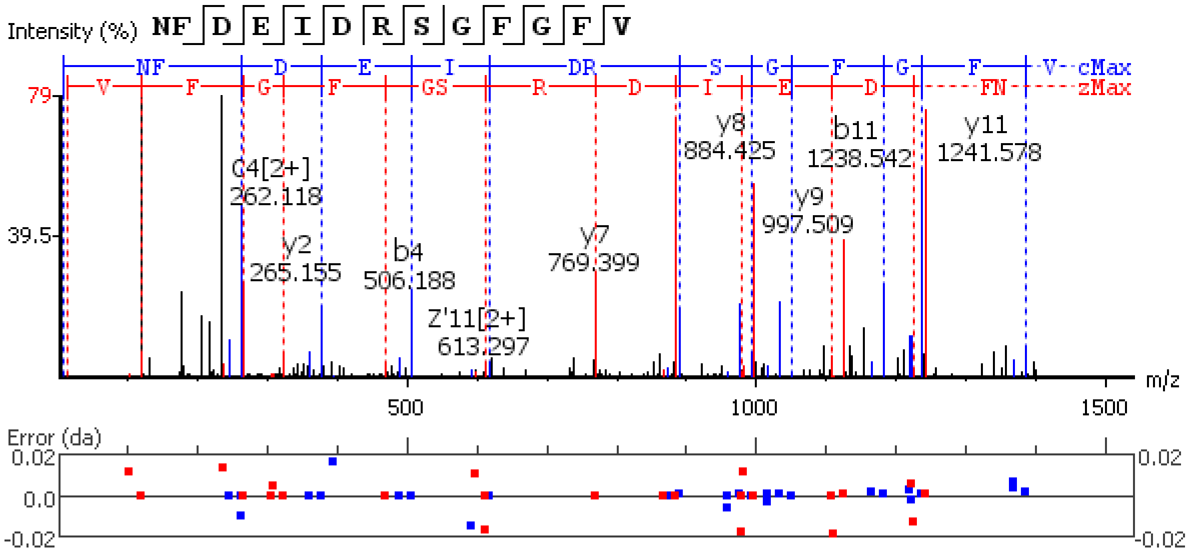


**Figure S5.75.** MS/MS spectrum of a Orcokinin neuropeptide with the sequence VYGPRDIANLY detected in the peptidergic signaling system of the American lobster *Homarus americanus* generated by PEAKS XPro software.

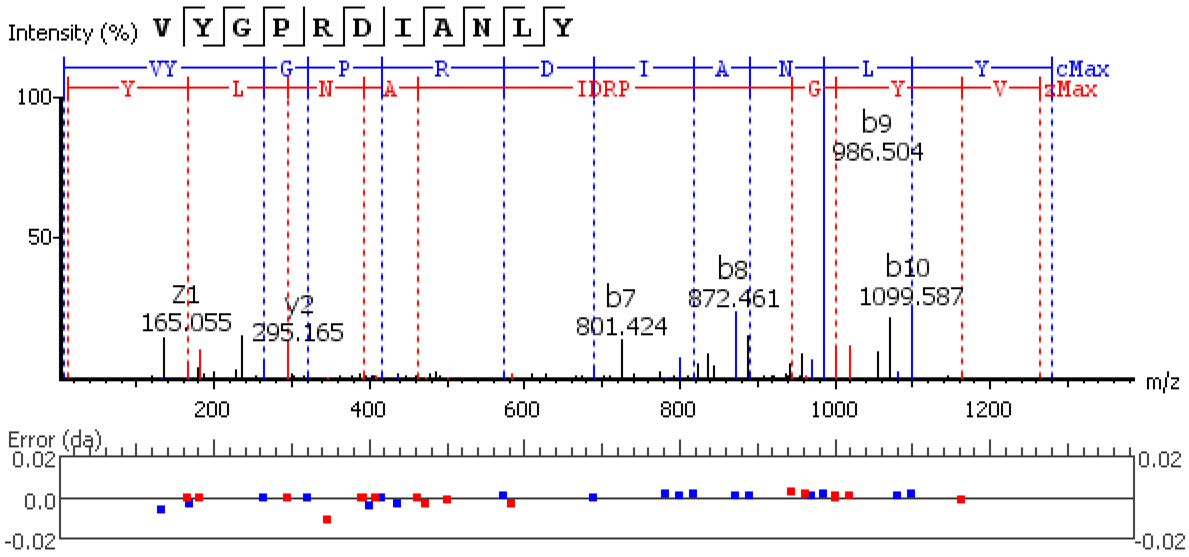


**Figure S5.76.** MS/MS spectrum of a pigment dispersing hormone (PDH) neuropeptide with the sequence Q(-17.03)ELKYPEREVVADMAAQILRVALGPWGSVAAVPR detected in the peptidergic signaling system of the American lobster *Homarus americanus* generated by PEAKS XPro software.

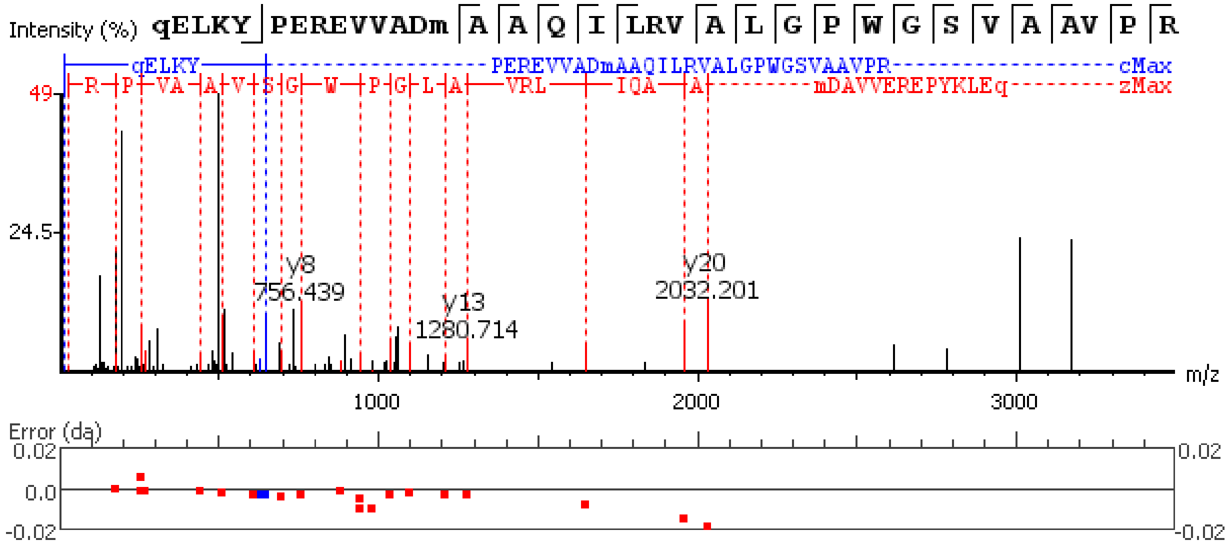


**Figure S5.77.** MS/MS spectrum of a pigment dispersing hormone (PDH) neuropeptide with the sequence Q(17.03)ELKYPEREVVADM(+15.99)AAQILRVALGPWGSVAAVPR detected in the peptidergic signaling system of the American lobster *Homarus american*us generated by PEAKS XPro software.

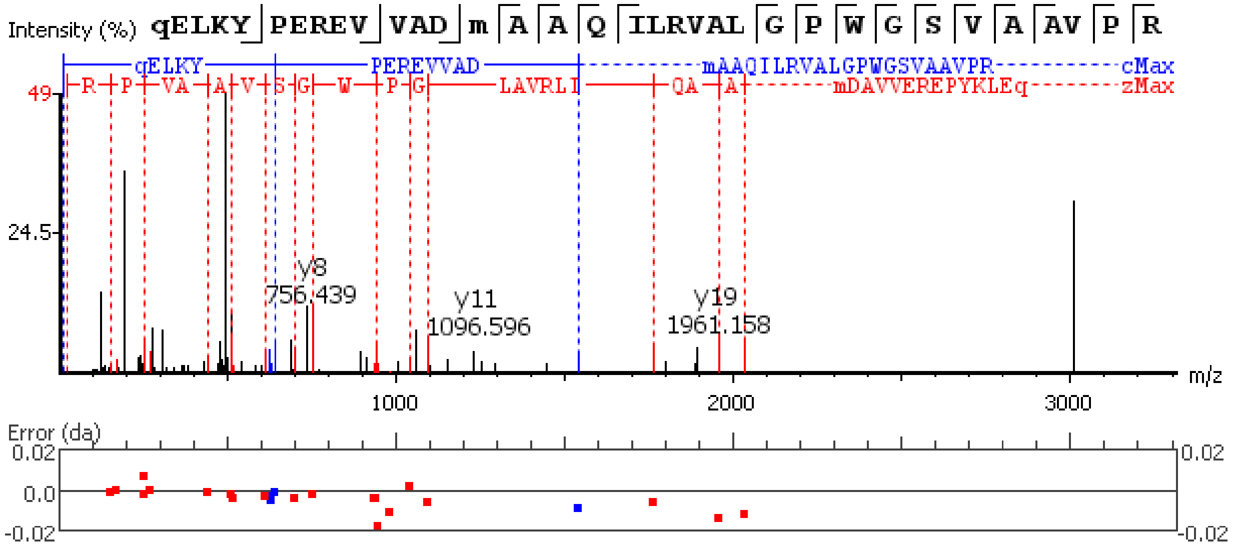


**Figure S5.78.** MS/MS spectrum of a pigment dispersing hormone (PDH) neuropeptide with the sequence Q(-17.03)ELKYPEREVVAELAAQIL detected in the peptidergic signaling system of the American lobster *Homarus american*us generated by PEAKS XPro software.

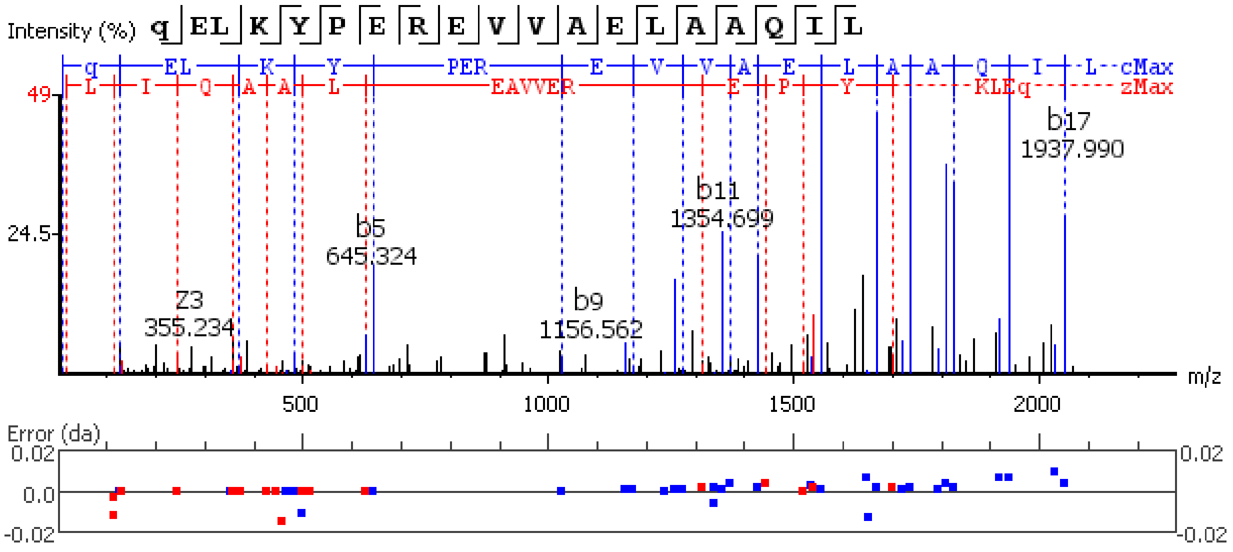


**Figure S5.79.** MS/MS spectrum of a pigment dispersing hormone (PDH) neuropeptide with the sequence Q(-17.03)ELKYPEREVVADMAAQIL detected in the peptidergic signaling system of the American lobster *Homarus american*us generated by PEAKS XPro software.

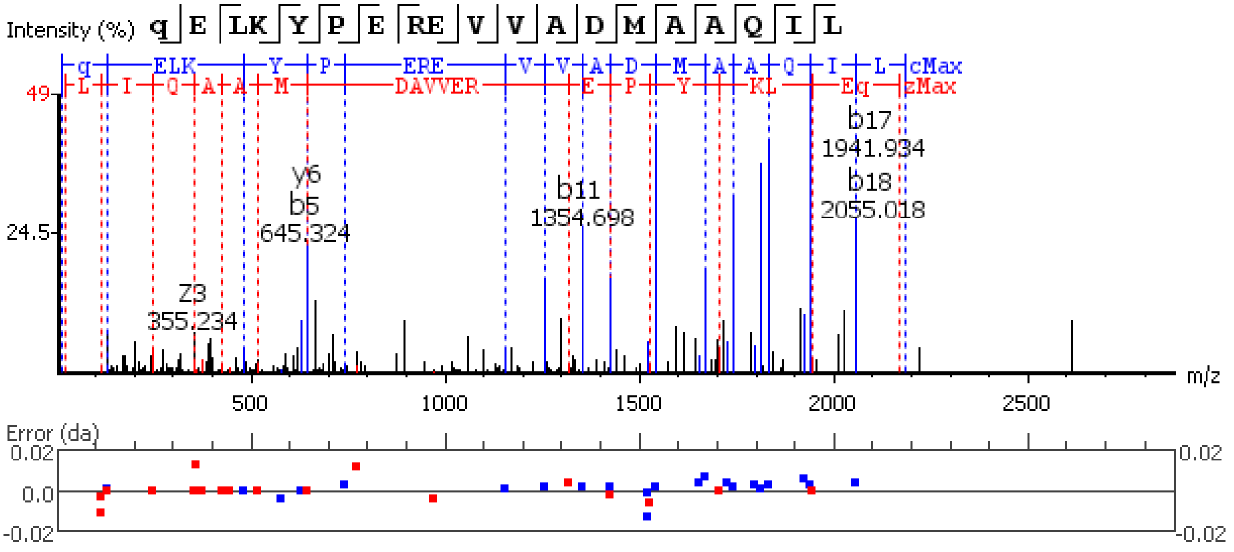


**Figure S5.80.** MS/MS spectrum of a pigment dispersing hormone (PDH) neuropeptide with the sequence Q(-17.03)ELKYPEREVVADM(+15.99)AAQIL detected in the peptidergic signaling system of the American lobster *Homarus american*us generated by PEAKS XPro software using data dependent acquisition (DDA) method.

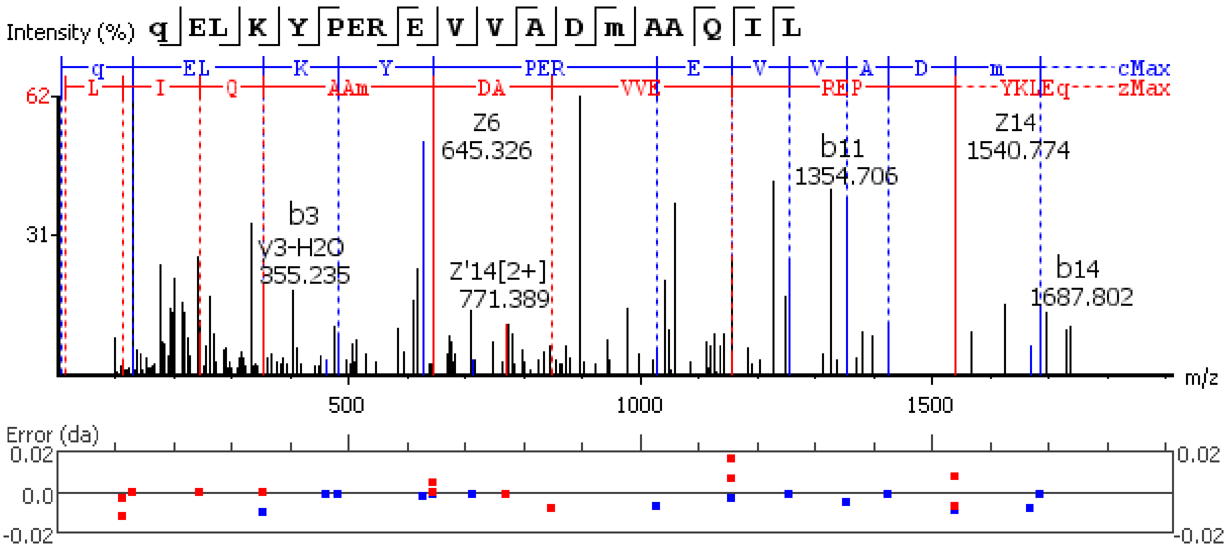


**Figure S5.81.** MS/MS spectrum of a pigment dispersing hormone (PDH) neuropeptide with the sequence NSELINSLLGIPKVMNDA(-.98) detected in the peptidergic signaling system of the American lobster *Homarus american*us generated by PEAKS XPro software.

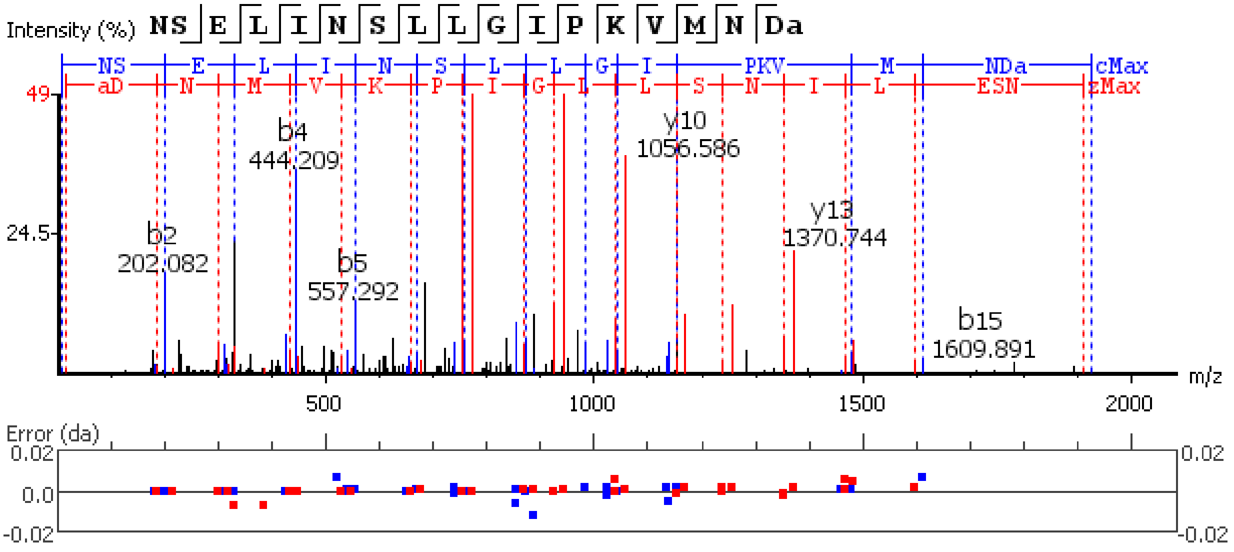


**Figure S5.82.** MS/MS spectrum of a pigment dispersing hormone (PDH) neuropeptide with the sequence NSELINSLLGIPKVM(+15.99)NDA(-.98) detected in the peptidergic signaling system of the American lobster *Homarus american*us generated by PEAKS XPro software.

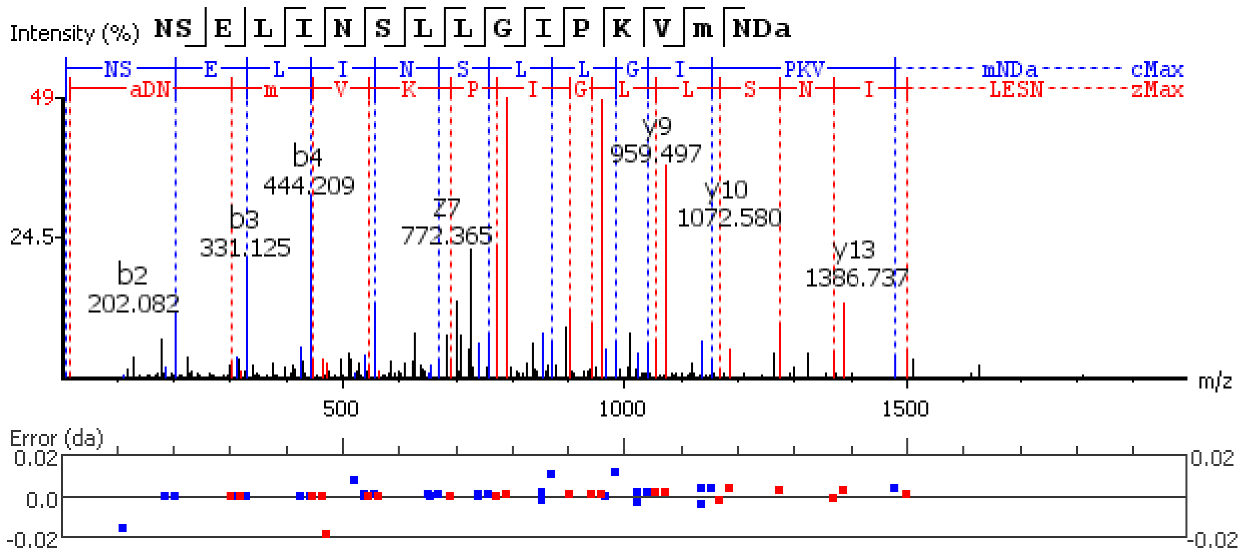


**Figure S5.83.** MS/MS spectrum of a pigment dispersing hormone (PDH) neuropeptide with the sequence NSELINSILGLPKVMNDA(-.98) detected in the peptidergic signaling system of the American lobster *Homarus american*us generated by PEAKS XPro software.

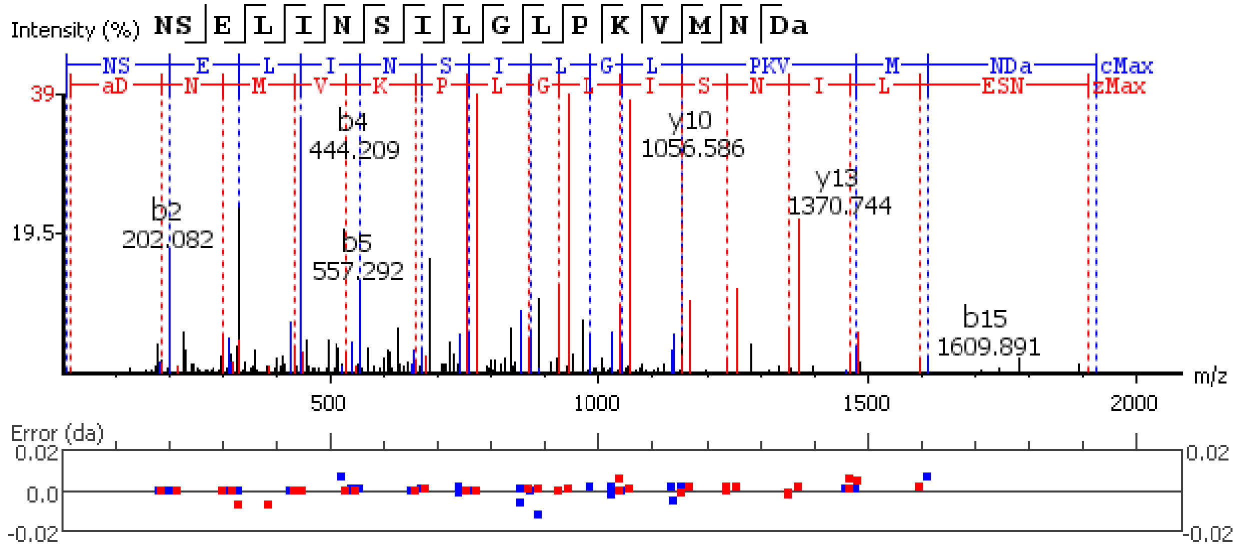


**Figure S5.84.** MS/MS spectrum of a pigment dispersing hormone (PDH) neuropeptide with the sequence NSELINSILGLPKVM(+15.99)NDA(-.98) detected in the peptidergic signaling system of the American lobster *Homarus american*us generated by PEAKS XPro software.

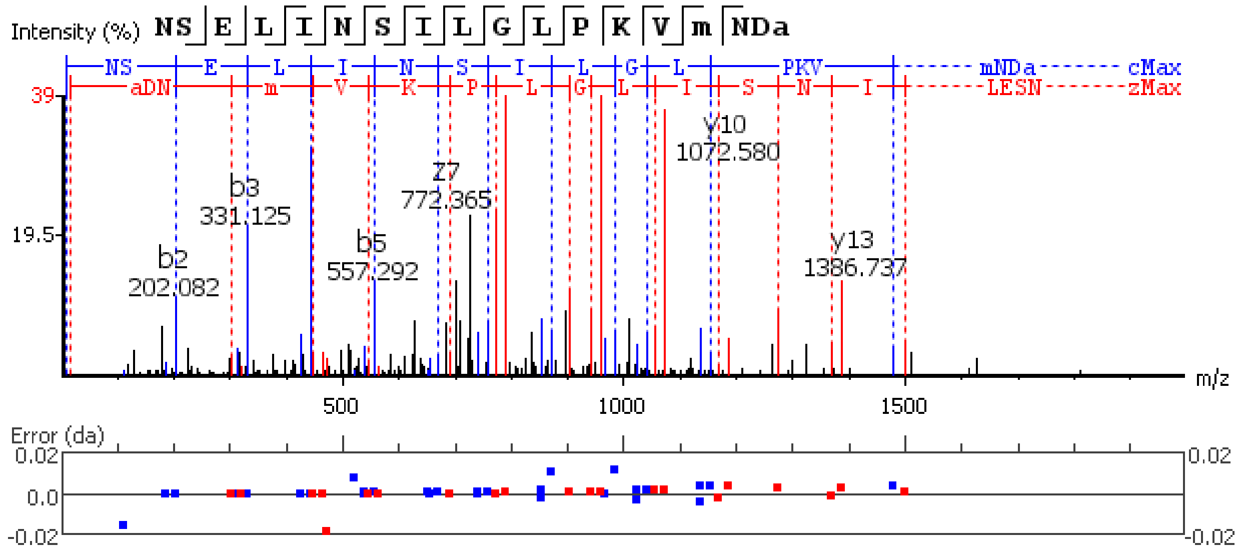


**Figure S5.85.** MS/MS spectrum of a pigment dispersing hormone (PDH) neuropeptide with the sequence NSEILNTLLGSQDLSNMRSA(-.98) detected in the peptidergic signaling system of the American lobster *Homarus american*us generated by PEAKS XPro software.

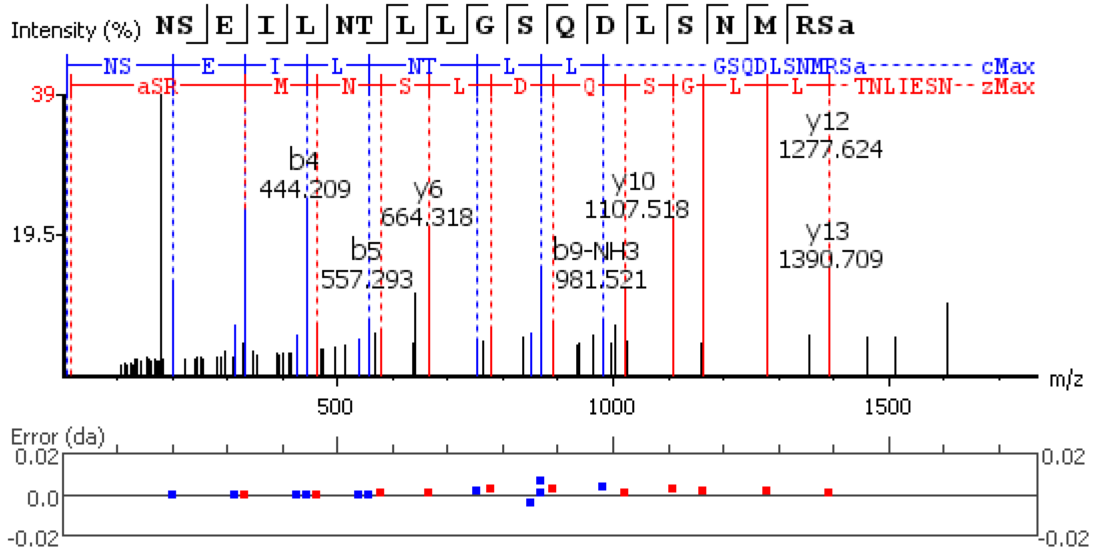


**Figure S5.86.** MS/MS spectrum of a Pyrokinin neuropeptide with the sequence GDGFAFSPRL(-.98) detected in the peptidergic signaling system of the American lobster *Homarus american*us generated by PEAKS XPro software.

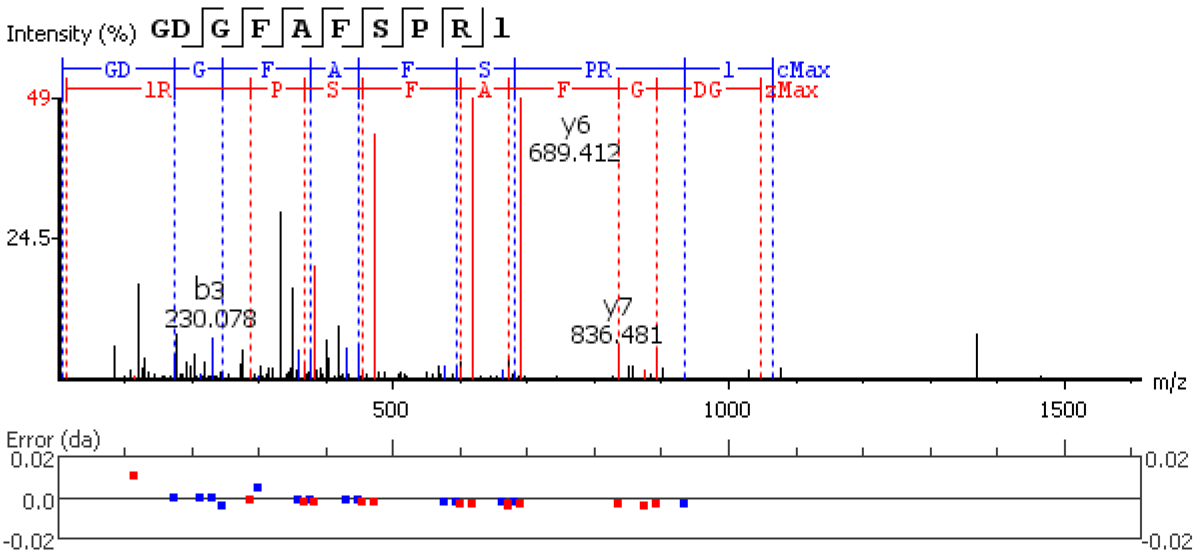


**Figure S5.87.** MS/MS spectrum of a Pyrokinin neuropeptide with the sequence GADFAFSPRL(-.98) detected in the peptidergic signaling system of the American lobster *Homarus american*us generated by PEAKS XPro software.

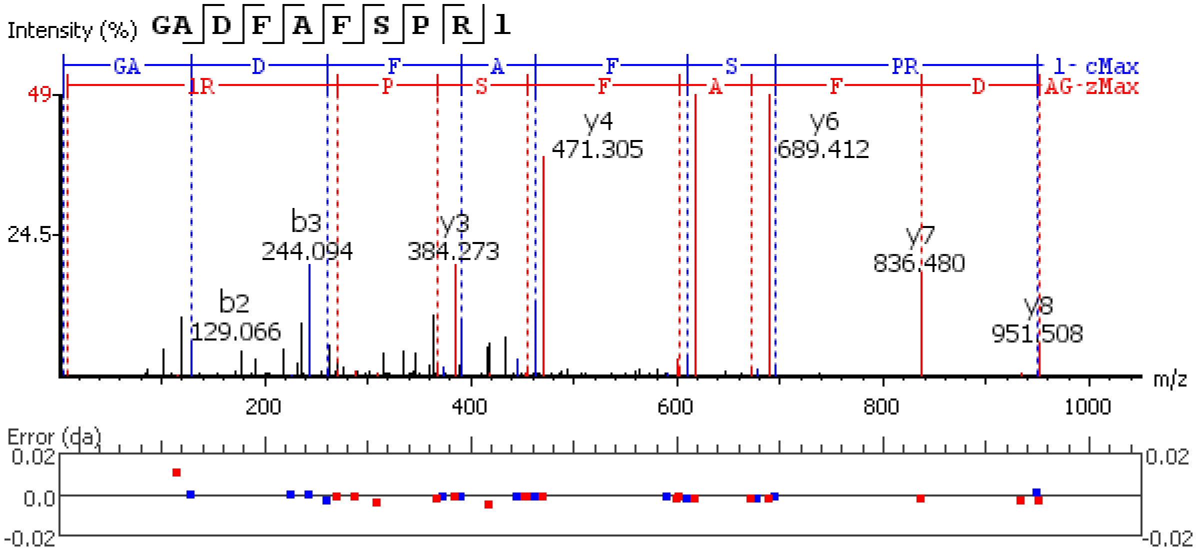


**Figure S5.88.** MS/MS spectrum of a Pyrokinin neuropeptide with the sequence SEFVFSSRP(-.98) detected in the peptidergic signaling system of the American lobster *Homarus american*us generated by PEAKS XPro software.

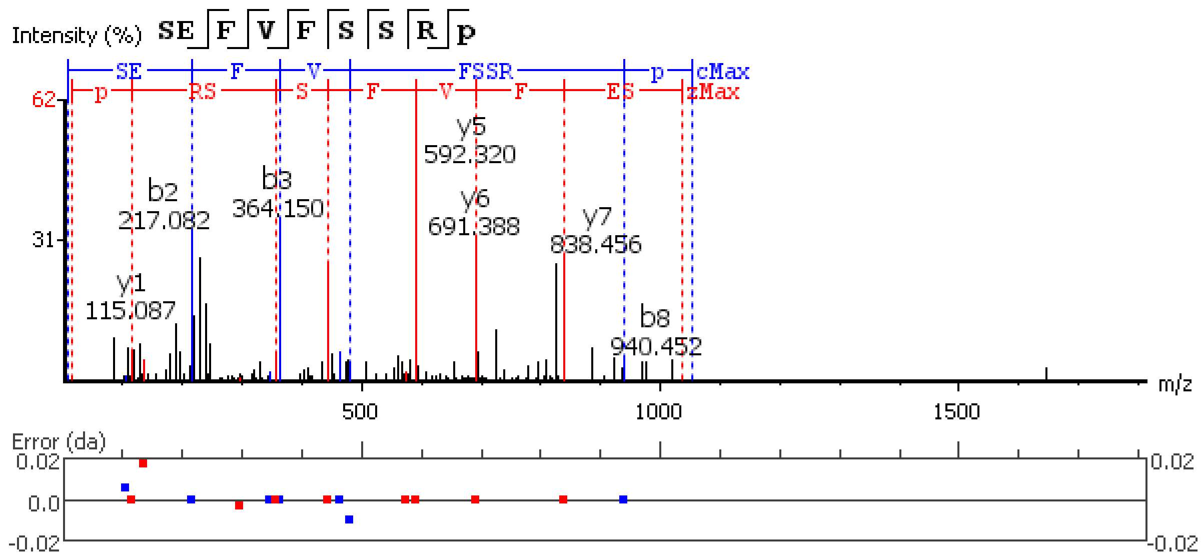


**Figure S5.89.** MS/MS spectrum of a Pyrokinin neuropeptide with the sequence VRRSLFSPRL(-.98) detected in the peptidergic signaling system of the American lobster *Homarus americanus* generated by PEAKS XPro software.

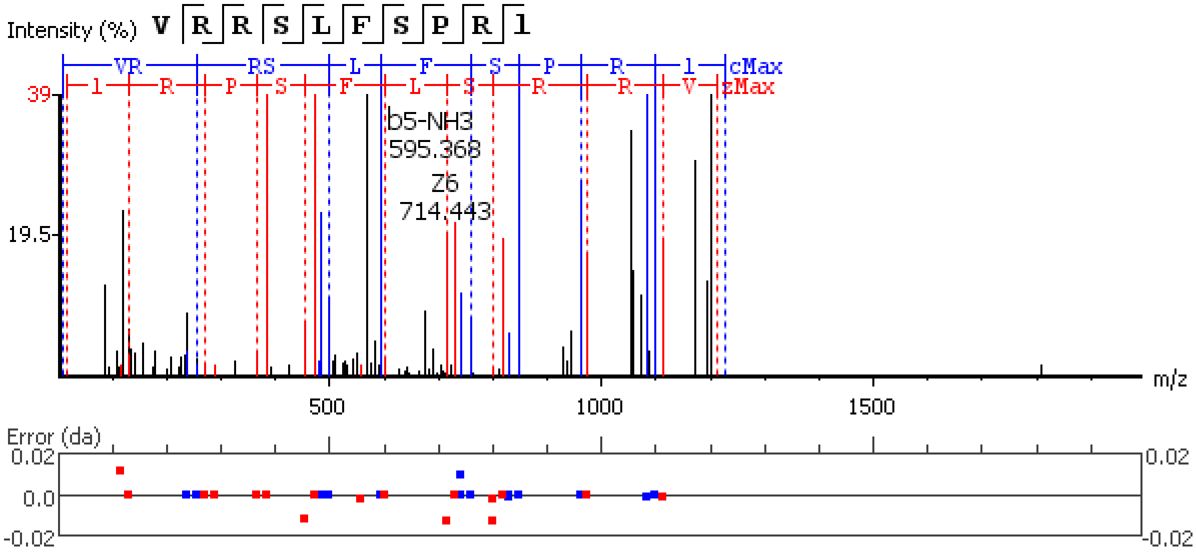


**Figure S5.90.** MS/MS spectrum of a Pyrokinin neuropeptide with the sequence AYFSPRL(-.98) detected in the peptidergic signaling system of the American lobster *Homarus americanus* generated by PEAKS XPro software.

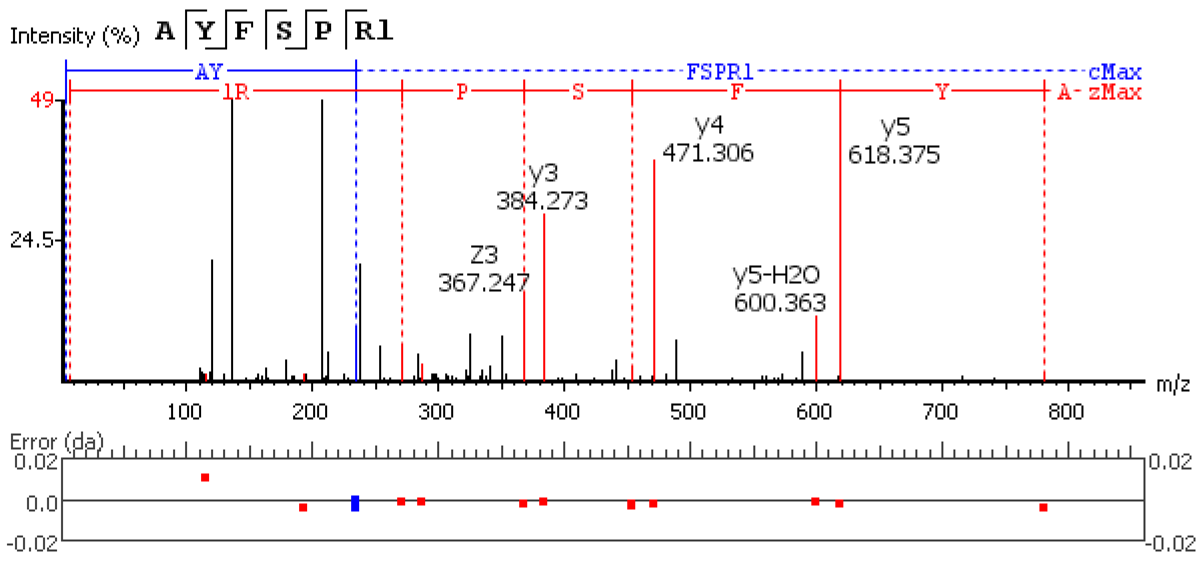


**Figure S5.91.** MS/MS spectrum of a Pyrokinin neuropeptide with the sequence LYYSQRP(-.98) detected in the peptidergic signaling system of the American lobster *Homarus americanus* generated by PEAKS XPro software.

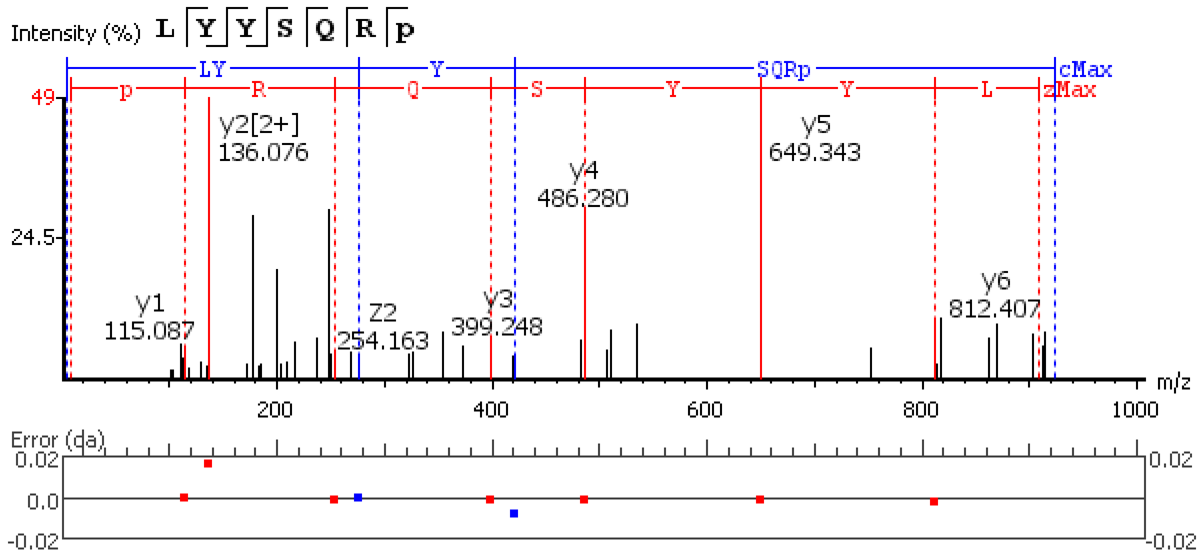


**Figure S5.92.** MS/MS spectrum of a Pyrokinin neuropeptide with the sequence SLFSPRL(-.98) detected in the peptidergic signaling system of the American lobster *Homarus americanus* generated by PEAKS XPro software.

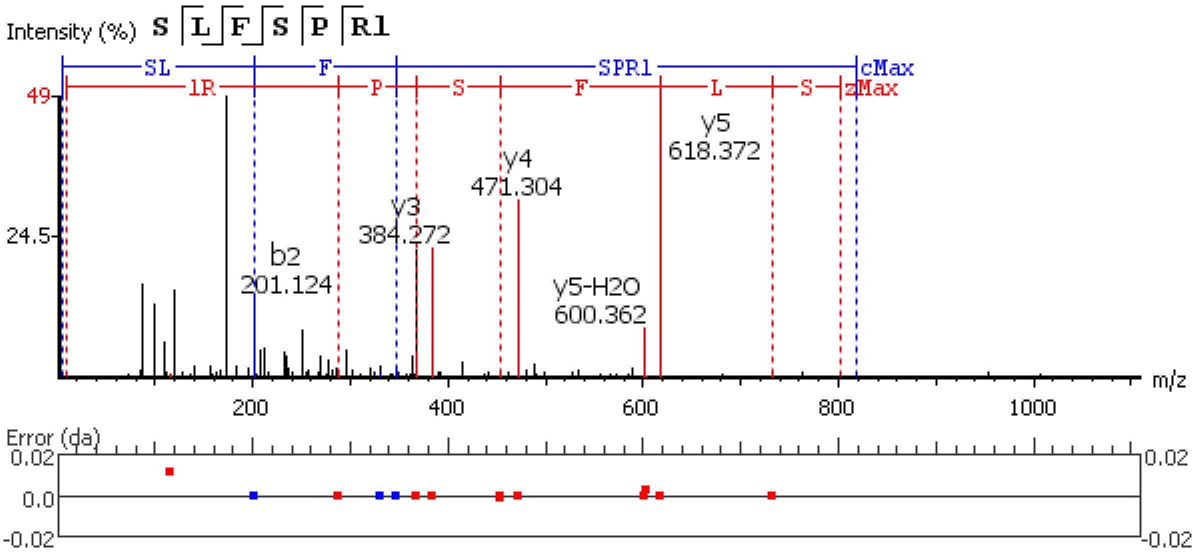


**Figure S5.93.** MS/MS spectrum of a Pyrokinin neuropeptide with the sequence SDFAFSPRL(-.98) detected in the peptidergic signaling system of the American lobster *Homarus american*us generated by PEAKS XPro software.

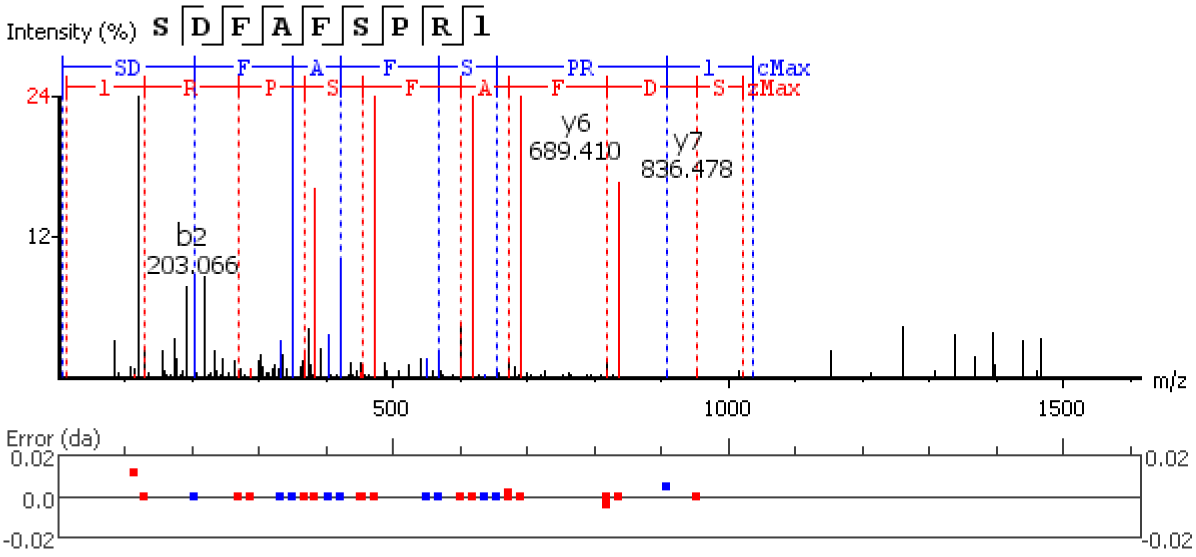


**Figure S5.94.** MS/MS spectrum of a SIFamide neuropeptide with the sequence VYRKPPFNGSIF(-.98) detected in the peptidergic signaling system of the American lobster *Homarus american*us generated by PEAKS XPro software.

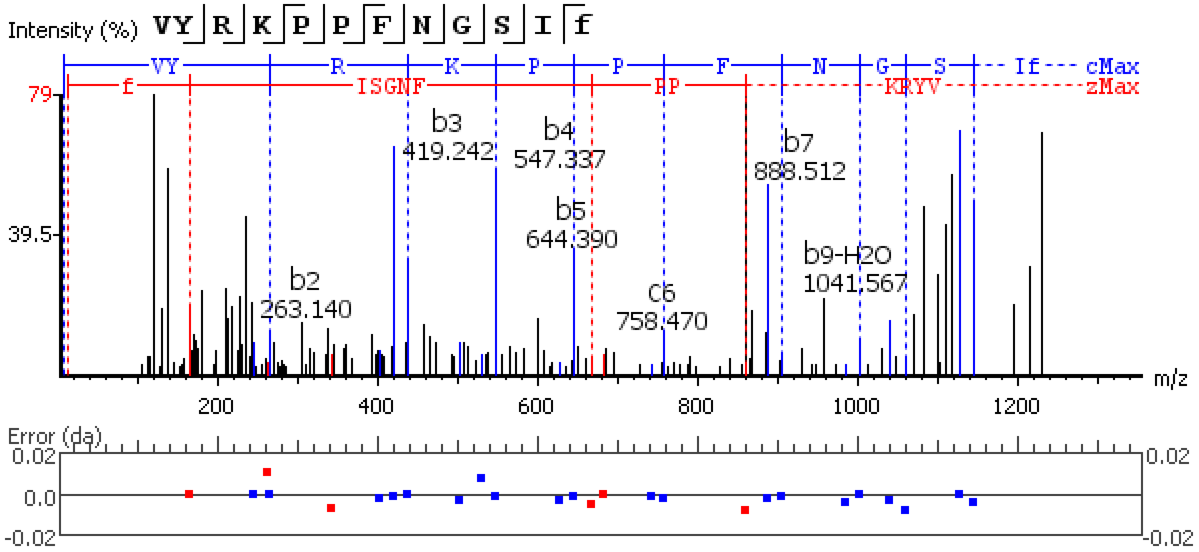


**Figure S5.95.** MS/MS spectrum of a SIFamide neuropeptide with the sequence RKPPFNGSIF(-.98) detected in the peptidergic signaling system of the American lobster *Homarus american*us generated by PEAKS XPro software.

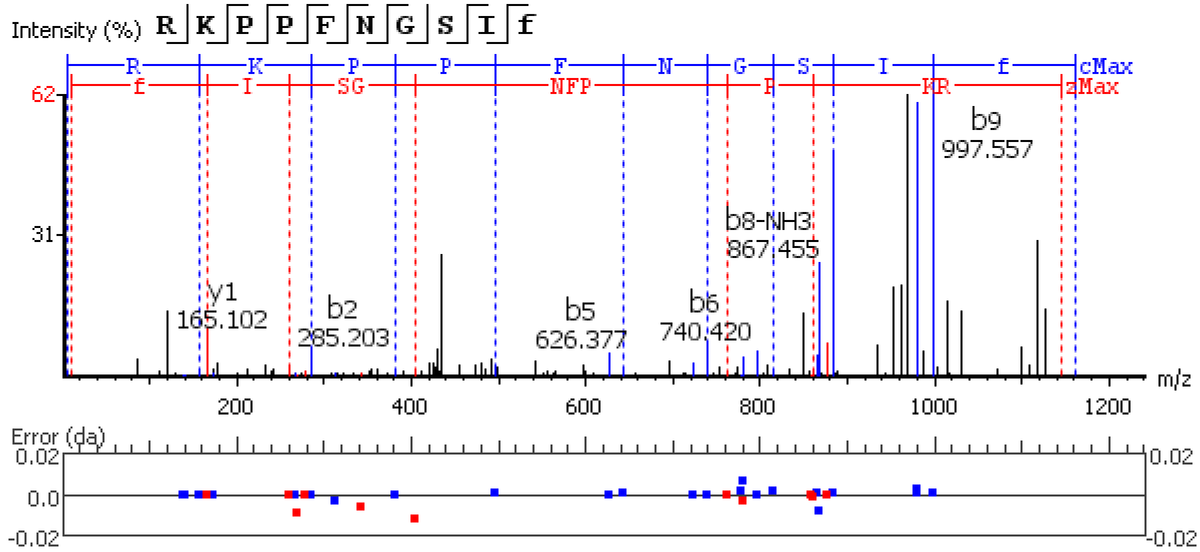


**Figure S5.96.** MS/MS spectrum of a Tachykinin neuropeptide with the sequence APSGFLGMR(-.98) detected in the peptidergic signaling system of the American lobster *Homarus american*us generated by PEAKS XPro software.

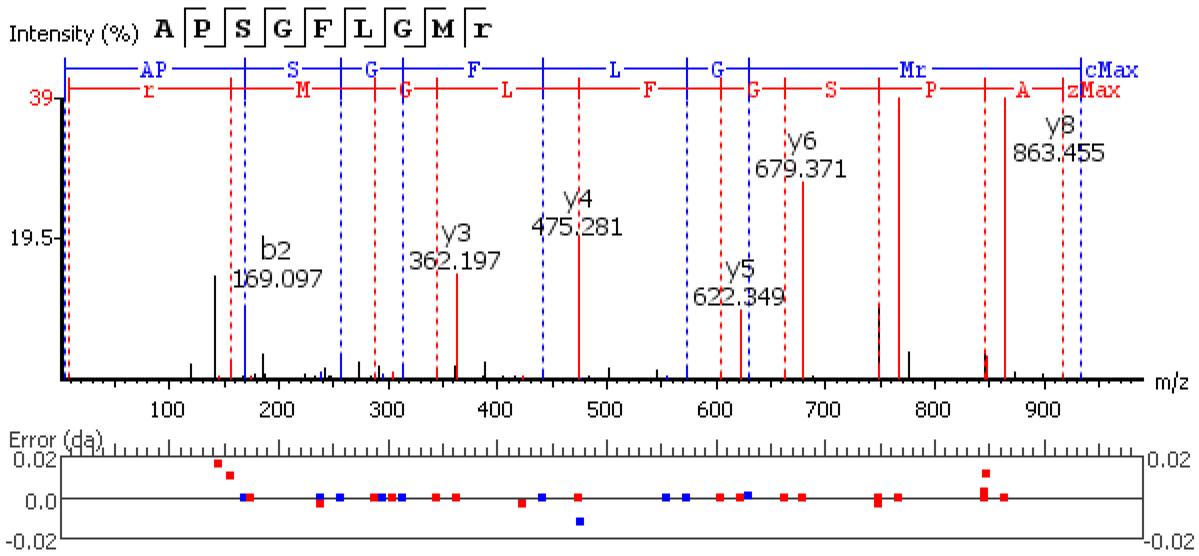


**Figure S5.97.** MS/MS spectrum of a Tachykinin neuropeptide with the sequence APSGFLGM(+15.99)R(-.98) detected in the peptidergic signaling system of the American lobster *Homarus american*us generated by PEAKS XPro software.

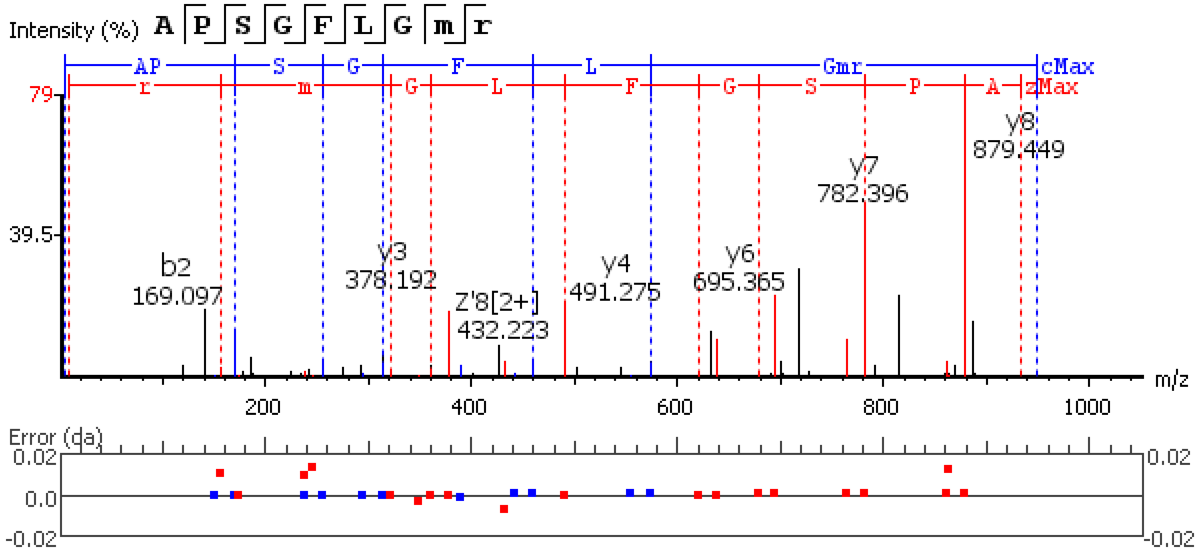


**Figure S5.98.** MS/MS spectrum of a Tachykinin neuropeptide with the sequence AGEGQDTPQDRE detected in the peptidergic signaling system of the American lobster *Homarus american*us generated by PEAKS XPro software.

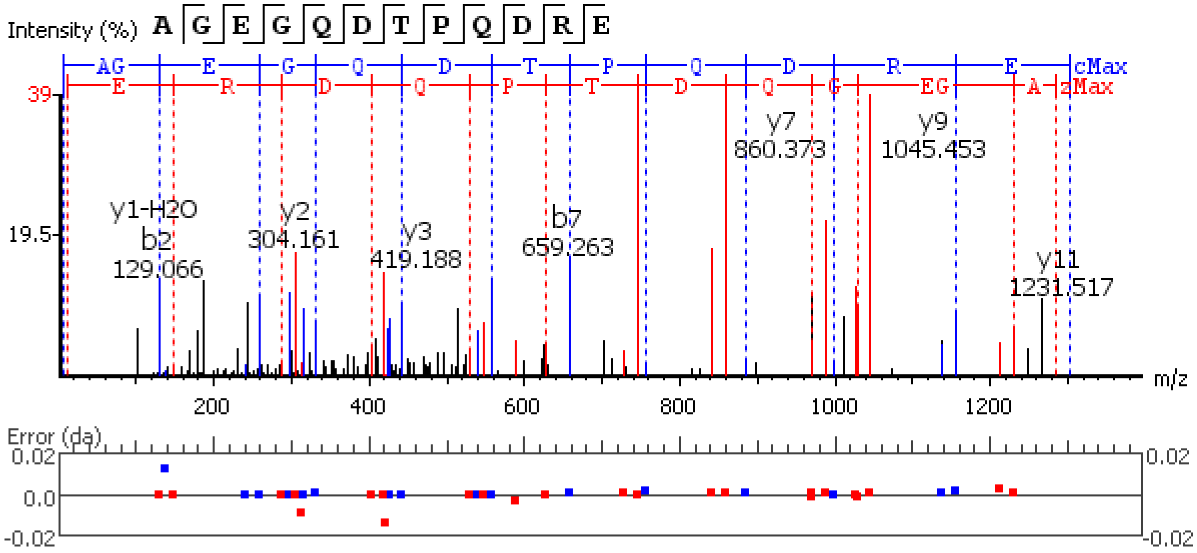


**Figure S5.99.** MS/MS spectrum of a Tachykinin neuropeptide with the sequence SDEEVFSDATADNDLEILL detected in the peptidergic signaling system of the American lobster *Homarus american*us generated by PEAKS XPro software.

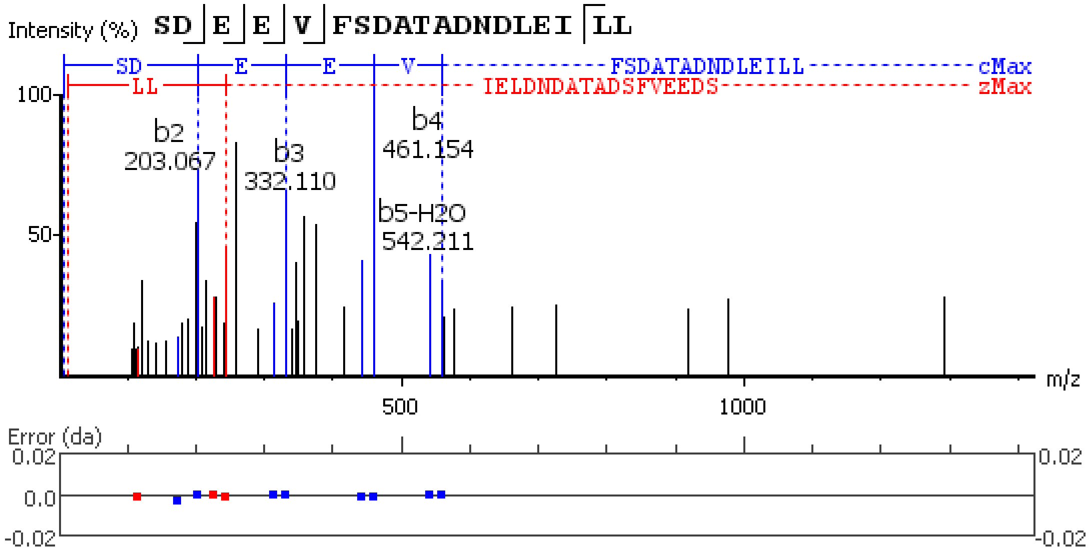


**Figure S5.100.** MS/MS spectrum of a Tachykinin neuropeptide with the sequence YYDDDSDMDAYIQALTAVVDGQQQQ detected in the peptidergic signaling system of the American lobster *Homarus american*us generated by PEAKS XPro software.

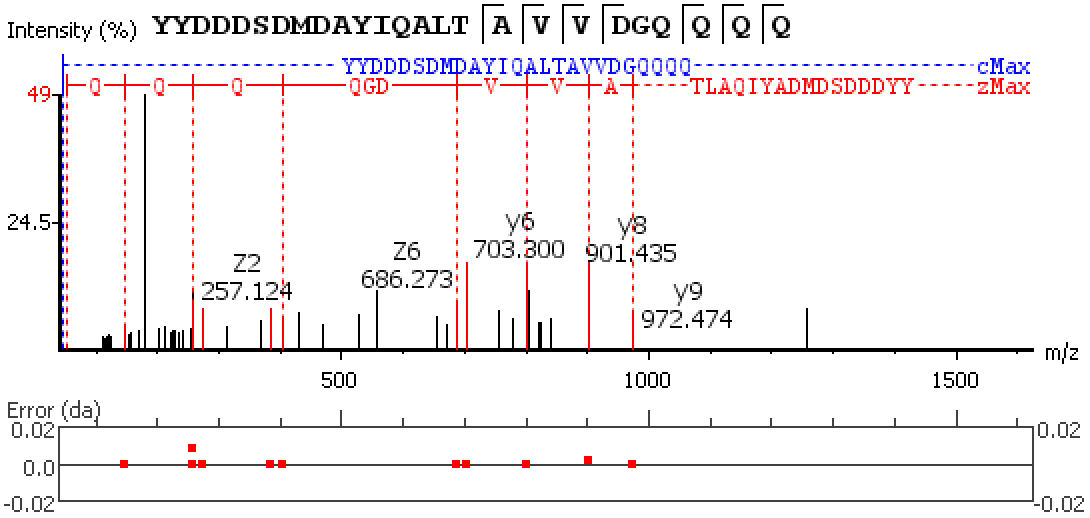


**Figure S5.101.** MS/MS spectrum of a Tachykinin neuropeptide with the sequence AYYSENPDEEISM(+15.99)TGVD detected in the peptidergic signaling system of the American lobster *Homarus american*us generated by PEAKS XPro software.

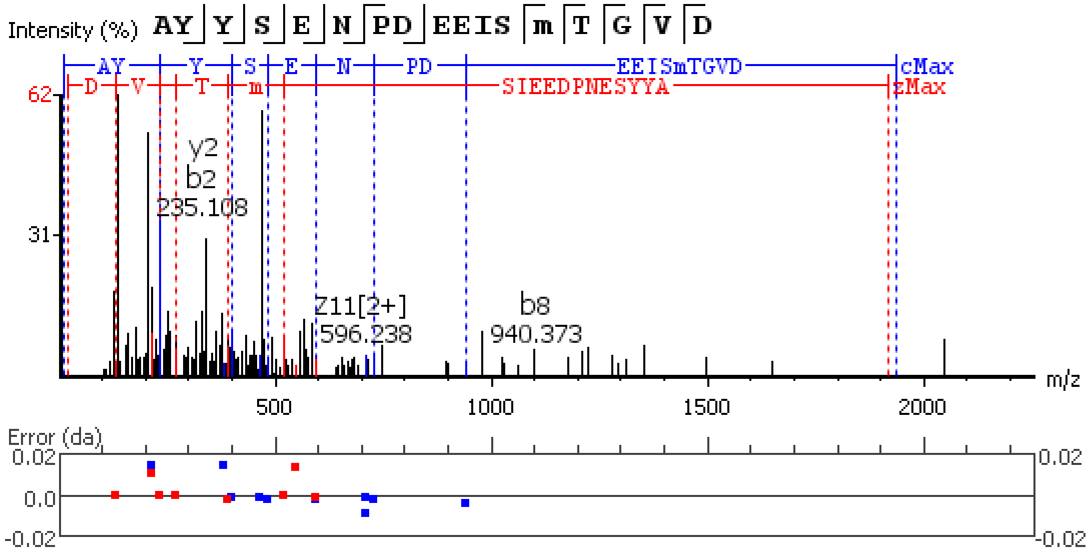


**Figure S5.102.** MS/MS spectrum of a Tachykinin neuropeptide with the sequence AYYSENPDEEISMTGVD detected in the peptidergic signaling system of the American lobster *Homarus american*us generated by PEAKS XPro software.

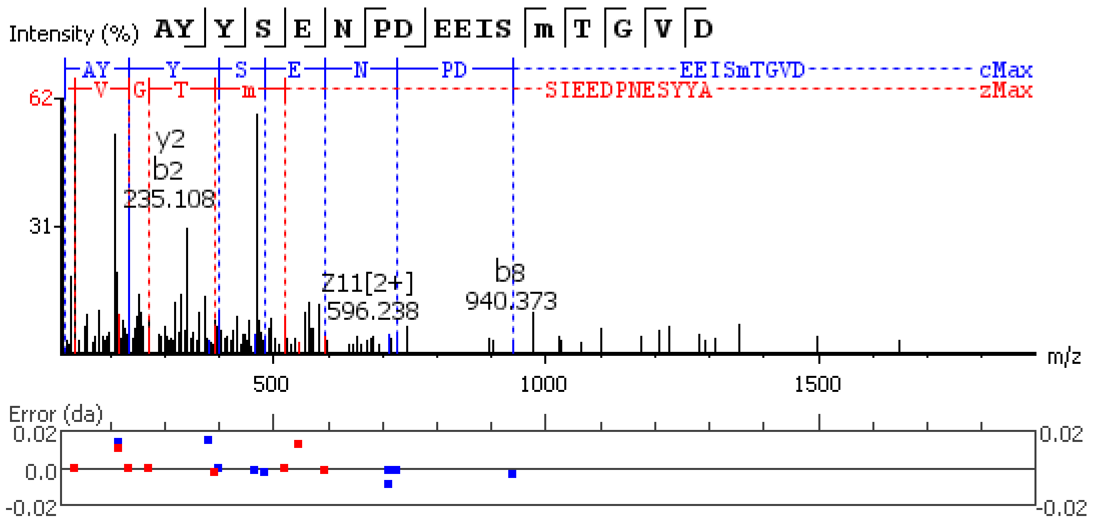


**Figure S5.103.** MS/MS spectrum of a Tachykinin neuropeptide with the sequence TPSGFLGMR(-.98) detected in the peptidergic signaling system of the American lobster *Homarus americanus* generated by PEAKS XPro software.

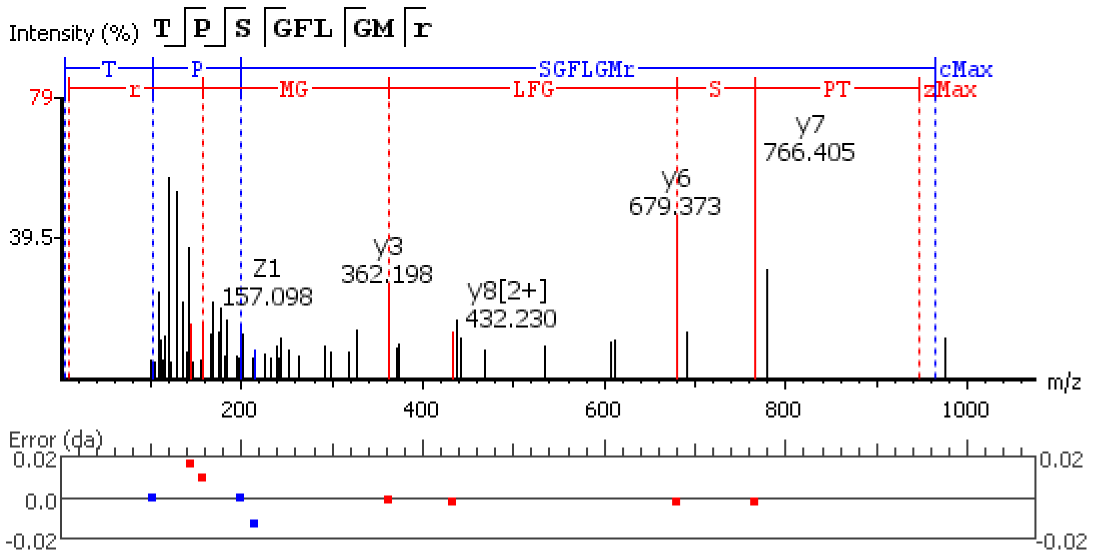


**Figure S5.104.** MS/MS spectrum of a Tachykinin neuropeptide with the sequence TPSGFLGM(+15.99)R(-.98) detected in the peptidergic signaling system of the American lobster *Homarus americanus* generated by PEAKS XPro software.

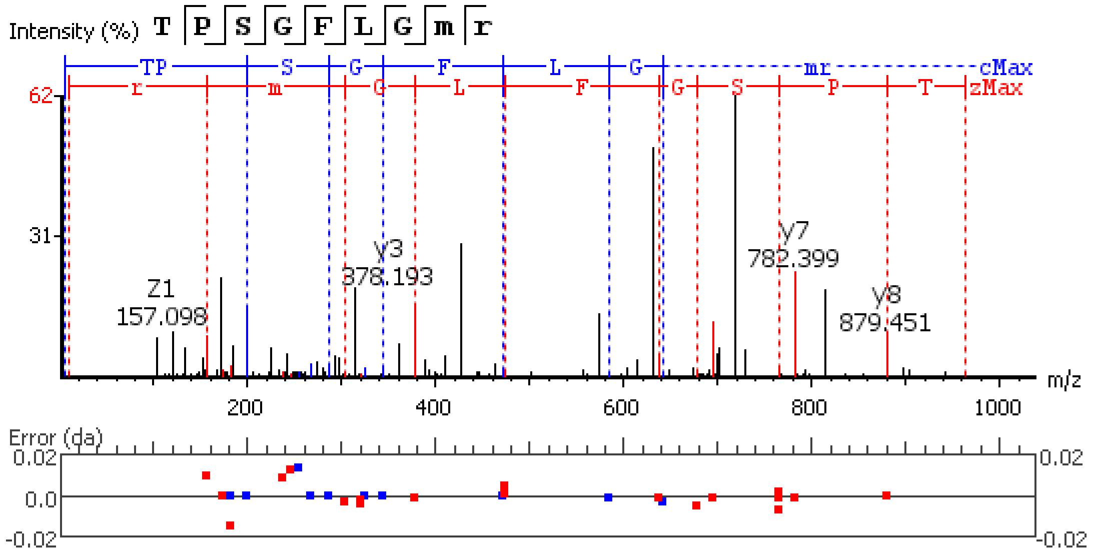


**Figure S5.105.** MS/MS spectrum of a Natalisin neuropeptide with the sequence PSSELLHQHHQ detected in the peptidergic signaling system of the American lobster *Homarus americanus* generated by PEAKS XPro software.

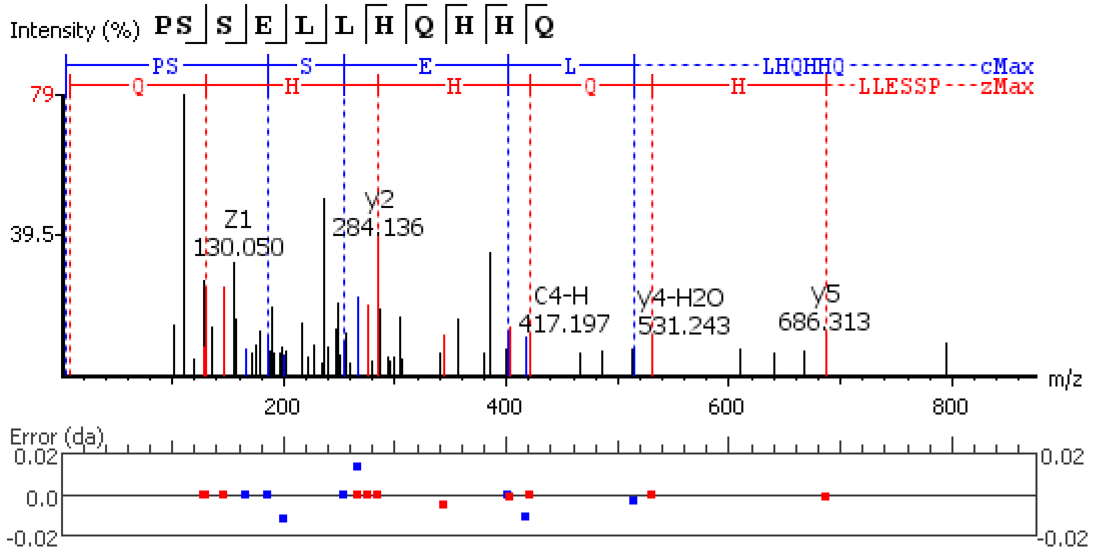


**Figure S5.106.** MS/MS spectrum of a Natalisin neuropeptide with the sequence DGGGPFWIAR(-.98) detected in the peptidergic signaling system of the American lobster *Homarus americanus* generated by PEAKS XPro software.

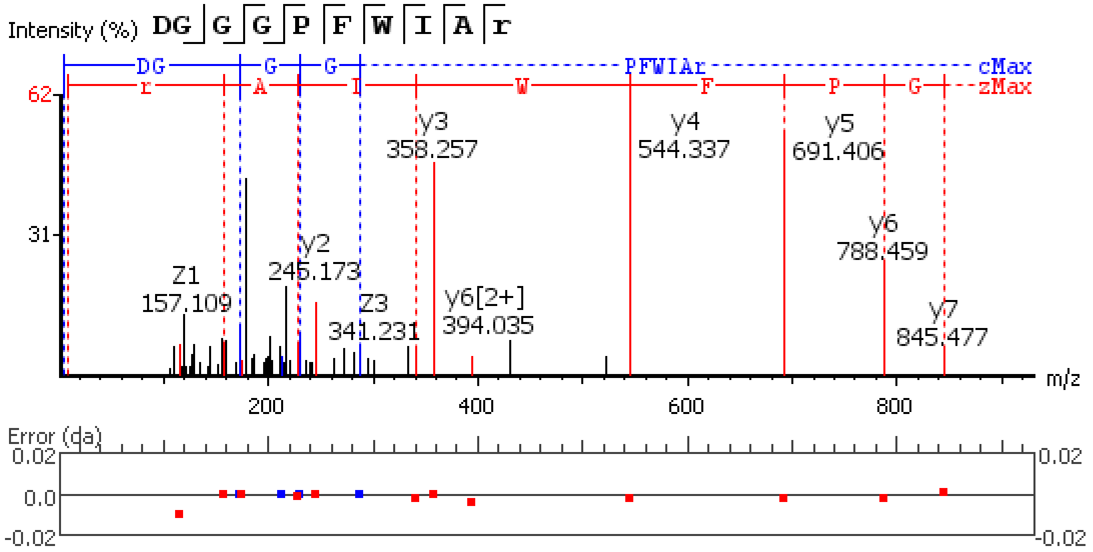


**Figure S5.107.** MS/MS spectrum of a Natalisin neuropeptide with the sequence

Q(-17.03)ETEGNGGPFWIAR(-.98) detected in the peptidergic signaling system of the American lobster *Homarus american*us generated by PEAKS XPro software.

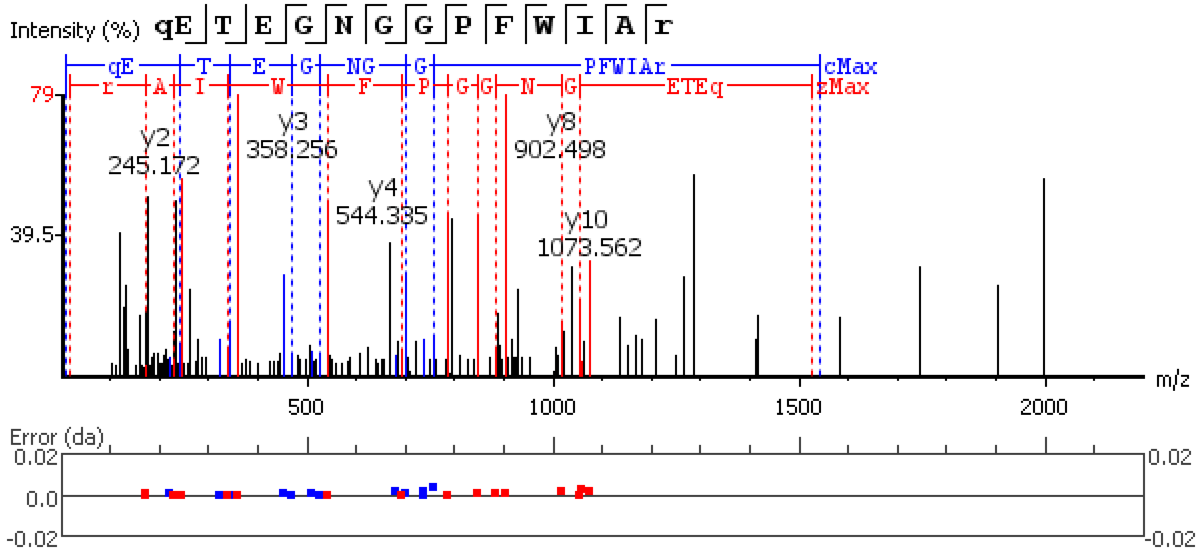


**Figure S5.108.** MS/MS spectrum of a Natalisin neuropeptide with the sequence

Q(-17.03)DGGPFWISR(-.98) detected in the peptidergic signaling system of the American lobster *Homarus american*us generated by PEAKS XPro software.

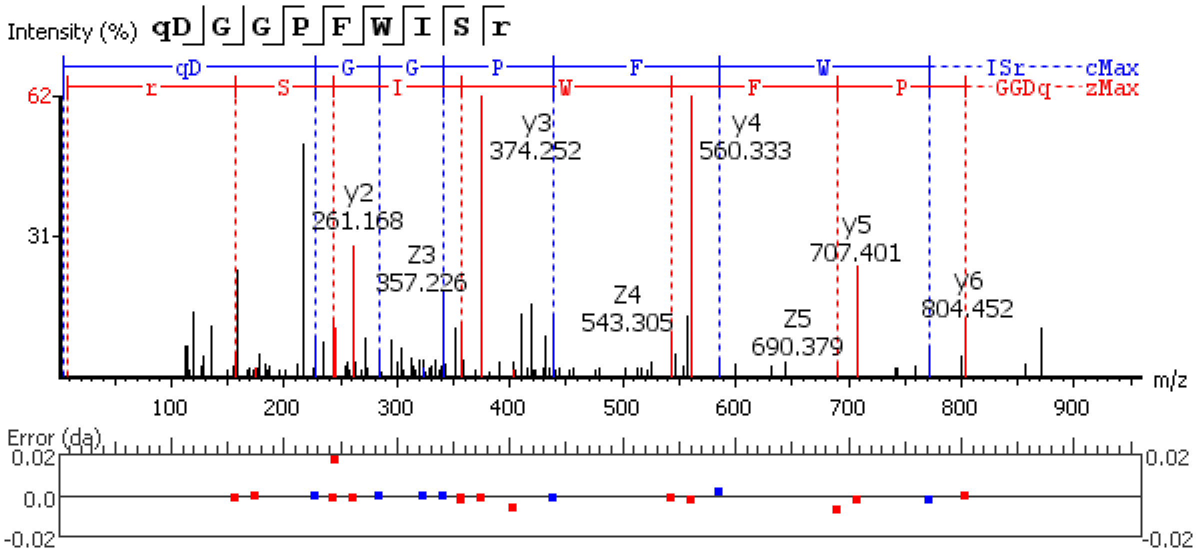


**Figure S5.109.** MS/MS spectrum of a Natalisin neuropeptide with the sequence QDGGPFWISR(-.98) detected in the peptidergic signaling system of the American lobster *Homarus americanus* generated by PEAKS XPro software.

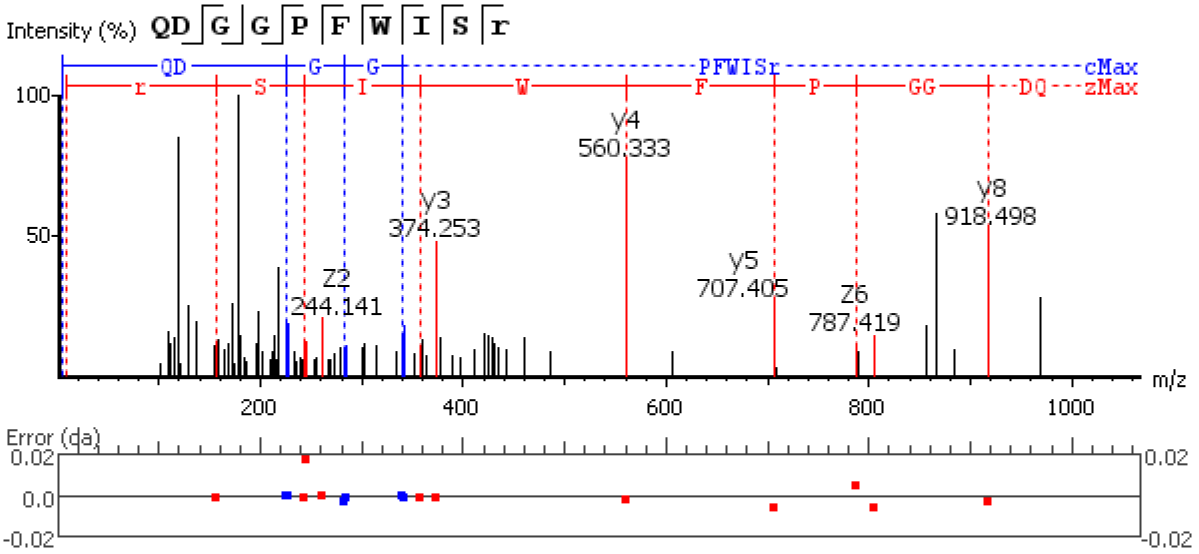


**Figure S5.110.** MS/MS spectrum of a Natalisin neuropeptide with the sequence EGEAPPFWVSR(-.98) detected in the peptidergic signaling system of the American lobster *Homarus americanus* generated by PEAKS XPro software.

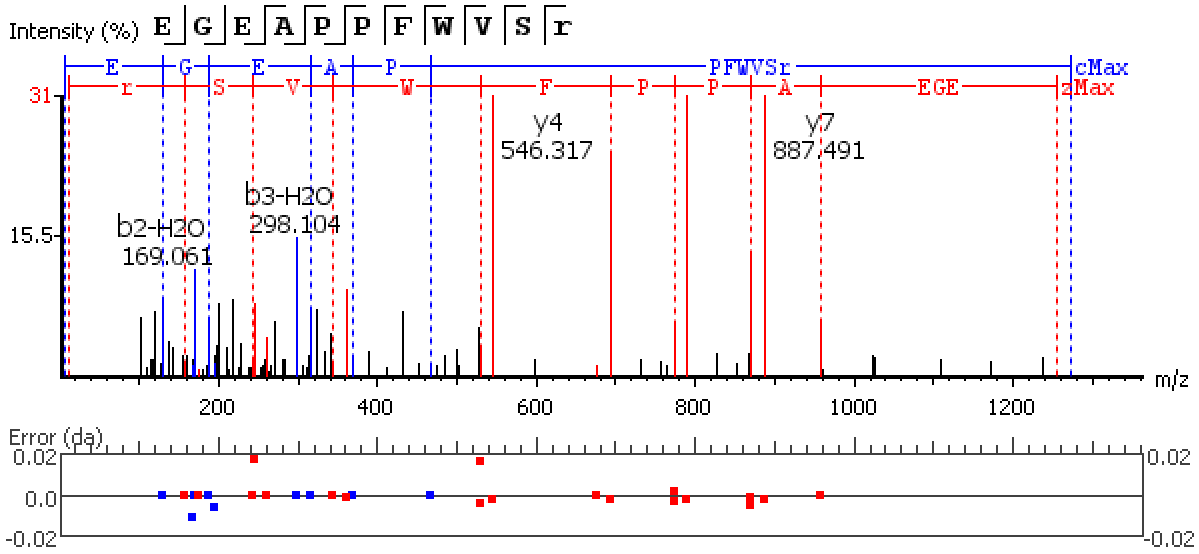


**Figure S5.111.** MS/MS spectrum of a Natalisin neuropeptide with the sequence EGEETHPFWVSR(-.98) detected in the peptidergic signaling system of the American lobster *Homarus american*us generated by PEAKS XPro software.

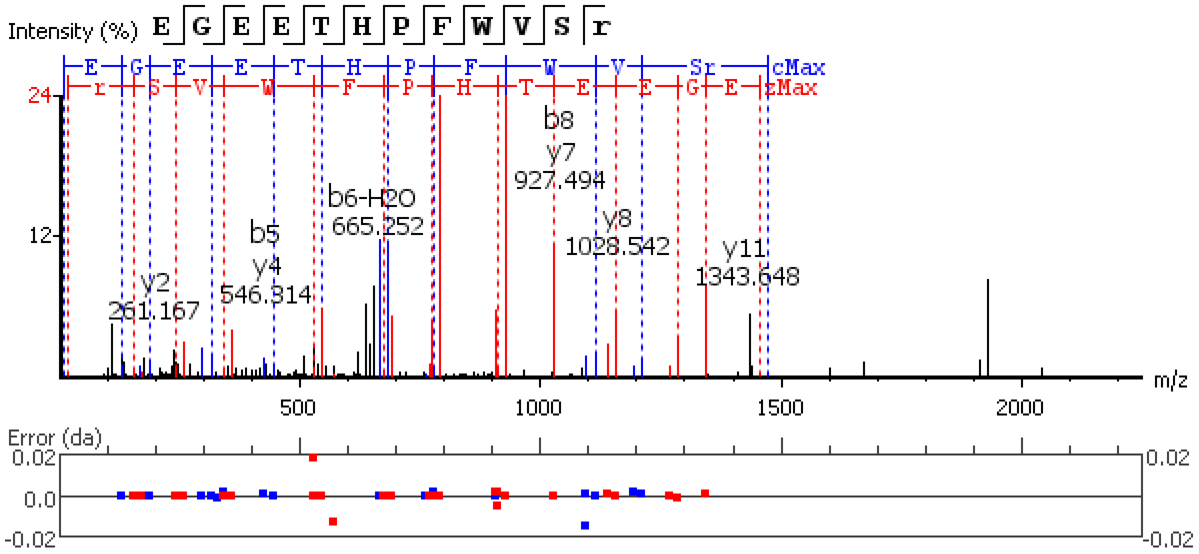


**Figure S5.112.** MS/MS spectrum of a Natalisin neuropeptide with the sequence E(-18.01)GEETHPFWVSR(-.98) detected in the peptidergic signaling system of the American lobster *Homarus american*us generated by PEAKS XPro software.

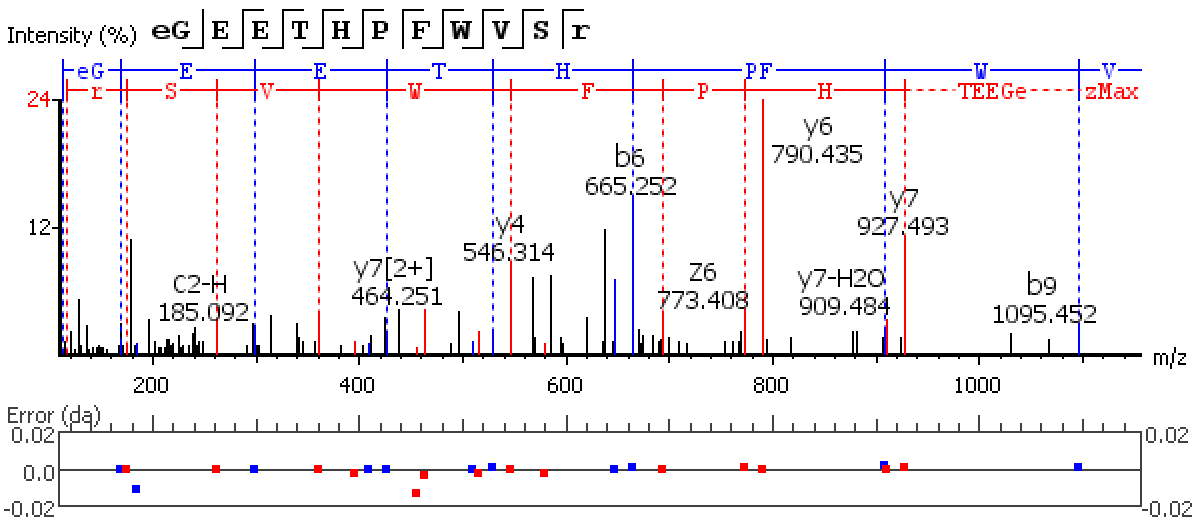


**Figure S5.113.** MS/MS spectrum of a Natalisin neuropeptide with the sequence DAVDGRAPFWISR(-.98) detected in the peptidergic signaling system of the American lobster *Homarus american*us generated by PEAKS XPro software.

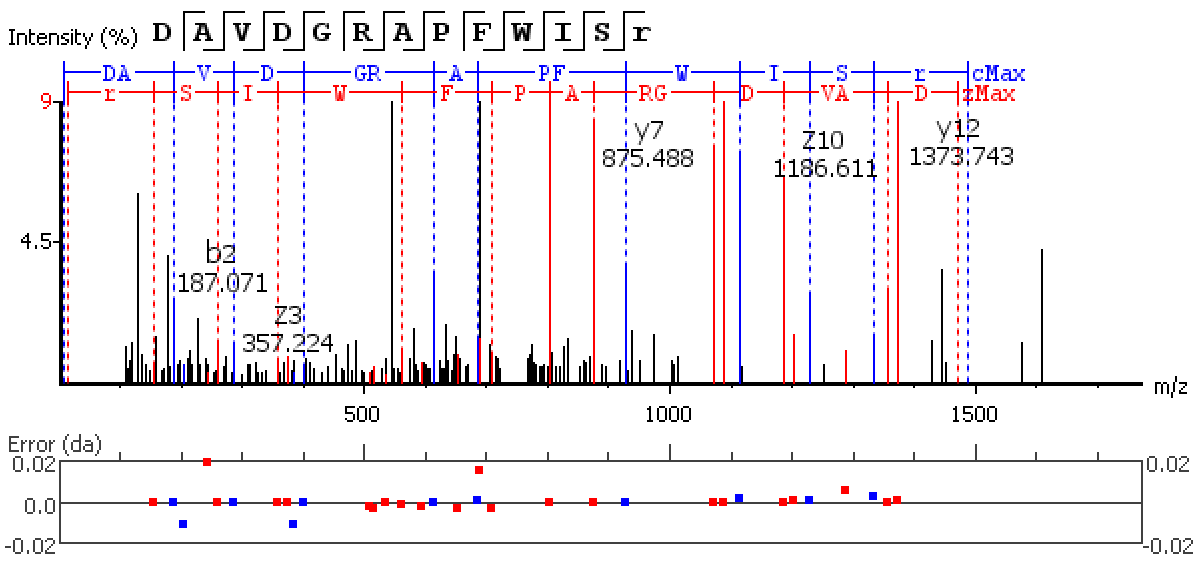


**Figure S5.114.** MS/MS spectrum of a Natalisin neuropeptide with the sequence DTPALLPVGHPSLWGNR(-.98) detected in the peptidergic signaling system of the American lobster *Homarus american*us generated by PEAKS XPro software.

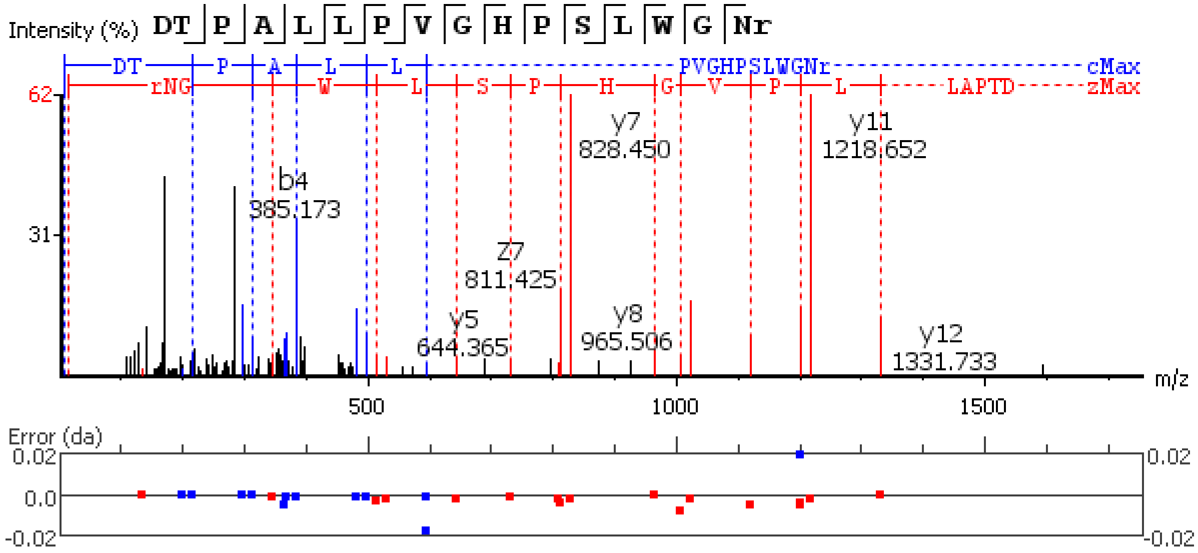

**Figure S5.115.** MS/MS spectrum of a Natalisin neuropeptide with the sequence DTTYGPIDDPFVKGFLALR(-.98) detected in the peptidergic signaling system of the American lobster *Homarus american*us generated by PEAKS XPro software.

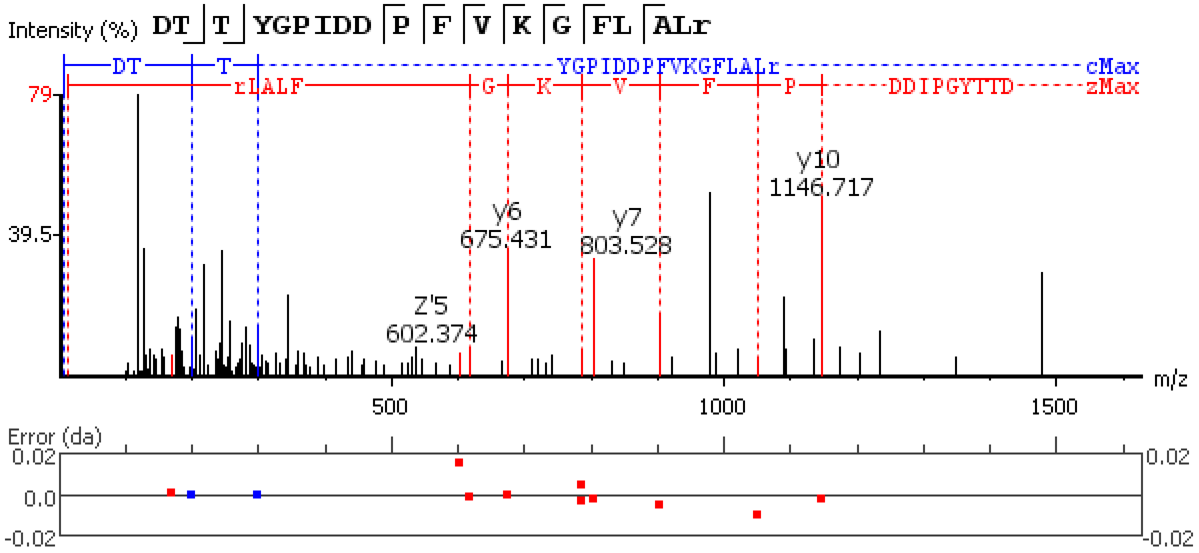

**Figure S5.116.** MS/MS spectrum of a RYamide neuropeptide with the sequence

Q(-17.03)GFYTQRY(-.98) detected in the peptidergic signaling system of the American lobster *Homarus american*us generated by PEAKS XPro software.

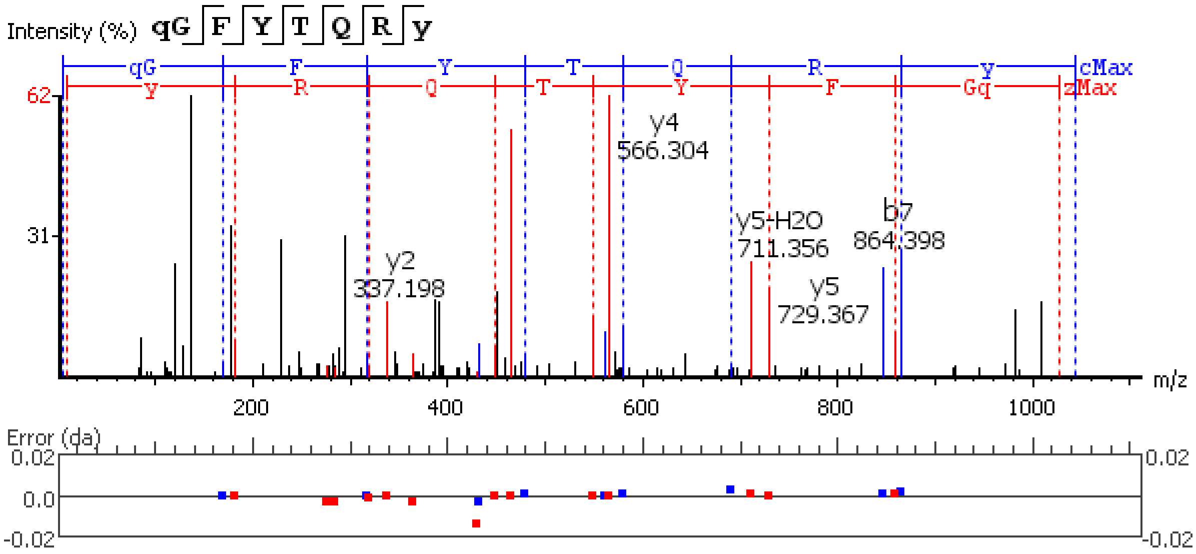


**Figure S5.117.** MS/MS spectrum of a RYamide neuropeptide with the sequence QGFYTQRY(-.98) detected in the peptidergic signaling system of the American lobster *Homarus american*us generated by PEAKS XPro software.

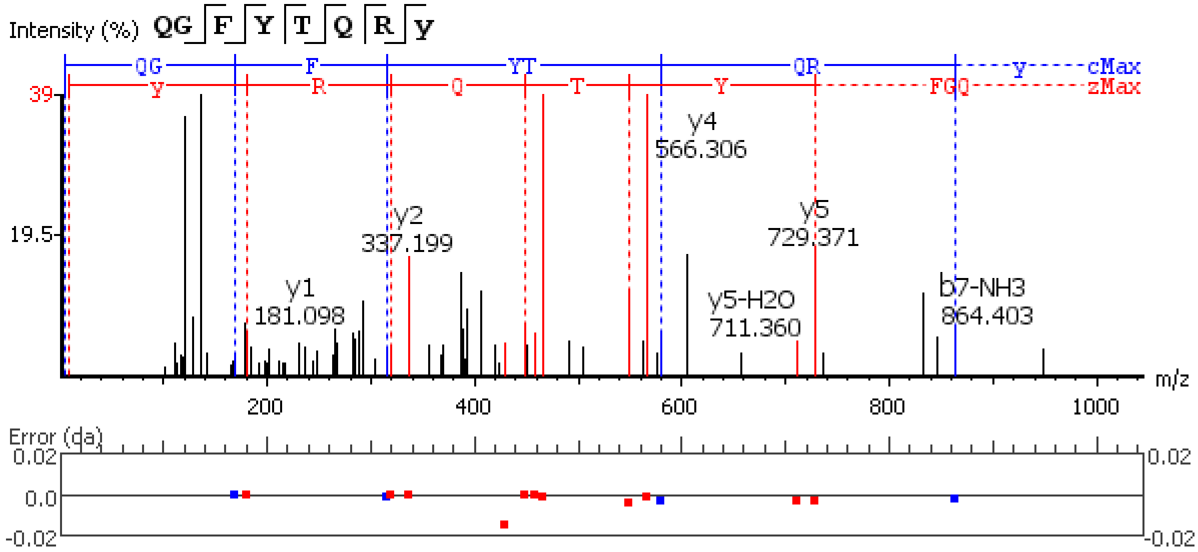


**Figure S5.118.** MS/MS spectrum of a RYamide neuropeptide with the sequence SGFYANRN(-.98) detected in the peptidergic signaling system of the American lobster *Homarus american*us generated by PEAKS XPro software.

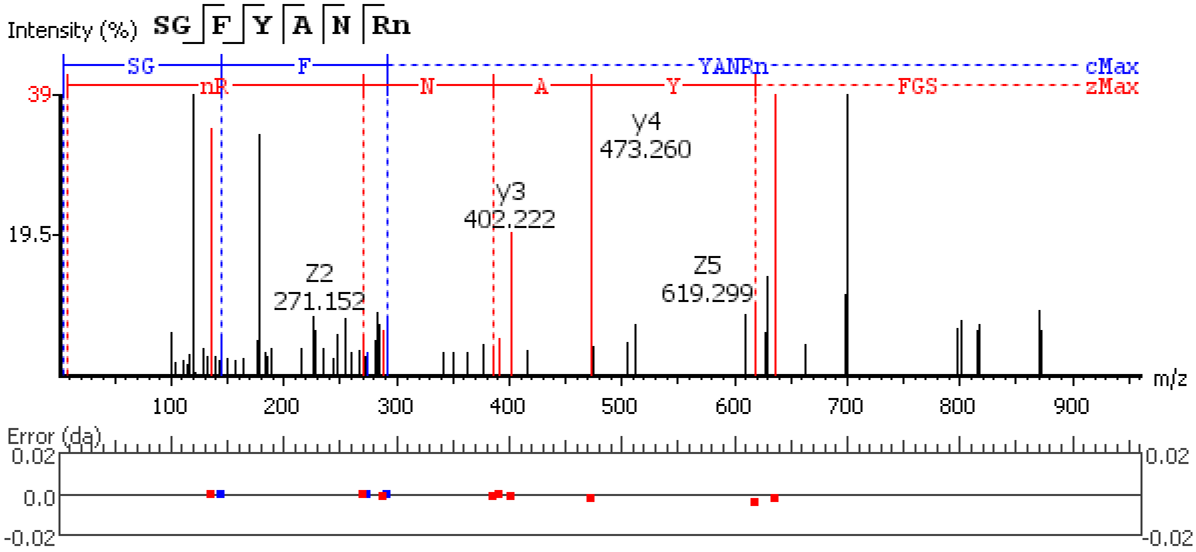

**Figure S5.119.** MS/MS spectrum of a RYamide neuropeptide with the sequence SSPSQGLPEIKIRSSRFIGGSRY(-.98) detected in the peptidergic signaling system of the American lobster *Homarus americanus* generated by PEAKS XPro software.

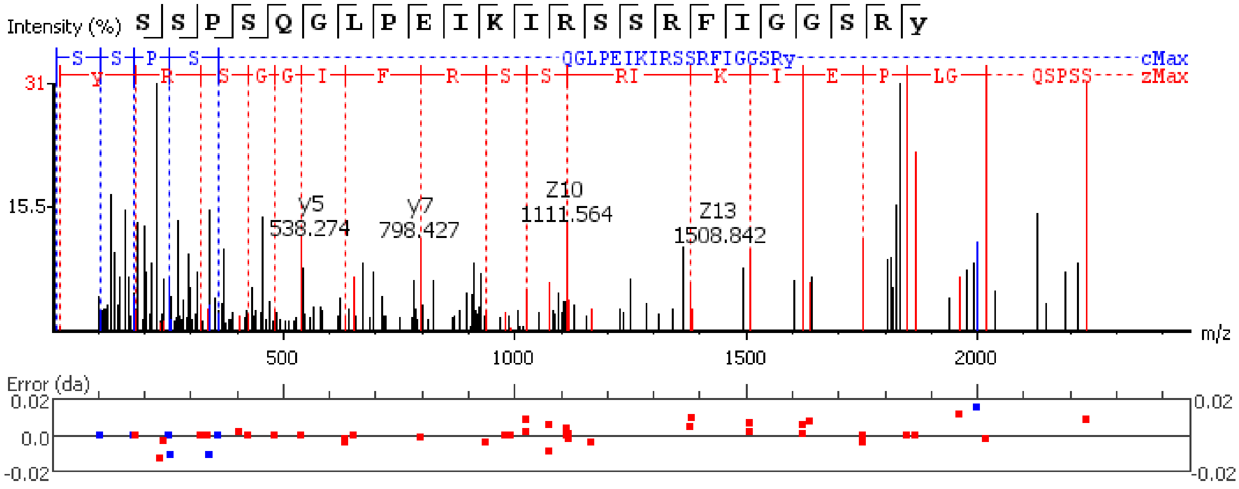

**Figure S5.120.** MS/MS spectrum of a RYamide neuropeptide with the sequence SSPSQGLPEI detected in the peptidergic signaling system of the American lobster *Homarus americanus* generated by PEAKS XPro software.

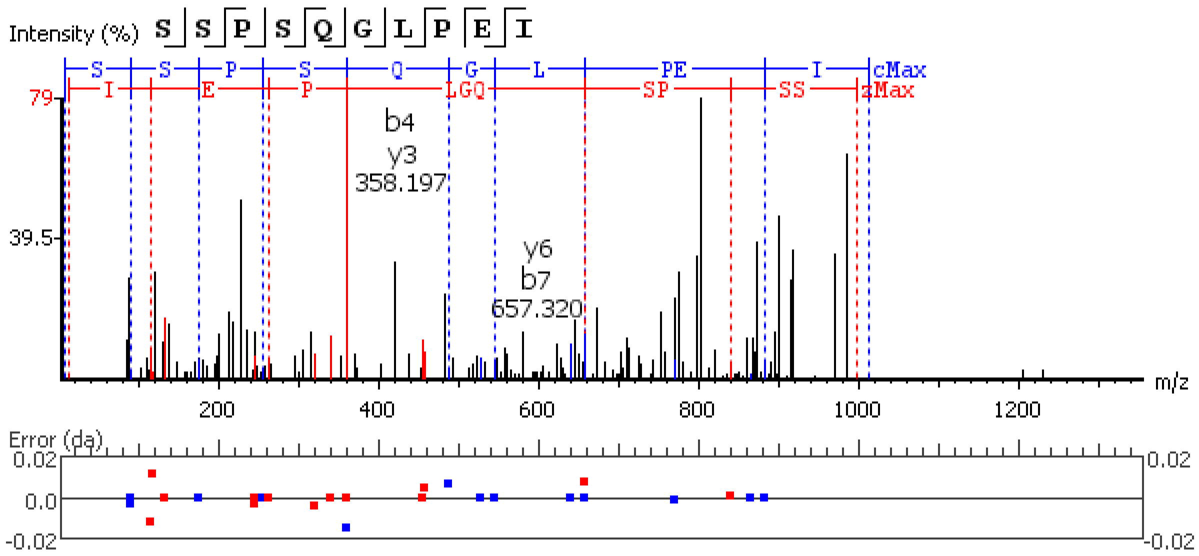


**Figure S5.121.** MS/MS spectrum of a RYamide neuropeptide with the sequence SSRFIGGSRY(-.98) detected in the peptidergic signaling system of the American lobster *Homarus americanus* generated by PEAKS XPro software.

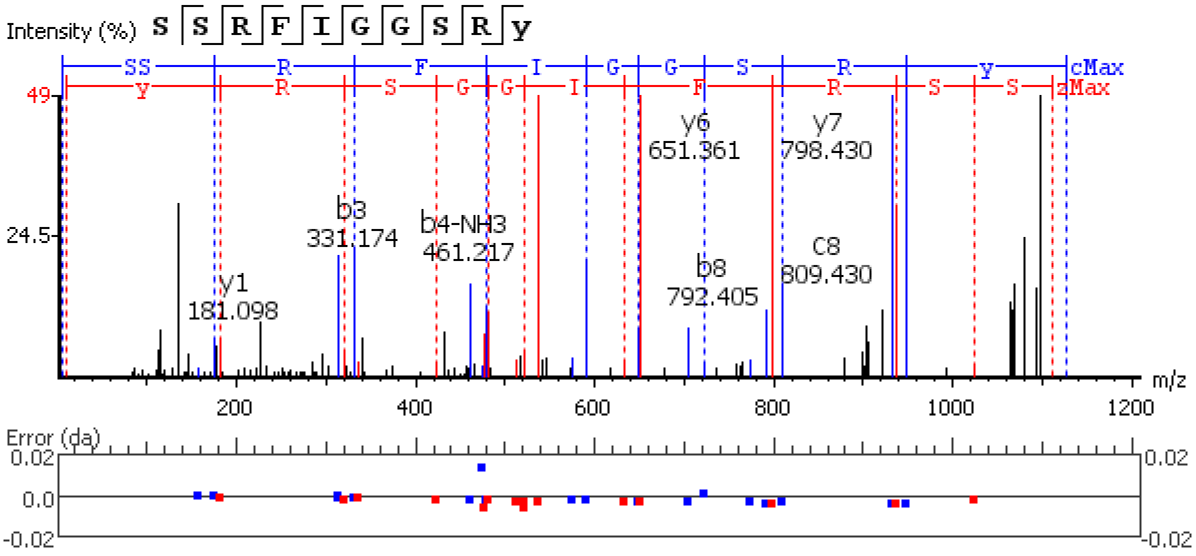


**Figure S5.122.** MS/MS spectrum of a Gonadoliberin neuropeptide with the sequence

Q(-17.03)IHWNRGWGAGGSM(+15.99)(-.98) detected in the peptidergic signaling system of the American lobster *Homarus americanus* generated by PEAKS XPro software.

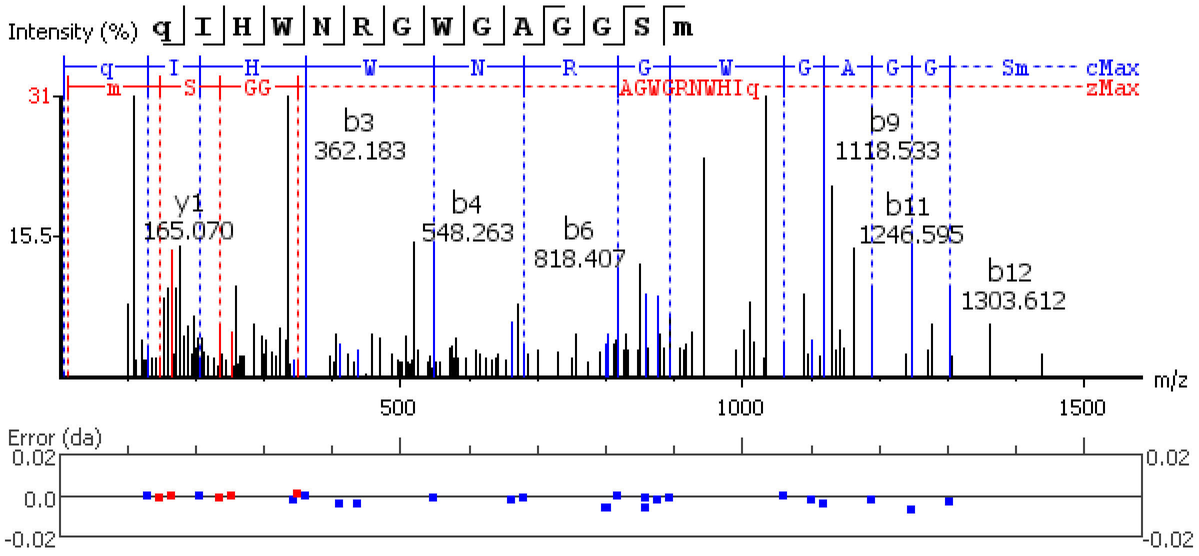


**Figure S5.123.** MS/MS spectrum of a Gonadoliberin neuropeptide with the sequence

Q(-17.03)IHWNRGWGAGGSM(-.98) detected in the peptidergic signaling system of the American lobster *Homarus american*us generated by PEAKS XPro software.

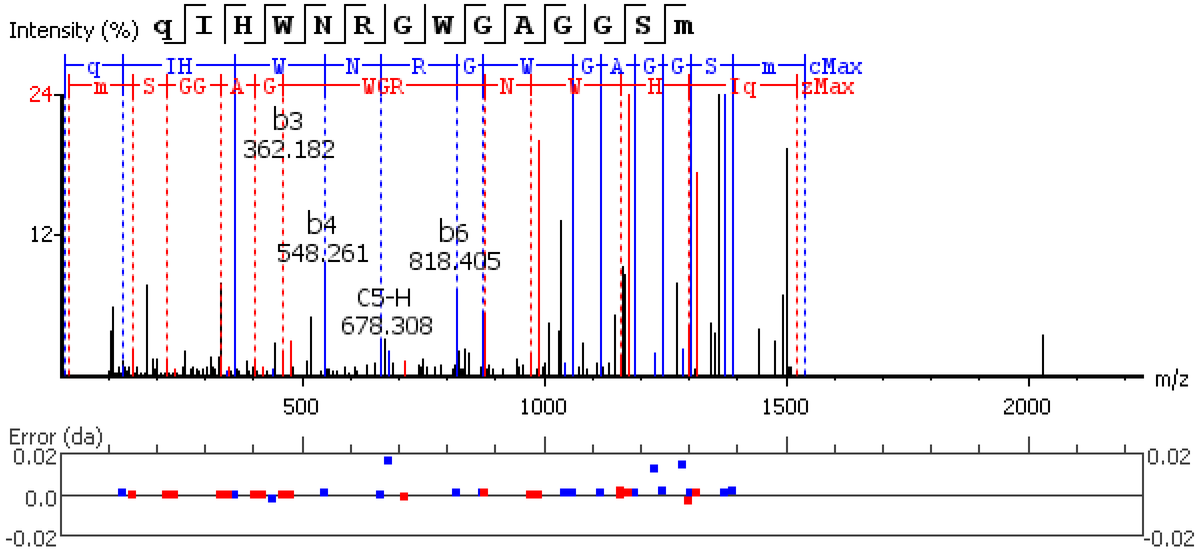


**Figure S5.124.** MS/MS spectrum of a Adipokinetic hormone corazonin-like (ACP) neuropeptide with the sequence Q(-17.03)ITFSRSWVPQ(-.98) detected in the peptidergic signaling system of the American lobster *Homarus american*us generated by PEAKS XPro software.

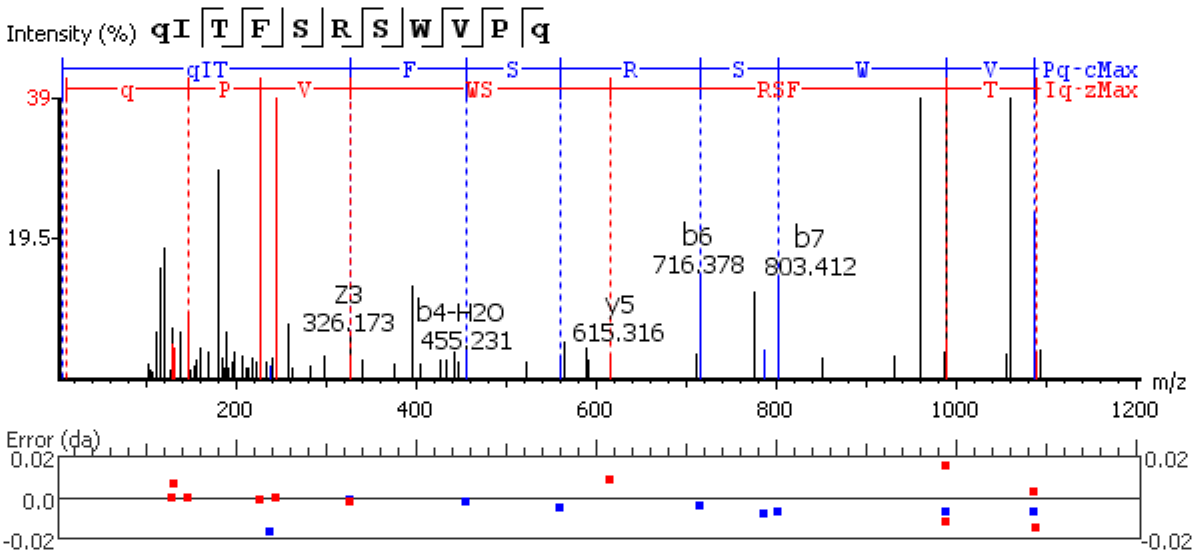


**Figure S5.125.** MS/MS spectrum of a Red Pigment Concentrating Hormone neuropeptide with the sequence Q(-17.03)LNFSPGW(-.98) detected in the peptidergic signaling system of the American lobster *Homarus american*us generated by PEAKS XPro software.

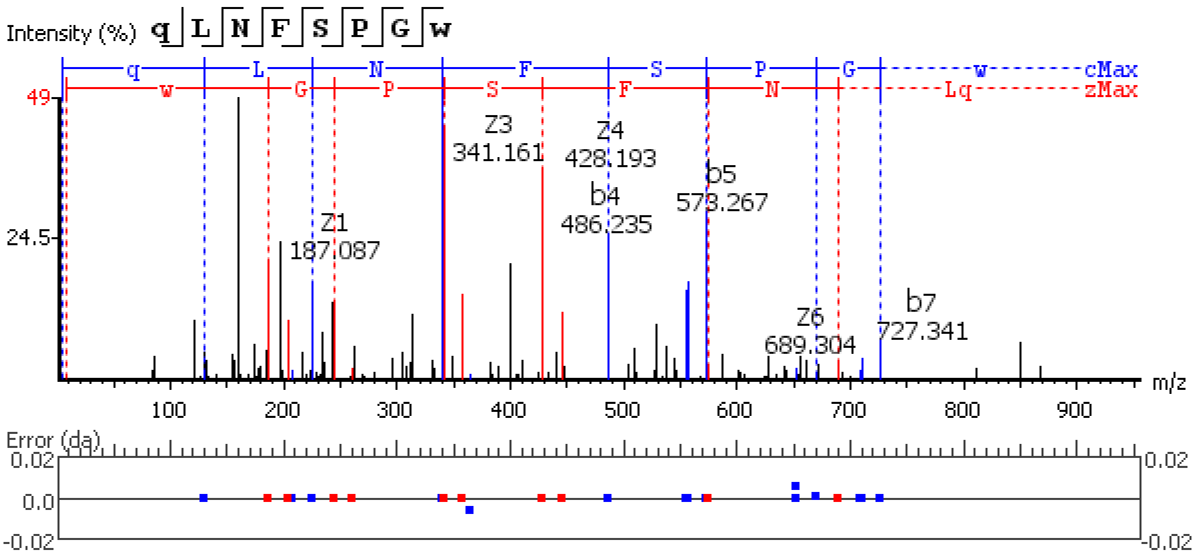


**Figure S5.126.** MS/MS spectrum of a Agatoxin-like peptide (ALP) neuropeptide with the sequence DDVAGSDPIK detected in the peptidergic signaling system of the American lobster *Homarus american*us generated by PEAKS XPro software.

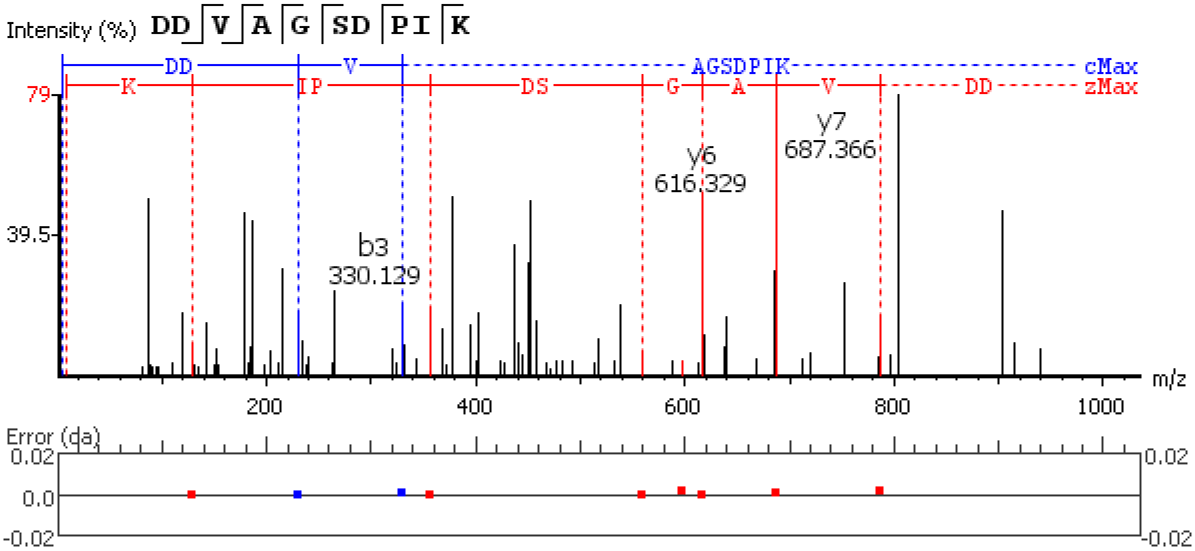


**Figure S5.127.** MS/MS spectrum of a Agatoxin-like peptide (ALP) neuropeptide with the sequence Q(-17.03)PLLEEGREEDGVQQAEPDYAADLLERLLARTQ detected in the peptidergic signaling system of the American lobster *Homarus american*us generated by PEAKS XPro software.

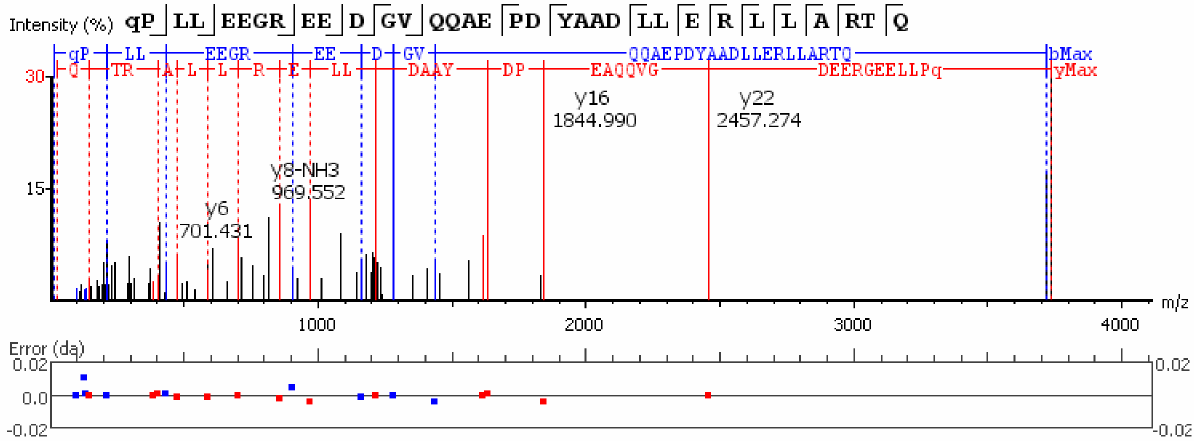


**Figure S5.128.** MS/MS spectrum of a Agatoxin-like peptide (ALP) neuropeptide with the sequence M(+15.99)GIFQQW(-.98) detected in the peptidergic signaling system of the American lobster *Homarus american*us generated by PEAKS XPro software.

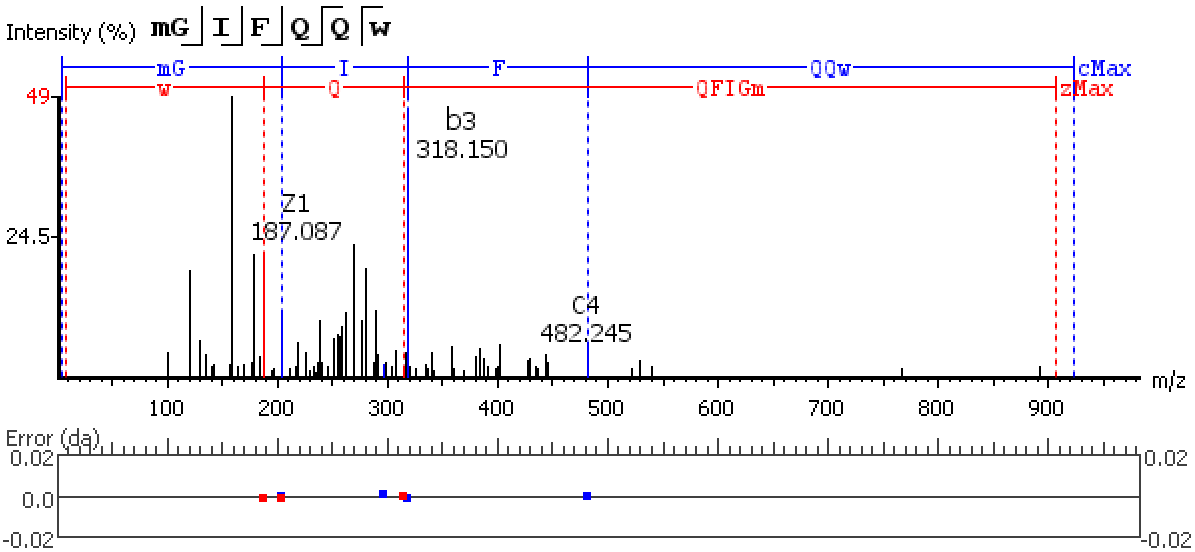


**Figure S5.129.** MS/MS spectrum of a Agatoxin-like peptide (ALP) neuropeptide with the sequence MGIFQQW(-.98) detected in the peptidergic signaling system of the American lobster *Homarus american*us generated by PEAKS XPro software.

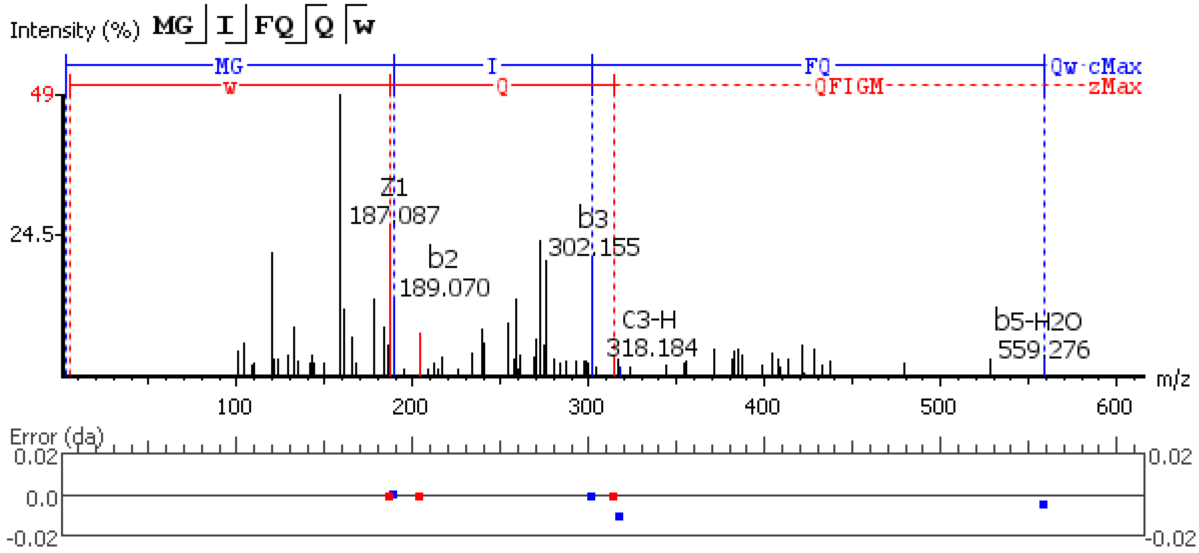


**Figure S5.130.** MS/MS spectrum of a Periviscerokinin neuropeptide with the sequence QDLIPFPRV(-.98) detected in the peptidergic signaling system of the American lobster *Homarus american*us generated by PEAKS XPro software.

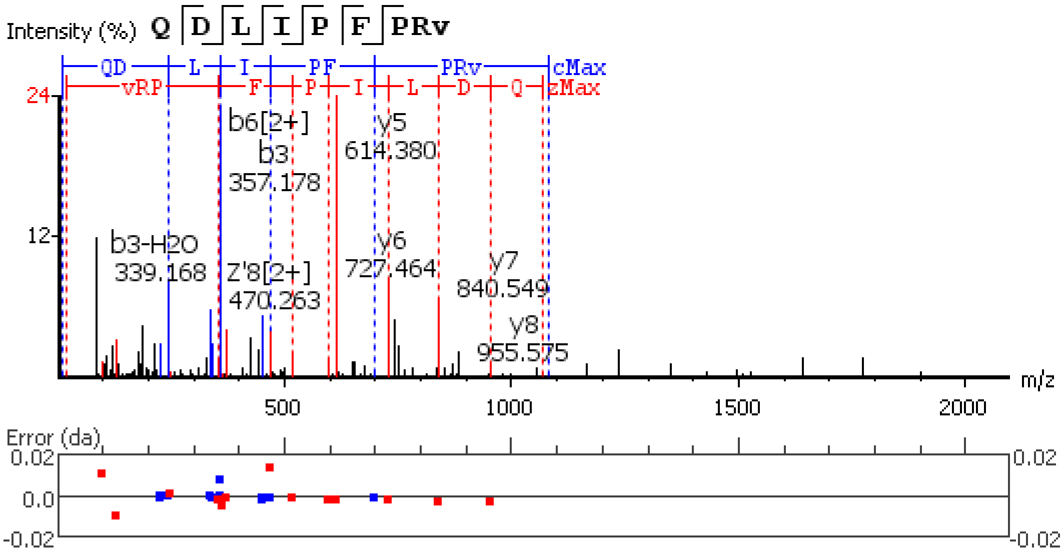


**Figure S5.131.** MS/MS spectrum of a Periviscerokinin neuropeptide with the sequence Q(-17.03)DLIPFPRV(-.98) detected in the peptidergic signaling system of the American lobster *Homarus american*us generated by PEAKS XPro software.

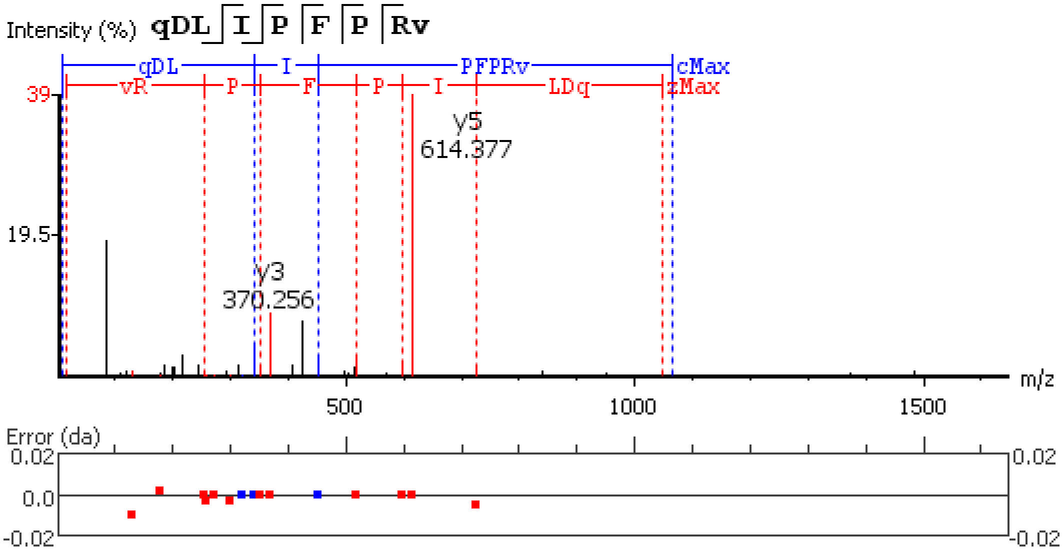


**Figure S5.132.** MS/MS spectrum of a Sulfakinin neuropeptide with the sequence EFDEYGHMRF(-.98) detected in the peptidergic signaling system of the American lobster *Homarus american*us generated by PEAKS XPro software.

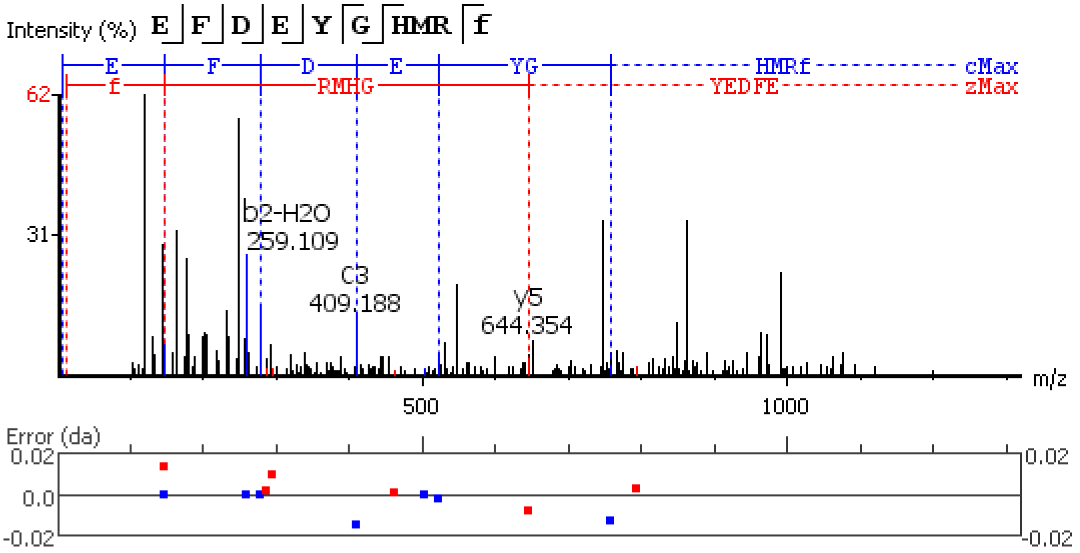


**Figure S5.133.** MS/MS spectrum of a Sulfakinin neuropeptide with the sequence EFDEYGHM(+15.99)RF(-.98) detected in the peptidergic signaling system of the American lobster *Homarus americanus* generated by PEAKS XPro software.

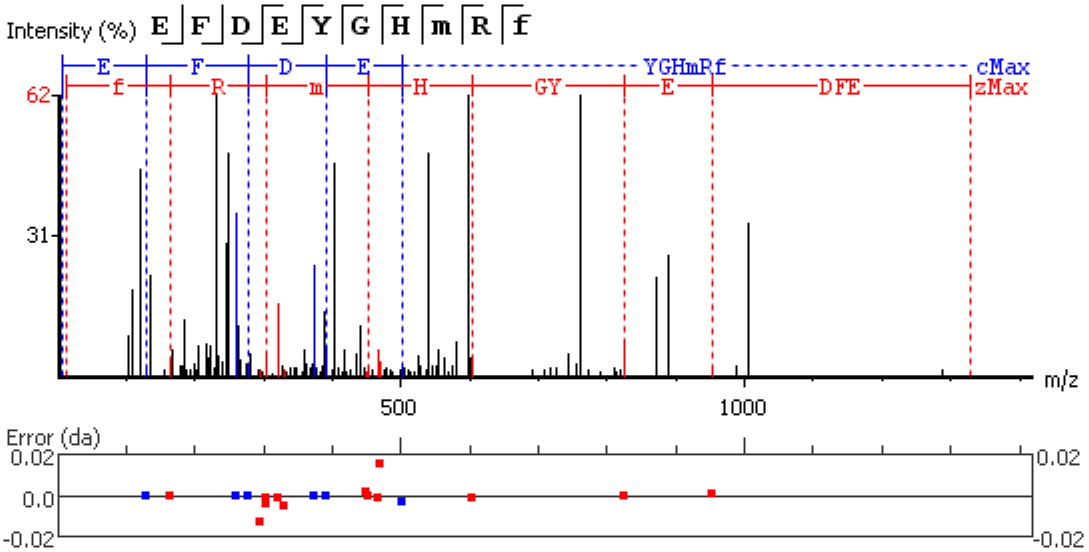


**Figure S5.134.** MS/MS spectrum of a Leucokinin neuropeptide with the sequence SDPLLPASQHEPNT detected in the peptidergic signaling system of the American lobster *Homarus americanus* generated by PEAKS XPro software.

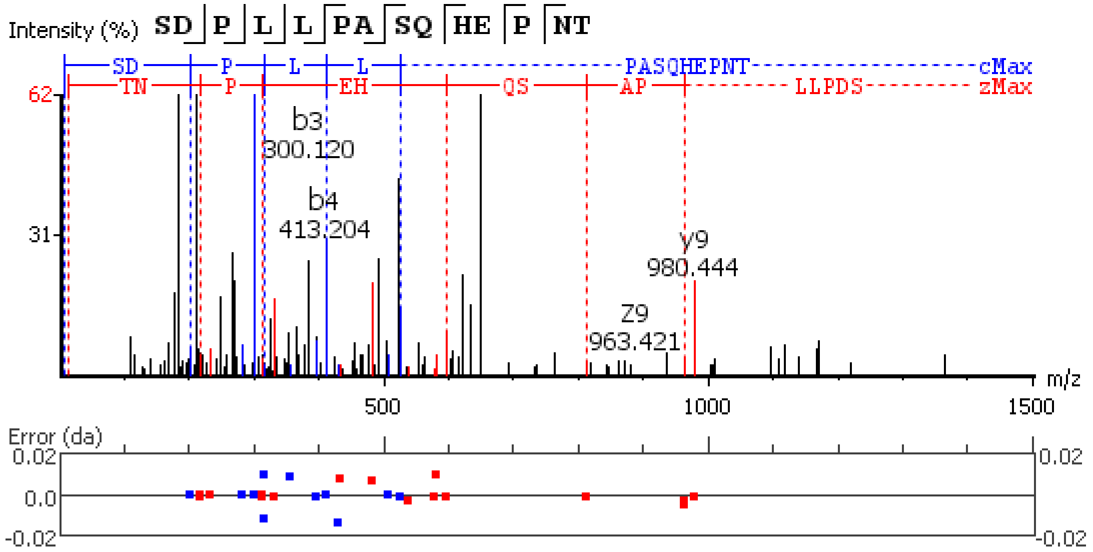


**Figure S5.135.** MS/MS spectrum of a Leucokinin neuropeptide with the sequence SPSMDLSGNQD detected in the peptidergic signaling system of the American lobster *Homarus american*us generated by PEAKS XPro software.

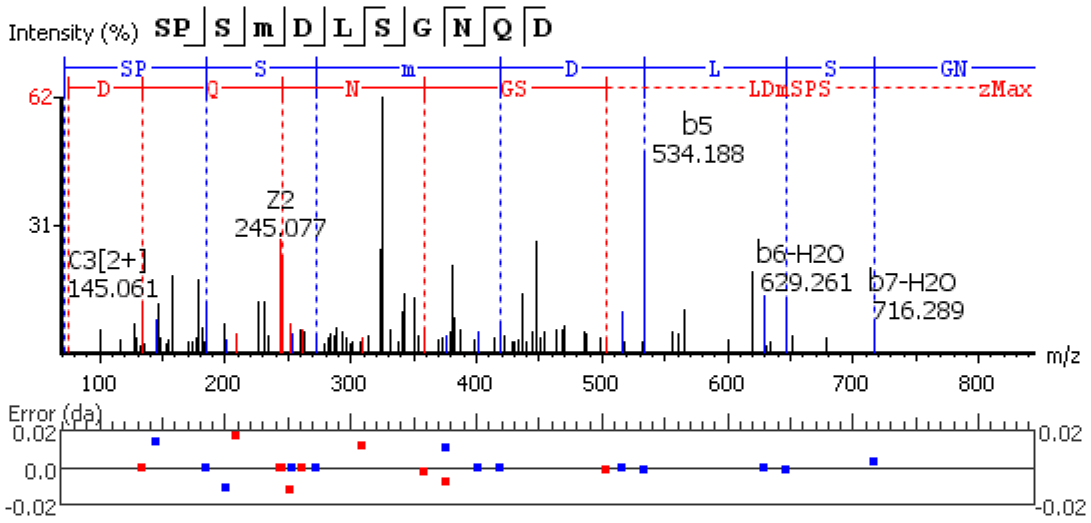


**Figure S5.136.** MS/MS spectrum of a Leucokinin neuropeptide with the sequence SPSM(+15.99)DLSGNQD detected in the peptidergic signaling system of the American lobster *Homarus american*us generated by PEAKS XPro software.

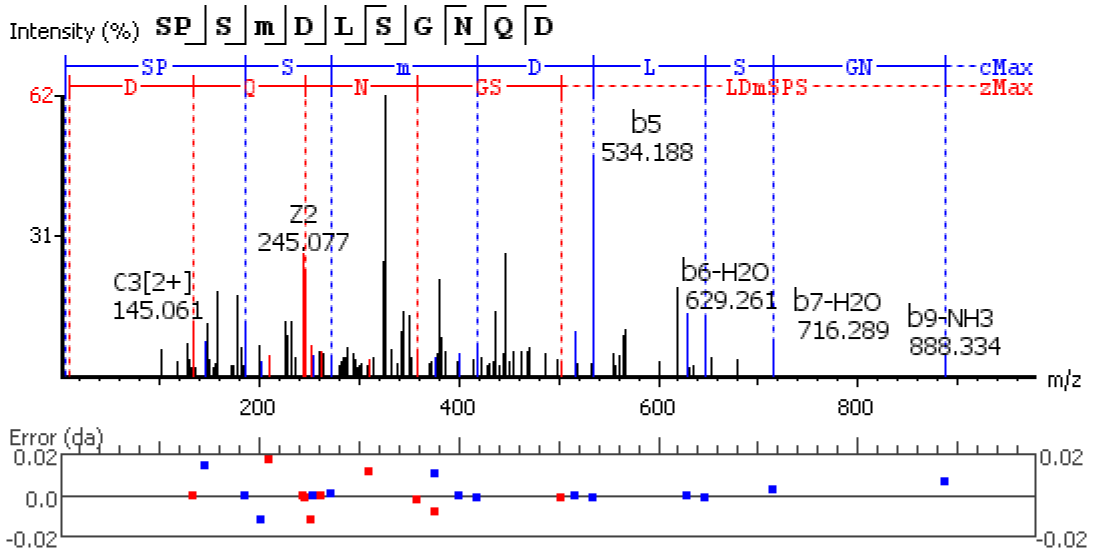


**Figure S5.137.** MS/MS spectrum of a Leucokinin neuropeptide with the sequence SSGDELDDHFLD detected in the peptidergic signaling system of the American lobster *Homarus americanus* generated by PEAKS XPro software.

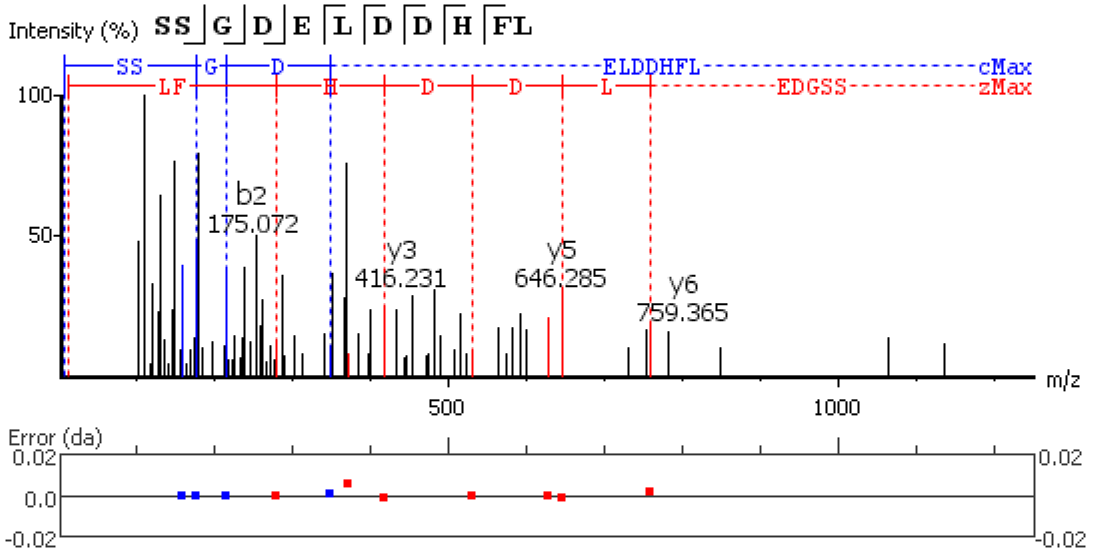


**Figure S5.138.** MS/MS spectrum of a Leucokinin neuropeptide with the sequence TRFSAWA(-.98) detected in the peptidergic signaling system of the American lobster *Homarus americanus* generated by PEAKS XPro software.

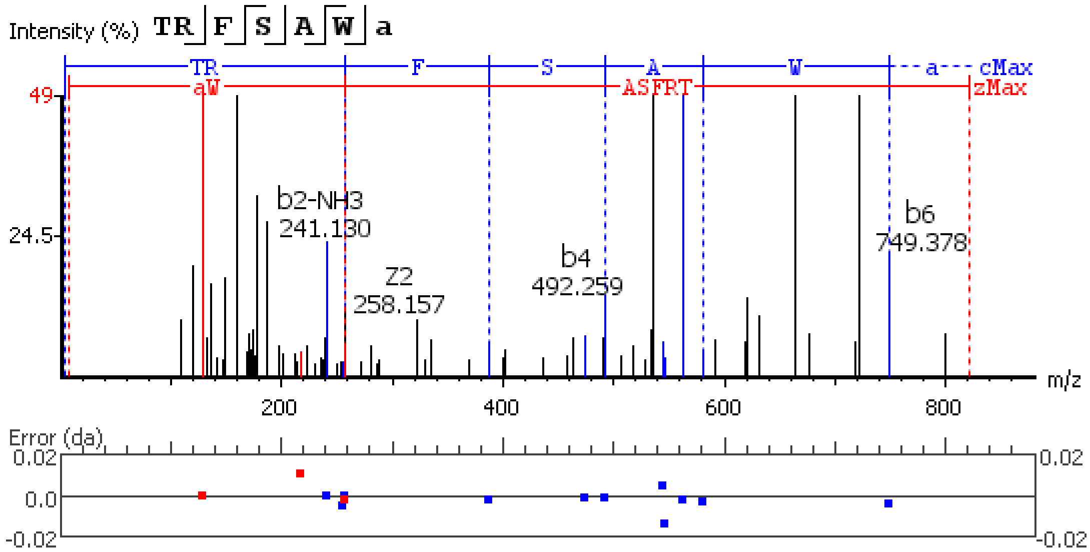


**Figure S5.139.** MS/MS spectrum of a Leucokinin neuropeptide with the sequence RTFSAWA(-.98) detected in the peptidergic signaling system of the American lobster *Homarus american*us generated by PEAKS XPro software.

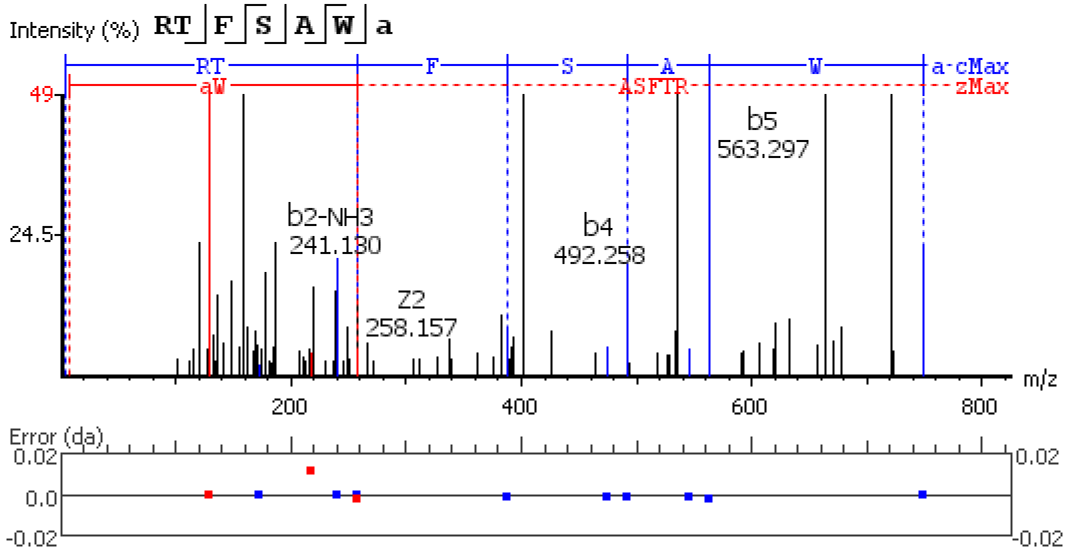


**Figure S5.140.** MS/MS spectrum of a Leucokinin neuropeptide with the sequence TRFSPWA(-.98) detected in the peptidergic signaling system of the American lobster *Homarus american*us generated by PEAKS XPro software.

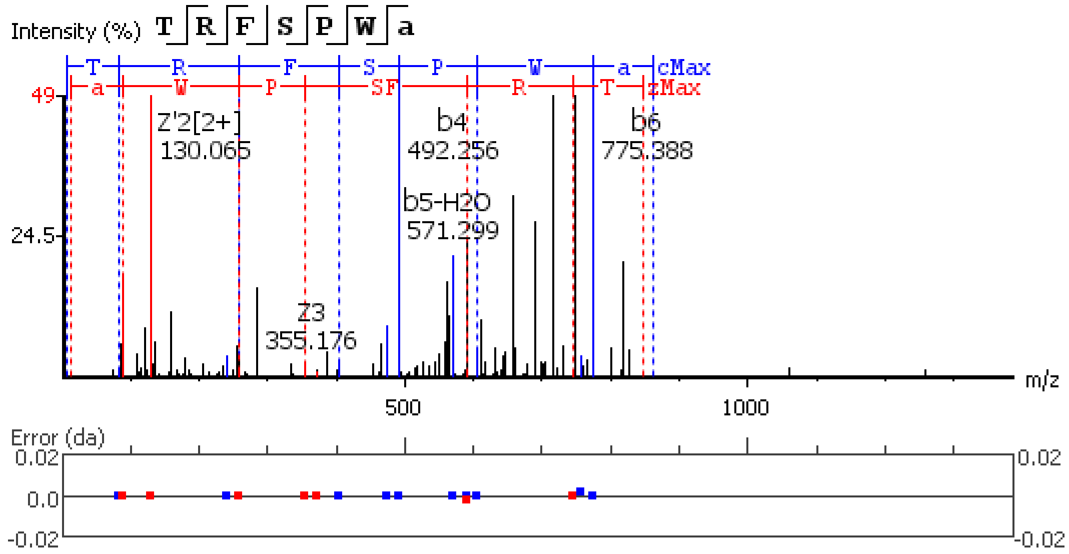


**Figure S5.141.** MS/MS spectrum of a Leucokinin neuropeptide with the sequence QAFHPWG(-.98) detected in the peptidergic signaling system of the American lobster *Homarus american*us generated by PEAKS XPro software.

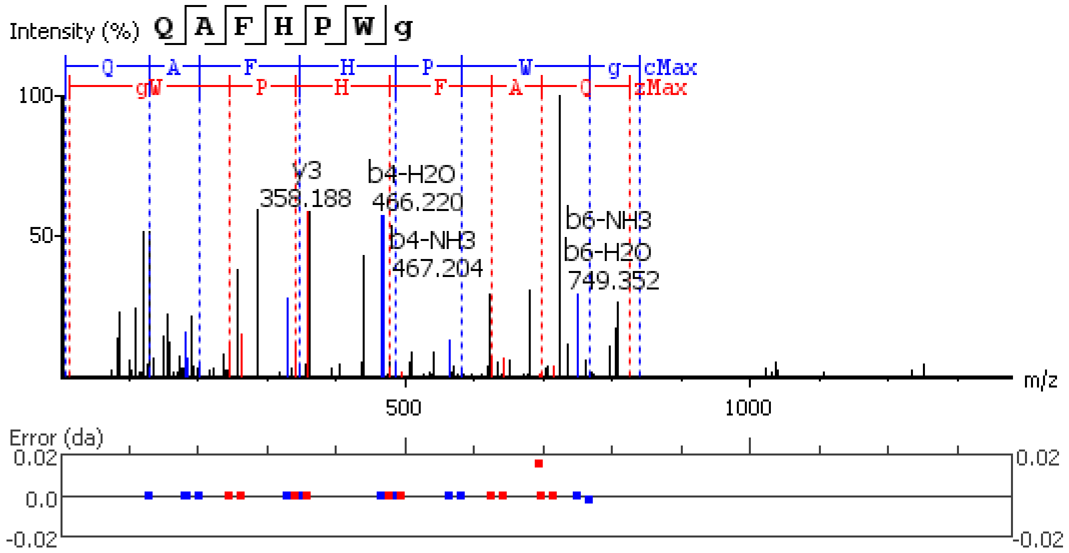


**Figure S5.142.** MS/MS spectrum of a Leucokinin neuropeptide with the sequence SSFKTAPGLPLSL detected in the peptidergic signaling system of the American lobster *Homarus american*us generated by PEAKS XPro software.

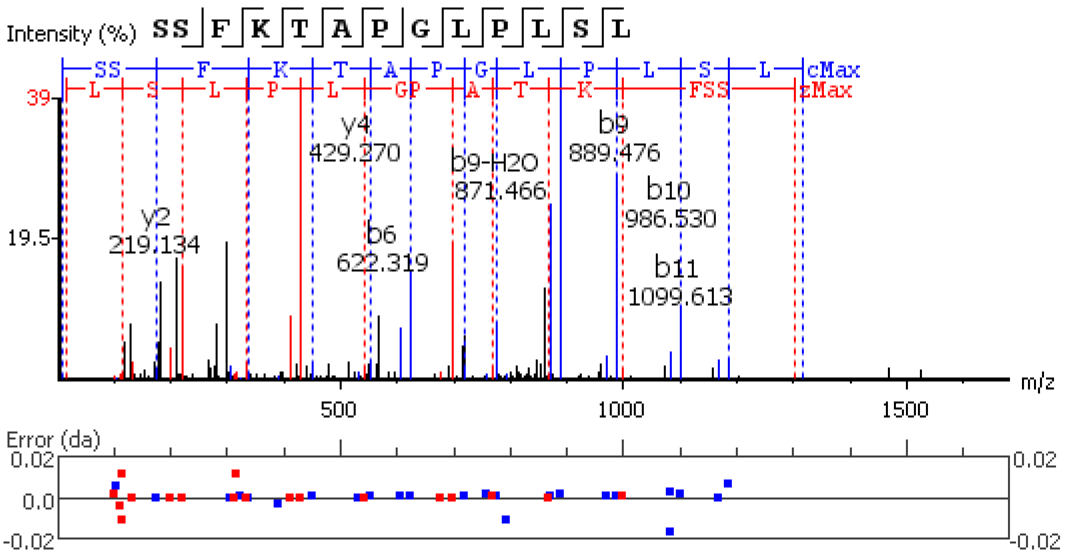


**Figure S5.143.** MS/MS spectrum of a Leucokinin neuropeptide with the sequence SDIDEKRPSFNAWA(-.98) detected in the peptidergic signaling system of the American lobster *Homarus american*us generated by PEAKS XPro software using data dependent acquisition (DDA)

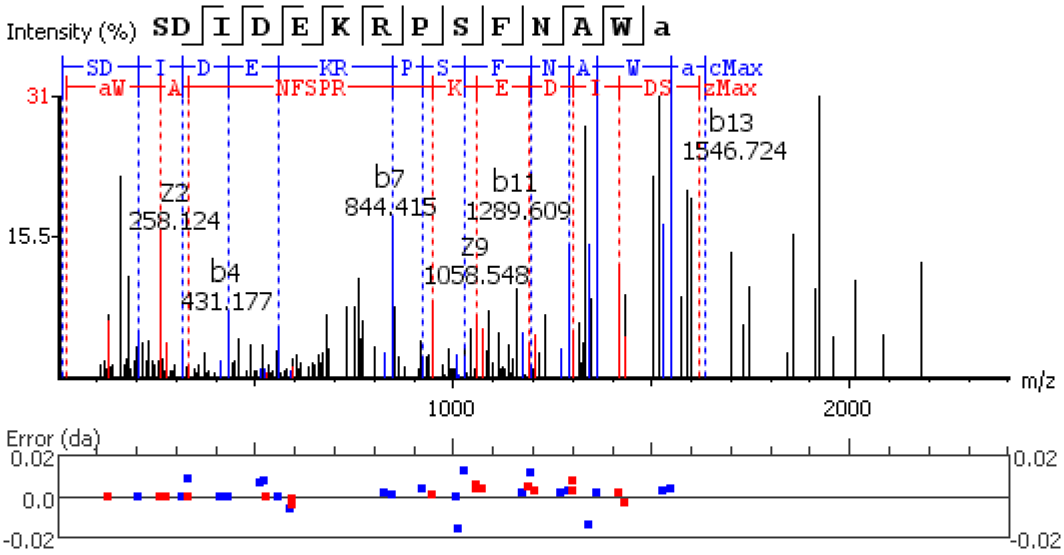


method

**Figure S5.144.** MS/MS spectrum of a Leucokinin neuropeptide with the sequence SDSDEKRPSFSAWA(-.98) detected in the peptidergic signaling system of the American lobster *Homarus american*us generated by PEAKS XPro software.

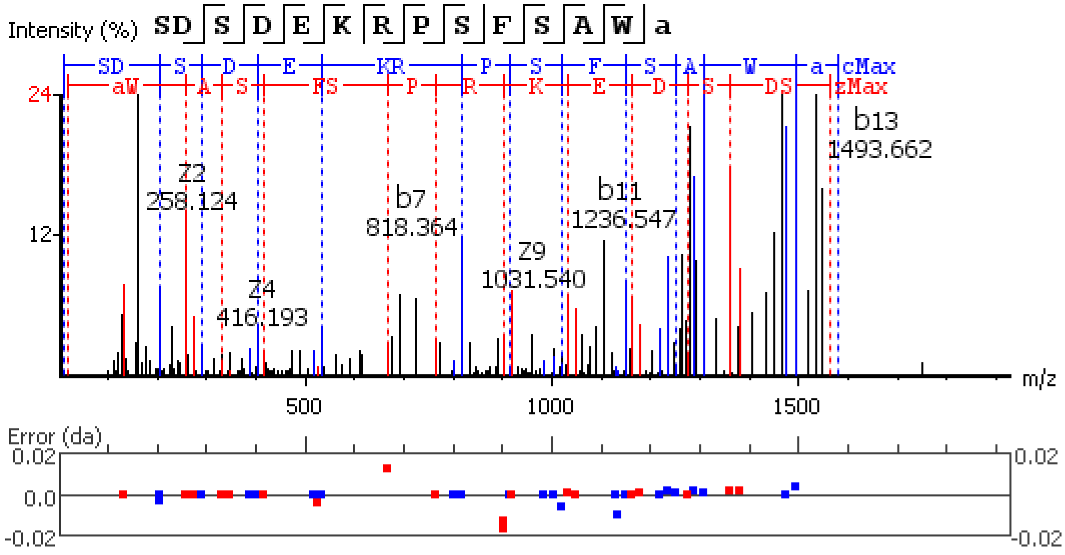

Intensity (%)
SD S D E K R P S F S A W a
SD S D E KR P S F S A W a cMax
aW A S FS P R K E D S DS zMax
24
12
Z2
258.124
Z4
416.193
b7
818.364
Z9
1031.540
b11
1236.547
b13
1493.662
m/z
500
1000
1500
Error (da)
0.02
0.0
-0.02
0.02
-0.02

**Figure S6.1.-24.** MS/MS spectrum of glycosylated neuropeptide sequences detected in the peptidergic signaling system of the American lobster *Homarus americanus* generated by PEAKSGlycanFinder v.2.5. software.

**Figure S6.1.** MS/MS spectrum of a AST-B glycosylated neuropeptide with the sequence <u>**S**</u>(+365.13)SSSPQQDDPASSPSHIEE detected in the peptidergic signaling system of the American lobster *Homarus americanus* generated by PEAKSGlycanFinder v.2.5. software.

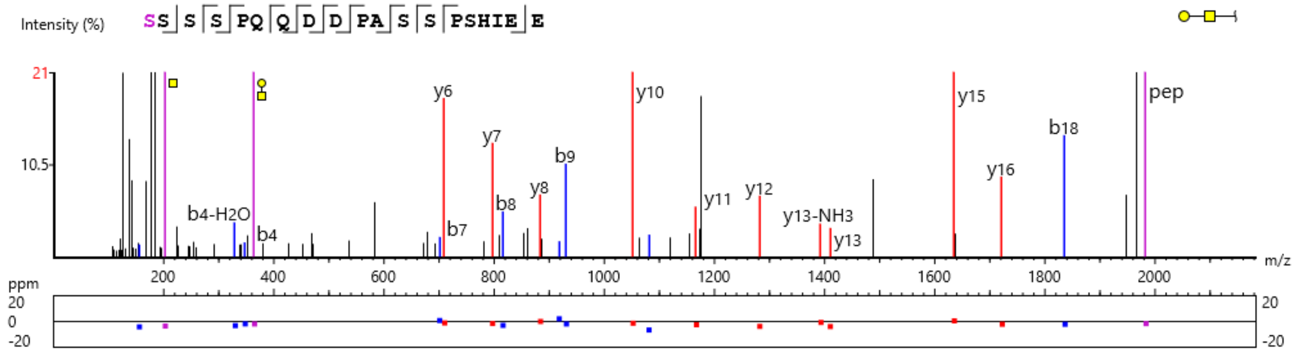


**Figure S6.2.** MS/MS spectrum of a CHH-B glycosylated neuropeptide with the sequence L<u>**S**</u>(+203.08)SISPSSTPLGFLSQDHSVN detected in the peptidergic signaling system of the American lobster *Homarus americanus* generated by PEAKSGlycanFinder v.2.5. The *m/z* 204.09 ion corresponds to a HexNAc residue. Although the software by default assigns this as GlcNAc (blue square), the glycan annotation was manually revised to GalNAc (yellow square) based on the observed GlcNAc/GalNAc oxonium ion intensity ratio.

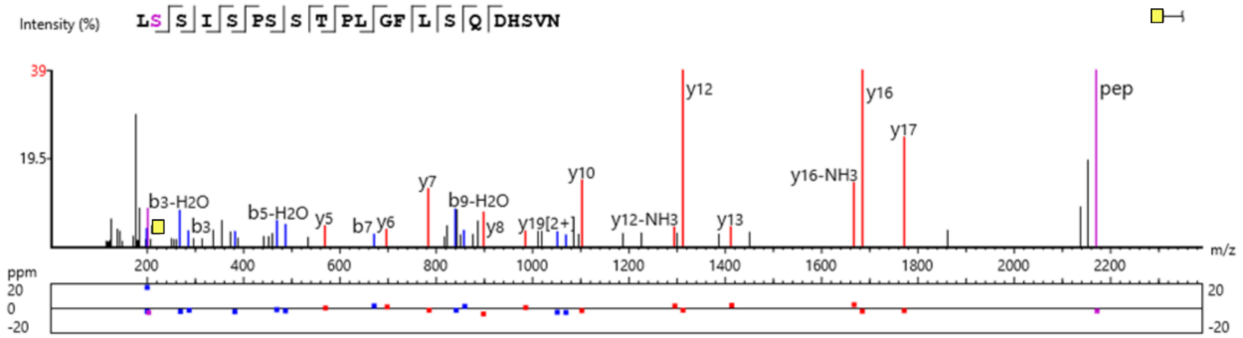


**Figure S6.3.** MS/MS spectrum of a CHH-B glycosylated neuropeptide with the sequence LSSISP<u>**S**</u>(+365.13)STPLGFLSQDHSV detected in the peptidergic signaling system of the American lobster *Homarus americanus* generated by PEAKSGlycanFinder v.2.5.

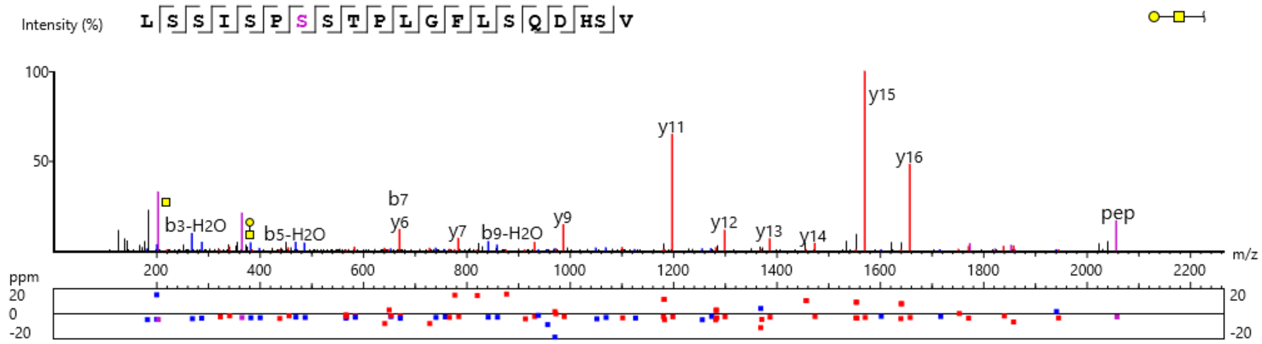

**Figure S6.4.** MS/MS spectrum of a CHH-B glycosylated neuropeptide with the sequence LLSSISP**S**(+365.13)STPLGFLSQDHSV detected in the peptidergic signaling system of the American lobster *Homarus americanus* generated by PEAKSGlycanFinder v.2.5.

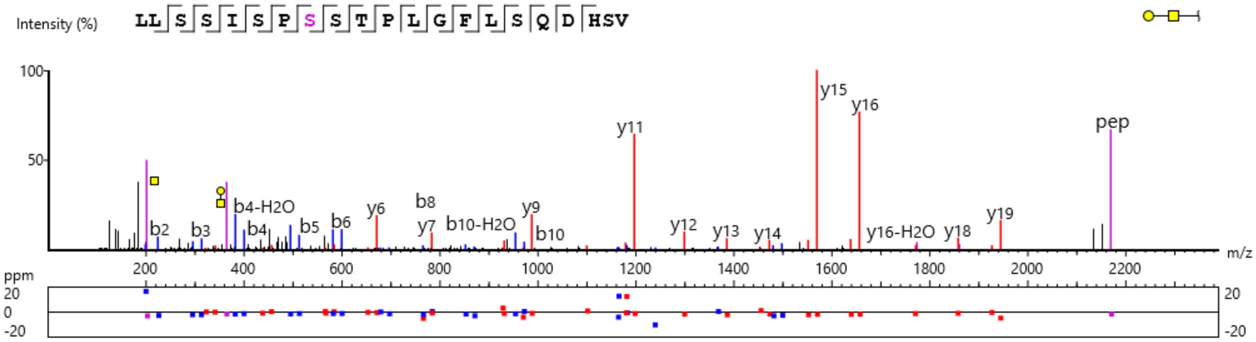


**Figure S6.5.** MS/MS spectrum of a CHH-B glycosylated neuropeptide with the sequence LLSSISP**S**(+203.08)STPLGFLSQDHSV detected in the peptidergic signaling system of the American lobster *Homarus americanus* generated by PEAKSGlycanFinder v.2.5. The *m/z* 204.09 ion corresponds to a HexNAc residue. Although the software by default assigns this as GlcNAc (blue square), the glycan annotation was manually revised to GalNAc (yellow square) based on the observed GlcNAc/GalNAc oxonium ion intensity ratio.

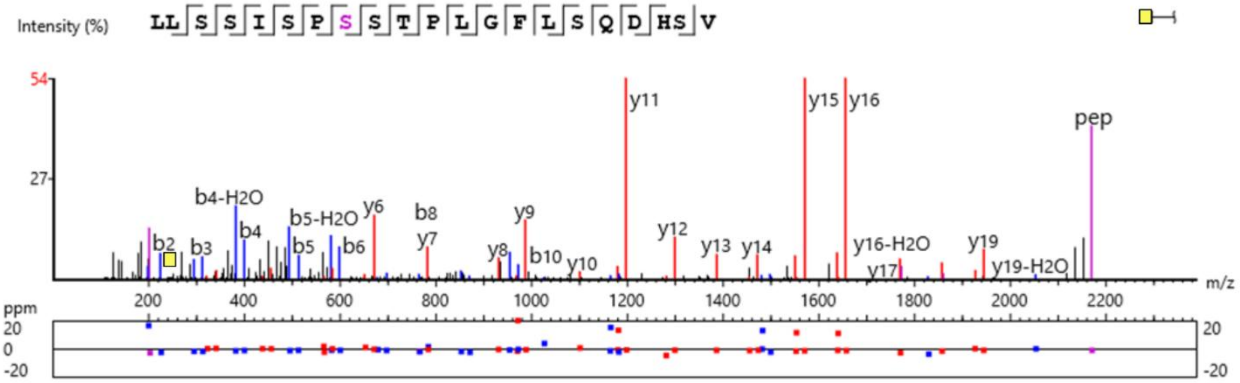


**Figure S6.6.** MS/MS spectrum of a CHH-B glycosylated neuropeptide with the sequence LSSISP**S**(+365.13)STPLGFLSQDHS detected in the peptidergic signaling system of the American lobster *Homarus americanus* generated by PEAKSGlycanFinder v.2.5.

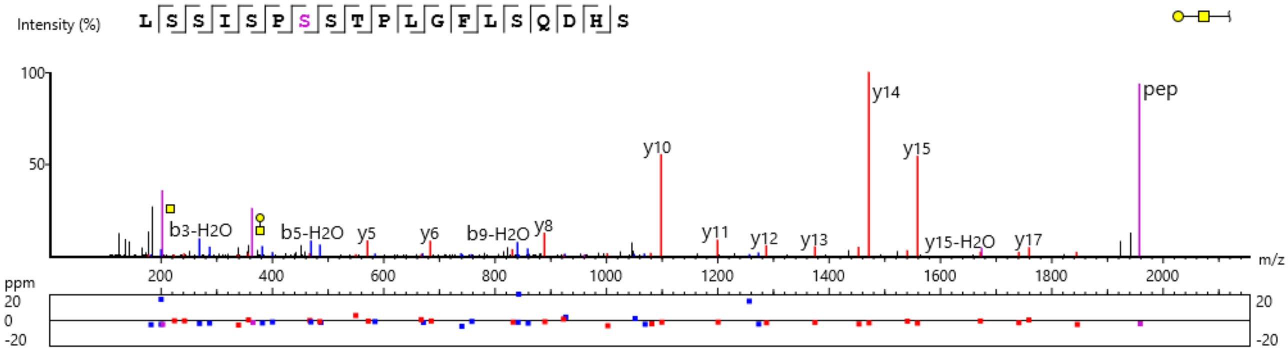

**Figure S6.7.** MS/MS spectrum of a CHH-B glycosylated neuropeptide with the sequence SSISP<u>**S**</u>(+365.13)STPLGFLSQDHSV detected in the peptidergic signaling system of the American lobster *Homarus american*us generated by PEAKSGlycanFinder v.2.5.

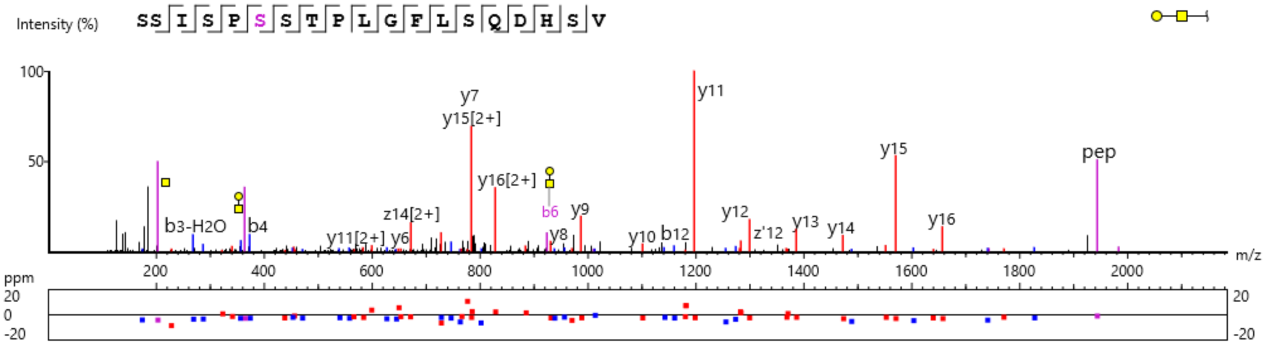


**Figure S6.8.** MS/MS spectrum of a CHH-B glycosylated neuropeptide with the sequence SSISP<u>**S**</u>(+365.13)STPLGFLSQDHSVN detected in the peptidergic signaling system of the American lobster *Homarus american*us generated by PEAKSGlycanFinder v.2.5.

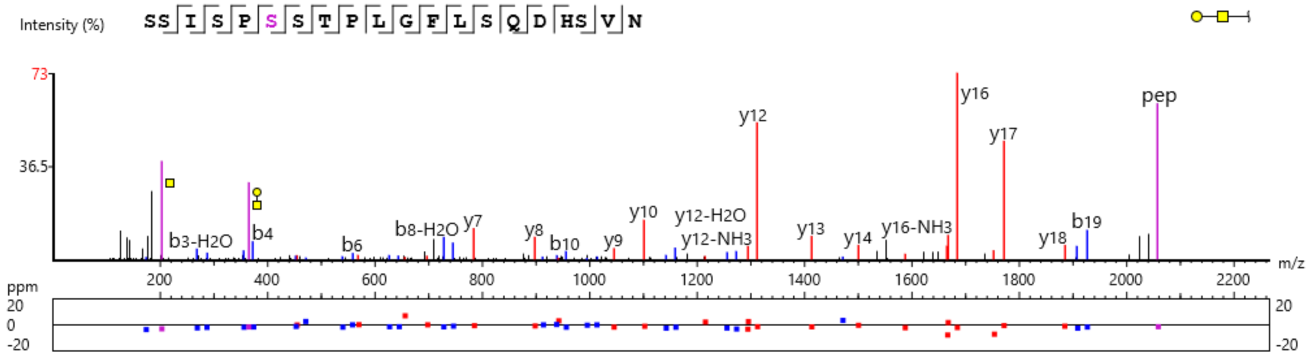


**Figure S6.9.** MS/MS spectrum of a CHH-B glycosylated neuropeptide with the sequence SSISP<u>**S**</u>(+365.13)STPLGFLSQDHS detected in the peptidergic signaling system of the American lobster *Homarus american*us generated by PEAKSGlycanFinder v.2.5.

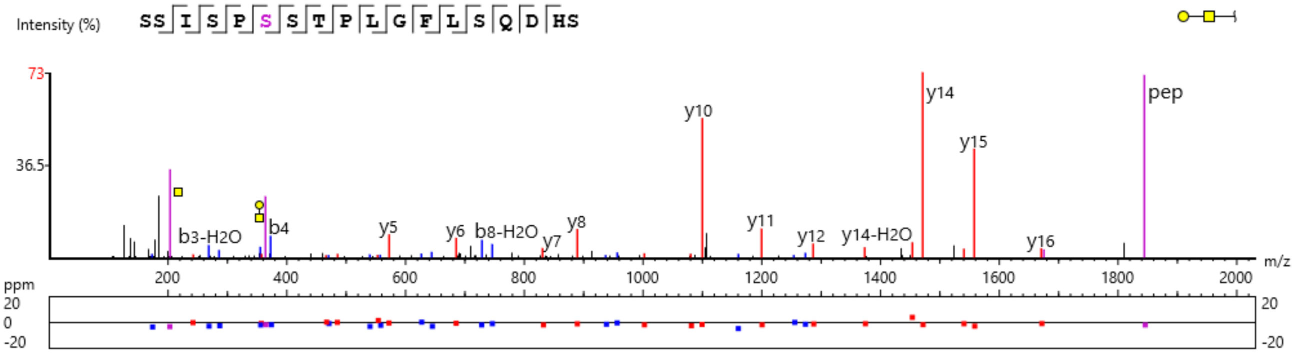


**Figure S6.10.** MS/MS spectrum of a CHH-B glycosylated neuropeptide with the sequence LSSISP<u>**S**</u>(+365.13)STPLGFLSQDHSVN detected in the peptidergic signaling system of the American lobster *Homarus american*us generated by PEAKSGlycanFinder v.2.5.

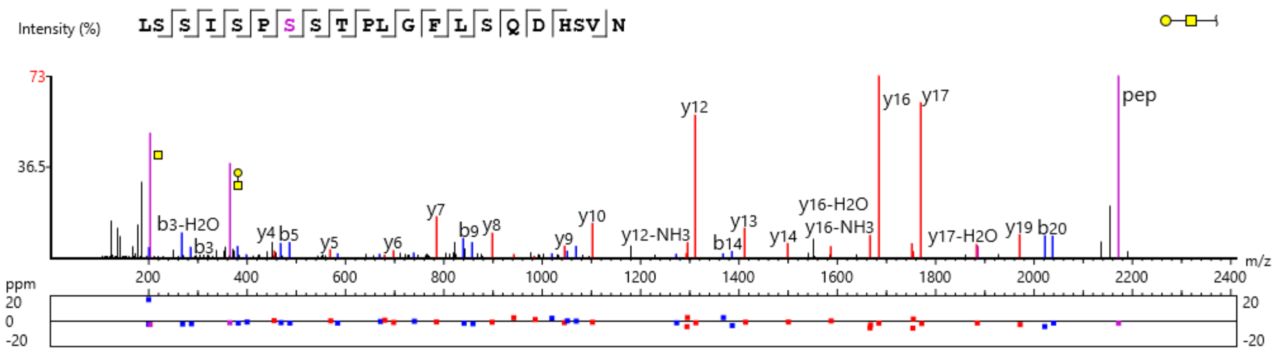


**Figure S6.11.** MS/MS spectrum of a CHH-B glycosylated neuropeptide with the sequence LSSISPS(+203.08)STPLGFLSQDHSVN detected in the peptidergic signaling system of the American lobster *Homarus american*us generated by PEAKSGlycanFinder v.2.5. The *m/z* 204.09 ion corresponds to a HexNAc residue. Although the software by default assigns this as GlcNAc (blue square), the glycan annotation was manually revised to GalNAc (yellow square) based on the observed GlcNAc/GalNAc oxonium ion intensity ratio.

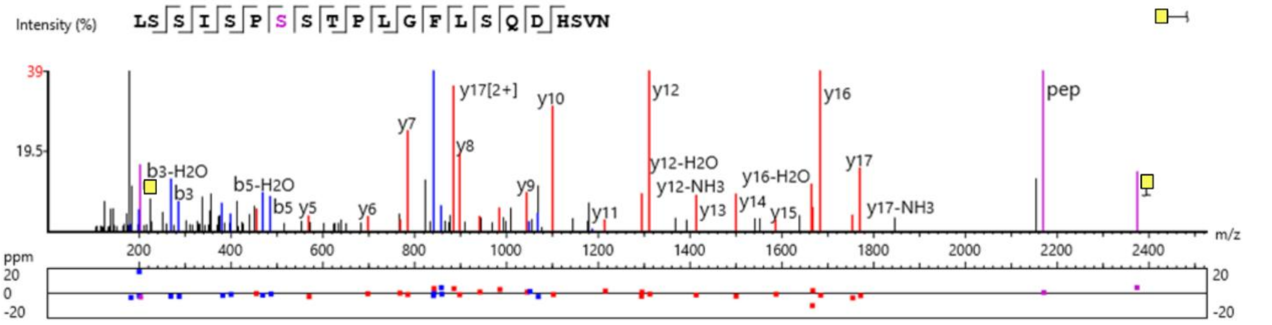


**Figure S6.12.** MS/MS spectrum of a CHH-B glycosylated neuropeptide with the sequence LSSISPS(+203.08)STPLGFLSQDHS detected in the peptidergic signaling system of the American lobster *Homarus american*us generated by PEAKSGlycanFinder v.2.5. The *m/z* 204.09 ion corresponds to a HexNAc residue. Although the software by default assigns this as GlcNAc (blue square), the glycan annotation was manually revised to GalNAc (yellow square) based on the observed GlcNAc/GalNAc oxonium ion intensity ratio.

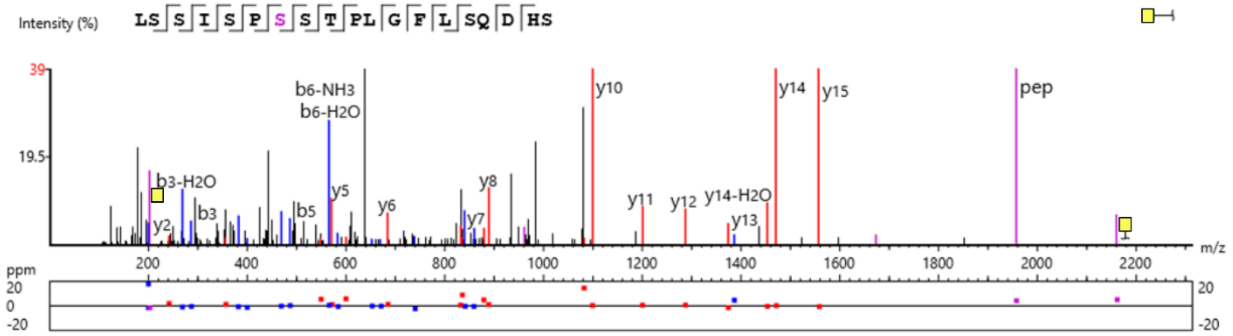


**Figure S6.13.** MS/MS spectrum of a CHH-B glycosylated neuropeptide with the sequence LSSISPS(+203.08)STPLGFLSQDHSV detected in the peptidergic signaling system of the American lobster *Homarus american*us generated by PEAKSGlycanFinder v.2.5. The *m/z* 204.09 ion corresponds to a HexNAc residue. Although the software by default assigns this as GlcNAc (blue square), the glycan annotation was manually revised to GalNAc (yellow square) based on the observed GlcNAc/GalNAc oxonium ion intensity ratio.

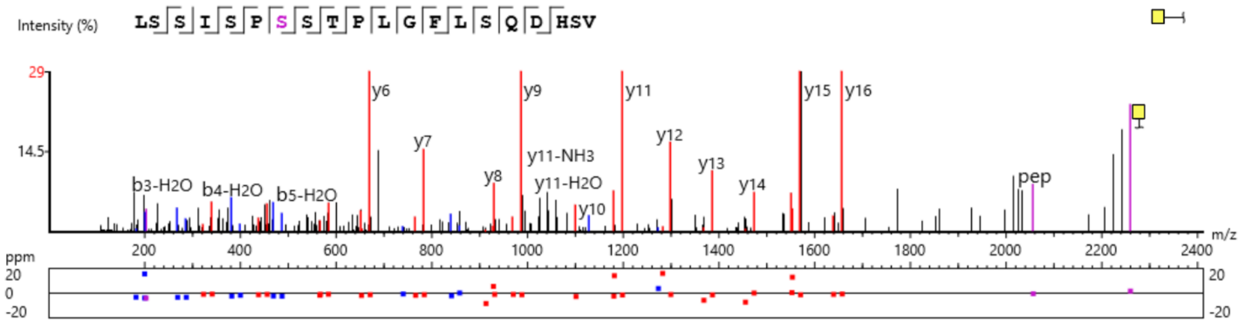


**Figure S6.14.** MS/MS spectrum of a CHH-B glycosylated neuropeptide with the sequence SSISP**S**(+203.08)STPLGFLSQDHSV detected in the peptidergic signaling system of the American lobster *Homarus americanus* generated by PEAKSGlycanFinder v.2.5. The *m/z* 204.09 ion corresponds to a HexNAc residue. Although the software by default assigns this as GlcNAc (blue square), the glycan annotation was manually revised to GalNAc (yellow square) based on the observed GlcNAc/GalNAc oxonium ion intensity ratio.

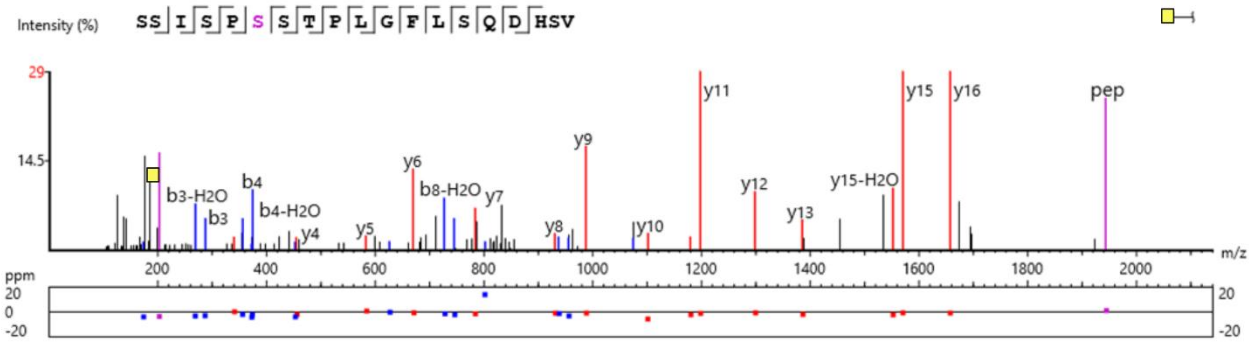


**Figure S6.15.** MS/MS spectrum of a CHH-B glycosylated neuropeptide with the sequence LSSISP**S**(+203.08)STPLGFLSQDHSV(-0.98) detected in the peptidergic signaling system of the American lobster *Homarus americanus* generated by PEAKSGlycanFinder v.2.5. The *m/z* 204.09 ion corresponds to a HexNAc residue. Although the software by default assigns this as GlcNAc (blue square), the glycan annotation was manually revised to GalNAc (yellow square) based on the observed GlcNAc/GalNAc oxonium ion intensity ratio.

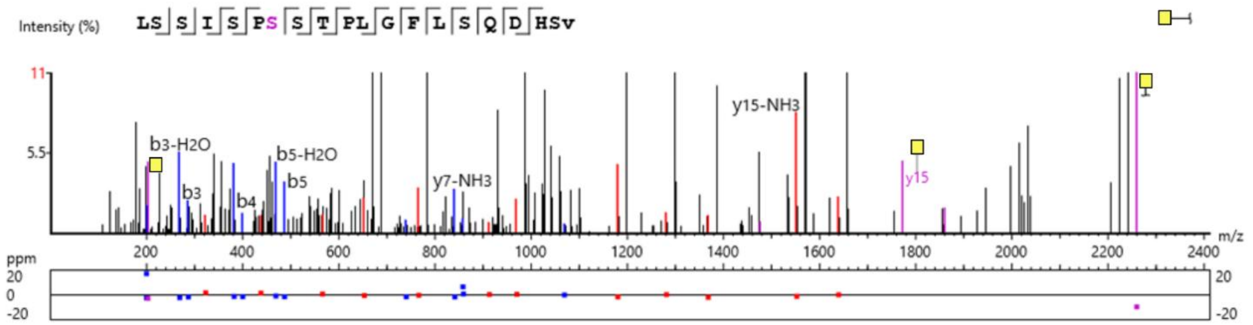


**Figure S6.16.** MS/MS spectrum of a CHH-B glycosylated neuropeptide with the sequence LSSISP**S**(+552.22)STPLGFLSQDH(-0.98) detected in the peptidergic signaling system of the American lobster *Homarus american*us generated by PEAKSGlycanFinder v.2.5. The glycan structure annotation was revised to an undetermined color to reflect the ambiguous identity of the distal HexNAc residue.

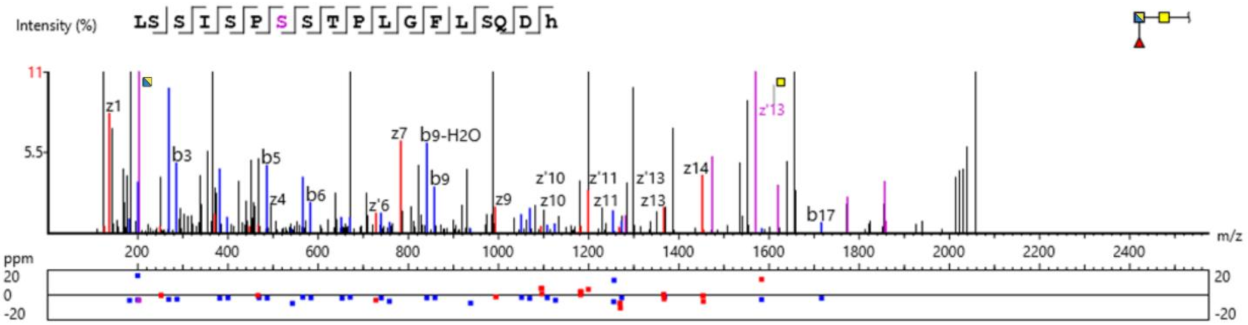


**Figure S6.17.** MS/MS spectrum of a CHH-B glycosylated neuropeptide with the sequence LSSISP**S**(+365.13)STPLGFLSQDHSV(-0.98) detected in the peptidergic signaling system of the American lobster *Homarus american*us generated by PEAKSGlycanFinder v.2.5.

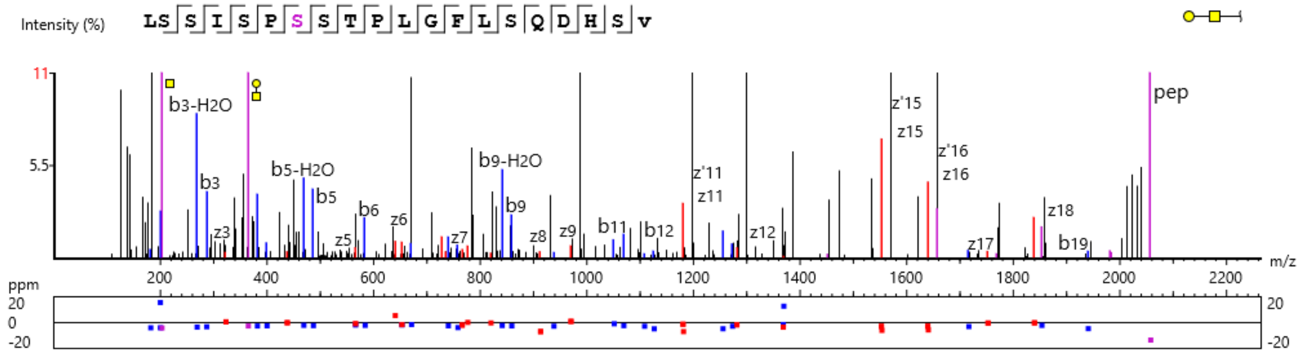


**Figure S6.18.** MS/MS spectrum of a CHH-B glycosylated neuropeptide with the sequence LLSSISP**S**(+552.22)STPLGFLSQDH(-0.98) detected in the peptidergic signaling system of the American lobster *Homarus american*us generated by PEAKSGlycanFinder v.2.5. The glycan structure annotation was revised to an undetermined color to reflect the ambiguous identity of the distal HexNAc residue.

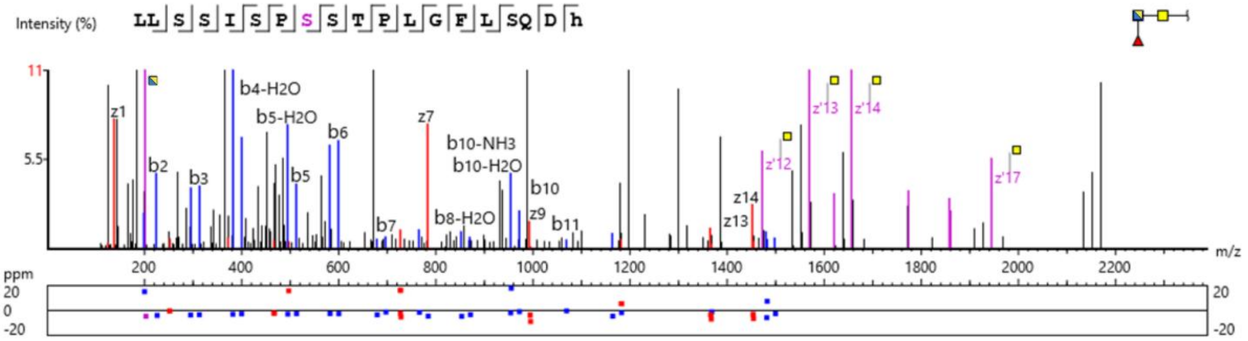


**Figure S6.19.** MS/MS spectrum of a CHH-B glycosylated neuropeptide with the sequence LL**S**(+365.13)SISPSSTPLGFLSQDHSV detected in the peptidergic signaling system of the American lobster *Homarus american*us generated by PEAKSGlycanFinder v.2.5.

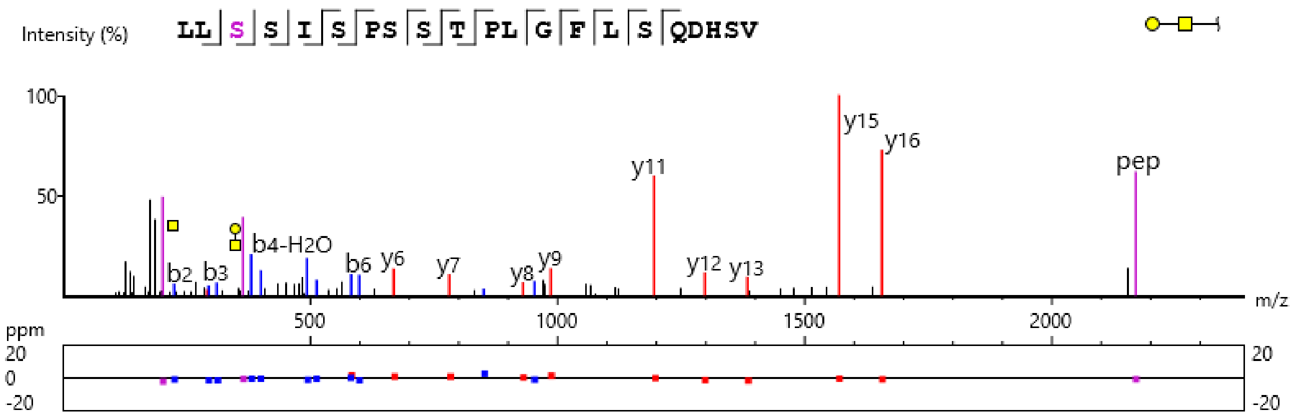


**Figure S6.20.** MS/MS spectrum of a CHH-B glycosylated neuropeptide with the sequence RSVEGVSRMEKLLSSISPSSTPLGFL**S**(+365.13)QDHSV(-0.98) detected in the peptidergic signaling system of the American lobster *Homarus americanus* generated by PEAKSGlycanFinder v.2.5.

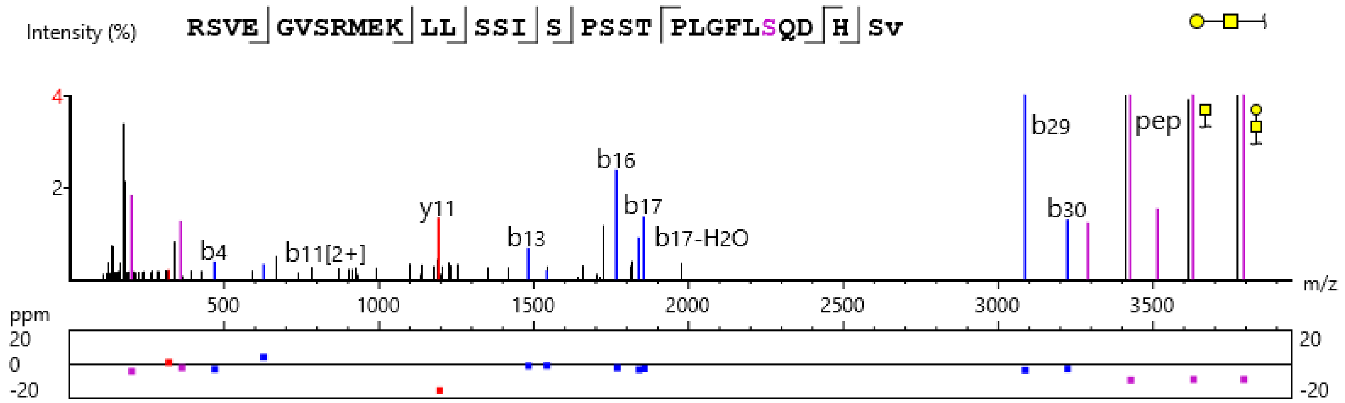


**Figure S6.21.** MS/MS spectrum of a CHH-B glycosylated neuropeptide with the sequence L**S**(+365.13)SISPSSTPLGFLSQDHSV detected in the peptidergic signaling system of the American lobster *Homarus americanus* generated by PEAKSGlycanFinder v.2.5.

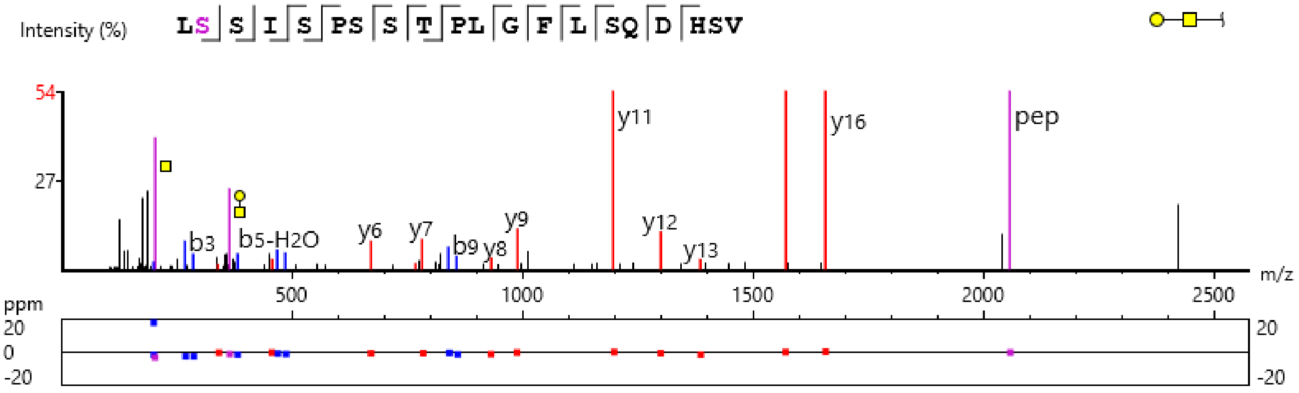


**Figure S6.22.** MS/MS spectrum of a Orcokinin glycosylated neuropeptide with the sequence GPIKAAPARSSPQQDAAAGY**T**(+365.13)DGAPV detected in the peptidergic signaling system of the American lobster *Homarus americanus* generated by PEAKSGlycanFinder v.2.5.

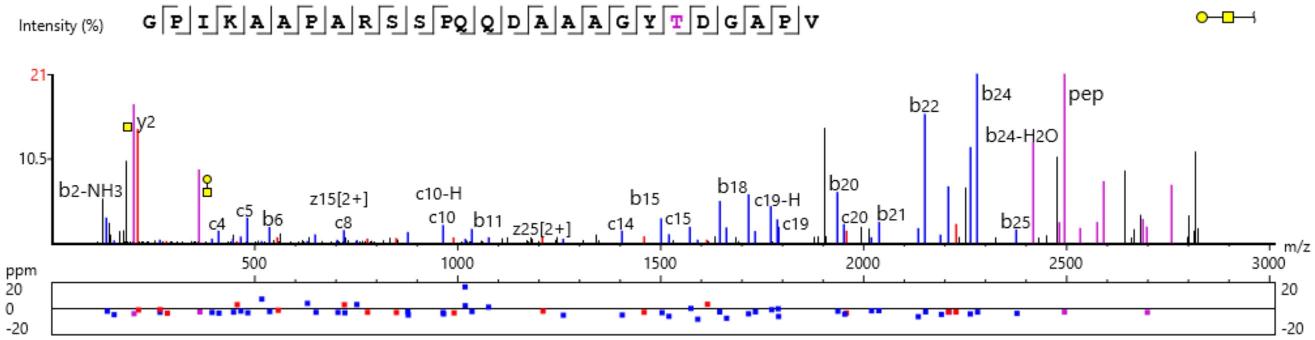


**Figure S6.23.** MS/MS spectrum of a Orcokinin glycosylated neuropeptide with the sequence AAPARS(+568.21)SPQQDAAAGY(-0.98) detected in the peptidergic signaling system of the American lobster *Homarus american*us generated by PEAKSGlycanFinder v.2.5. Although the software initially assigned the glycan as $GlcNac_2Man_1$-O-Ser, the glycan annotation was manually revised to Core 2-type O-glycan structure based on the observed GlcNAc/GalNAc oxonium ion intensity ratio and the known biological prevalence of Core 2 O-glycans in mucin-type glycosylation.

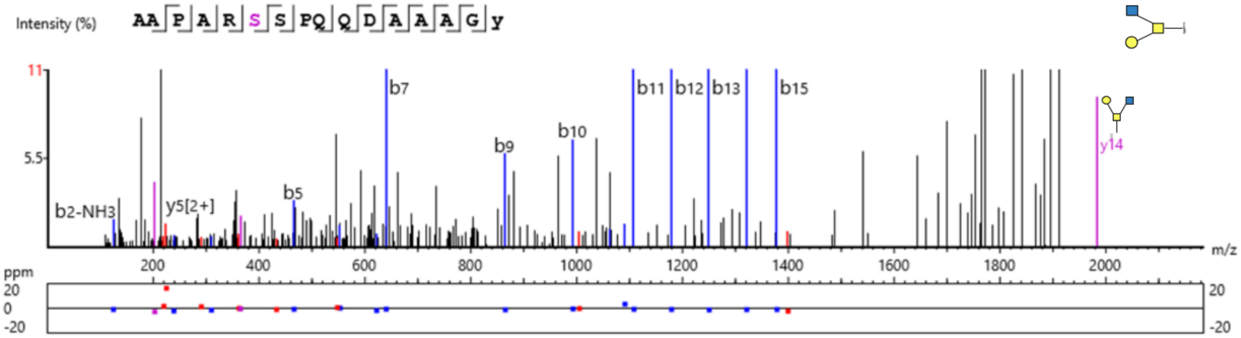


**Figure S6.24.** MS/MS spectrum of a SIFamide glycosylated neuropeptide with the sequence VYRKPPFNGS(+1022.38)IF(-0.98) detected in the peptidergic signaling system of the American lobster *Homarus americanus* generated by PEAKSGlycanFinder v.2.5. While the software initially assigned the glycosylation site to S10, manual inspection and annotation (as shown in **Figure 5** of the main article) support that the correct glycosylation site is N8.

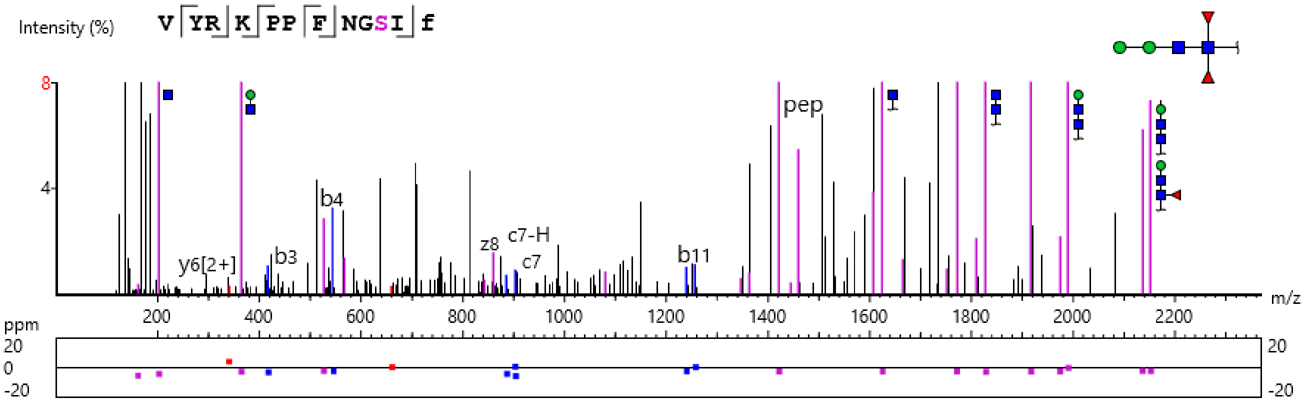
